# Steroid‑based Tide Quencher 1 probes enable real‑time mapping of novel non‑canonical cholesterol sites on the M1 muscarinic receptor

**DOI:** 10.64898/2026.03.26.714567

**Authors:** Nikolai Chetverikov, Eszter Szánti-Pintér, Jan Jurica, Mariia Vodolazhenko, Miloš Buděšínský, Václav Zima, Martin Svoboda, Eva Dolejší, Alena Janoušková-Randáková, Anna Urbánková, Jan Jakubík, Eva Kudová

**Affiliations:** Institute of Physiology, Academy of Sciences of the Czech Republic, Prague, Czech Republic; Institute of Organic Chemistry and Biochemistry, Academy of Sciences of the Czech Republic, Prague, Czech Republic

## Abstract

Steroid-based fluorescent-quencher probes now enable real-time, residue-level mapping of previously inaccessible cholesterol-binding sites on G-protein-coupled receptors. We designed Tide Quencher 1 (TQ1) conjugated steroids that target two distinct peripheral sites on the M_1_ muscarinic receptor. One near the extracellular N-terminus and another adjacent to the intracellular C-terminus. Using pregnanolone glutamate as a versatile scaffold, we synthesised a library of probes varying in C-3 linker length (γ-aminobutyric acid vs. *L*-glutamic acid) and C-3/C-5 stereochemistry (3α/3β/5α/5β). Fluorescence-quenching assays with CFP-tagged receptors revealed that TQ1 probes consistently outperformed Dabcyl, delivering up to 40 % quenching within minutes and sub-micromolar EC_50_ values. The most potent N-terminal probe (3α5α-PRG-Glu-TQ1 (5)) achieved 300 nM potency, while the best C-terminal probe (3α5β-PRG-Glu-TQ1 (3)) reached 1 µM potency with rapid association. Molecular docking and MD simulations identified key residues (K20, Q24, W405 at the N-site; K57, Y62, W150 at the C-site) mediating binding, a prediction confirmed by alanine-scan mutagenesis that markedly reduced quenching at the N-terminus and only modestly affected the C-terminus. Competition experiments with non-quenching analogues further validated probe specificity. Crucially, the pregnane core proved essential; alternative steroid backbones failed to generate robust quenching. This fluorescence-quenching platform overcomes the limitations of traditional radioligand assays, providing kinetic insight, high-throughput compatibility, and the ability to dissect lipid-GPCR interactions in native membranes. The approach is readily extensible to other GPCR families, opening new avenues for structure-guided drug discovery targeting allosteric cholesterol sites.

## Introduction

G protein-coupled receptors (GPCRs) are one of the most extensively studied and targeted families in pharmacotherapy, primarily due to their significant role in human pathophysiology and their pharmacological tractability. GPCRs are the target for approximately 30-36% of all approved drugs, with about 516 drugs acting on 121 unique GPCR targets (Lorente et al., 2025). Cholesterol has been found co-crystallised with several GPCRs (Gimpl, 2016). Today, the number of structures has already surpassed 200 (rcbs.org). Membrane cholesterol directly allosterically modulates ligand binding to and activation of GPCRs and affects the pharmacology of GPCRs (Jakubík & El-Fakahany, 2021).

Two binding motifs for membrane cholesterol were postulated. First, the so-called ‘cholesterol recognition amino acid consensus (CRAC) domain’ common for all membrane proteins that contain a (–L/V-(X)1–5-Y-(X)1–5-R/K-) pattern was described(Li & Papadopoulos, 1998). Subsequently, a (K/R)-X1−5-(Y/F)-X1−5-(L/V) reversed pattern (CARC) was found in the nicotinic acetylcholine receptor(Baier et al., 2011). The so-called ‘cholesterol consensus motif’ (CCM) was identified in the structure of the β_2_-adrenergic receptor (3D4S) and predicted for a plethora of other GPCRs (Hanson et al., 2008). Yet a current extensive study of cryo-EM and X-ray structures has shown that almost all cholesterol binding to GPCRs occurs in predictable locations that, however, lack perceivable cholesterol-binding motifs (Taghon et al., 2021). Cholesterol can enhance the conformational stability of GPCRs, either by promoting active or inactive states, which is crucial for their signalling functions. This stabilisation is often achieved through direct interactions with specific residues rather than relying solely on conserved motifs (Sarkar & Chattopadhyay, 2020). For example, membrane cholesterol prevents persistent activation of muscarinic receptors by wash-resistant xanomeline by interaction with R^6.35^ and L/I^6.46^(Ballesteros and Weinstein numbering (Ballesteros & Weinstein, 1995) in the TM6 (Randáková et al., 2018). The high plasticity of cholesterol interaction sites and the lack of universal motifs highlight the importance of detailed, residue-level analysis to understand how cholesterol influences GPCR function. For example, molecular dynamics simulations and structural bioinformatics elucidate cholesterol-GPCR interactions at a finer scale(Geiger et al., 2021; Sarkar & Chattopadhyay, 2020).

Neurosteroids and steroid hormones exhibit nanomolar-range allosteric modulation of muscarinic receptors via these cholesterol-binding sites, suggesting novel therapeutic opportunities (Dolejší et al., 2021). Understanding the location of the binding site and the structure-activity relationship is crucial for structure-based drug design. The affinity of conventional modulators of GPCRs can be easily and precisely determined in a cheap radioligand binding assay based on quantifying the amount of bound radioligand after separating it from the free radioligand. Unfortunately, radiolabelled steroids are not suitable for radioligand binding assays because steroids dissolve instantly in biological membranes, and a steroid bound to a GPCR cannot then be separated from its free form. To overcome this limitation, we developed a fluorescence-quenching assay addressing the technical limitations of traditional radioligand methods. Fluorescently tagged steroid derivatives with γ-aminobutyric acid /*L*-glutamate linkers conjugated with fluorescence quenchers enabled direct binding measurements to GPCRs fused with fluorescent proteins. This approach establishes a generalizable platform for studying lipid-GPCR interactions, overcoming historical challenges in separating membrane-embedded ligands. Future applications could expand to other receptor families and facilitate structure-based drug design targeting allosteric cholesterol sites.

## Results

### Influence of Steroid Scaffold Geometry on Muscarinic Receptor Recognition

Our previous comprehensive structure–activity relationship (SAR) investigation of endogenous steroid hormones and neurosteroids demonstrated that endogenous neurosteroid 3α-hydroxy-5β-pregnan-20-one (3α5β-pregnanolone, **3α5β-PRG 23**), in contrast to its C-3 and C-5 stereoisomers (**Figure 1B, C**), exclusively exhibited nanomolar affinities across M_1_, M_2_, M_4_, and M_5_ acetylcholine muscarinic receptors in the binding experiments measuring the allosteric modulation of [^3^H]N-methylscopolamine binding ([^3^H]NMS) (Dolejší, Szánti-Pintér, et al., 2021). In the initial phase of this study, we found that C-3 glutamate substitution was highly favourable, as the C-3 glutamate derivative of pregnanolone retained nanomolar binding affinity at the M_1_ receptor. Specifically, **3α5β-PRG (23)** and 3α,5β-pregnanolone glutamate (**3α5β-PRG-Glu 35**) displayed M_1_ receptor affinities of 180 nM and 150 nM, respectively (**Figure 1D**). Further, our data demonstrated that the steroid skeleton can be further modified by removal of the C-17 substituent, yielding highly active derivatives across muscarinic subtypes, particularly when combined with C-3 aspartate or glutamate substituents. Among these, 5β-androstan-3α-yl *L*-aspartate exhibited an affinity of 180 nM for M_1_ receptors, 50 nM at M_2_, and 400 nM at M_5_, while 5β-androstan-3α-yl *L*-glutamate maintained consistent nanomolar binding (150 nM) across all subtypes (Dolejší, Chetverikov, et al., 2021). These results indicate that polar amino-acid substituents at C-3 can compensate for modifications of the steroidal skeleton. Within this framework, pregnanolone glutamate (**Figure 1D**) was selected as the lead molecule for quencher dye tagging due to its combination of functional groups; the amino group, keto group, and carboxylic acid all provide suitable sites for further chemical modification. This chemical flexibility facilitates efficient conjugation to quencher dyes, enabling advanced biophysical assays.

**Figure 1.**
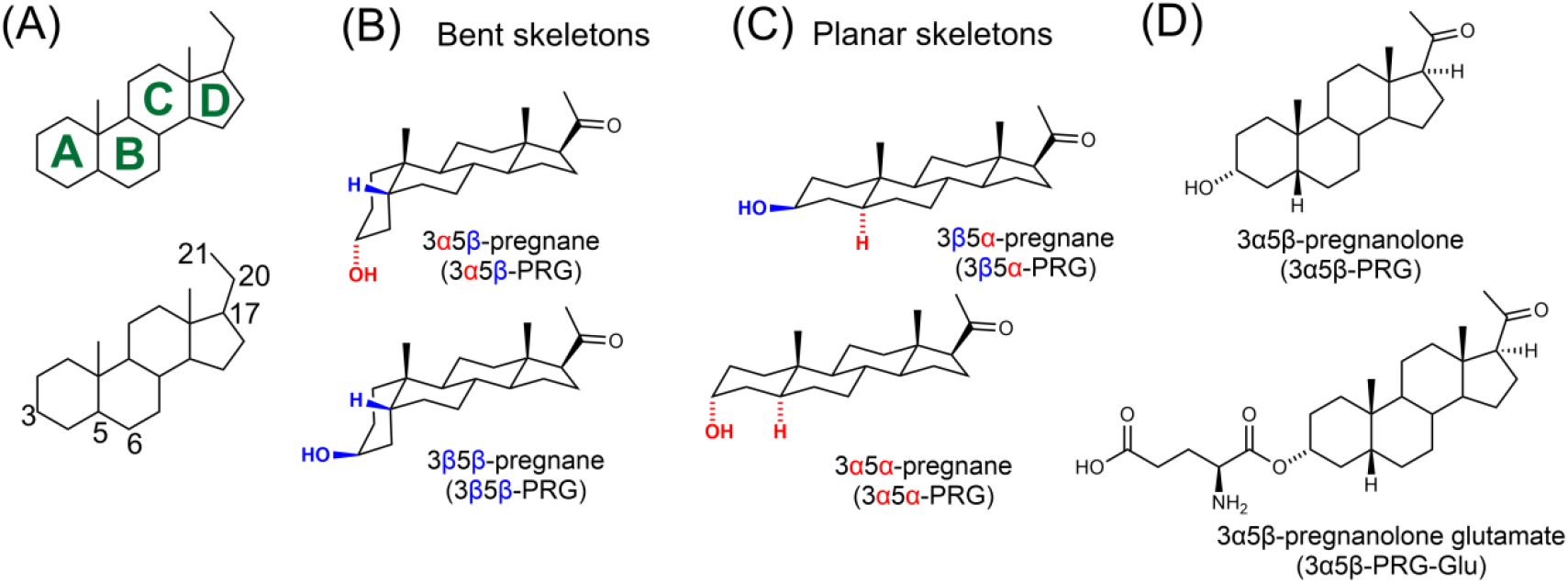
Stereochemical definitions, steroidal skeleton conformations, and representative 3α5β-pregnanolone-based ligands. *(A)* Numbering and lettering of the rings of steroid compounds *(B)* Representation of the bent 3α,5β- and 3β,5β-pregnane skeletons used in this study. *(C)* Representation of the planar 3β,5α- and 3α,5α-pregnane skeletons used in this study. *(D)* Structures of high-affinity compounds – 3α,5β-pregnanolone and 3α,5β-pregnanolone glutamate for M_1_ muscarinic receptor.

### Fluorescence Quenching Assay

To quantitatively assess ligand–receptor interactions, this study evaluated synthesised compounds for their affinity for the M_1_ muscarinic receptor using a fluorescence quenching assay. Recombinant M_1_ receptor constructs fused with CFP at either the N-or C-terminus were expressed in Sf9 cell membranes using a baculovirus system, ensuring high receptor expression with minimal interference from endogenous GPCRs. Membrane preparations containing receptor-CFP fusion proteins were incubated with either steroid-quencher probes or a control solution (KHB). Fluorescence quenching was monitored over time by measuring CFP fluorescence intensity at 485 nm, with excitation at 435 nm, providing a measure of ligand-binding dynamics.

#### Selection of Tide Quencher 1 (TQ1) as an optimal dark quencher

First, only the 5β-pregnanolone 3-yl glutamate esters were functionalized with Dabcyl and TQ1 probes to generate four conjugates: **3α5β-PRG-Glu-Dabcyl (1), 3β5β-PRG-Glu-Dabcyl (2), 3α5β-PRG-Glu-TQ1 (3), 3β5β-PRG-Glu-TQ1 (4)** (**Scheme 1A**). Dabcyl is a well-known, reliable dark quencher used in FRET assays. However, TQ1 offers superior absorption and quenching efficiency, improving signal discrimination and enabling more sensitive detection across various spectral conditions, enhancing assay performance and reliability. Synthesis (**Scheme 1A**) involved the conversion of the 3-hydroxysteroid into the Boc- and benzyl-protected *L*-glutamic acid esters, followed by palladium-catalysed hydrogenolysis of the benzyl group and Boc removal with trifluoroacetic acid. This afforded the free amino glutamate ester intermediates. The intermediates were then coupled with the succinimidyl esters of the quenchers (Dabcyl-OSu or TQ1-OSu) to form the final amide-linked conjugates.

Next, we tested the two Dabcyl probes (**3α5β-PRG-Glu-Dabcyl (1), 3β5β-PRG-Glu-Dabcyl (2)**) and the two TQ1 probes (**3α5β-PRG-Glu-TQ1 (3), 3β5β-PRG-Glu-TQ1 (4)**) in quenching assays (**Figure 2**). We first determined the time course and maximal quenching efficacy (E_max_) at a probe concentration of 1 µM. Potencies (half-maximal effective concentrations, EC_50_) were then derived from quenching responses over a probe concentration range of 100 nM to 1 μM (Supplementary Information Figure S1) (**Table 1**). Finally, binding affinity was assessed in radioligand displacement experiments using [^3^H]NMS. The binding affinity (K_A_) reflects the overall interaction at all binding sites, including both proximal to the N-and C-terminal proximal sites.

**Table 1.** Affinities (K_A_), potencies (EC_50_), and maximum quenching (E_MAX_) at 1 µM concentration. Affinities (K_A_) were obtained from competition with [^3^H]NMS. Potencies (EC_50_) and maximal quenching at 1 μM concentration of the probe were obtained from quenching experiments. ^[a]^E_MAX_ values are expressed in per cent of fluorescence quenched.

| Compound | N-terminal proximal site |  | C-terminal proximal site |  | Affinity |
| --- | --- | --- | --- | --- | --- |
| | $EC_{50}^a$ | $E_{MAX}$ @ 1 μM | $EC_{50}^a$ | $E_{MAX}$ @ 1 μM | $K_A^b$ |
| 3α5β-PRG-Glu-Dabcyl (1) | > 3 μM | 15% | cnbd | 0% | 10 μM |
| 3β5β-PRG-Glu-Dabcyl (2) | ≈ 3 μM | 33% | > 3 μM | 18% | 1 μM |
| 3α5β-PRG-Glu-TQ1 (3) | 3 μM | 15% | 1 μM | 35% | 3 μM |
| 3β5β-PRG-Glu-TQ1 (4) | 1 μM | 40% | 1 μM | 28% | 1 μM |
| 3α5α-PRG-Glu-TQ1 (5) | 300 nM | 23% | 3 μM | 17% | cnbd |
| 3β5α-PRG-Glu-TQ1 (6) | 1 μM | 41% | 1 μM | 30% | 1 μM |
| 3α5β-PRG-GABA-TQ1 (7) | 500 nM | 40% | > 3 μM | 20% | cnbd |
| 3β5β-PRG-GABA-TQ1 (8) | > 3 μM | 30% | > 3 μM | 10% | 2 μM |
| 3α5α-PRG-GABA-TQ1 (9) | 300 nM | 37% | 750 nM | 21% | 1 μM |
| 3β5α-PRG-GABA-TQ1 (10) | 500 nM | 12% | cnbd | 8% | 1 μM |
| 3α5β-PRG-TQ1 (11) | cnbd | 0% | cnbd | 0% | 1 μM |
| 3β5β-PRG-TQ1 (12) | cnbd | 0% | cnbd | 0% | 2 μM |
| 3α5α-PRG-TQ1 (13) | 1 μM | 30% | 3 μM | 18% | 1 μM |
| 3β5α-PRG-TQ1 (14) | cnbd | 0% | cnbd | 0% | cnbd |
| Pregnenolone-GABA-TQ1 (15) | cnbd | 8% | cnbd | 5% | 3 μM |
| DHEA-GABA-TQ1 (16) | cnbd | 5%<sup>c</sup> | cnbd | 10%<sup>c</sup> | 500 nM |
| 17β-Me-5β-AND-3β-GABA-TQ1 (17) | cnbd | 0% | cnbd | 0% | 10 μM |

[a] Determined from quenching experiments. [b] Determined from radioligand binding experiments. [c] Transient decrease. [d] cnbd, cannot be determined either because of lack of quenching ( $EC_{50}$ ) or effect on [ $^3$ H]NMS binding.

**Figure 2.**
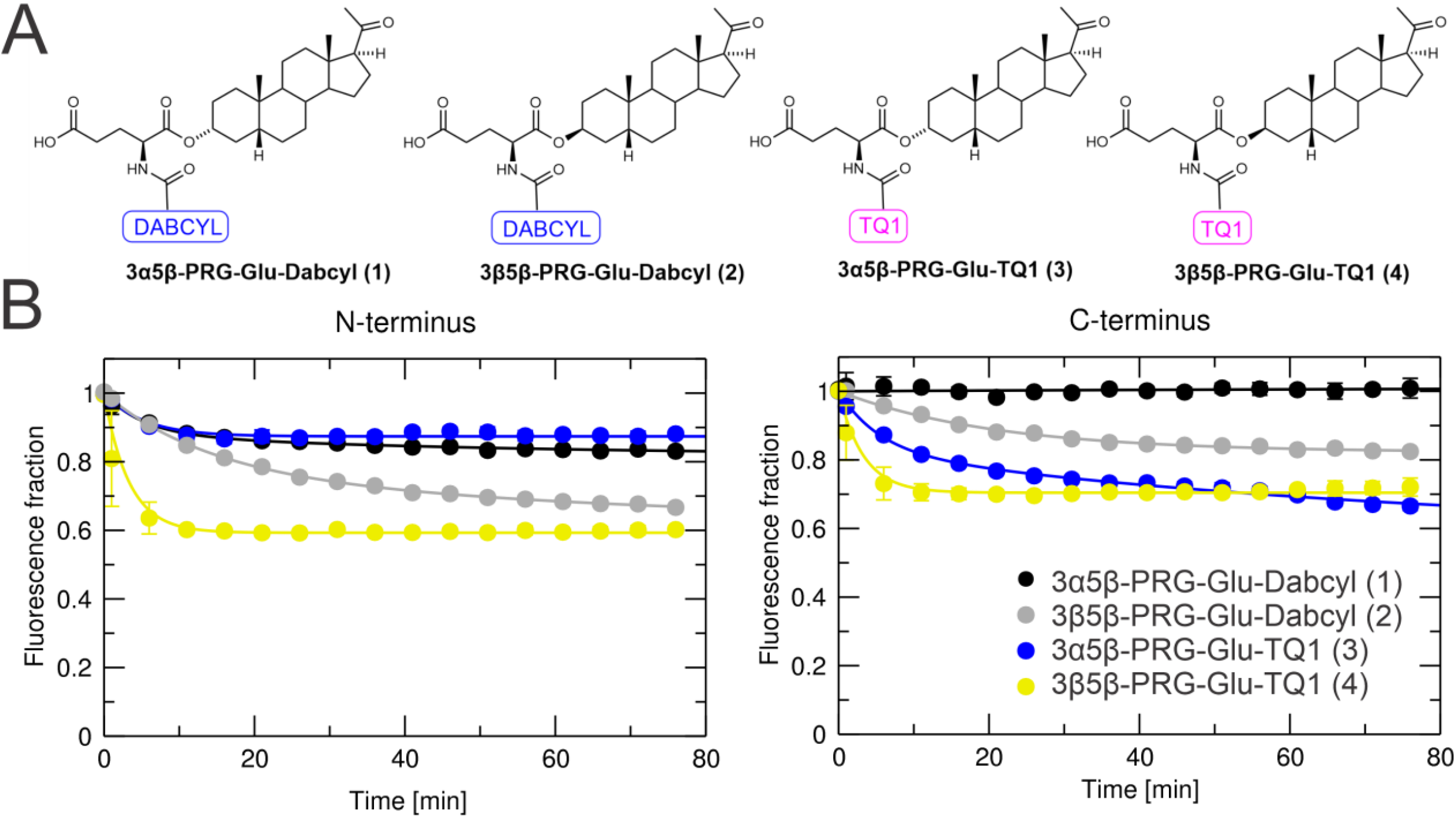
Fluorescence quenching by Dabcyl probes and TQ1 probes. (A) Structures of Dabcyl and TQ1 probes with 5β-skeletons. (B) Time courses (in minutes) of fluorescence quenching of CFP attached to the N-terminus (left) or C-terminus (right) of M_1_ muscarinic receptor expressed as a fraction of fluorescence before the addition of quencher, indicated in legend at final 1 μM concentration. Data are means ± SD from 3 independent experiments performed in dodecaplicates. Curves are fits of two exponential decays to the data.

In direct comparison, TQ1 probes outperformed Dabcyl probes in quenching experiments. The Dabcyl probe **3α5β-PRG-Glu-Dabcyl (1)** showed only minor quenching at the N-terminal-proximal site and none at the C-terminal-proximal site. The TQ1 probe **3α5β-PRG-Glu-TQ1 (3)** showed weak quenching at the N-terminus, but, unlike the Dabcyl analogue, was highly effective at the C-terminus. Interestingly, both 3β5β-substituted probes quenched fluorescence at the N-and C-terminal sites. Determined potencies for both TQ1 probes were approximately 1 μM at the C-terminal-proximal site, and 1 μM (**3β5β-PRG-Glu-TQ1 (4)**) or 3 μM (**3α5β-PRG-Glu-TQ1 (3)**) at the N-terminal-proximal site (Table 1). Both Dabcyl probes had potencies of 3 μM or less at both sites. The binding affinities of TQ1 probes ranged from 10 μM to 1 μM for **3β5β-PRG-Glu-TQ1 (4)**. Notably, **3β5β-PRG-Glu-TQ1 (4)** exhibited rapid association, reaching maximal quenching within 10 minutes, whereas **3β5β-PRG-Glu-Dabcyl (2)** at the N-terminal-proximal site and **3α5β-PRG-Glu-TQ1 (3)** at the C-terminal-proximal site showed slow association, not reaching equilibrium even after 75 minutes. These data identify TQ1 as the preferred quencher for further studies and indicate that the 3β5β stereochemistry markedly improves probe performance relative to the 3α5β configuration.

The stereochemistry of the steroid skeleton, particularly at the C-5 position, plays a pivotal role in defining the overall three-dimensional architecture. A 5β-configuration imparts a bent steroid backbone, whereas the 5α-configuration yields a planar structure (**Figure 1B, C**). These contrasting conformations strongly influence both physicochemical properties and receptor recognition. In addition, the 5α/5β orientation governs the spatial positioning of other substituents, including the functionally critical C-3 site. Specifically, in 5α-steroids, the C-3α substituent assumes an axial orientation relative to the backbone, while the C-3β orientation is equatorial. These differences in substituent placement dictate how chemical modifications are translated to the receptor. Notably, whereas previous results highlighted 3α5β as the most active configuration, the incorporation of TQ1 shifted the preference towards the 3β5β configuration, underscoring the interdependence of steroid stereochemistry and quencher selection. This finding demonstrates that the effect of stereochemistry and quencher choice is not simply additive. Consequently, all C-3/C-5 stereochemical combinations in the context of TQ1 were systematically evaluated to fully define the optimal architecture for neurosteroid-based allosteric modulation.

**scheme 1.**
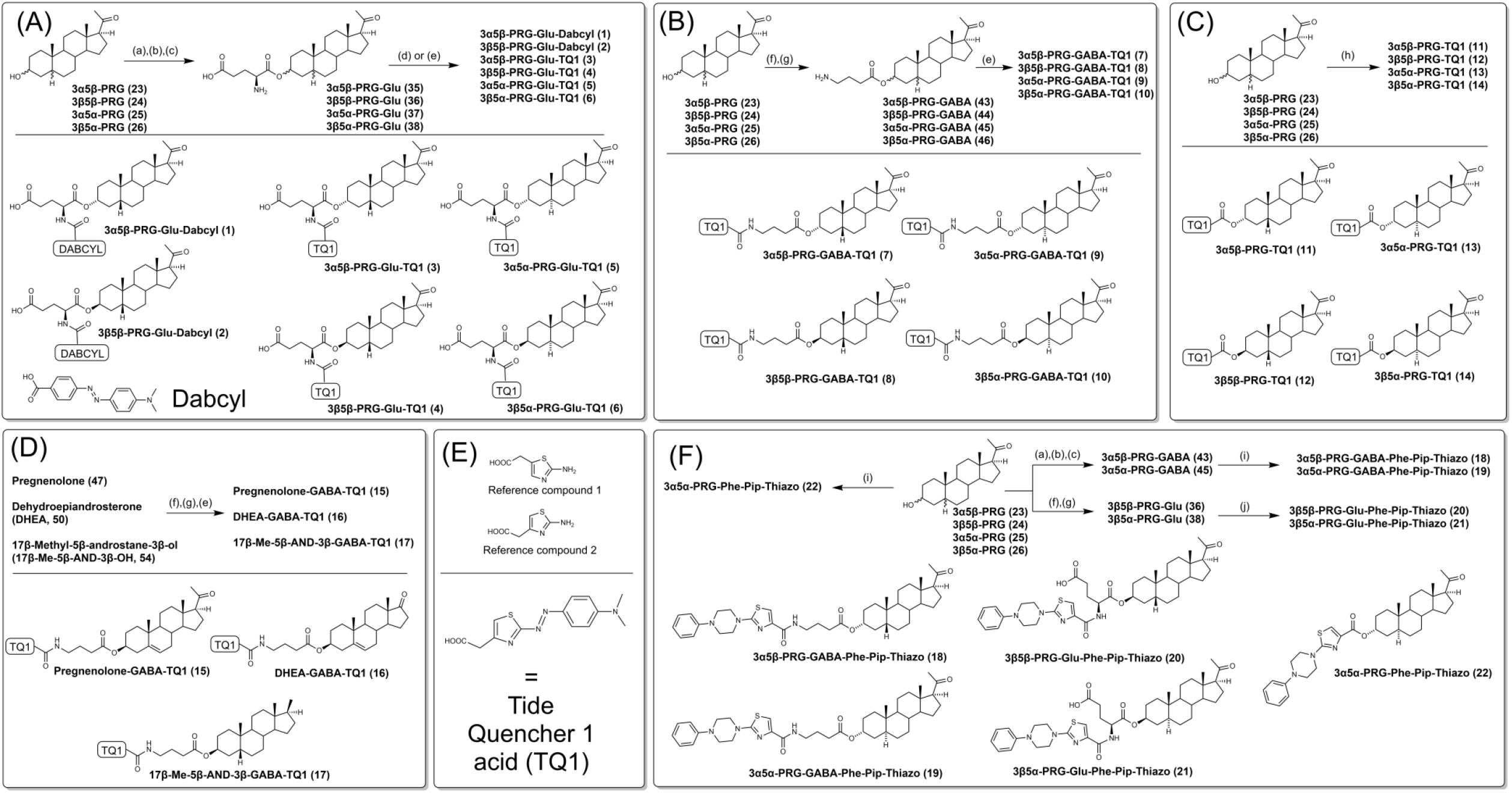
Synthesis of Dabcyl and TQ1 probes and their structures. Reagents and conditions: (a) Boc-Glu(OBzl)-OH, EDCI.HCl, DMAP, CH_2_Cl_2_, Ar, 0°C to rt, overnight; (b) Pd/C (10%), EtOH, 1 atm H_2_, overnight; (c) TFA, CH_2_Cl_2_, rt, 1 h; (d) Dabcyl succinimidyl ester, DIPEA, CH_2_Cl_2_, Ar, rt, overnight, in dark; (e) TQ1 succinimidyl ester, DIPEA, CH_2_Cl_2_, Ar, rt, overnight, in dark; (f) Boc-GABA-OH, EDCI.HCl, DIPEA, DMAP, CH_2_Cl_2_, Ar, rt, overnight; (g) TFA, CH_2_Cl_2_, 3 h, rt; (h) TQ1 acid, EDCI.HCl, DIPEA, DMAP, CH_2_Cl_2_, Ar, rt, overnight, in dark; (i) 2-(4-phenylpiperazino)-1,3-thiazole-4-carboxylic acid, EDCI.HCl, DIPEA, DMAP, CH_2_Cl_2_, Ar, overnight; (j) Phe-Pip-Thiazo succinimidyl ester (**57**), DIPEA, CH_2_Cl_2_, Ar, rt, overnight.

#### The effect of C-3/C-5 pregnane stereochemistry on FRET quenching

To fully describe the effect of C-3/C-5 stereochemistry, we have synthesised TQ1 glutamate derivatives with a 5α-skeleton (**Scheme 1A**). Subsequently, TQ1 probes with 5β-stereochemistry (**3α5β-PRG-Glu-TQ1 (3), 3β5β-PRG-Glu-TQ1 (4)**) and TQ1 probes with 5α-stereochemistry (**3α5α-PRG-Glu-TQ1 (5), 3β5α-PRG-Glu-TQ1 (6)**) were comparatively evaluated in quenching assays (**Figure 3B**). Interestingly, the 3β-probes (**3β5β-PRG-Glu-TQ1 (4)** and **3β5α-PRG-Glu-TQ1 (6)**) were more efficacious than the 3α-probes (**3α5β-PRG-Glu-TQ1 (3)** and **3α5α-PRG-Glu-TQ1 (5)**) at the N-terminal proximal site (**Figure 3B**). At the C-terminal proximal site, however, **3α5β-PRG-Glu-TQ1 (3)** and **3β5α-PRG-Glu-TQ1 (6)** showed a similar efficacy to **3β5β-PRG-Glu-TQ1 (4)**. Among this series, **3α5α-PRG-Glu-TQ1 (5)** exhibited one of the highest potencies (300 nM) at the N-terminal site (**Table 1**), but its potency at the C-terminal site was markedly lower (3 μM), indicating site preference for the N-terminus. By contrast, **3β5α-PRG-Glu-TQ1 (6)** showed comparable potencies (1 μM) at both sites. The potencies of **3β5β-PRG-Glu-TQ1 (4)** and **3α5β-PRG-Glu-TQ1 (3)** were 1 μM at the C-terminal site and 1 μM (3β5β) or 3 μM (3α5β) at the N-terminal site. Binding studies revealed affinities of 1 μM for **3β5α-PRG-Glu-TQ1 (6)** and **3β5β-PRG-Glu-TQ1 (4)**, while **3α5β-PRG-Glu-TQ1 (3)** displayed a weaker affinity (3 μM). For **3α5α-PRG-Glu-TQ1 (5)**, affinity could not be determined because it did not affect [^3^H]NMS binding. Association kinetics were fast for 3α5α-, 3β5α-, and 3β5β-probes, reaching equilibrium within 10 minutes, similar across these variants. Together, these data demonstrate that probe orientation at the 3-position strongly influences both site selectivity and potency, with 3β-substituted probes generally favouring higher efficacy, while **3α5α-PRG-Glu-TQ1 (5)** uniquely prefers the N-terminal site.

**Figure 3.**
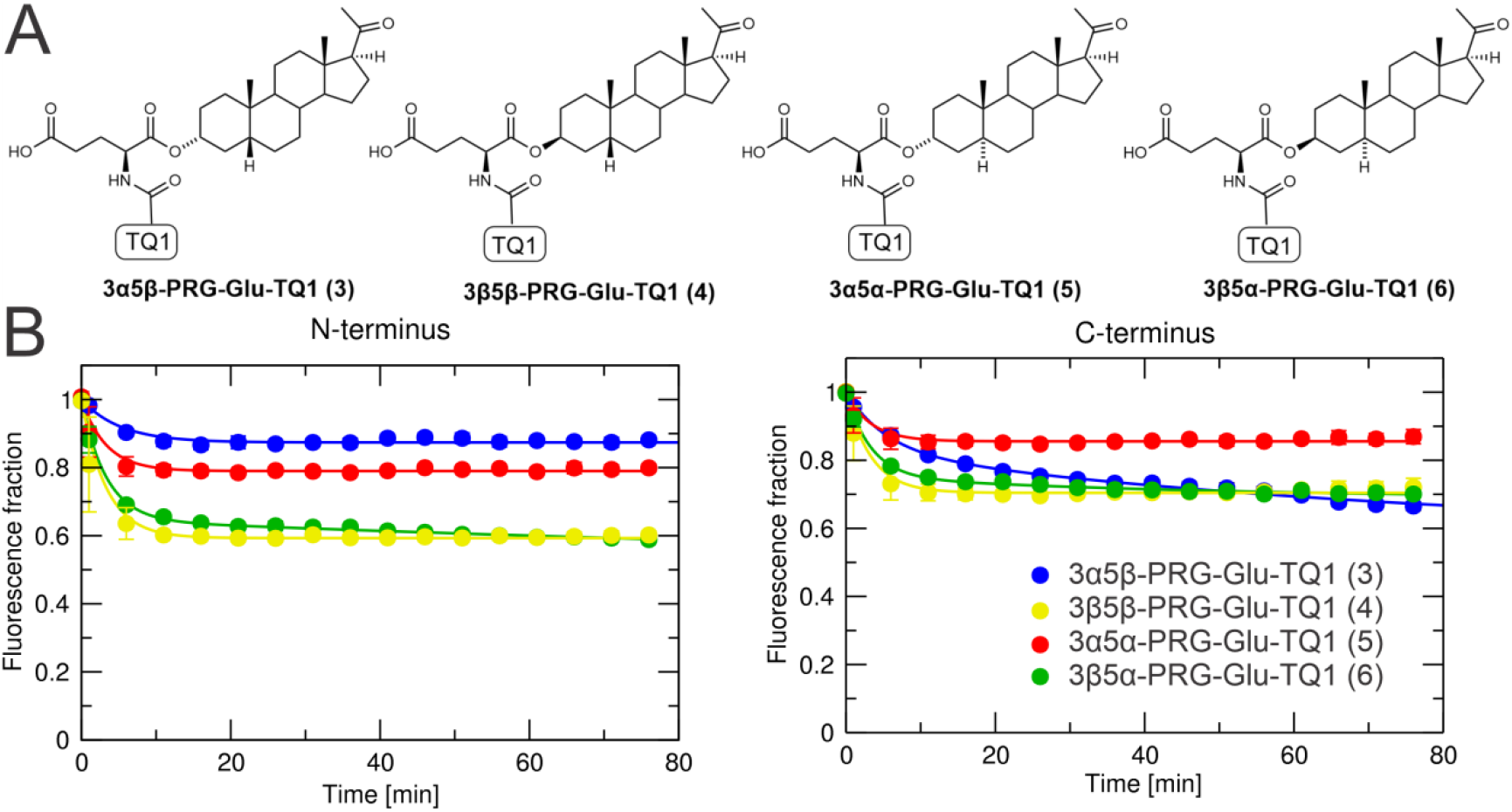
Fluorescence quenching by 5α- and 5β-pregnane glutamate esters tagged with TQ1. (A) Structures of TQ1 probes with 5α- and 5β-skeletons. (B) Time courses (in minutes) of fluorescence quenching of CFP attached to the N-terminus (left) or C-terminus (right) of M_1_ muscarinic receptor expressed as a fraction of fluorescence before the addition of quencher, indicated in legend at final 1 μM concentration. Data are means ± SD from 3 independent experiments performed in dodecaplicates. Curves are fits of two exponential decays to the data.

### Linker identity controls fluorescence quenching efficiency and site preference

To evaluate how the C-3 linker influences efficacy and potency, we designed two series of probes: one in which a linear γ-aminobutyric acid linker was conjugated to TQ1 (**Figure 4A**) and a second in which TQ1 was directly attached to the C-3 hydroxyl group (**Figure 4C**). The synthetic approach is outlined in **Scheme 1**. The 3-hydroxysteroid was first esterified with Boc-protected γ-aminobutyric acid (**Scheme 1B**), and subsequent Boc deprotection with trifluoroacetic acid provided the free amine. This intermediate was then coupled with the succinimidyl ester of TQ1. In the second approach, the TQ1 fluorophore was directly coupled to the steroid backbone (**Scheme 1C**). The 3-hydroxysteroid was directly esterified with the TQ1 dye, affording four distinct steroid-TQ1 conjugates.

**Figure 4.**
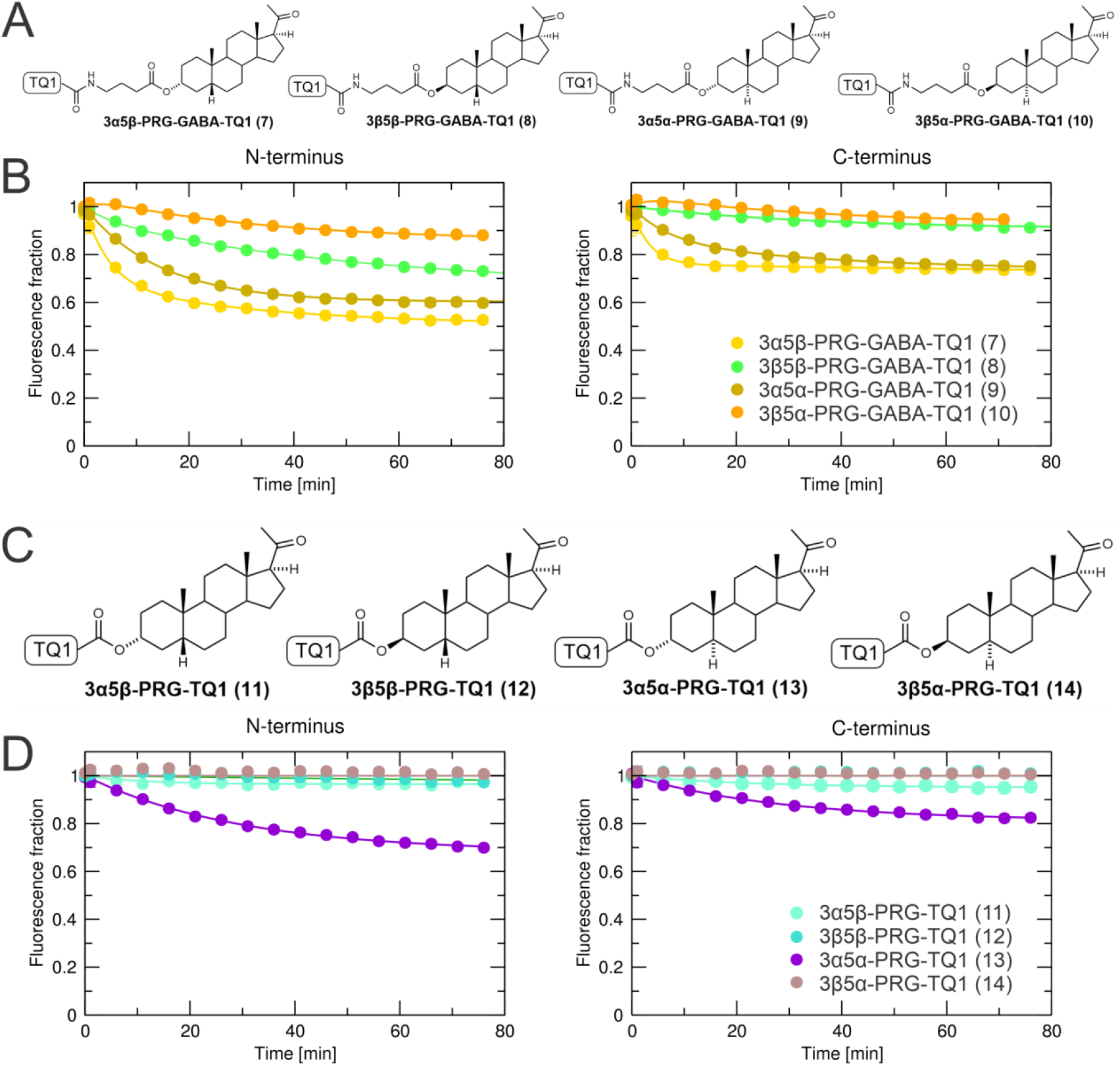
Fluorescence quenching by GABA-TQ1 steroids and steroids tagged with TQ1 directly at the C-3 position. (A, C) Structures of TQ1 probes tagged by a GABA linker or directly coupled to a steroid. (B, D) Time courses (in minutes) of fluorescence quenching of CFP attached to the N-terminus (left) or the C-terminus (right) of M_1_ muscarinic receptor expressed as a fraction of fluorescence before the addition of quencher, indicated in legend at final 1 μM concentration. Data are means ± SD from 3 independent experiments performed in dodecaplicates. Curves are fits of two exponential decays to the data

Among GABA-linked probes, **3α5β-PRG-GABA-TQ1 (7)** showed the highest efficiency, whereas **3β5α-PRG-GABA-TQ1 (10)** was the least effective (**Figure 4B**). Based on the concentration dependence of quenching effects, the potency of **3α5β-PRG-GABA-TQ1 (7)** at the N-terminal site was estimated at 500 nM, whereas its potency at the C-terminal site was 3 μM or more, as 100 nM concentrations produced no measurable quenching **(Table 1)**. Thus, **3α5β-PRG-GABA-TQ1 (7)** displayed potency and efficacy patterns comparable to those of the glutamate analogue **3α5α-PRG-Glu-TQ1 (5)**. By contrast, **3α5α-PRG-GABA-TQ1 (9)** exhibited higher potencies of 300 nM at the N-terminal site and 750 nM at the C-terminal site. Affinity estimates for the GABA-TQ1 probes ranged from 2 μM for **3β5β-PRG-GABA-TQ1 (8) and** 1 μM for **3β5α-PRG-GABA-TQ1 (10)** and **3α5α-PRG-GABA-TQ1 (9)**. The affinity of **3α5β-PRG-GABA-TQ1 (7)** could not be determined due to the absence of detectable effects on [^3^H]NMS binding. Except for **3α5β-PRG-GABA-TQ1 (7)**, all GABA esters exhibited slow association kinetics, reaching the equilibration was slower than with the glutamate-based conjugates **3β5α-PRG-Glu-TQ1 (6)** and **3β5β-PRG-Glu-TQ1 (4)**, requiring 20 min (C-terminus) to 40 min (N-terminus) to approach steady state.

For directly labelled steroids, only **3α5α-PRG-TQ1 (13)** produced detectable CFP quenching (**Figure 4**), and its association was markedly slow. The estimated potencies of **3α5α-PRG-TQ1 (13)** were 1 μM at the N-terminal and 3 μM at the C-terminal site (**Table 1**). Due to insufficient efficacy, potencies for other directly coupled probes could not be derived from fluorescence quenching assays. Radioligand binding, however, allowed determination of affinities for **3α5β-PRG-TQ1 (11)** (1 μM) and **3β5β-PRG-TQ1 (12)** (2 μM), whereas the affinity of **3β5α-PRG-TQ1 (14)** could not be assessed due to solubility limitations.

In summary, at the N-terminal site, the most efficacious probes were **3α5β-PRG-GABA-TQ1, 3α5α-PRG-GABA-TQ1 (9), 3β5α-PRG-Glu-TQ1 (6)**, and **3β5β-PRG-Glu-TQ1 (4)**, each quenching up to 40% of CFP fluorescence. The most potent probes at this site were **3α5α-PRG-Glu-TQ1 (5)** and **3α5α-PRG-GABA-TQ1 (9)**, both with EC_50_ values around 300 nM. Overall, **3α5β-PRG-GABA-TQ1 (7)** was identified as the most effective probe for N-terminal quenching. At the C-terminal site, **3α5β-PRG-Glu-TQ1 (3)** was the most efficacious probe (35% quenching), followed by **3α5β-PRG-GABA-TQ1 (7), 3α5α-PRG-TQ1 (13)**, and **3α5α-PRG-GABA-TQ1 (9)**, each achieving 20% quenching. Among these, **3α5α-PRG-GABA-TQ1 (9)** displayed the highest potency, with an EC_50_ of 750 nM. Thus, represents the most efficacious probe at the C-terminus, although its slow association kinetics may limit practicality due to the long incubation times required.

#### The pregnane skeleton is required for efficient fluorescence quenching

As the pregnane skeleton bearing a γ-aminobutyric acid linker was identified as the most effective skeleton in our study, we evaluated whether this structural element is a critical determinant for the quenching. Therefore, probes based on endogenous steroid skeletons (pregnenolone, dehydroepiandrosterone) and a synthetic 5β-androstane analogue, each functionalized with a γ-aminobutyric acid linker and conjugated to TQ1, were synthesised and evaluated (**Figure S1, Supporting Information**). None of these alternative skeletons produced robust quenching of CFP fluorescence at either site. The pregnenolone derivative induced only weak and transient quenching (~10% at the N-terminal site and 5% at the C-terminal site, appearing 5–15 minutes after addition), suggesting that **Pregnenolone-GABA-TQ1 (15)** interacts with the receptor but that its final binding state is not compatible with a stable quenching configuration. These data indicate that the pregnanolone core is critical for efficient receptor engagement and CFP quenching.

Comprehensive structure-activity relationship analysis indicated that steroidal probe performance at muscarinic receptors is governed by the interplay of three parameters: skeleton nature, C-3 linker identity, and steroidal stereochemistry. Direct coupling of TQ1 to the steroid diminished activity, except for **3α5α-PRG-TQ1 (13)**, whereas incorporation of a γ-aminobutyric acid linker consistently afforded high potency and balanced efficacy. Among steroidal frameworks, 5β-steroids supported robust receptor interaction, as both glutamate- and γ–aminobutyric acid–linked derivatives displayed high affinity toward both receptor N-terminus and at the C-terminus. In contrast, 5α-steroids exhibited site selectivity, exemplified by **3α5α-PRG-Glu-TQ1 (5)**, which retained high potency at the N-terminus but showed markedly reduced activity at the C-terminus. Collectively, these findings show that optimal activity of neurosteroid-based allosteric modulators arises from the convergence of linker chemistry, pregnane stereochemistry, and substituent orientation, which together define the geometry required for efficient muscarinic receptor modulation.

#### Deciphering of the Tide Quencher 1 structure

Tide Quencher− dyes, developed by AAT Bioquest, represent a commercially available family of chromophores notable for exceptionally high FRET efficiency and have been promoted as alternatives to DABCYL dyes. Despite their widespread use, the chemical structures of these dyes are not disclosed by the manufacturer. To enable the synthesis of non-quenching analogues for our further mechanistic studies, we therefore undertook structural elucidation of the Tide Quencher 1 (TQ1) acid. Comprehensive spectroscopic analyses - including 1D (1H and 13C) and 2D (COSY, ROESY, HSQC, HMBC) NMR experiments, supported by elemental analysis and high-resolution mass spectrometry - were used to assign the structure of TQ1 acid. The definitive localisation of the acetic acid group was established through comparative examination of the NMR spectra of reference compounds **1** and **2** (**Figure S2, Supporting Information**). Our combined data confirm that Tide Quencher 1 acid corresponds to 3-[(N,N-dimethylaminophenyl)-4′-diazenyl]thiazoleacetic acid, thereby providing a defined chemical basis for further analogue design (**Scheme 1E**). The 1H NMR spectrum of TQ1 acid matches that reported in the literature for this compound (Fraga et al., 2004); however, prior publications do not explicitly identify this structure as Tide Quencher 1.

#### Competition experiments between quencher probes and their non-quenching analogues

With the structure of Tide Quencher 1 established, we evaluated the specificity of TQ1 probe binding in competition experiments using corresponding non-quenching analogues. As non-specific binding sites have effectively unlimited capacity, they are not saturated and therefore cannot be efficiently competed by a non-quenching analogue; in such a scenario, quenching would remain unchanged. In contrast, a decrease in quenching in the presence of a non-quenching analogue indicates competition for particular specific sites. For competition experiments, we selected the most potent and efficacious TQ1 probes **(3α5β-PRG-GABA-TQ1 (7), 3α5α-PRG-GABA-TQ1 (9), 3β5α-PRG-Glu-TQ1 (6), 3β5β-PRG-Glu-TQ1 (4)**, and **3α5α-PRG-TQ1 (13)**), spanning all linker types (GABA-linked, Glu-linked, and directly coupled steroids), and paired each with its corresponding non-quenching analogue (**Figure 5**). The selection of structures for synthesising these non-quenching analogues was guided by the potency and affinity profiles of the corresponding TQ1 probes to ensure their suitability for competition studies. As the non-quenching moiety, we selected 2-(4-phenylpiperazino)-1,3-thiazole-4-carboxylic acid, which lacks the azo group but retains the phenyl and thiazole rings of TQ1 acid and exhibits distinct spectral properties (λ_max_ = 248 nm in MeOH), avoiding interference with quencher absorbance.

**Figure 5.**
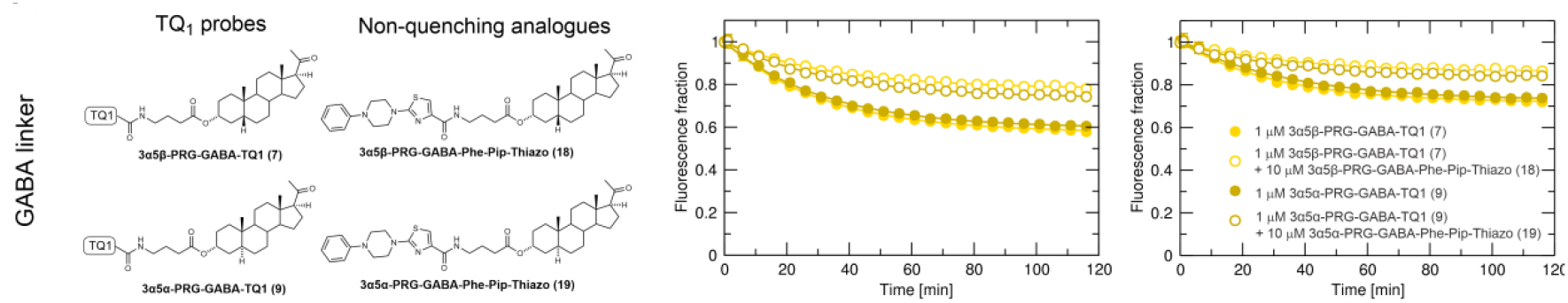
Competition between quenchers and their non-quenching analogues. Structures of TQ1 probes and their non-quenching analogues tagged by a GABA linker and their time courses (in minutes) of fluorescence quenching of CFP attached to the N-terminus (left) or the C-terminus (right) of M_1_ muscarinic receptor expressed as a fraction of fluorescence. Data are means ± SD from 3 independent experiments performed in dodecaplicates. Curves are fits of two exponential decays to the data.

Quenching of **3α5β-PRG-GABA-TQ1 (7)** and **3α5α-PRG-GABA-TQ1 (9)**, respectively, at 100 nM was weak and not suitable for competition experiments. Therefore, the quenchers were used at a concentration of 1 μM. The non-quenching GABA analogues **3α5β-PRG-GABA-Phe-Pip-Thiazo (18)** and **3α5α-PRG-GABA-Phe-Pip-Thiazo (22)** (**Figure 5A**) reduced quenching of the corresponding TQ1 probes by approximately 50% at both the N- and C-terminal labelling sites, when applied at a 10-fold molar excess. This decrease in quenching indicates that quencher and non-quencher steroids compete for the same specific binding sites, rather than producing quenching through non-specific interactions.

Importantly, 10 μM **3α5β-PRG-GABA-Phe-Pip-Thiazo (18)** completely competes out 100 nM **3β5α-PRG-Glu-TQ1 (6)** at both sites (Supplementary Information Figure S2). These data indicate that a compound with a GABA linker and a different scaffold competes with the directly coupled quencher at these sites. However, 10 μM **3α5α-PRG-GABA-Phe-Pip-Thiazo (19)** only weakly decreases 100 nM **3α5α-PRG-TQ1 (13)**. These data indicate a very low affinity of **3α5α-PRG-GABA-Phe-Pip-Thiazo (19)**.

### Mutagenesis study

#### Identification of residues for mutagenesis by docking and molecular dynamics analysis

In our recent study (Chetverikov et al., 2025), we have identified six potential steroid-binding sites on muscarinic receptors. Among these, the e1-e2-e7 site in the extracellular leaflet of the membrane is the closest to the N-terminus, and the i1-i2-i4 site in the intracellular leaflet of the membrane is the closest to the C-terminus. Therefore, probes **3α5β-PRG-GABA-TQ1 (7)** (the most effective probe for both C- and N-terminal quenching) and **3α5α-PRG-GABA-TQ1 (9)** (the most potent probe for C-terminal quenching) were docked to these two sites. Both probes adopted favourable poses at each site (**Figure 6**). At the N-terminal proximal site, both probes formed a hydrogen bond with Q24 and a π–π interaction with W405, whereas at the C-terminal proximal site, they formed a hydrogen bond with K57 and a π–π interaction with W150. Interestingly, R140, a key residue of the cholesterol-consensus motif (CCM), was oriented outside the binding pocket. Top poses were subjected to three independent runs of molecular dynamics (MD) simulations. Analysis of MD trajectories for both probes at the N- and C-terminal sites indicated major contributions of K20, Q24, and W405 to binding at the N-terminal site, and of K54, K57, Y62, and W150 at the C-terminal site (Supplementary Information Figure S3). Lysine and glutamine residues primarily interacted with probe oxygen atoms via direct and water-mediated hydrogen bonds, whereas tyrosine and tryptophan residues contacted the steroid core through hydrophobic and π–π interactions. To further test these modelling-based predictions, K20 and Q24 of the N-terminal CFP-tagged receptor and K51 and K57 of the C-terminal CFP-tagged receptor were mutated to alanine. In contrast, Y62, W150, and W405 cannot be mutated to study hydrophobic interaction (mutated by charged amino acid) with probes, as such mutagenesis would cause substantial disturbance of the receptor structure.

**Figure 6.**
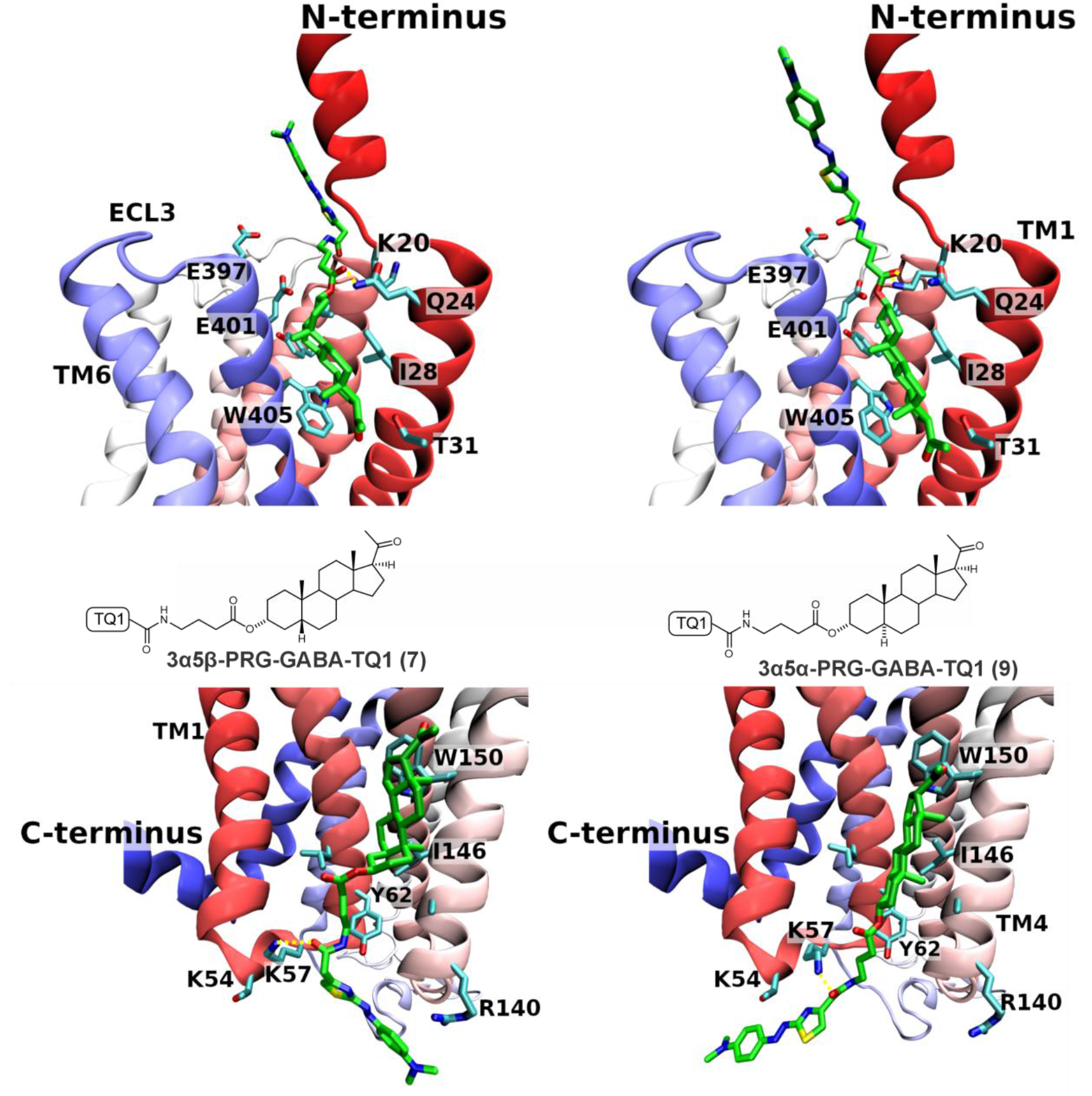
Docking of probes 3α5β-PRG-GABA-TQ1 (7) and 3α5α-PRG-GABA-TQ1 (9). Top-ranked docking poses of probes 3α5β-PRG-GABA-TQ1 (7) (left) and 3α5α-PRG-GABA-TQ1 (9) (right) at the N-terminus (top) and C-terminus (bottom) proximal sites of the M_1_ receptor are shown. Binding site residues are displayed. Dashed yellow lines denote hydrogen bonds between receptor and ligand. The receptor backbone is coloured with a red-white-blue gradient. Atom colours are as follows: green (ligand carbon), cyan (receptor carbon), blue (nitrogen), red (oxygen), yellow (sulphur).

#### Key binding resides in the N- and C-terminal proximal site

Both mutations in the putative N-terminal site (K20A and Q24A) markedly reduced quenching by **3α5β-PRG-GABA-TQ1 (7)** and **3α5α-PRG-GABA-TQ1 (9)** probes, confirming their substantial contribution to the probe binding at this site, as predicted by molecular modelling (**Figure 7B**, left). In contrast, neither single mutation in the putative C-terminal site (K51A or K57A) affected quenching by either probe (Supplementary Information Figure S4). Therefore, the double mutation K51A K57A was constructed. It produced a small reduction in quenching, indicating a partial contribution of these residues to binding at the C-terminal proximal site (**Figure 7B**, right). This result is therefore consistent with the modelling, which predicts that binding at this site is dominated by hydrophobic contacts with Y62 and W150 (Supplementary Information Figure S3). Together, these mutagenesis data demonstrate the N-terminal site as a steroid-binding epitope centred on K20 and Q24, while indicating that the C-terminal site involves a more distributed interaction network in which K51 and K57 make only a partial contribution.

**Figure 7.**
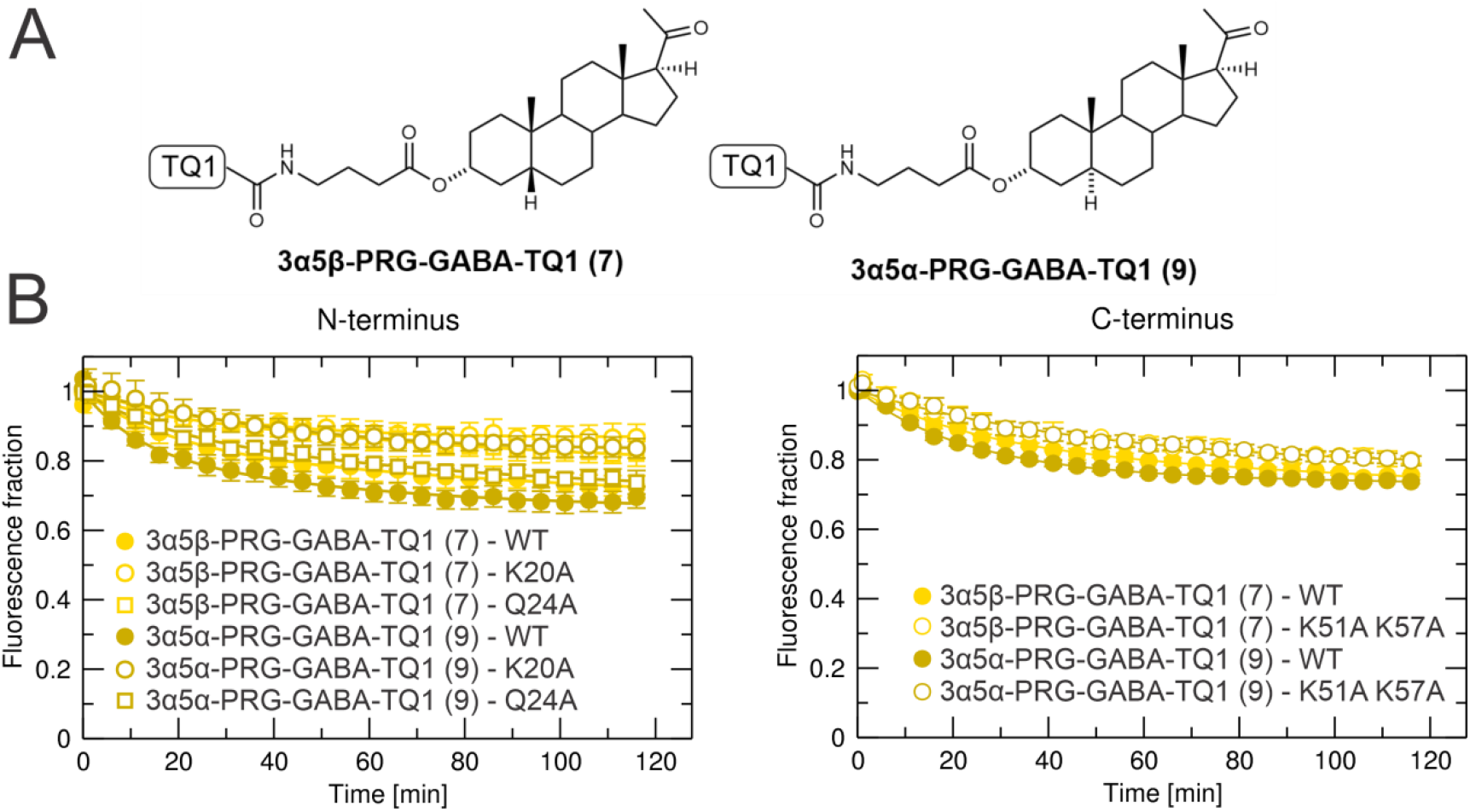
Effect of mutations on quenching by 3α5β-PRG-GABA-TQ1 (7) and 3α5α-PRG-GABA-TQ1 (9). Time courses (in minutes) of fluorescence quenching of CFP attached to the N-terminal site (left) and C-terminal site (right) of M_1_ muscarinic receptor expressed as a fraction of fluorescence before addition of quenchers (**3α5β-PRG-GABA-TQ1 (7)**, yellow, or **3α5α-PRG-GABA-TQ1 (9)**, gold) at the final 1 μM concentration. Quenching was measured at wild-type receptors (WT, closed symbols) or mutated receptors (open symbols). Data are means ± SD from 3 independent experiments performed in dodecaplicates. Curves are fits of two exponential decays to the data.

## Discussion

G protein-coupled receptors (GPCRs) are pivotal drug targets, with cholesterol playing a key but mechanistically unconventional role in their modulation. There are approximately 337 agents in clinical trials targeting 133 GPCRs, including 30 novel targets. Many of these agents are allosteric modulators or biologics, offering new therapeutic strategies (Lorente et al., 2025). Despite the absence of conserved cholesterol-recognition motifs, cholesterol stabilises GPCR conformations through site-specific interactions (Sengupta & Chattopadhyay, 2015). This challenges traditional motif-based predictions and highlights the need for residue-level analysis in lipid-receptor crosstalk (Geiger et al., 2021; Gimpl, 2016).

Structural studies reveal cholesterol binding to GPCRs at specific transmembrane sites (e.g., TM6 residues like R^6.35^ in muscarinic receptors (Randáková et al., 2018)) without relying on classical cholesterol-binding motifs (CRAC/CARC or CCM) (Baier et al., 2011; Hanson et al., 2008; Li & Papadopoulos, 1998). Neurosteroids and steroid hormones exploit these cholesterol-binding pockets to allosterically modulate GPCRs at nanomolar concentrations (Dolejší, Szánti-Pintér, et al., 2021), offering untapped therapeutic potential. To study these interactions, we have developed a fluorescence-quenching assay using steroid-dye conjugates, overcoming limitations of traditional radioligand methods by the possibility of separating free radioligand from radioligand bound to the receptor. The fluorescence-quenching assay brings further advantages over radioligand methods like real-time binding dynamics, enabling kinetic studies of lipid-GPCR interactions, or mutational mapping, directly linking cholesterol-site residues to functional effects. Moreover, unlike in FRET measurements, quencher concentration may exceed the concentration of the receptor, allowing detection of ligands with relatively low (micromolar) affinity. However, unlike FRET, quenching is not ratiometric, and thus a concentration of donor (CFP-tagged receptor in our case) affects measurement and requires data normalisation and precaution. In this assay, we took advantage of *Spodoptera frugiperda* (Sf9)-insect cells that express receptors at high densities after baculovirus infection, which is desired for fluorescent techniques. Moreover, the cholesterol/phospholipid ratio in Sf9 cells is about 10-times lower than in mammalian cell lines (Dawaliby et al., 2016; Marheineke et al., 1998). This diminishes the competition of membrane cholesterol with steroid probes for receptor binding.

Our structure–activity analysis shows that steroidal quencher probe performance at M_1_ is governed by a combination of steroid skeleton, C-3 linker identity, and C-3/C-5 stereochemistry. Among dark quenchers, TQ1 clearly outperformed Dabcyl, with TQ1 conjugates displaying higher E_max_ values, faster association kinetics, and lower EC_50_ values at both N- and C-terminal sites. Within the TQ1 series, 3β-pregnane substitution with glutamate-conjugated TQ1 exhibits higher efficacy in quenching, while **3α5α-PRG-Glu-TQ1 (5)** stands out as a highly potent probe that selectively favours the N-terminal site (300 nM versus 3 µM for the C-terminal site). The introduction of GABA linkers significantly improved quenching potency at the N-terminal site for probes **3α5β-PRG-GABA-TQ1 (7)** (500 nM) and **3β5α-PRG-GABA-TQ1 (10)** (500 nM), while **3α5α-PRG-GABA-TQ1 (9)** exhibited higher potencies of 300 nM at the N-terminal site and 750 nM at the C-terminal site, yet the selectivity for the C-terminal site as compared to glutamate-conjugated was maintained. Direct coupling of TQ1 to the C-3 hydroxyl largely abolished quenching despite preserved radioligand affinity (Table 1), while introduction of a γ-aminobutyric acid linker consistently restored high potency and balanced efficacy, identifying pregnane–GABA–TQ1 as an optimal architecture. Alternative steroid skeletons (pregnenolone, DHEA, and a 5β-androstane analogue) failed to produce stable quenching, indicating that the pregnane core is critical for forming a quenching-competent complex with M_1_. Among synthesised probes, **3α5β-PRG-GABA-TQ1 (7), 3α5α-PRG-GABA-TQ1 (9)**, and **3α5α-PRG-Glu-TQ1 (5)** showed the highest quenching efficacy and potency at the N-terminal site, with **3α5α-PRG-GABA-TQ1 (9)** being the most effective at the C-terminal site.

Theoretically, quenching may arise from membrane microdomain partitioning or mere collisional interaction rather than from a specific binding. **3β5α-PRG-TQ1 (14)** did not quench the fluorescence of either the N-terminus or the C-terminus attached to CFP (**Figure 4**), being completely inactive. Therefore, collisional quenching or micro-domain partitioning can be excluded.

The two terminal sites differed in their SAR fingerprints. At the N-terminal site, **3α5β-PRG-GABA-TQ1 (7)** and **3α5α-PRG-GABA-TQ1 (9)**, together with **3β5α-(6)** and **3β5β-PRG-Glu-TQ1 (4)**, achieved up to ≈ 40% quenching with submicromolar EC_50_ values, indicating that both GABA and glutamate linkers can be present at the steroid probe and efficiently contribute to quenching at this site. In contrast, at the C-terminal proximal site, the most efficacious probes were **3α5β-PRG-Glu-TQ1 (3)**, which reached 35% quenching (EC_50_ = 1 µM) and **3α5α-PRG-GABA-TQ1 (9)**, which reached 21% quenching (EC_50_ = 750 nM). These patterns suggest that the N-terminal site tolerates multiple linker–stereochemistry combinations with robust, submicromolar activity, whereas the C-terminal site favours 3α-substituted probes and generally supports more modest efficacy and potency, which may be caused by a broader distance between CFP and the binding site.

Evaluation of non-quenching analogues provided an essential control for distinguishing specific from non-specific quenching. Because non-specific interactions are effectively high-capacity, they are not saturated and should not be competed by the particular non-quenching probe. The most potent and efficacious TQ1 probes (**3α5β-PRG-GABA-TQ1 (7), 3α5α-PRG-GABA-TQ1 (9), 3β5α-PRG-Glu-TQ1 (6), 3β5β-PRG-Glu-TQ1 (4)**, and **3α5α-PRG-TQ1 (13)**) were paired with their corresponding non-quenching analogue bearing the non-quenching moiety of 2-(4-phenylpiperazino)-1,3-thiazole-4-carboxylic acid that is structurally similar to TQ1. The non-quenching GABA analogues **3α5β-PRG-GABA-Phe-Pip-Thiazo (18)** and **3α5α-PRG-GABA-Phe-Pip-Thiazo (22)** significantly reduced quenching of the corresponding TQ1 probes by approximately 50% at both the N- and C-terminal labelling sites, when applied at a 10-fold molar excess (**Figure 5**), demonstrating that a substantial component of the signal arises from occupancy of specific binding sites rather than from non-specific collisional quenching.

To further validate the specificity of quenching, we designed mutations at putative binding sites aimed to impair the binding of probes and hence decrease their quenching. Docking of GABA probes to the N-proximal e1-e2-e7 site in the extracellular leaflet of the membrane and the C-proximal i1-i2-i4 site in the intracellular leaflet of the membrane confirmed possible binding of these probes to these sites (**Figure 6**). At the N-proximal e1-e2-e7 site, both probes formed a hydrogen bond with Q24. At the C-proximal i1-i2-i4 site, both probes formed a hydrogen bond with K57. The C-proximal i1-i2-i4 site corresponds to the binding of cholesterol hemisuccinate to the M_1_ receptor in X-ray and cryo-EM structures (5CXV, 6OIJ) (Maeda et al., 2019; Thal et al., 2016). This site is adjacent to the CCM motif (Hanson et al., 2008). However, none of the probes interacted with R140, a key residue of the CCM motif. Analysis of subsequent MD trajectories for both probes at the N- and C-terminal sites indicated major contributions of K20, Q24, and W405 to binding at the N-terminal site, and of K51, K57, Y62, and W150 at the C-terminal site. Charge elimination by mutation of K20 as well as Q24 to alanine strongly diminished quenching of **3α5β-PRG-GABA-TQ1 (7)** and **3α5α-PRG-GABA-TQ1 (9)**, respectively (**Figure 7B**, left), at the N-terminal site, indicating a crucial contribution of K20 and Q24 to the binding of these probes. On the other hand, charge elimination by mutation of K51 or K57 to alanine did not affect quenching of **3α5β-PRG-GABA-TQ1 (7)** nor **3α5α-PRG-GABA-TQ1 (9)** at the C-terminal site. The double mutation K51A and K57A only slightly decreased quenching of these probes (**Figure 7B**, right), indicating only partial contribution of K51 and K57 to binding of these probes, suggesting a dominant role of hydrophobic contacts with Y62 and W150 in binding to the C-terminal site. The data indicate that hydrogen bonding to K20 and Q24 primarily drives N-terminal site binding, while hydrophobic interactions are key for C-terminal site binding. These findings confirm the specificity of quenching and binding for both **3α5β-PRG-GABA-TQ1 (7)** and **3α5α-PRG-GABA-TQ1 (9)**. Additionally, the results validate existing models of N- and C-terminal binding sites, supporting their structural accuracy.

In summary, pregnane-based TQ1 probes with a γ-aminobutyric acid linker define a chemically and functionally optimised set of tools for the discovery of non-canonical steroid-binding sites on M_1_ muscarinic receptors. By integrating SAR, docking, molecular dynamics, competition assays, and mutagenesis, we delineated two distinct terminal sites that recognise neurosteroid-like ligands and couple their binding to robust fluorescence quenching. In principle, this framework may be broadly adaptable to other GPCRs with unresolved lipid-binding sites, offering a route to systematic, structure-based exploration of cholesterol and neurosteroid allostery of GPCRs in a membrane environment.

## Materials and Methods

### Synthesis – General description

Pregnanolone glutamate was selected for quencher dye conjugation owing to its accessible functional groups - an amino group, a keto group, and a carboxylic acid, which provide well-defined sites for chemical derivatisation. As shown in **Scheme 1A**, the conversion of the 3-hydroxysteroid afforded Boc- and benzyl-protected *L*-glutamic acid esters. Subsequent palladium-catalysed hydrogenolysis removed the benzyl protecting group, and Boc deprotection with trifluoroacetic acid afforded the free amino glutamate ester intermediates. These species were then coupled with succinimidyl esters of quenchers such as Dabcyl-OSu or TQ1-OSu to obtain the final amide-linked conjugates. In an alternative route (**Scheme 1B**), the 3-hydroxysteroid was esterified with Boc-protected γ-aminobutyric acid; after Boc removal, the resulting free amine was coupled with TQ1-OSu. **Scheme 1C** further details the direct attachment of the TQ1 fluorophore to the steroid skeleton, yielding four structurally distinct steroid– TQ1 conjugates. Full synthetic procedures are provided in the Supporting Information. All newly obtained probes were characterised by 1H and 13C NMR spectroscopy and HPLC-HRMS, confirming purities exceeding 95%. Fluorescence spectra of all conjugates are also included in the Supporting Information.

### Preparation of DNA constructs

The pcDNA3.1 plasmid containing the M_1_-muscarinic receptor fused with CFP at the C-terminus(Ziegler et al., 2011) was provided by Dr Carsten Hoffman. The pcDNA3.1 plasmid containing M_1_-muscarinic receptor fused with CFP at the N-terminus was derived from the previous construct and EmGFP-M1 plasmid bearing GFP at the N-terminus of the receptor using the Gibson Assembly® technique.

### Generation of recombinant baculovirus

Baculovirus/*Spodoptera frugiperda* (Sf9)-insect cells expression system represents a well-defined system enabling recombinant expression of a combination of given GPCRs with individual G-protein α-subunits in insect cells at a high level and without contamination by interfering GPCRs and with a limited set of endogenous G-proteins(Houston et al., 2002).

Baculoviral stocks for expression of M_1_ receptor fused with CFP at different positions (C-terminus, N-terminus) in SF9 cells were generated according to the Bac-to-Bac® baculovirus expression system— user guide. Briefly, pcDNA3.1 plasmids coding the receptor-fluorescent protein construct were subcloned into the pFastBac1 donor plasmid (Thermo Fisher Scientific; LOT: 2065047) using restriction endonucleases. pFastBac constructs were transformed into MAX Efficiency® DH10Bac− competent E. coli (Thermo Fisher Scientific; LOT: 2443297) to generate a recombinant bacmid. Bacmid DNA was transfected into adherent insect cell line Sf9 in a 12.5 cm^2^ flask in the number of 1.5 × 10^6^ cells per flask in 3 ml of Grace’s medium (Gibco; LOT: 2083249) using 10 µl of Cellfectin reagent (Thermo Fisher Scientific; LOT: 2094064) and 1.3 µg bacmid DNA. After 96 h at 27 °C, baculoviral particles released into the medium were harvested. After centrifugation at 500×g for 10 min to remove detached cells and cell debris, baculoviral stock P_0_ was ready for amplification. Suspension of Sf9 cells at a density of 2 × 10^6^ cells per ml was infected by baculoviral stock P_0_ in a ratio of 1:1000, and after 72 h of growth (in a shaking incubator at 135 rpm and 27 °C), the amplified baculoviral stock P_1_ was harvested. The cell suspension was centrifuged at 500×g for 10 min to remove cells and cell debris, and the supernatant containing a high level of baculoviral particles P_1_ was collected. FBS to the concentration of 0.5% was added, and the stock was stored at 4 °C, protected from light for up to 6 months. For the quantification of baculoviral particles, a plaque assay was used. The titre of baculoviral stock was 3–9 × 10^7^ pfu/ml (plaque-forming unit per ml).

### Cell culturing and transfection, membrane preparation

Sf9 cells (Gibco− Cat. No. 12659017) were purchased from Thermo Fisher Scientific (Waltham, MA, USA) and maintained according to provider guidelines (https://assets.thermofisher.com/TFS-Assets/LSG/manuals/Sf9_SFM_II_SFM_III_man.pdf). Cells were grown in suspension culture in 250 ml Erlenmeyer flasks in Sf-900− III serum-free medium in an aerated shaking incubator at 27 °C and 135 rpm. Cells were maintained at a density of 1–4 × 10^6^ cells per ml. Sf9 cells at a density of 2 × 10^6^ were infected with a baculoviral stock of either M1-CFP-C-terminus or M_1_-CFP-N-terminus receptor construct at the multiplicity of infection (MOI) = 1. Infected cells were harvested 96 hours after infection by centrifugation at 1000×g for 10 min. Cell pellets were kept at − 80 °C.

Cells harvested from 40 ml of cell suspension (Sf9) were resuspended in 20 ml of ice-cold incubation medium (100 mM NaCl, 20 mM Na-HEPES, 10 mM MgCl_2_, pH 7.4) supplemented with 10 mM EDTA and homogenized on ice by two 30-s strokes using a Polytron homogenizer (Ultra-Turrax; Janke & Kunkel GmbH & Co. KG, IKA-Labortechnik, Staufen, Germany) with a 30 s pause between strokes. Cell homogenates were centrifuged for 5 min at 1000×g to remove whole cells and cell nuclei. The resulting supernatants were centrifuged for 30 min at 30,000×g. Pellets were resuspended in a fresh incubation medium, incubated on ice for 30 min, and centrifuged again. The resulting membrane pellets were kept at − 80 °C until assayed within 10 weeks.

### Fluorescence quenching assay

Fluorescence quenching experiments were carried out on black 96-well plates (BRAND GmbH & Co KG, Wertheim, Germany). Fresh solutions of quencher probes dissolved in DMSO were prepared before each experiment. A suitable combination of the steroid-quencher probe and membrane suspension was added to the wells to a final volume of 100 µl. KHB assay buffer (final concentrations in mM: NaCl 138; KCl 4; CaCl_2_ 1.3; MgCl_2_ 1; NaH_2_PO_4_ 1.2; HEPES 20) was added to the control wells instead of the steroid-quencher probe. Fluorescence quenching measurements were performed at BioTek Cytation 3 Multi-Mode Reader (BioTek Instruments, Winooski, USA). Measurements were carried out at 5-minute intervals for 75 - 140 minutes at 25°C. The excitation and emission wavelengths were 435 nm and 485 nm, respectively.

### Experimental data and statistical analysis

The data and statistical analysis comply with the recommendations of the British Journal of Pharmacology on experimental design and analysis in pharmacology (Curtis et al., 2025). Experiments were independent, using different seedings and transfections of Sf-9 cells, followed by membrane preparation. Radioligand binding experiments as well as fluorescence assays were carried out in three to five independent experiments with samples in quadruplicate for radioligand binding and in dodecaplicates for fluorescence quenching experiments. Experimenters were blind to the chemical structures of the tested compounds. After subtraction of non-specific binding (binding experiments) or background/blank values (fluorescence experiments), data were normalised to control values determined in each experiment. IC_50_ and EC_50_ values were analysed after conversion to their logarithms. All data were included in the analysis, no outliers were excluded, and the normality of distribution was checked. In statistical analysis, a value of P < 0.05 was taken as significant for all data. When testing a single concentration, the one-sample t-test was used. In multiple comparison tests, ANOVA with P <.05 was followed by Tukey’s post-test (P < 0.05).

To determine the potency of steroid probes to quench CFP fluorescence (EC_50_), the observed maximal quenching E’_MAX_ (y) was plotted against concentration of the quencher probe (x) and Equation 1 was fitted to the data.

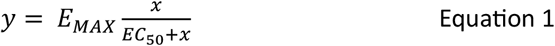

Where E_MAX_ is the maximum possible quenching by the probe.

### Molecular modelling

Structures of the M_1_-muscarinic acetylcholine receptor (5CXV) and α_2C_-adrenergic receptors (6KUW) were downloaded from the RCSB Protein Data Bank (https://www.rcsb.org/). Non-protein molecules (except ligands) and nanobody or hydrolase parts of protein were deleted, and the resulting receptor protein was processed in Maestro using Protein Preparation Wizard according to Sastry et al. guidelines (Sastry et al., 2013).

To model probes binding to the site proximal to the C-terminus, the steroid core of the probe was aligned with cholesterol hemisuccinate between TM1 and TM4 on the intracellular side of 5CXV. To model probes binding to the site proximal to the N-terminus, cholesterol between TM7 and TM1 on the extracellular side of 6KUW was mapped on 5CXV by alignment of receptor structures using MUSTANG (Konagurthu et al., 2006) followed by energy minimisation in YASARA (Krieger et al., 2002) The quenching probes were docked using the YASARA implementation of AutoDock (Trott & Olson, 2010). The binding site was defined as a 3 Å extension in all directions to the cholesterol molecule, and the AutoDock local search method was employed. All resulting poses were re-scored in YASARA by energy-minimisation of ligand pose and AutoDock VINA’s local search, confined closely to the original ligand pose. The top-scoring pose was taken for further evaluation.

## Supporting information

Supplemntary Information - Chemistry

