## Supplementary material for "Steroid‑based Tide Quencher 1 probes enable real‑time mapping of novel non‑canonical cholesterol sites on the M1 muscarinic receptor": Supplemtary Information - Data

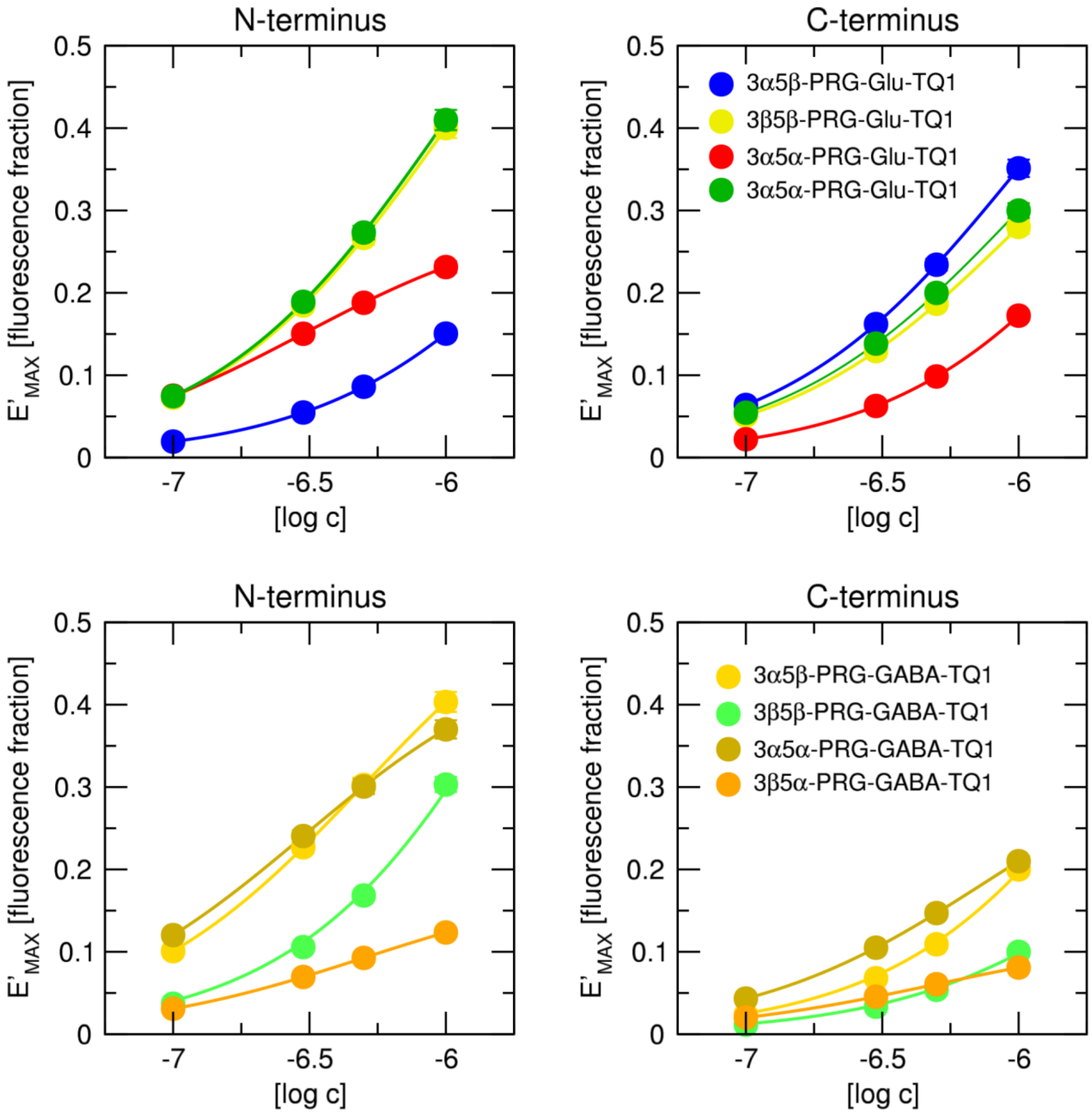


***Figure S1 Dependence of E'$_{MAX}$ of fluorescence quenching on the concentration of the probe.*** *Observed maximal quenching E'MAX at N-terminal site (left) or C-terminal site (right) is plotted against a decadic logarithm of molar concentration of the probe indicated in the legend. Data are means ± SD from 3 independent experiments performed in dodecaplicates. Curves are fits of Equation 1 to the data.*

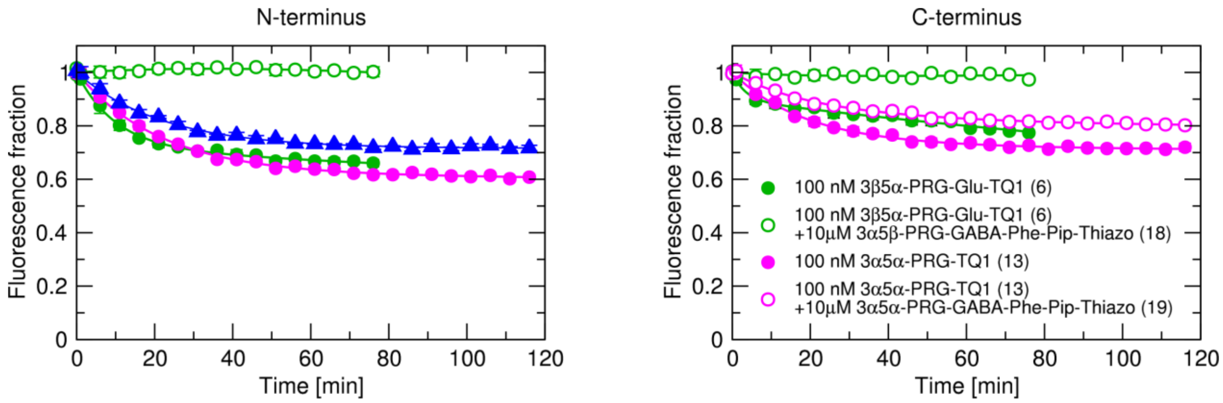


***Figure S2 Competition between quenchers and structurally different non-quenching probes.***
*Time courses (in minutes) of fluorescence quenching of CFP attached to N-terminus (left) and C-terminus (right) of $M_1$-muscarinic receptor expressed as fraction of fluorescence before addition of quencher (EST-256, green, EST-263, magenta) indicated in legend at final 100 nM concentration either in the absence (closed symbols) or presence of 10 µM non-quenching analogue (EST-296, EST-298 or EST-353 open symbols) as indicated in the legend. Data are means ± SD from 3 independent experiments performed in dodecaplicates.*

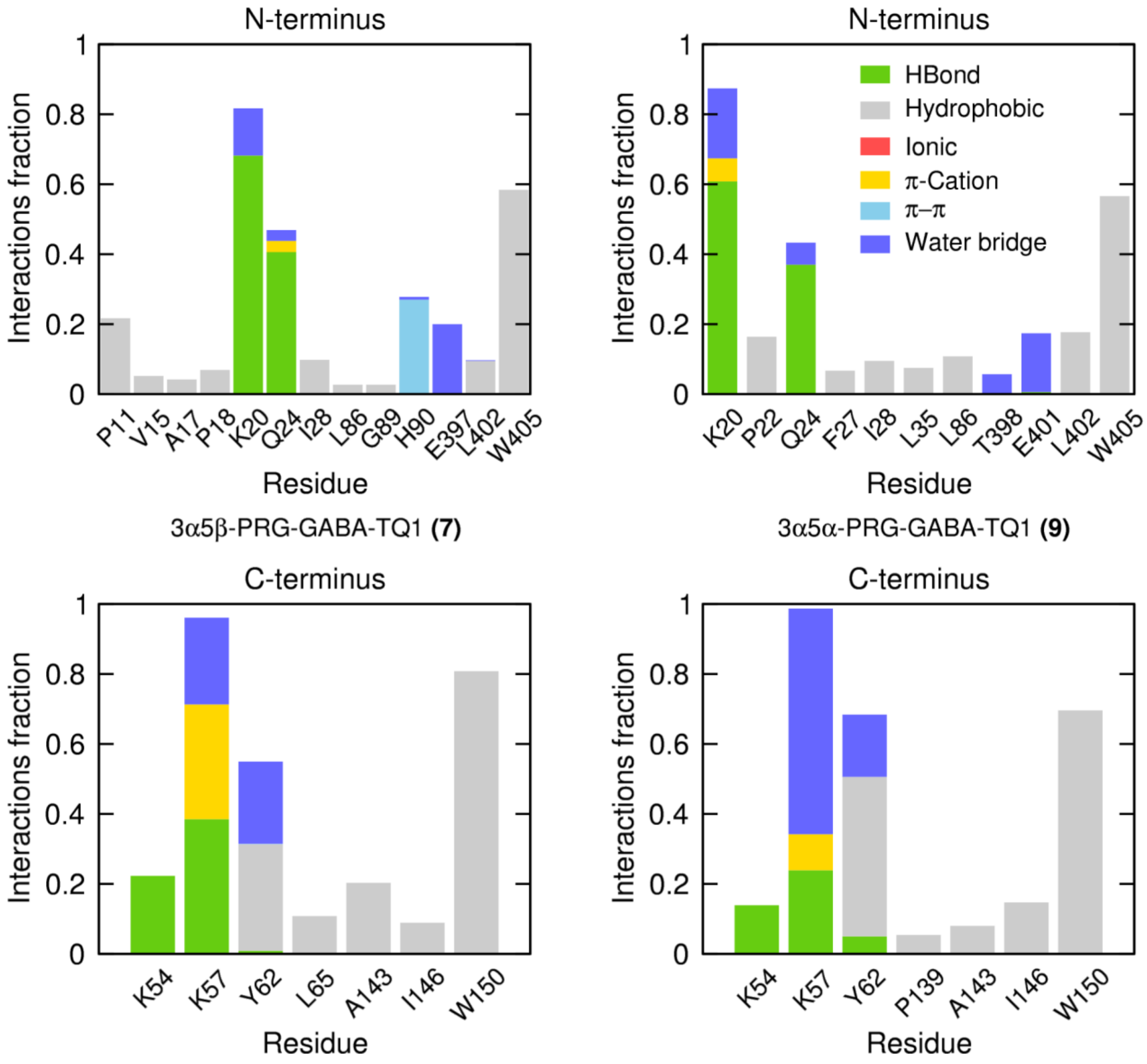


***Figure S3 Interaction of probes with $M_1$ receptor.***

*Histograms of 3α5β-PRG-GABA-TQ1 **(7)** (left), and 3α5α-PRG-GABA-TQ1 (right) interactions with the receptor in the putative N-terminus (top) and C-terminus (bottom) were calculated from MD trajectories using Ligand Interaction Diagram in Maestro. Ligand-receptor interactions are categorised into six types: Hydrogen bonds (green), hydrophobic (grey), ionic (red), π-cation (yellow), π-π (cyan) and water bridges (blue). The stacked bar charts are normalised over the course of the trajectory. Values over 1.0 are possible as some protein residues may make multiple contacts of the same type with the ligand within one time-frame.*

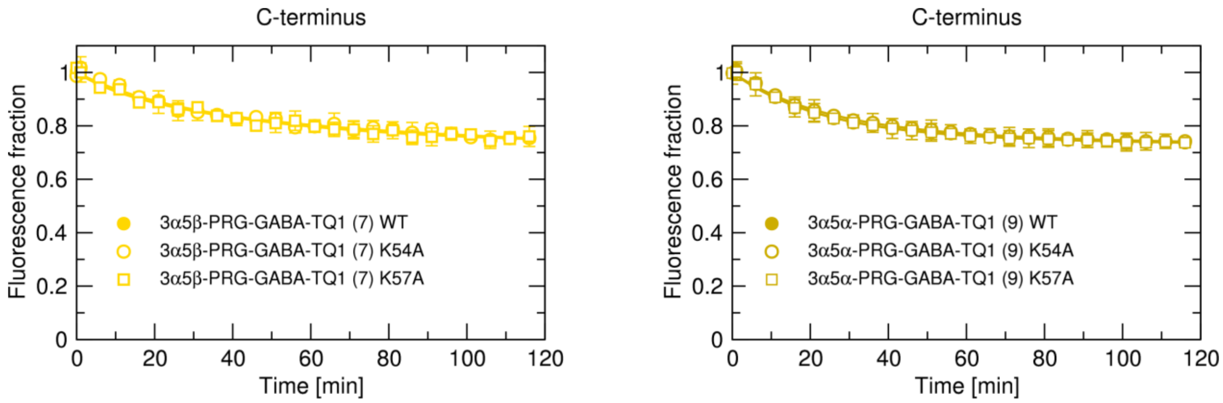


***Figure S4 Lack of effect of mutations on quenching by 3α5β-PRG-GABA-TQ1 (7) and 3α5α-PRG-GABA-TQ1 (9).***

*Time courses (in minutes) of fluorescence quenching of CFP attached to the C-terminal site of $M_1$ muscarinic receptor expressed as a fraction of fluorescence before addition of quenchers **3α5β-PRG-GABA-TQ1 (7)** (left) or **3α5α-PRG-GABA-TQ1 (9)** (right) at the final 1 µM concentration. Quenching was measured at wild-type receptors (WT, closed symbols) or mutated receptors (open symbols). Data are means ± SD from 3 independent experiments performed in dodecaplicates. Curves are fits of two exponential decays to the data.*
