## Supplementary material for "Steroid‑based Tide Quencher 1 probes enable real‑time mapping of novel non‑canonical cholesterol sites on the M1 muscarinic receptor": Supplemntary Information - Chemistry

### 1 Results

#### 1.1 The pregnane skeleton is required for efficient fluorescence quenching

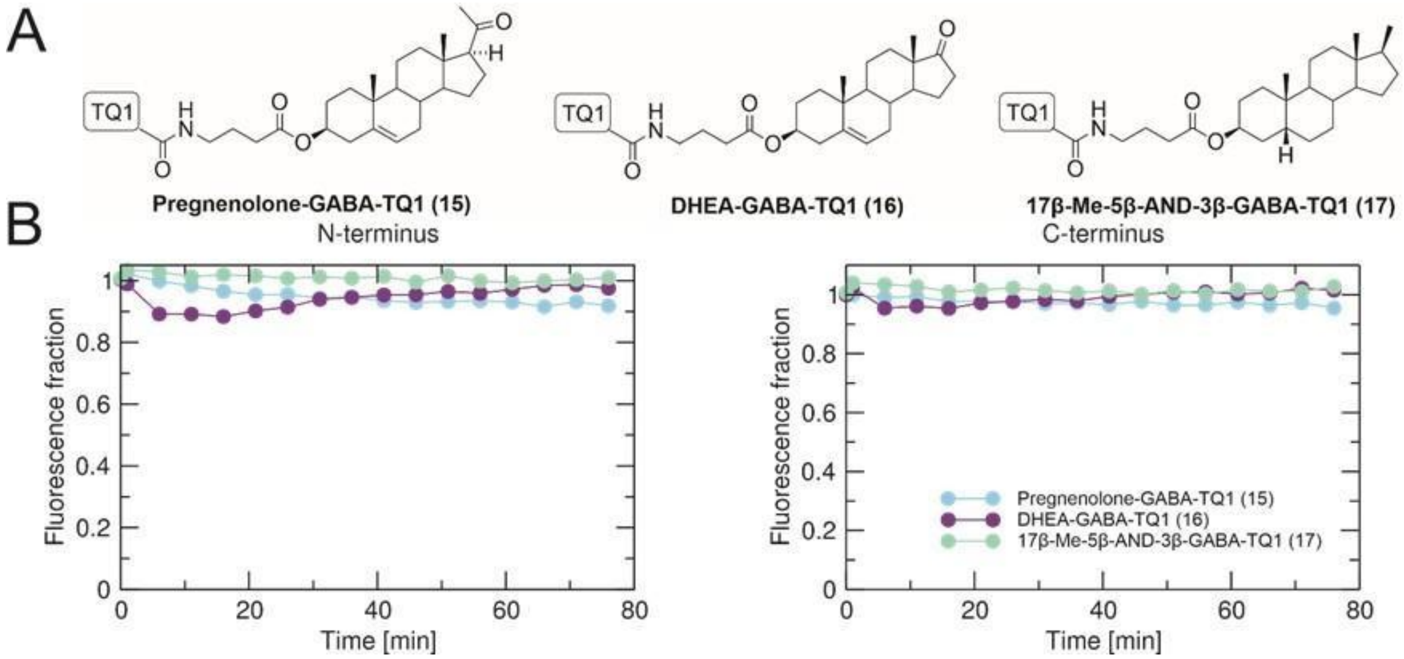

**Figure S1. Fluorescence quenching by γ-aminobutyric acid esters of various steroid skeletons tagged with TQ1.** (A) Structures of TQ1 probes tagged by GABA linker or directly coupled to steroid. (B) Time courses (in minutes) of fluorescence quenching of CFP attached to N terminus (left) or C terminus (right) of M1 muscarinic receptor expressed as a fraction of fluorescence before the addition of quencher indicated in legend at final 1 μM concentration. Data are means from the representative experiment performed in 24 technical replicates confirmed by at least two independent experiments. The relative SD of means was less than 3 %.

#### 1.2 Deciphering of the Tide Quencher 1 structure

The structural characterization of TQ1 acid was performed by 1D ($^{1}$H- and $^{13}$C-NMR) and 2D (H,H-COSY, H,HROESY, H,C-HSQC, H,C-HMBC) NMR spectroscopy, elemental analysis and HR-MS measurements. Also, the $^{1}$H NMR spectrum of TQ1 acid corresponds to a published structure[1] in the literature: $^{1}$H NMR (600 MHz, DMSO-*d*$_6$): δ 12.42 (bs, 1H, COOH), 7.80 (m, 2H, Ar-H meta $N(CH_3)_2$), 7.41 (t, $J$ = 0.8 Hz, 1H, CH-S), 6.88 (m, 2H, Ar-H ortho $N(CH_3)_2$), 3.76 (d, 2H, $J$ = 0.8 Hz, $CH_2$), 3.12 (s, 6H, $N(CH_3)_2$). $^{13}$C NMR (150.9 MHz, DMSO-d6): δ 176.8, 171.6, 154.0, 149.9, 141.7, 126.4 (2C), 117.2, 112.2 (2C), 40.1 (2C), 37.3. Anal. calcd for $C_{13}H_{14}O_2N_4S$: C, 53.78; H, 4.86; N, 19.30; S, 11.04. Found: C, 53.29; H, 4.95; N, 18.93; S, 10.60. MS ESI: *m/z* 245.1 (100%, M – $CO_2$), 289.1 (6%, M - H). HR-MS (ESI) *m/z*: for $C_{13}H_{13}O_2N_4S$ [M - H] calcd, 289.0765; found, 289.0762.

For further confirmation of the structure and the position of the heteroatoms in the heterocyclic ring, reference compounds were used.

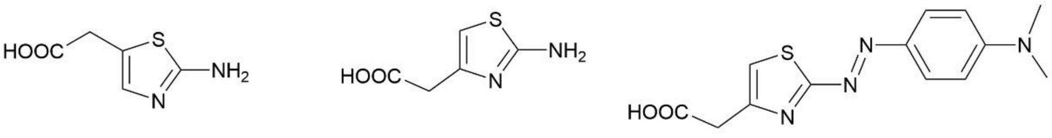

**Reference compound 1** **Reference compound 2** **Tide Quencher 1 acid (TQ1)**

**Figure S2. Structures of reference compound 1 a 2 and Tide Quencher 1 acid.**

**Table S1. Comparison of some carbon-13 chemical shifts (given in ppm) of reference compounds and Tide Quencher 1 acid in DMSO-$d_6$.** The proton chemical shifts are indicated in brackets.

| Compound | -CO- | -$CH_2$- | >C= | -CH= | S-C=N |
|---|---|---|---|---|---|
| 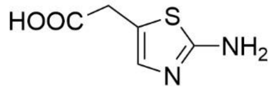  | 172.12 | 32.56 (3.55) | 117.42 | 137.43 (6.70) | 168.73 |
| 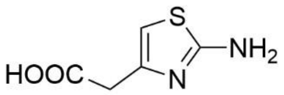  | 171.85 | 37.30 (3.36) | 145.10 | 103.02 (6.28) | 168.26 |
| Tide Quencher 1 acid | 171.62 | 37.28 (3.76) | 149.90 | 117.22 (7.41) | 176.79 |

#### 1.3 Competition experiments between quencher probes and their non-quenching analogues

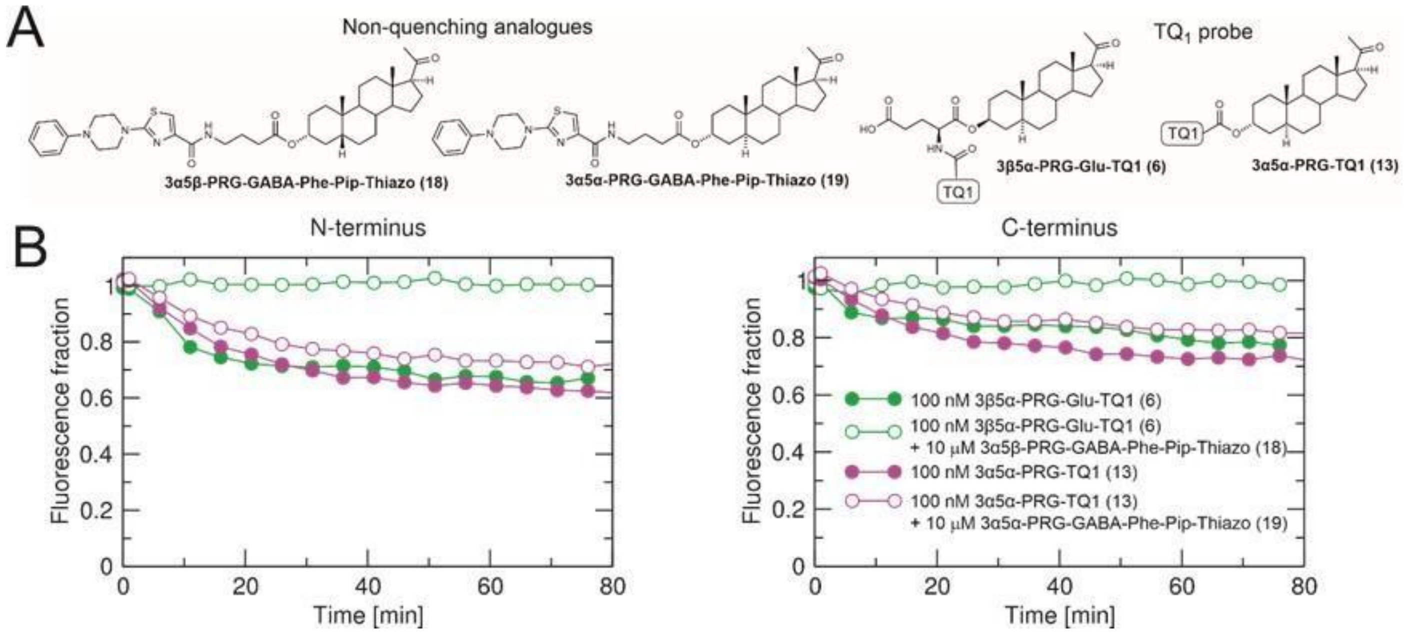

**Figure S3. Competition between quenchers and their non-quenching analogues.**
Time courses (in minutes) of fluorescence quenching of CFP attached to N-terminus (left) and C-terminus (right) of $M_1$-muscarinic receptor expressed as fraction of fluorescence before addition of TQ1-probe (green and purple, closed symbols) at final 100 nM concentration in the presence of 10 μM non-quenching analogue (open symbols). Data are means from the representative experiment performed in dodecaplicates confirmed by at least two independent experiments. The relative SD of means was less than 5 %.

### 2 Chemistry

#### 2.1 Synthesis

##### 2.1.1 Synthesis of PRG-Glu-Dabcyl and PRG-Glu-TQ1 conjugates

The synthesis of **PRG-Glu-Dabcyl** (**1** and **2**) and **PRG-Glu-TQ1** (**3-6**) conjugates (**Scheme S1**) involved the conversion of 3-hydroxysteroids **23-26** into the Boc- and benzyl-protected L-glutamic acid esters **27-30**, followed by palladiumcatalysed hydrogenolysis of the benzyl group and Boc removal with trifluoroacetic acid. This afforded the free amino glutamate ester intermediates **35-38**. The glutamate esters **35-38** were reacted with the corresonding succinimidyl esters to obtain quencher probes **1-6**.

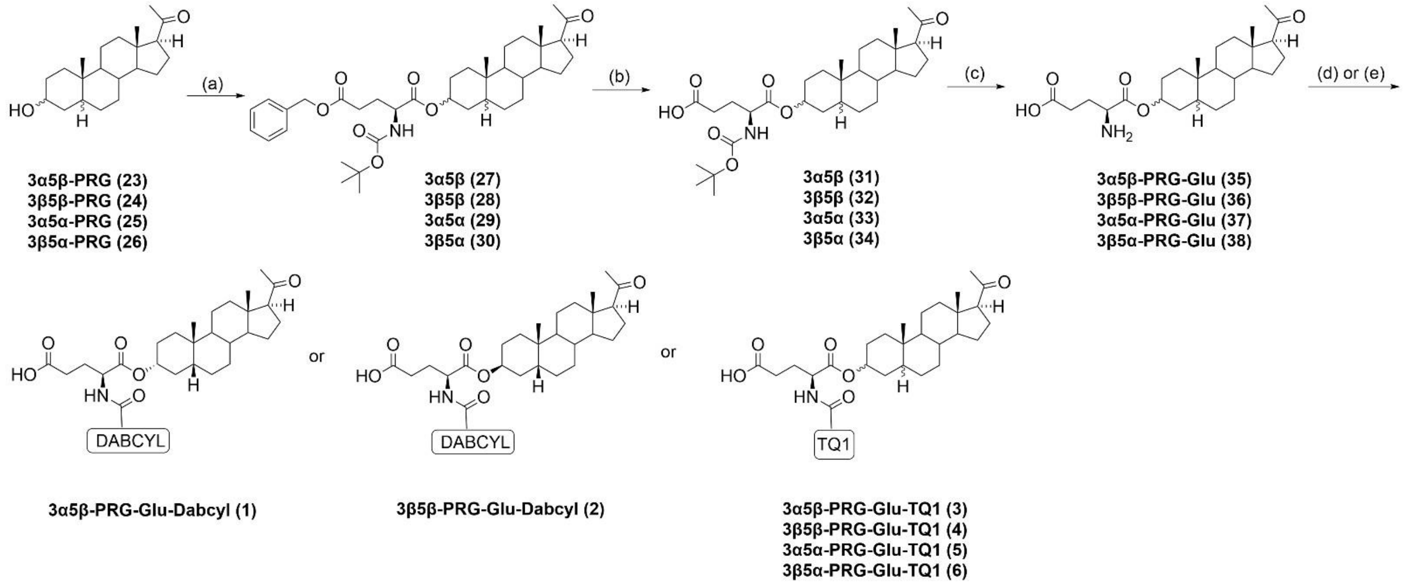

**Scheme S1.** Synthesis of compounds **1-6**. Reagents and conditions: (a) Boc-Glu(OBzl)-OH, EDCI.HCl, DMAP, $CH_2Cl_2$, Ar, 0 °C to rt, overnight; (b) Pd/C (10%), EtOH, 1 atm $H_2$, rt, overnight; (c) TFA, $CH_2Cl_2$, rt, 1h; (d) Dabcyl succinimidyl ester, DIPEA, $CH_2Cl_2$, Ar, rt, overnight, in dark; (e) TQ1 succinimidyl ester, DIPEA, $CH_2Cl_2$, Ar, rt, overnight, in dark.

##### 2.1.2 Synthesis of PRG-GABA-TQ1 conjugates

Steroid probes with GABA linker were synthesized in three consecutive steps (**Scheme S2**). In the first step, 3hydroxysteroids **23-26** were converted into steroid esters **39-42** in the presence of 4-(*tert*-butoxycarbonylamino)butyric acid. It was followed by the removal of Boc protecting group with trifluoroacetic acid or HCl in diethyl ether, leading to amines **43-46**. Lastly, the coupling of steroid amines **43-46** with Tide Quencher 1 succinimidyl ester afforded quencher probes **7-10**.

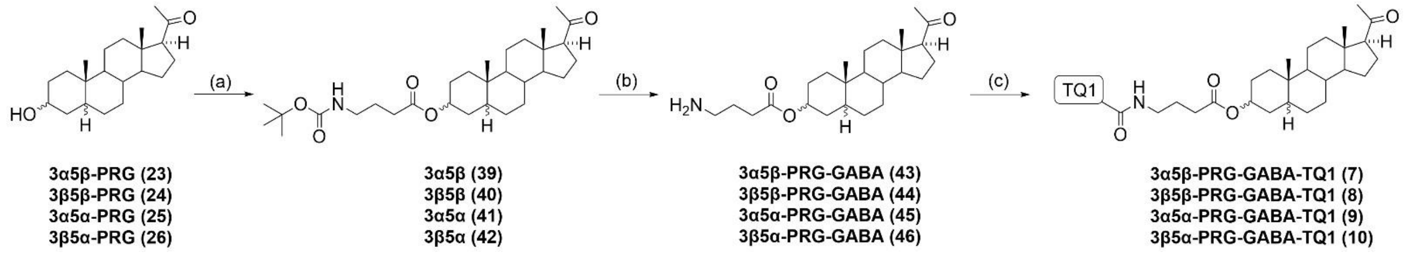

**Scheme S2.** Synthesis of compounds **7-10**. Reagents and conditions: (a) Boc-GABA-OH, EDCI.HCl, DIPEA, DMAP, $CH_2Cl_2$, Ar, rt, overnight; (b) TFA, $CH_2Cl_2$, rt, 3h, for compound **43** HCl in diethyl ether, 24h, Ar; (c) TQ1 succinimidyl ester, DIPEA, $CH_2Cl_2$, Ar, rt, overnight, in dark.

##### 2.1.3 Synthesis of PRG-TQ1 conjugates

To obtain directly coupled quencher probes **11-14**, 3-hydroxysteroids **23-26** were esterified with Tide Quencher 1 acid (**Scheme S3**).

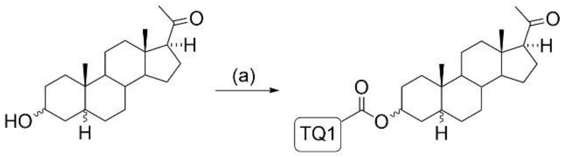

3α5β-PRG (23)
3β5β-PRG (24)
3α5α-PRG (25)
3β5α-PRG (26)

3α5β-PRG-TQ1 (11)
3β5β-PRG-TQ1 (12)
3α5α-PRG-TQ1 (13)
3β5α-PRG-TQ1 (14)

**Scheme S3.** Synthesis of compounds **11-14**. Reagents and conditions: (a) TQ1 acid, EDCI.HCl, DIPEA, DMAP, $CH_2Cl_2$, Ar, rt, overnight, in dark.

##### 2.1.4 Synthesis of pregnenolone-GABA-TQ1 conjugate

To obtain pregnenolone-GABA-TQ1 conjugate **15** (**Scheme S4**), pregnenolone (**47**) was converted to a 4-(*tert*butoxycarbonylamino)butyric acid ester **48**, followed by Boc deprotection with trifluoroacetice acid. The resulting amine **49** was coupled with Tide Quencher succinimidyl ester to afford quencher probe **15**.

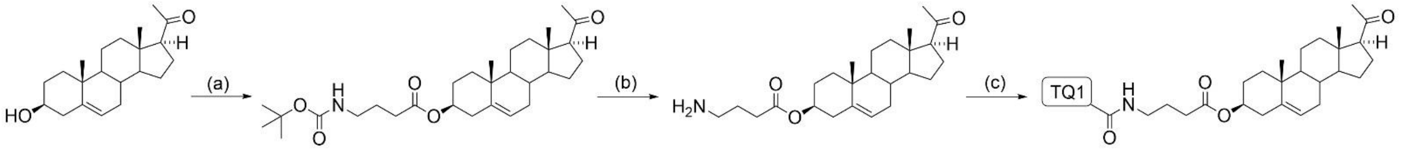

Pregnenolone (47) 48 49 Pregnenolone-GABA-TQ1 (15)

**Scheme S4.** Synthesis of compound **15**. Reagents and conditions: (a) Boc-GABA-OH, EDCI.HCl, DIPEA, DMAP, $CH_2Cl_2$, Ar, rt., overnight; (b) TFA, $CH_2Cl_2$, rt, 3h; (c) TQ1 succinimidyl ester, DIPEA, $CH_2Cl_2$, Ar, rt, overnight, in dark.

##### 2.1.5 Synthesis of DHEA-GABA-TQ1 conjugate

To obtain DHEA-GABA-TQ1 conjugate **16** (**Scheme S5**), dehydroepiandrosterone (**50**) was converted to a 4-(*tert*butoxycarbonylamino)butyric acid ester **51**, followed by Boc deprotection with trifluoroacetice acid. The resulting amine **52** was coupled with Tide Quencher succinimidyl ester to afford quencher probe **16**.

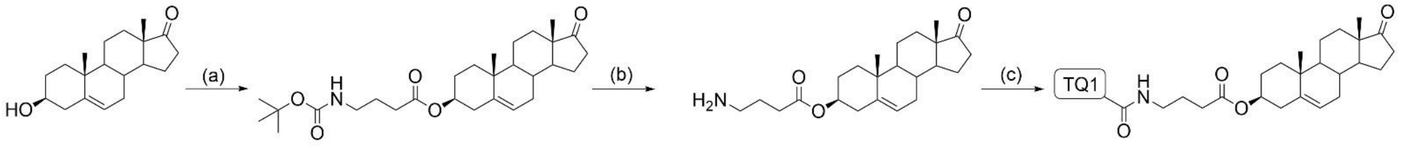

DHEA (50) 51 52 DHEA-GABA-TQ1 (16)

**Scheme S5.** Synthesis of compound **16**. Reagents and conditions: (a) Boc-GABA-OH, EDCI.HCl, DIPEA, DMAP, $CH_2Cl_2$, Ar, rt, overnight; (b) TFA, $CH_2Cl_2$, rt, 1.5h; (c) TQ1 succinimidyl ester, DIPEA, $CH_2Cl_2$, Ar, rt, overnight in dark.

##### 2.1.6 Synthesis of 17β-Me-5β-AND-3β-GABA-TQ1 conjugate

To obtain 17β-Me-5β-AND-3β-GABA-TQ1 conjugate **17** (**Scheme S6**), first, 3-keto steroid **53** was converted to 3βhydroxysteroid **54** in the presence of L-selectride. It was followed by the ester formation of 3β-hydroxysteroid **54** with 4-(*tert*-butoxycarbonylamino)butyric acid affording steroid ester **55**. Then, Boc protecting group was removed in the presence of trifluoroacetic acid. The resulting amine **52** was coupled with Tide Quencher 1 succinimidyl ester to afford quencher probe **17**.

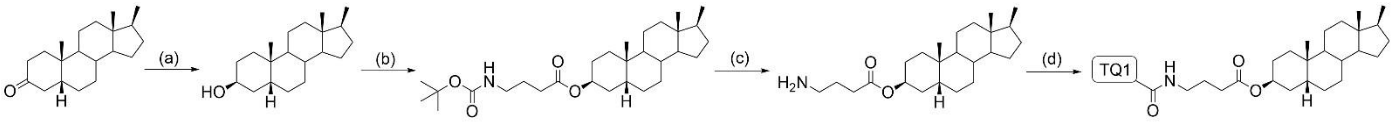

**Scheme S6.** Synthesis of compound **17**. Reagents and conditions: (a) L-selectride, THF, Ar, -78 °C, 1h; (b) Boc-GABAOH, EDCI.HCl, DIPEA, DMAP, $CH_2Cl_2$, Ar, rt, overnight; (c) TFA, $CH_2Cl_2$, rt, 3h; (d) TQ1 succinimidyl ester, DIPEA, $CH_2Cl_2$, Ar, rt, overnight, in dark.

##### 2.1.7 Synthesis of non-quenching analogues

The starting materials of non-quenching analogues with GABA linkers (**Scheme S7**, compounds **43** and **45**) were synthesized in the same manner as described in **Section 2.1.2.** Compounds **43** and **45** were further transformed to nonquenching analogues **18** and **19** by ester formation with 2-(4-phenylpiperazino)-1,3-thiazole-4-carboxylic acid.

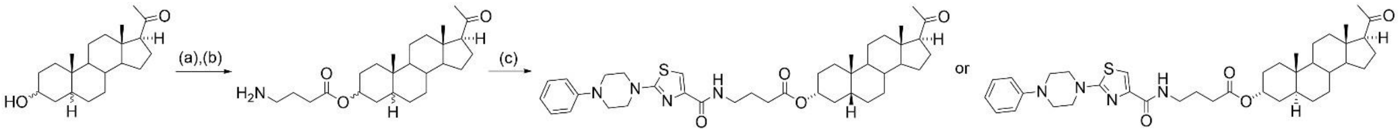

**Scheme S7.** Synthesis of compounds **18-19**. Reagents and conditions: (a) Boc-GABA-OH, EDCI.HCl, DIPEA, DMAP, $CH_2Cl_2$, Ar, rt, overnight; (b) TFA, $CH_2Cl_2$, rt, 3h; (c) 2-(4-phenylpiperazino)-1,3-thiazole-4-carboxylic acid, EDCI.HCl, DIPEA, DMAP, $CH_2Cl_2$, Ar, rt, overnight.

The starting materials of non-quenching analogues with glutamate linkers (**Scheme S8**, compounds **36** and **38**) were synthesized in the same manner as described in **Section 2.1.1.** Compounds **36** and **38** were further transformed to nonquenching analogues **20** and **21** by coupling with Phe-Pip-Thiazo succinimidyl ester (**57**).

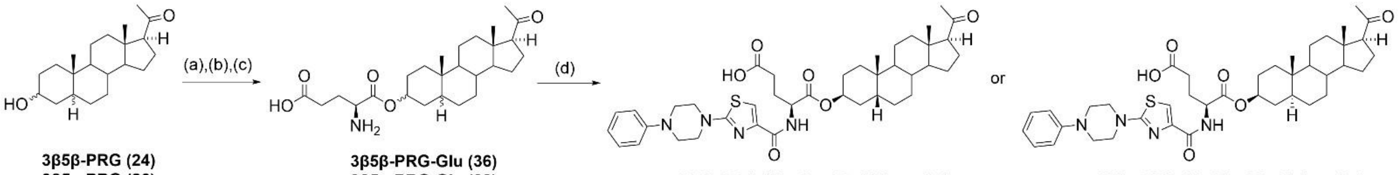

**Scheme S8.** Synthesis of compounds **20-21**. Reagents and conditions: (a) Boc-Glu(OBzl)-OH, EDCI.HCl, DMAP, $CH_2Cl_2$, Ar, 0 °C to rt, overnight; (b) Pd/C (10%), EtOH, 1 atm $H_2$, rt, overnight; (c) TFA, $CH_2Cl_2$, rt, 1h; (d) Phe-PipThiazo succinimidyl ester (**57**), DIPEA, $CH_2Cl_2$, Ar, rt, overnight.

To obtain non-quenching analogue **22**, 3-hydroxysteroid **25** was esterified with 2-(4-phenylpiperazino)-1,3-thiazole-4carboxylic acid (**Scheme S9**).

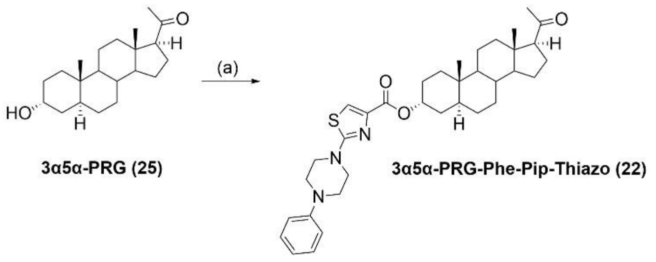

**Scheme S9.** Synthesis of compound **22**. Reagents and conditions: (a) 2-(4-phenylpiperazino)-1,3-thiazole-4-carboxylic acid, EDCI.HCl, DIPEA, DMAP, $CH_2Cl_2$, Ar, rt, overnight.

#### 2.2 Materials and methods

For the optical rotation measurements, an AUTOPOL IV (Rudolph Research Analytical, Hacketts-town, NJ) was used. All samples were measured at 20 °C at a given concentration in a given solvent at 589 nm.

The $^{1}$H and $^{13}$C NMR spectra were measured on a Bruker Avance III HD 400 instrument (400 MHz for $^{1}$H and 101 MHz for $^{13}$C) and on a Bruker Avance III 600 MHz spectrometer at 25 °C (600 MHz for $^{1}$H and 150.9 MHz for $^{13}$C). The $^{13}$C NMR spectra of compounds **29** and **30** were recorded on a JEOL ECZR 500 MHz spectrometer (126 MHz for $^{13}$C). Chemical shifts are given in ppm (δ) and are referenced to the solvent signal ($CDCl_3$: δ 7.26 for $^{1}$H NMR and δ 77.16 for $^{13}$C NMR, DMSO-$d_6$: δ 2.50 for $^{1}$H NMR and 39.70 for $^{13}$C NMR, $CD_3OD$: δ 3.31 for $^{1}$H NMR and δ 49.00 for $^{13}$C NMR). The coupling constants *J* are given in Hz. Structures of selected starting materials, quencher probes and nonquenching analogues were established by 1D ($^{1}$H- and $^{13}$C-NMR) and 2D (H,H-COSY, H,H-ROESY, H,C-HSQC, H,CHMBC) NMR spectroscopy.

The purity of final compounds **1-19** was measured using LC-MS system equipped with UV-VIS detector. Equipment: TSP quaternary pump P4000, TSP autosampler AS3000, UV/VIS detector UV6000LP and ion-trap mass spectrometer Advantage. Solvent A was 98% water / 2% acetonitrile and B was 95% acetonitrile / 3% isopropanol/ 2% water with 5 mM ammonium formate both. UV6000LP is diode array detector witch 512 diodes. UV/VIS setup: wavelength range 190-700 nm, bandwith 1 nm, step 1 nm and scan rate 1 Hz. Ions have been detected in positive ion mode with m/z range from 250 to 1500 Da. Gradient setup: 0-25-30-30.1-45 min 50-100-100-50-50% solvent B and flow rate 150 uL/min. For separation has been used C4 column: Phenomenex 250 mm length, 2 mm width, Jupiter 300-5 C4.
The analytical HPLC measurements of compounds **20-22** were performed on Shimadzu (Tokyo, Japan) Nexera LC-40 equipped with an SCL-40 communication module, two degasser units DGU-403 and DGU-405, two LC-40 D X3 dual solvent delivery modules, an autosampler, and a thermostated column oven CTO-40S. An ELSD-LT II, evaporative light scattering detector (ELSD) (Shimadzu, Japan) was utilized for the analysis of the purity. The conditions for the analysis were 60 °C nebulization temperature by nitrogen and 10 the gain factor.
Mass spectrometric detection was performed on a single quadrupole LC-MS-2020 (Shimadzu, Japan) with dual ion source by an electrospray ionization (ESI) probe, and simultaneously ionized by an atmospheric pressure chemical ionization (APCI) corona discharge needle located below the ESI outlet. All acquisitions were performed in both positive and negative ionization modes. The nebulizing gas had a flow of 1.5 L/min, the drying gas was set to 15 L/min, the corona needle voltage was 4500 V (for positive mode), and the heat block had a temperature of 400°C. Full scan mass spectra were acquired from m/z 200-800. The data was processed by software LabSolutions™.

FTIR spectra were recorded on a Nicolet 6700 spectrometer (Thermo Scientific, USA). The HRMS spectra were performed with an LTQ Orbitrap XL instrument (Thermo Fischer Scientific, Waltham, MA) using electrospray ionization (ESI). For elemental analysis, a PE 2400 Series II CHNS/O Analyzer (PerkinElmer, Waltham, MA) was used with a microbalance MX5 (Mettler Toledo, Switzerland).

TLC was performed on Silica gel 60 F254-coated aluminium sheets (VWR). The TLC plates were visualized by exposure to ultraviolet light, then the spots were stained by chemical reagent: the TLC plate was immersed into methanol/sulfuric acid (10% v/v) mixture, then was heated briefly to 200 °C with a heat gun. Flash chromatography was performed on a puriFlash 5.250 instrument (Interchim, Montluçon, France) using neutral silica gel (Merck, 40–63 μm) and an ELSD detector. All commercially available solvents and reagents were used as received.

*N*,*N*-Diisopropylethylamine (DIPEA), 4-dimethylaminopyridin (DMAP), *N*-(3-dimethylaminopropyl)-*N'*ethylcarbodiimide hydrochloride (EDCI.HCl), Boc-*L*-glutamic acid 5-benzyl ester (Boc-Glu(OBzl)-OH), 4-(*tert*butoxycarbonylamino)butyric acid (Boc-GABA-OH), L-selectride (1M in THF), Pd/C (10 wt%), HCl in diethyl ether (1 M), trifluoroacetic acid, succinic acid and 4-[4-(dimethylamino)phenylazo]benzoic acid *N*-succinimidyl ester (Dabcyl-OSu) were purchased from Sigma-Aldrich. 2-(4-Phenylpiperazino)-1,3-thiazole-4-carboxylic acid is a Key Organics Limited product. Tide Quencher 1 succinimidyl ester (TQ1-SE) and Tide Quencher 1 acid (TQ1 acid) were purchased from Scintila, s.r.o., Jihlava. Reference compounds for the characterization of TQ1 acid, 2-(2-aminothiazol5-yl)acetic acid and 2-amino-4-thiazoleacetic acid were Asta Tech Inc. and Sigma Aldrich products, respectively. Dichloromethane was distilled from $P_2O_5$ under an inert atmosphere before use.

#### 2.3 Synthetic procedures

##### 2.3.1 Synthesis of steroid-DABCYL and steroid-TQ1 conjugates via a glutamate linker

###### 2.3.1.1 General procedure for the Boc- and benzyl-protected *L*-glutamic acid ester formation

DMAP (0.15 eq., 0.15 mmol, 18 mg) and Boc-Glu(OBzl)-OH (1.1 eq., 1.1 mmol, 371 mg) were placed in a roundbottom flask equipped with a septum inlet. The flask was evacuated and refilled with argon two times. It was followed by the addition of freshly distilled dichloromethane (9 ml) and the mixture was stirred while all solids were dissolved. Then, the solution was cooled to 0°C in an ice bath and EDCI.HCl (1 eq., 1 mmol, 192 mg) was added under an argon stream. Steroid alcohol **3α5α-PRG** (**25**) or **3β5α-PRG** (**26**) (1 mmol, 319 mg) was dissolved separately in 7 ml dichloromethane under argon atmosphere, and the solution was added dropwise to the reaction mixture at 0°C. The reaction mixture was warmed to room temperature upon stirring and further stirred at room temperature overnight. After 24 hours, starting material was still present, therefore, Boc-Glu(OBzl)-OH (0.3 eq., 0.3 mmol, 101 mg) and EDCI.HCl (0.4 eq., 0.4 mmol, 77 mg) were added. The reaction was complete after additional 24 hours.
In the case of **3β5β-PRG** (**24**), using higher excess of Boc-Glu(OBzl)-OH (1.4 eq., 1.4 mmol, 472 mg) and EDCI.HCl (1.4 eq., 1.4 mmol, 268 mg), the reaction was complete after 24 hours. Then, the solvent was evaporated in vacuo. The crude material was re-dissolved in dichloromethane (70 ml), washed with saturated $NaHCO_3$ solution (25 ml), water (25 ml) and brine (25 ml). The organic phase was dried over anhydrous $Na_2SO_4$. It was filtered off and the solvent was removed in vacuo. The crude material was purified by flash chromatography on silica gel using a gradient of petroleum ether/ethyl-acetate (5% to 50% in 20 column volumes) gave products **28** (546 mg, 86%), **29** (505 mg, 79%), **30** (578 mg, 91%), as a viscous oil.

**1-((3*R*,5*R*,8*R*,9*S*,10*S*,13*S*,14*S*,17*S*)-17-Acetyl-10,13-dimethylhexadecahydro-1*H*-cyclopenta[*a*]phenanthren-3-yl) 5-benzyl (tert-butoxycarbonyl)-*L*-glutamate (27)**

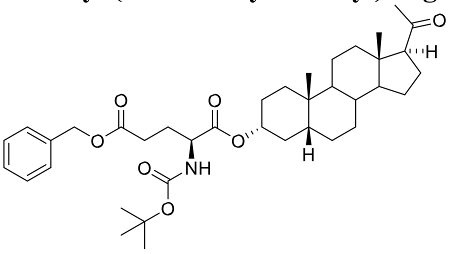

Compound **27** was synthesized according to the previously reported procedure.[2]

**1-((3*S*,5*R*,8*R*,9*S*,10*S*,13*S*,14*S*,17*S*)-17-Acetyl-10,13-dimethylhexadecahydro-1*H*-cyclopenta[*a*]phenanthren-3-yl) 5-benzyl (*tert*-butoxycarbonyl)-*L*-glutamate (28)**

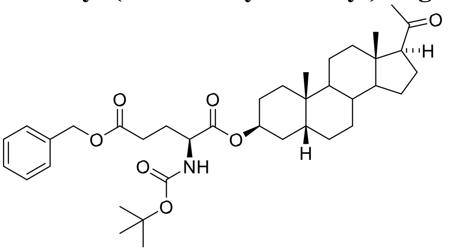

$[\alpha]_D$ +58.2, c 0.295, $CHCl_3$. $^1H$ NMR (400 MHz, $CDCl_3$): $\delta$ 7.41-7.29 (m, 5H, $C_6H_5$), 5.18-5.05 (m, 4H, H-3, NH, $OCH_2$-Ph), 4.38-4.27 (m, 1H, C*H*-NH), 2.53 (t, $J$ = 9.3 Hz, 1H, H-17), 2.11 (s, 3H, H-21), 1.43 (s, 9H, $(CH_3)_3$C-O), 0.95 (s, 3H, H-19), 0.60 (s, 3H, H-18). $^{13}C$ NMR (101 MHz, $CDCl_3$): $\delta$ 209.8, 172.7, 171.7, 155.5, 135.9, 128.7 (2C), 128.4 (3C), 80.1, 72.5, 66.6, 64.0, 56.9, 53.2, 44.5, 40.1, 39.4, 37.5, 35.8, 35.0, 31.7, 30.9, 30.7, 30.5, 28.5 (3C), 28.1, 26.5, 26.3, 25.1, 24.6, 23.9, 23.0, 21.2, 13.6. IR ($CHCl_3$, $\nu$($cm^{-1}$)): 3435 (NH), 2979 ($CH_3$), 2939 ($CH_3$), 2876 ($CH_2$), 1729 (C=O), 1704 (C=O), 1171 (C-O, Boc). MS ESI: m/z 660.4 (100%, M + Na). HR-MS (ESI) m/z: for $C_{38}H_{56}O_7N$ [M + H] calcd, 638.4051; found, 638.4048. **1-((3*R*,5*S*,8*R*,9*S*,10*S*,13*S*,14*S*,17*S*)-17-Acetyl-10,13-dimethylhexadecahydro-1*H*-cyclopenta[*a*]phenanthren-3-yl) 5-benzyl (*tert*-butoxycarbonyl)-*L*-glutamate (29)**

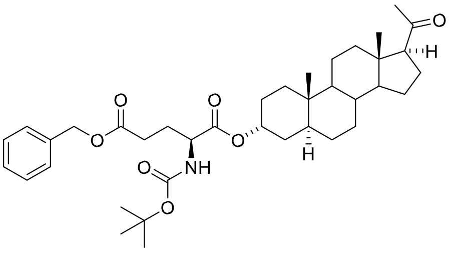

[α]D +62.3, c 0.239, $CHCl_3$. [1]H NMR (400 MHz, $CDCl_3$): *δ* 7.40-7.29 (m, 5H, $C_6H_5$), 5.18-5.09 (m, 3H, NH, O-$CH_2$Ph), 5.07 (p, *J* = 2.7 Hz, 1H, H-3), 4.39-4.28 (m, 1H, C*H*-NH), 2.11 (s, 3H, H-21), 1.44 (s, 9H, $(CH_3)_3$C-O), 0.78 (s, 3H, H-19), 0.60 (s, 3H, H-18). $^{13}$C NMR (126 MHz, $CDCl_3$): *δ* 209.9, 172.8, 171.8, 155.6, 135.9, 128.7 (2C), 128.4, 128.3 (2C), 80.1, 71.8, 66.6, 64.0, 56.8, 54.1, 53.1, 44.4, 40.1, 39.1, 35.9, 35.5, 33.0, 32.8, 31.9, 31.7, 30.4, 28.5 (3C), 28.3, 28.2, 26.2, 24.5, 22.9, 20.9, 13.6, 11.4. IR ($CHCl_3$, *ν*($cm^{-1}$)): 3434 (NH), 2971 ($CH_3$), 2936 ($CH_2$), 1730 (C=O), 1704 (C=O), 1162 (C-O, Boc). MS ESI: m/z 660.4 (100%, M + Na). HR-MS (ESI) m/z: for $C_{38}H_{55}O_7NNa$ [M + Na] calcd, 660.3871; found, 660.3870.

**1-((3*S*,5*S*,8*R*,9*S*,10*S*,13*S*,14*S*,17*S*)-17-Acetyl-10,13-dimethylhexadecahydro-1*H*-cyclopenta[*a*]phenanthren-3-yl) 5-benzyl (*tert*-butoxycarbonyl)-*L*-glutamate (30)**

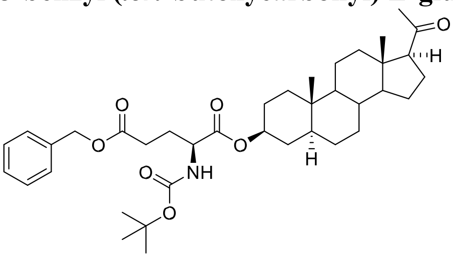

[α]D +48.2, c 0.226, $CHCl_3$. [1]H NMR (400 MHz, $CDCl_3$): *δ* 7.39-7.29 (m, 5H, $C_6H_5$), 5.15-5.04 (m, 3H, NH, O-$CH_2$Ph), 4.73 (tt, *J* = 11.4 Hz, 4.9 Hz, 1H, H-3), 4.34-4.22 (m, 1H, C*H*-NH), 2.52 (t, *J* = 9.1 Hz, 1H, H-17), 2.11 (s, 3H, H21), 1.43 (s, 9H, $(CH_3)_3$C-O), 0.81 (s, 3H, H-19), 0.60 (s, 3H, H-18). $^{13}$C NMR (126 MHz, $CDCl_3$): *δ* 209.8, 172.7, 171.8, 155.5, 135.9, 128.7 (2C), 128.4 (3C), 80.1, 75.2, 66.6, 63.9, 56.7, 54.2, 53.1, 44.7, 44.4, 39.1, 36.8, 35.6, 35.6, 33.9, 32.0, 31.7, 30.5, 28.5, 28.4 (3C), 28.1, 27.5, 24.5, 22.9, 21.3, 13.6, 12.3. IR ($CHCl_3$, *ν*($cm^{-1}$)): 3436 (NH), 2972 ($CH_3$), 2937 ($CH_2$), 1730 (C=O), 1704 (C=O), 1167 (C-O, Boc). MS ESI: m/z 660.4 (100%, M + Na). HR-MS (ESI) m/z: for $C_{38}H_{55}O_7NNa$ [M + Na] calcd, 660.3871; found, 660.3866.

##### 2.3.1.2 General procedure for the deprotection of benzyl group

In a round-bottom flask equipped with an adapter with a two-way tap, 1 mmol steroid was dissolved in 42 ml EtOH. Then, Pd/C (10 wt%, 63 mg) was added under an argon atmosphere. The equipment was carefully evacuated and refilled with hydrogen gas (three short evacuation and refill cycles) with the help of a gas balloon. The mixture was stirred at room temperature overnight. After completion of the reaction, the equipment was evacuated and refilled with argon three times. The catalyst was filtered off on a celite/silica pad and was washed with a small amount of ethyl acetate. The solvent was evaporated.

**(*S*)-5-(((3*R*,5*R*,8*R*,9*S*,10*S*,13*S*,14*S*,17*S*)-17-Acetyl-10,13-dimethylhexadecahydro-1*H*-cyclopenta[*a*]phenanthren3-yl)oxy)-4-((*tert*-butoxycarbonyl)amino)-5-oxopentanoic acid (31)**

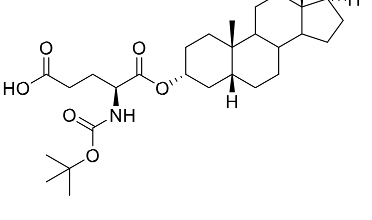

Compound **31** was synthesized according to the previously reported procedure.[2]

**(*S*)-5-(((3*S*,5*R*,8*R*,9*S*,10*S*,13*S*,14*S*,17*S*)-17-Acetyl-10,13-dimethylhexadecahydro-1*H*-cyclopenta[*a*]phenanthren3-yl)oxy)-4-((*tert*-butoxycarbonyl)amino)-5-oxopentanoic acid (32)**

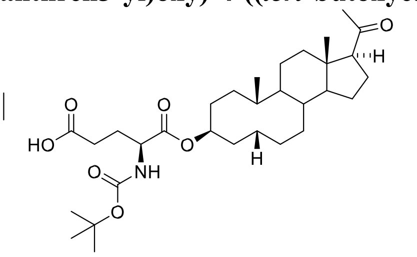

Starting from **28** (0.677 mmol, 432 mg), compound **32** was synthesized according to general procedure and isolated as amorphous solid (370 mg, 99%).

[α]D +65.8, c 0.316, $CHCl_3$. $^1H$ NMR (400 MHz, $CDCl_3$): $\delta$ 5.24-5.09 (m, 2H, overlapping signals of H-3 and NH), 4.40-4.28 (m, 1H, C*H*-NH), 2.53 (t, *J* = 9.0 Hz, 1H, H-17), 2.11 (s, 3H, H-21), 1.44 (s, 9H, $(CH_3)_3C$-O), 0.97 (s, 3H, H19), 0.60 (s, 3H, H-18). $^{13}C$ NMR (101 MHz, $CDCl_3$): $\delta$ 209.9, 177.1, 171.7, 155.8, 80.4, 72.7, 64.0, 56.9, 53.0, 44.5, 40.1, 39.4, 37.5, 35.8, 35.0, 31.7, 30.8, 30.7, 30.2, 28.4 (3C), 28.2, 26.5, 26.3, 25.1, 24.6, 23.9, 23.0, 21.2, 13.6. IR ($CHCl_3$, $\nu$($cm^{-1}$)): 3514 (OH), 3434 (NH), 2978 ($CH_3$), 2939 ($CH_2$), 2876 ($CH_3$), 1731 (C=O), 1711 (C=O), 1174 (C-O, Boc). MS ESI: m/z 570.3 (100%, M + Na). HR-MS (ESI) m/z: for $C_{31}H_{49}O_7NNa$ [M + Na] calcd, 570.3401; found, 570.3398.

**(*S*)-5-(((3*R*,5*S*,8*R*,9*S*,10*S*,13*S*,14*S*,17*S*)-17-Acetyl-10,13-dimethylhexadecahydro-1*H*-cyclopenta[*a*]phenanthren3-yl)oxy)-4-((*tert*-butoxycarbonyl)amino)-5-oxopentanoic acid (33)**

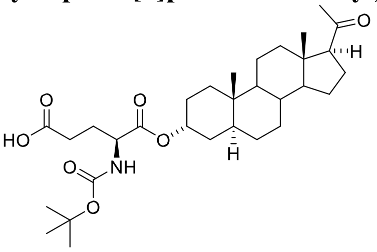

Starting from **29** (0.561 mmol, 358 mg), compound **33** was synthesized according to general procedure and isolated as amorphous solid (293 mg, 95%).

[α]D +72.6, c 0.215, $CHCl_3$. $^1H$ NMR (400 MHz, $CDCl_3$): $\delta$ 5.23 (d, *J* = 8.1 Hz, 1H, NH), 5.12-5.05 (m, 1H, H-3), 4.414.30 (m, 1H, C*H*-NH), 2.53 (t, *J* = 8.5 Hz, 1H, H-17), 2.12 (s, 3H, H-21), 1.45 (s, 9H, $(CH_3)_3C$-O), 0.79 (s, 3H, H-19),

0.60 (s, 3H, H-18). $^{13}C$ NMR (101 MHz, $CDCl_3$): $\delta$ 210.0, 176.7, 171.6, 155.9, 80.4, 72.1, 64.0, 56.8, 54.1, 53.0, 44.4, 40.1, 39.1, 35.9, 35.6, 33.0, 32.9, 31.9, 31.7, 30.2, 28.5 (3C), 28.3, 26.2, 24.5, 23.0, 20.9, 13.6, 11.5. 1 carbon could not be detected. IR ($CHCl_3$, $\nu$($cm^{-1}$)): 3516 (OH), 3434 (NH), 2972 ($CH_3$), 2935 ($CH_2$), 1730 (C=O), 1711 (C=O), 1162 (CO, Boc). MS ESI: m/z 546.3 (100%, M - H). HR-MS (ESI) m/z: for $C_{31}H_{49}O_7NNa$ [M + Na] calcd, 570.3401; found, 570.3398.

**(*S*)-5-(((3*S*,5*S*,8*R*,9*S*,10*S*,13*S*,14*S*,17*S*)-17-Acetyl-10,13-dimethylhexadecahydro-1*H*-cyclopenta[*a*]phenanthren3-yl)oxy)-4-((*tert*-butoxycarbonyl)amino)-5-oxopentanoic acid (34)**

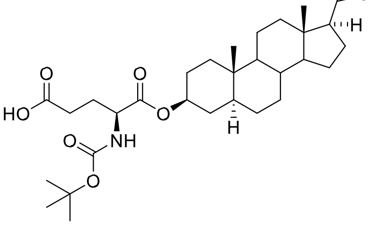

Starting from **30** (0.688 mmol, 439 mg), compound **34** was synthesized according to general procedure and isolated as amorphous solid (373 mg, 99%).

[α]D +51.4, c 0.218, $CHCl_3$. $^1H$ NMR (400 MHz, $CDCl_3$): $\delta$ 5.17 (d, *J* = 8.2 Hz, 1H, NH), 4.74 (tt, *J* = 11.4 Hz, 4.9 Hz, 1H, H-3), 4.35-4.22 (m, 1H, C*H*-NH), 2.52 (t, *J* = 8.8 Hz, 1H, H-17), 2.11 (s, 3H, H-21), 1.44 (s, 9H, $(CH_3)_3C$-O), 0.82

(s, 3H, H-19), 0.60 (s, 3H, H-18). $^{13}$C NMR (101 MHz, $CDCl_3$): $\delta$ 209.9, 176.9, 171.7, 155.8, 80.4, 75.3, 64.0, 56.8, 54.2, 53.0, 44.8, 44.4, 39.1, 36.8, 35.6, 35.6, 33.9, 32.0, 31.7, 30.2, 28.6, 28.4 (3C), 28.3, 27.5, 24.5, 22.9, 21.3, 13.6, 12.4. IR ($CHCl_3$, $\nu$(cm$^{-1}$)): 3516 (OH), 3434 (NH), 2972 ($CH_3$), 2937 ($CH_2$), 1731 (C=O), 1711 (C=O), 1161 (C-O, Boc). MS ESI: m/z 546.3 (100%, M - H). HR-MS (ESI) m/z: for $C_{31}H_{49}O_7NNa$ [M + Na] calcd, 570.3401; found, 570.3400.

##### 2.3.1.3 General procedure for the deprotection of Boc group

In a round-bottom flask, 0.2 mmol steroid was dissolved in 3 ml dichloromethane, followed by the addition of 1.4 ml trifluoroacetic acid (1.2-times diluted with dichloromethane). The reaction mixture was stirred at room temperature for 1 hour. Dichloromethane was evaporated, the residue of trifluoroacetic acid was flushed out in a nitrogen stream. The residue was dissolved in 1 ml pyridine and it was added dropwise to 60 ml ice-cold water. Precipitate was formed, and it was kept at 4°C in fridge overnight. The precipitate was filtered off and was washed with ice-cold water (3 x 2 ml). The obtained material was dried by suction and then in a high vacuum at 40°C.

**(*S*)-5-(((3*R*,5*R*,8*R*,9*S*,10*S*,13*S*,14*S*,17*S*)-17-Acetyl-10,13-dimethylhexadecahydro-1*H*-cyclopenta[*a*]phenanthren3-yl)oxy)-4-amino-5-oxopentanoic acid (35)**

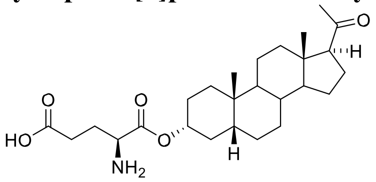

Compound **35** was synthesized according to the previously reported procedure.[2] The $^{1}$H and $^{13}$C NMR chemical shifts and corresponding signal assignments are reported in **Table S2** and **Table S3**, respectively.

**(*S*)-5-(((3*S*,5*R*,8*R*,9*S*,10*S*,13*S*,14*S*,17*S*)-17-Acetyl-10,13-dimethylhexadecahydro-1*H*-cyclopenta[*a*]phenanthren3-yl)oxy)-4-amino-5-oxopentanoic acid (36)**

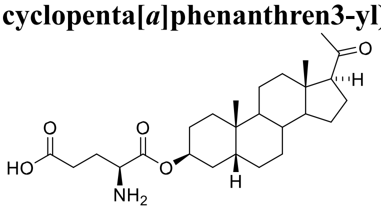

Starting from **32** (0.257 mmol, 141 mg), compound **36** was synthesized according to general procedure and isolated as white solid (89 mg, 77%).

The $^{1}$H and $^{13}$C NMR chemical shifts and corresponding signal assignments are reported in **Table S2** and **Table S3**, respectively. Due to limited solubility in DMSO and MeOH, $[\alpha]_D$ was not measured. IR (KBr, $\nu$(cm$^{-1}$)): 2932 ($CH_2$), 2874 ($CH_3$), 2660 ($NH_3^+$), 2050 (indicator band, $NH_3^+$), 1739 (C=O), 1705 (C=O), 1580 ($NH_3^+$,$COO^-$), 1227 (C-O), zwitterionic compound. MS ESI: m/z 446.3 (100%, M - H). HR-MS (ESI) m/z: for $C_{26}H_{40}O_5N$ [M - H] calcd, 446.2912; found, 446.2908.

**(*S*)-5-(((3*R*,5*S*,8*R*,9*S*,10*S*,13*S*,14*S*,17*S*)-17-Acetyl-10,13-dimethylhexadecahydro-1*H*-cyclopenta[*a*]phenanthren3-yl)oxy)-4-amino-5-oxopentanoic acid (37)**

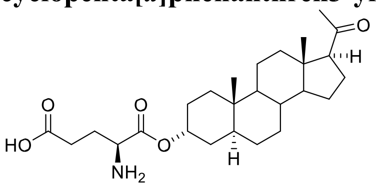

Starting from **33** (0.1095 mmol, 60 mg), compound **37** was synthesized according to general procedure and isolated as white solid (42 mg, 86%).

The $^{1}$H and $^{13}$C NMR chemical shifts and corresponding signal assignments are reported in **Table S2** and **Table S3**, respectively. $[\alpha]_D$ +73.9, c 0.165, MeOH. IR (MeOH, film, $\nu$(cm$^{-1}$)): 2966 ($CH_3$), 2926 ($CH_2$), 2687 ($NH_3^+$), 2163 (indicator band, $NH_3^+$) 1744 (C=O), 1712 (C=O), 1605 ($NH_3^+$), 1576 ($COO^-$), 1387 ($CH_3$), zwitterionic compound. MS ESI: m/z 446.3 (100%, M - H). HR-MS (ESI) m/z: for $C_{26}H_{40}O_5N$ [M - H] calcd, 446.2912; found, 446.2908. **(*S*)-5-(((3*S*,5*S*,8*R*,9*S*,10*S*,13*S*,14*S*,17*S*)-17-Acetyl-10,13-dimethylhexadecahydro-1*H*-cyclopenta[*a*]phenanthren3-yl)oxy)-4-amino-5-oxopentanoic acid (38)**

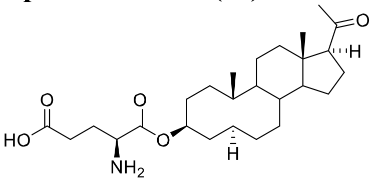

Starting from **34** (0.482 mmol, 264 mg), compound **38** was synthesized according to general procedure and isolated as white solid (175 mg, 81%).

The $^1H$ and $^{13}C$ NMR chemical shifts and corresponding signal assignments are reported in **Table S2** and **Table S3**, respectively. $[\alpha]_D$ +64.6, c 0.169, DMSO. IR (MeOH, film, $\nu$($cm^{-1}$)): 2938 ($CH_2$), 2090 (indicator band, $NH_3^+$), 1742 (C=O), 1705 (C=O), 1582 ($COO^-$), 1537 ($NH_3^+$), 1254 (C-O). MS ESI: m/z 446.3 (6%, M - H). HR-MS (ESI) m/z: for $C_{26}H_{40}O_5N$ [M - H] calcd, 446.2912; found, 446.2909.

###### 2.3.1.4 Couplings of glutamate esters with DABCYL and TQ1

###### 2.3.1.4.1 General procedure for the coupling of glutamate esters with DABCYL

Steroid glutamate ester (0.056 mmol) was placed in a round-bottom flask equipped with a septum inlet and covered by aluminium foil. It was evacuated and refilled with argon two times. Freshly distilled dichloromethane (2 ml) was added upon stirring. It was followed by the addition of DIPEA (1 eq, 0.056 mmol, 10 µl) and Dabcyl succinimidyl ester (1 eq., 0.056 mmol, 20 mg) was added under argon stream. The reaction mixture was stirred at room temperature overnight, in dark. The solvent was evaporated in vacuo and the crude material was re-dissolved in dichloromethane (30 ml). The organic phase was washed with water (2 x 5 ml) and dried over anhydrous $Na_2SO_4$. It was filtered and the solvent was evaporated.

**(*S*)-5-(((3*R*,5*R*,8*R*,9*S*,10*S*,13*S*,14*S*,17*S*)-17-Acetyl-10,13-dimethylhexadecahydro-1*H*-cyclopenta[*a*]phenanthren3-yl)oxy)-4-(4-((*E*)-(4-(dimethylamino)phenyl)diazenyl)benzamido)-5-oxopentanoic acid (1)**

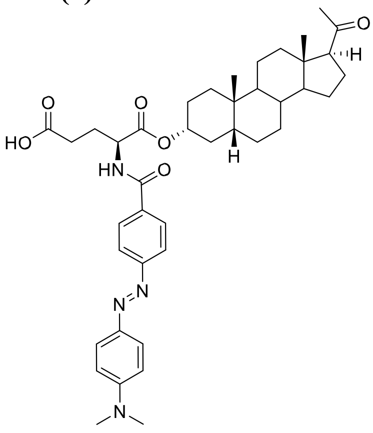

Starting from **35** (0.056 mmol, 25 mg), compound **1** was synthesized according to general procedure. The crude material was purified by preparative TLC on silica gel in eluent dichloromethane/methanol (6%) to give compound **1** (28 mg, 72%).

The $^1H$ and $^{13}C$ NMR chemical shifts and corresponding signal assignments are reported in **Table S2** and **Table S3**, respectively. MS ESI: m/z 697.4 (100%, M - H). HR-MS (ESI) m/z: for $C_{41}H_{53}O_6N_4$ [M - H] calcd, 697.3971; found, 697.3968.

**(*S*)-5-(((3*S*,5*R*,8*R*,9*S*,10*S*,13*S*,14*S*,17*S*)-17-Acetyl-10,13-dimethylhexadecahydro-1*H*-cyclopenta[*a*]phenanthren3-yl)oxy)-4-(4-((*E*)-(4-(dimethylamino)phenyl)diazenyl)benzamido)-5-oxopentanoic acid (2)**

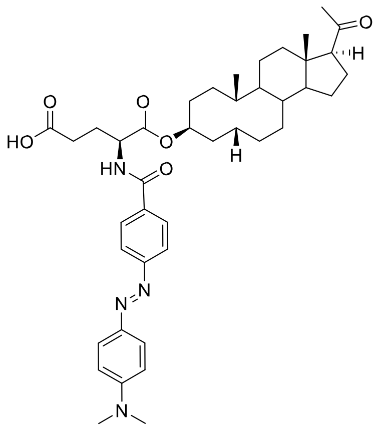

Starting from **36** (0.0715 mmol, 32 mg), compound **2** was synthesized according to general procedure, but using larger excess of DIPEA (6 eq., 74 µl) to enhance the dissolution of the starting steroid. The crude material was purified purified by flash chromatography on silica gel using a gradient of dichloromethane/EtOH (1% to 20%) in 20 column volumes. Compound **2** was isolated as a red solid (41 mg, 82%).
The $^{1}H$ and $^{13}C$ NMR chemical shifts and corresponding signal assignments are reported in **Table S2** and **Table S3**, respectively. MS ESI: m/z 697.4 (100%, M - H). HR-MS (ESI) m/z: for $C_{41}H_{53}O_6N_4$ [M - H] calcd, 697.3971; found, 697.3970.

**2.3.1.4.2 General procedure for the coupling of glutamate esters with TQ1**

Steroid glutamate ester (0.07 mmol, 32 mg) was placed in a round-bottom flask equipped with a septum inlet and covered by aluminium foil. It was evacuated and refilled with argon two times. Freshly distilled dichloromethane (2.5 ml) was added upon stirring. It was followed by the addition of DIPEA (1 eq., 0.07 mmol, 12 µl) and Tide Quencher 1 succinimidyl ester (1 eq., 0.07 mmol, 27 mg) was added under an argon stream. The reaction mixture was stirred at room temperature overnight, in dark. The solvent was evaporated in vacuo. The obtained crude material was purified by flash chromatography on silica gel using a gradient of dichloromethane/EtOH (1% to 20%) in 20 column volumes.

**(*S*)-5-(((3*R*,5*R*,8*R*,9*S*,10*S*,13*S*,14*S*,17*S*)-17-Acetyl-10,13-dimethylhexadecahydro-1*H*-cyclopenta[*a*]phenanthren3-yl)oxy)-4-(2-(2-((*E*)-(4-(dimethylamino)phenyl)diazenyl)thiazol-4-yl)acetamido)-5-oxopentanoic acid (3)**

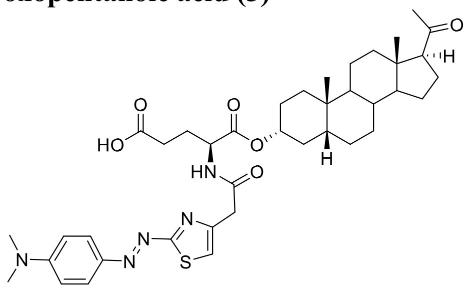

Starting from **35** (0.067 mmol, 30 mg), compound **3** was isolated as deep red solid (35 mg, 73%).
The $^{1}H$ and $^{13}C$ NMR chemical shifts and corresponding signal assignments are reported in **Table S2** and **Table S3**, respectively. MS ESI: m/z 742.4 (100%, M + Na). HR-MS (ESI) m/z: for $C_{39}H_{54}O_6N_5S$ [M + H] calcd, 720.3789; found, 720.3793. **(*S*)-5-(((3*S*,5*R*,8*R*,9*S*,10*S*,13*S*,14*S*,17*S*)-17-Acetyl-10,13-dimethylhexadecahydro-1*H*-cyclopenta[*a*]phenanthren-**

**3-yl)oxy)-4-(2-(2-((*E*)-(4-(dimethylamino)phenyl)diazenyl)thiazol-4-yl)acetamido)-5-oxopentanoic acid (4)**

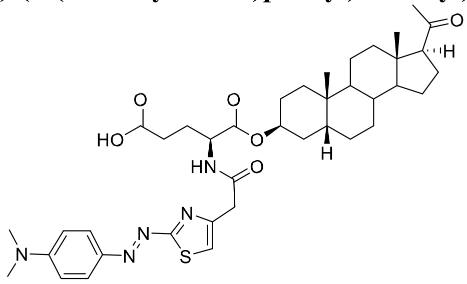

Starting from **36** (0.0706 mmol, 32 mg), the reaction was carried out according to the general procedure, but using larger excess of DIPEA (6 eq., 73 µl) to enhance the dissolution of the starting steroid. Compound **4** was isolated as deep red solid (49.2 mg, 97%).

The $^{1}H$ and $^{13}C$ NMR chemical shifts and corresponding signal assignments are reported in **Table S2** and **Table S3**, respectively. MS ESI: m/z 742.4 (100%, M + Na). HR-MS (ESI) m/z: for $C_{39}H_{54}O_6N_5S$ [M + H] calcd, 720.3789; found, 720.3792.

**(*S*)-5-(((3*R*,5*S*,8*R*,9*S*,10*S*,13*S*,14*S*,17*S*)-17-Acetyl-10,13-dimethylhexadecahydro-1*H*-cyclopenta[*a*]phenanthren3-yl)oxy)-4-(2-(2-((*E*)-(4-(dimethylamino)phenyl)diazenyl)thiazol-4-yl)acetamido)-5-oxopentanoic acid (5)**

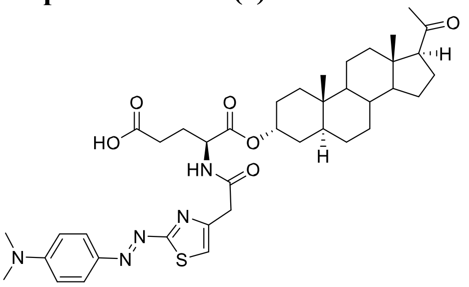

Starting from **37** (0.0324 mmol, 15 mg), compound **5** was isolated as deep red solid (23 mg, 99%).

The $^{1}H$ and $^{13}C$ NMR chemical shifts and corresponding signal assignments are reported in **Table S2** and **Table S3**, respectively. MS ESI: m/z 742.4 (100%, M + Na). HR-MS (ESI) m/z: for $C_{39}H_{54}O_6N_5S$ [M + H] calcd, 720.3789; found, 720.3787.

**(*S*)-5-(((3*S*,5*S*,8*R*,9*S*,10*S*,13*S*,14*S*,17*S*)-17-Acetyl-10,13-dimethylhexadecahydro-1*H*-cyclopenta[*a*]phenanthren3-yl)oxy)-4-(2-(2-((*E*)-(4-(dimethylamino)phenyl)diazenyl)thiazol-4-yl)acetamido)-5-oxopentanoic acid (6)**

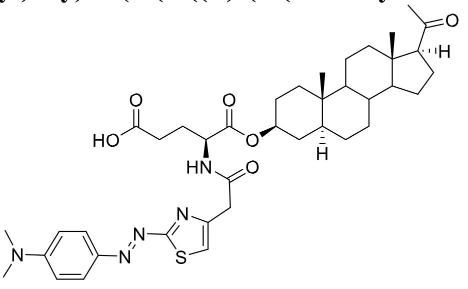

Starting from **38** (0.067 mmol, 30 mg), compound **6** was isolated as deep red solid (47.5 mg, 99%).

The $^{1}H$ and $^{13}C$ NMR chemical shifts and corresponding signal assignments are reported in **Table S2** and **Table S3**, respectively. MS ESI: m/z 742.4 (100%, M + Na). HR-MS (ESI) m/z: for $C_{39}H_{54}O_6N_5S$ [M + H] calcd, 720.3789; found, 720.3793.

##### 2.3.2 Synthesis of steroid-TQ1 conjugates via a GABA linker

###### 2.3.2.1 General procedure for the formation of *N*-Boc-GABA esters

Steroid (0.8 mmol, 255 mg) and DMAP (1 eq., 0.8 mmol, 98 mg) were placed in a round-bottom flask equipped with a septum inlet. The flask was evacuated and refilled with argon two times. It was followed by the addition of freshly distilled dichloromethane (16 ml) and the mixture was stirred. DIPEA (2.5 eq., 2 mmol, 348 µl) was added, after under argon stream the following solids: EDCI.HCl (2.5 eq., 2 mmol, 383 mg) and finally Boc-GABA-OH (2.5 eq., 2 mmol, 406.5 mg). The reaction mixture was stirred at room temperature overnight. After completion of the reaction, the solvent was evaporated in vacuo. The crude material was re-dissolved in dichloromethane (60 ml), washed with water (2 x 15 ml). The organic phase was dried over anhydrous $Na_2SO_4$. It was filtered off and the solvent was removed in vacuo.

**(3*R*,5*R*,8*R*,9*S*,10*S*,13*S*,14*S*,17*S*)-17-Acetyl-10,13-dimethylhexadecahydro-1*H*-cyclopenta[*a*]phenanthren-3-yl 4((*tert*-butoxycarbonyl)amino)butanoate (39)**

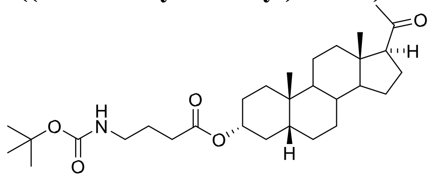

Starting from **3α5β-PRG (23)** (0.8 mmol, 255 mg), compound **39** was synthesized according to general procedure. The crude material was purified by flash chromatography on silica gel using a gradient n-hexane/ethyl-acetate (2% to 30% in 20 column volumes). The product **39** (369 mg, 92%) was isolated as viscous oil.
$[\alpha]_D$ +90.4, c 0.316, $CHCl_3$. Selected signals of **39** in $^1$H NMR (400 MHz, $CDCl_3$): $\delta$ 4.73 (tt, $J$ = 11.4 Hz, 4.7 Hz, 1H, H-3), 4.61 (br, 1H, NH), 3.16 (q, $J$ = 6.4 Hz, 2H, $CH_2$), 2.53 (t, $J$ = 8.9 Hz, 1H, H-17), 2.32 (t, $J$ = 7.3 Hz, 2H, $CH_2$), 2.11 (s, 3H, H-21), 1.44 (s, 9H, $(CH_3)_3C$-O), 0.93 (s, 3H, H-19), 0.59 (s, 3H, H-18). $^{13}$C NMR (101 MHz, $CDCl_3$): $\delta$ 209.7, 172.9, 156.1, 79.5, 74.5, 64.0, 56.8, 44.5, 42.0, 40.5, 40.3, 39.3, 35.9, 35.2, 34.8, 32.4, 32.2, 31.7, 28.6 (3C), 27.0, 26.8, 26.4, 25.5, 24.6, 23.4, 23.0, 21.0, 13.6. IR ($CHCl_3$, $\nu$(cm$^{-1}$)): 3456 (NH), 2959 ($CH_3$), 2937 ($CH_2$), 1725 (C=O), 1712 (C=O), 1702 (C=O), 1170 (C-O, Boc). MS ESI: m/z 404.3 (100%, M – Boc + H), 504.4 (77%, M + H), 526.3 (16%, M + Na). HR-MS (ESI) m/z: for $C_{30}H_{49}O_5NNa$ [M + Na] calcd, 526.3503; found, 526.3501.

**(3*S*,5*R*,8*R*,9*S*,10*S*,13*S*,14*S*,17*S*)-17-Acetyl-10,13-dimethylhexadecahydro-1*H*-cyclopenta[*a*]phenanthren-3-yl 4((*tert*-butoxycarbonyl)amino)butanoate (40)**

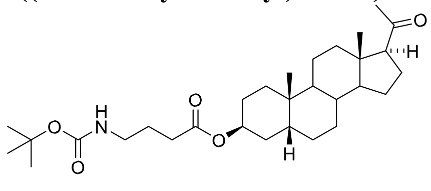

Starting from **3β5β-PRG (24)**, (0.628 mmol, 200 mg), compound **40** was synthesized according to general procedure. The crude material was purified by flash chromatography on silica gel using a gradient petroleum ether/ethyl-acetate (1% to 18% in 25 column volumes). The product **40** (174 mg, 55%) was isolated as viscous oil.
$[\alpha]_D$ +64.5, c 0.474, $CHCl_3$. Selected signals of **40** in $^1$H NMR (400 MHz, $CDCl_3$): $\delta$ 5.09 (p, $J$ = 2.6 Hz, 1H, H-3), 4.62 (br, 1H, NH), 3.17 (q, $J$ = 6.7 Hz, 2H, $CH_2$), 2.53 (t, $J$ = 8.9 Hz, 1H, H-17), 2.35 (t, $J$ = 7.3 Hz, 2H, $CH_2$), 2.11 (s, 3H, H-21), 1.44 (s, 9H, $(CH_3)_3C$-O), 0.96 (s, 3H, H-19), 0.60 (s, 3H, H-18). $^{13}$C NMR (101 MHz, $CDCl_3$): $\delta$ 209.8, 172.9, 156.1, 79.4, 70.9, 64.0, 56.9, 44.5, 40.1, 40.0, 39.4, 37.5, 35.8, 35.1, 32.2, 31.7, 30.9, 30.8, 28.6 (3C), 26.5, 26.3, 25.6, 25.2, 24.6, 24.0, 23.0, 21.2, 13.6. IR ($CHCl_3$, $\nu$(cm$^{-1}$)): 3456 (NH), 2978 ($CH_3$), 2939 ($CH_2$), 1725 (C=O), 1709 (C=O), 1703 (C=O), 1172 (C-O, Boc). MS ESI: m/z 404.3 (2%, M – Boc + H), 504.4 (7%, M + H), 526.4 (100%, M + Na). HR-MS (ESI) m/z: for $C_{30}H_{49}O_5NNa$ [M + Na] calcd, 526.3503; found, 526.3504.

**(3*R*,5*S*,8*R*,9*S*,10*S*,13*S*,14*S*,17*S*)-17-Acetyl-10,13-dimethylhexadecahydro-1*H*-cyclopenta[*a*]phenanthren-3-yl 4((*tert*-butoxycarbonyl)amino)butanoate (41)**

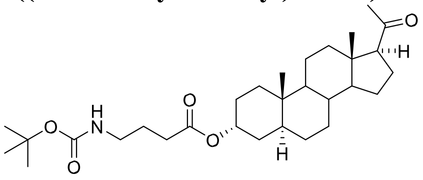

Starting from **3α5α-PRG (25)** (1 mmol, 318.5 mg), **41** was synthesized according to general procedure. The crude material was purified by flash chromatography on silica gel using a gradient petroleum ether/ethyl-acetate (1% to 20% in 20 column volumes). The product **41** (447 mg, 89%) was isolated as viscous oil.

$[\alpha]_D$ +65.4, c 0.286, $CHCl_3$. Selected signals of **41** in $^1$H NMR (400 MHz, $CDCl_3$): *δ* 5.03 (p, *J* = 2.6 Hz, 1H, H-3), 4.63 (br, 1H, NH), 3.17 (q, *J* = 6.6 Hz, 2H, $CH_2$), 2.52 (t, *J* = 8.9 Hz, 1H, H-17), 2.35 (t, *J* = 7.4 Hz, 2H, $CH_2$), 2.11 (s, 3H, H-21), 1.44 (s, 9H, $(CH_3)_3$C-O), 0.79 (s, 3H, H-19), 0.60 (s, 3H, H-18). $^{13}$C NMR (101 MHz, $CDCl_3$): *δ* 209.9, 172.9, 156.1, 79.4, 70.3, 64.0, 56.9, 54.2, 44.4, 40.2, 40.2, 39.2, 36.0, 35.6, 33.1, 33.0, 32.3, 32.0, 31.7, 28.6 (3C), 28.4, 26.2, 25.6, 24.5, 22.9, 21.0, 13.6, 11.5. IR ($CHCl_3$, $\nu$($cm^{-1}$)): 3455 (NH), 2972 ($CH_3$), 1730 (C=O), 1711 (C=O), 1703 (C=O), 1162 (C-O, Boc). MS ESI: m/z 404.3 (3%, M – Boc + H), 504.4 (22%, M + H), 526.4 (100%, M + Na). HR-MS (ESI) m/z: for $C_{30}H_{49}O_5NNa$ [M + Na] calcd, 526.3503; found, 526.3505.

**(3*S*,5*S*,8*R*,9*S*,10*S*,13*S*,14*S*,17*S*)-17-Acetyl-10,13-dimethylhexadecahydro-1*H*-cyclopenta[*a*]phenanthren-3-yl 4((*tert*-butoxycarbonyl)amino)butanoate (42)**

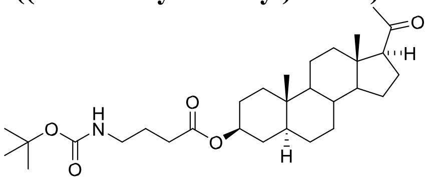

Starting from **3β5α-PRG (26)** (1 mmol, 318.5 mg), **42** was synthesized according to general procedure. The crude material was purified by flash chromatography on silica gel using a gradient petroleum ether/ethyl-acetate (1% to 20% in 20 column volumes). The product **42** (478 mg, 95%) was isolated as viscous oil.

$[\alpha]_D$ +54.9, c 0.272, $CHCl_3$. Selected signals of **42** in $^1$H NMR (400 MHz, $CDCl_3$): *δ* 4.70 (tt, *J* = 11.4 Hz, 4.9 Hz, 1H, H-3), 4.61 (br, 1H, NH), 3.15 (q, *J* = 6.6 Hz, 2H, $CH_2$), 2.52 (t, *J* = 8.9 Hz, 1H, H-17), 2.31 (t, *J* = 7.3 Hz, 2H, $CH_2$), 2.11 (s, 3H, H-21), 1.43 (s, 9H, $(CH_3)_3$C-O), 0.82 (s, 3H, H-19), 0.60 (s, 3H, H-18). $^{13}$C NMR (101 MHz, $CDCl_3$): *δ* 209.8, 173.0, 156.1, 79.4, 73.9, 64.0, 56.8, 54.2, 44.8, 44.4, 40.1, 39.2, 36.9, 35.7, 35.6, 34.1, 32.2, 32.1, 31.7, 28.6 (3C), 27.6, 25.5, 24.5, 22.9, 21.4, 13.6, 12.4. 1 carbon could not be detected. IR ($CHCl_3$, $\nu$($cm^{-1}$)): 3455 (NH), 2977 ($CH_3$), 1730 (C=O), 1710 (C=O), 1703 (C=O), 1170 (C-O, Boc). MS ESI: m/z 504.4 (23%, M + H), 526.3 (100%, M + Na). HR-MS (ESI) m/z: for $C_{30}H_{49}O_5NNa$ [M + Na] calcd, 526.3503; found, 526.3501.

**(3*S*,8*S*,9*S*,10*R*,13*S*,14*S*,17*S*)-17-Acetyl-10,13-dimethyl-2,3,4,7,8,9,10,11,12,13,14,15,16,17-tetradecahydro-1*H*cyclopenta[*a*]phenanthren-3-yl 4-((*tert*-butoxycarbonyl)amino)butanoate (48)**

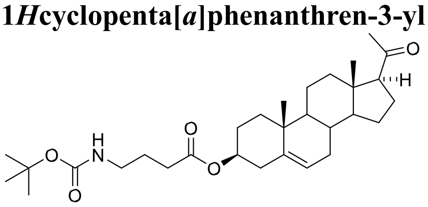

Starting from pregnenolone (**47**) (3.16 mmol, 1 g), compound **48** was synthesized according to general procedure. The crude material was purified by flash chromatography on silica gel using a gradient of petroleum ether/ether (0% to 63% in 22 column volumes). The product **48** (1.38 g, 87%) was isolated as white solid.

$[\alpha]_D$ +14.4, c 0.421, $CHCl_3$. Selected signals of **48** in $^1$H NMR (400 MHz, $CDCl_3$): *δ* 5.45-5.30 (m, 1H, H-6), 4.70-4.51 (m, 2H, NH, H-3), 3.16 (q, *J* = 6.6 Hz, 2H, $CH_2$), 2.53 (t, *J* = 8.9 Hz, 1H, H-17), 2.39-2.25 (m, 2H, $CH_2$, overlapping signal), 2.12 (s, 3H, H-21), 1.44 (s, 9H, $(CH_3)_3$C-O), 1.02 (s, 3H, H-19), 0.63 (s, 3H, H-18). $^{13}$C NMR (101 MHz, $CDCl_3$): *δ* 209.7, 172.9, 156.1, 139.8, 122.5, 79.4, 74.1, 63.8, 57.0, 50.0, 44.1, 40.1, 38.9, 38.2, 37.1, 36.7, 32.1, 32.0,

31.9, 31.7, 28.6 (3C), 27.9, 25.5, 24.6, 23.0, 21.2, 19.5, 13.4. IR ($CHCl_3$, $\nu$($cm^{-1}$)): 3456 (NH), 2971 ($CH_3$), 1730 (C=O), 1712 (C=O), 1703 (C=O), 1170 (C-O, Boc). MS ESI: m/z 524.3 (100%, M + Na). HR-MS (ESI) m/z: for $C_{30}H_{47}O_5NNa$ [M + Na] calcd, 524.3346; found, 526.3348.

**(3*S*,8*R*,9*S*,10*R*,13*S*,14*S*)-10,13-Dimethyl-17-oxo-2,3,4,7,8,9,10,11,12,13,14,15,16,17-tetradecahydro-1*H*cyclopenta[*a*]phenanthren-3-yl 4-((*tert*-butoxycarbonyl)amino)butanoate (51)**

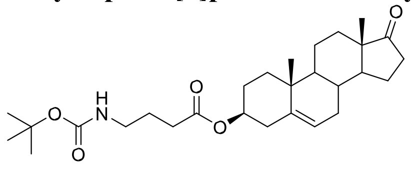

Starting from dehydroepiandrosterone (**50**) (3.47 mmol, 1 g), **51** was synthesized according to general procedure. The crude material was purified by flash chromatography on silica gel using a gradient of petroleum ether/ethyl acetate (0% to 25%) resulting in 2.09 g of white solid (theoretical mass 1.64 g). According to NMR analysis of the obtained solid, it contained the desired product **51** and *tert*-butyl 2-oxopyrrolidine-1-carboxylate as a byproduct. According to TLC analysis in different eluents, the product **51** and *tert*-butyl 2-oxopyrrolidine-1-carboxylate have identical $R_f$ values. As it was not possible to separate them by column chromatography, or by crystallization, the obtained mixture was used in the next reaction step.
Selected signals of **51** in $^1$H NMR (400 MHz, $CDCl_3$): $\delta$ 5.45-5.36 (m, 1H, 6-H), 4.70-4.52 (m, 2H, 3-H and NH), 3.16 (q, $J$ = 6.6 Hz, 2H, $CH_2$), 2.32 (t, $J$ = 7.4 Hz, 2H, $CH_2$, overlapping signal), 1.43 (s, 9H, $(CH_3)_3$C-O), 1.04 (s, 3H, H-19), 0.88 (s, 3H, H-18). $^{13}$C NMR (101 MHz, $CDCl_3$): $\delta$ 172.8, 156.1, 140.0, 122.1, 79.4, 74.0, 51.8, 50.3, 47.7, 40.1, 38.2, 37.1, 36.9, 36.0, 32.1, 31.6, 31.5, 30.9, 28.6 (3C), 27.8, 25.5, 22.0, 20.5, 19.5, 13.7. C=O could not be detected. HR-MS (ESI) m/z: for $C_{28}H_{44}NO_5$ [M + H] calcd, 474.3214; found, 474.3213.

In the $^1$H and $^{13}$C NMR spectra, signals of *tert*-butyl 2-oxopyrrolidine-1-carboxylate correspond to the literature.[3]

**(3*S*,5*R*,8*S*,9*S*,10*S*,13*R*,14*S*,17*S*)-10,13,17-Trimethylhexadecahydro-1*H*-cyclopenta[*a*]phenanthren-3-ol (54)**

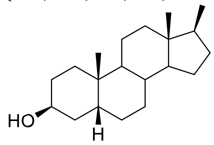

17β-Methyl-5β-androstan-3-one (**53**) (1 eq., 4.5 mmol, 1.3 g) was dissolved in 20 ml of dry THF and cooled to -78 °C. L-selectride (1.2 eq., 5.4 mmol, 5.4 ml, 1.0 M in THF) was added dropwise to the steroid solution. The reaction mixture was stirred for two hours at -78 °C, then one hour at room temperature. After completion of the reaction, the solvent was removed in vacuo. The crude material was redissolved in ethyl acetate and was washed with saturated $NH_4Cl$ (10 ml), brine (10 mL) and dried with $MgSO_4$. It was filtered off and the solvent was evaporated. The crude material was purified by flash chromatography on silica gel using a gradient *n*-hexane/ethyl-acetate (0% to 15% in 35 column volumes). The product 17β-methyl-5β-androstan-3β-ol was isolated as an off-white powder (1.28 g, 97%).
$[\alpha]_D$ +16.2, c 0.245, $CHCl_3$. Selected signals of **54** in $^1$H NMR (400 MHz, $CDCl_3$): $\delta$ 4.11 (p, $J$ = 2.8 Hz, 1H, H-3), 0.97 (s, 3H, H-19), 0.82 (d, $J$ = 6.8 Hz, 3H, H-20), 0.53 (d, $J$ = 0.7 Hz, 3H, H-18). $^{13}$C NMR (101 MHz, $CDCl_3$): $\delta$ 67.4, 56.1, 45.3, 42.4, 40.3, 37.9, 36.9, 36.0, 35.4, 33.7, 30.4, 30.2, 28.0, 26.8, 26.6, 24.9, 24.1, 21.0, 14.0, 12.2. IR ($CHCl_3$, $\nu$($cm^{-1}$)): 3616 (OH), 2982 ($CH_3$), 2978 ($CH_3$). HR-MS (EI) m/z: for $C_{20}H_{34}O$ [M+] calcd, 290.2604; found, 290.2602.

**(3*S*,5*R*,8*S*,9*S*,10*S*,13*R*,14*S*,17*S*)-10,13,17-Trimethylhexadecahydro-1*H*-cyclopenta[*a*]phenanthren-3-yl 4-((*tert*butoxycarbonyl)amino)butanoate (55)**

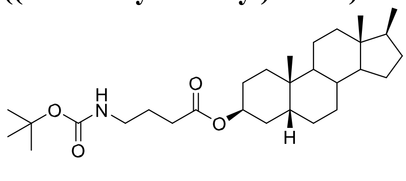

Starting from compound **54** (0.69 mmol, 200 mg), **55** was synthesized according to general procedure. The crude material was purified by flash chromatography on silica gel using a gradient n-hexane/ethyl-acetate (0% to 20% in 30 column volumes). The product **55** was isolated as an off-white powder (200 mg, 61%).
$[\alpha]_D$ +9.7, c 0.420, $CHCl_3$. Selected signals of **55** in $^1$H NMR (400 MHz, $CDCl_3$): $\delta$ 5.08 (p, $J$ = 3.1 Hz, 1H, H-3), 4.62 (br, 1H, NH), 3.16 (q, $J$ = 6.7 Hz, 2H, $CH_2$), 2.34 (t, $J$ = 7.4 Hz, 2H, $CH_2$), 1.43 (s, 9H, $(CH_3)_3$C-O), 0.97 (s, 3H, H-19), 0.81 (d, $J$ = 6.8 Hz, 3H, H-20), 0.52 (d, $J$ = 0.6 Hz, 3H, H-18). $^{13}$C NMR (101 MHz, $CDCl_3$): $\delta$ 172.9, 156.1, 79.3, 71.1, 56.0, 45.3, 42.3, 40.4, 37.9, 37.7, 36.0, 35.2, 31.1, 30.8, 30.4, 28.5 (3C), 26.7, 26.5, 25.2, 24.9, 24.1, 21.0, 14.0, 12.2. IR ($CHCl_3$, $\nu$(cm$^{-1}$)): 3455 (NH), 2960 ($CH_3$), 1725 (C=O). MS ESI: m/z 498.4 (100%, M + Na). HR-MS (ESI) m/z: for $C_{29}H_{49}NO_4Na$ [M + Na] calcd, 498.3554; found, 498.3549.

###### 2.3.2.2 General procedure for the Boc deprotection of compound 39 using HCl in diethyl ether 4-(((3*R*,5*R*,8*R*,9*S*,10*S*,13*S*,14*S*,17*S*)-17-Acetyl-10,13-dimethylhexadecahydro-1*H*-cyclopenta[*a*]phenanthren-3yl)oxy)-4-oxobutan-1-aminium chloride (43)

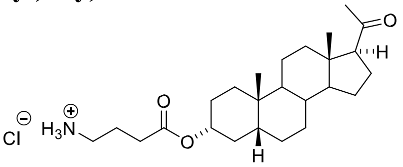

Steroid **39** (0.641 mmol, 323 mg) was placed in a round bottom-flask equipped with a septum inlet. It was evacuated and refilled with argon. To the steroid, HCl in diethyl-ether (3.2 ml, 1 M) was added slowly upon stirring. The reaction mixture was stirred overnight at room temperature. After 24 h, starting material was detected by TLC, therefore additional 1.5 ml HCl in diethyl ether (1 M) was added. After completion of the reaction, the solvent was removed in vacuo. To the crude solid material 2 ml n-heptane was added and stirred for 20 minutes. The solid was filtered off and was washed with ice-cold n-heptane (3 x 1 ml). The product was dried further in high vacuum and isolated as a white solid **43** (169 mg, 60%).
The $^1$H and $^{13}$C NMR chemical shifts and corresponding signal assignments are reported in **Table S4** and **Table S5**, respectively. $[\alpha]_D$ +90.2, c 0.224, MeOH. IR (KBr, $\nu$(cm$^{-1}$)): 3435 (NH), 2933 ($CH_2$), 2869 ($CH_2$), 1729 (C=O), 1706 (C=O), 1605 ($NH_3^+$), 1195 (C-O). MS ESI: m/z 404.3 (100%, M + H). HR-MS (ESI) m/z: for $C_{25}H_{42}O_3N$ [M + H] calcd, 404.3159; found, 404.3155.

###### 2.3.2.3 General procedure for the Boc deprotection of compounds 40-42 and 48, 51, 55 in the presence of trifluoroacetic acid

Steroid ester (0.3 mmol) was dissolved in 4.5 ml dichloromethane and upon stirring, 1 ml trifluoroacetic acid diluted with 1.3 ml dichloromethane was added dropwise. The reaction mixture was stirred at room temperature for 3 hours. The dichloromethane was evaporated, and the residue was further dried under nitrogen stream. The crude material was dissolved in dichloromethane (40 ml) and washed with saturated $NaHCO_3$ solution (15 ml), then with water (15 ml). The solvent was evaporated and the obtained material was used in the next step without further purification. As the obtained amines showed limited solubility in various organic solvents, they were converted and characterized as succinate salts (see Section 2.3.2.2.3.).

- Starting from compound **40** (0.3 mmol, 150 mg), compound **44** was isolated as off white solid (115 mg, 96%). For characterization of its salts form (**44a**) see **Section 2.3.2.2.3.**
- Starting from compound **41** (0.7 mmol, 350 mg), compound **45** was isolated as off white solid (262 mg, 94%). For characterization of its salts form (**45a**) see **Section 2.3.2.2.3.**
- Starting from compound **42** (0.75 mmol, 380 mg), compound **46** was isolated as off white solid (287 mg, 94%). For characterization of its salts form (**46a**) see **Section 2.3.2.2.3.**

###### 2.3.2.4 General procedure for the succinate salt formation of steroid amines 44, 45 and 46

For the synthesis of succinate salts, the steroid amine (0.193 mmol, 78 mg) was suspended in 2.5 ml ethanol/ethyl acetate 4/1 (v/v) mixture followed by the addition of succinic acid (1 eq., 0.193 mmol, 22.8 mg). After stirring at room temperature for 20 minutes (**44**) or overnight (**45** and **46**), the solvent was evaporated. The isolated material was recrystallized from ethyl acetate/methanol 9/1 (v/v) mixture. **4-(((3*S*,5*R*,8*R*,9*S*,10*S*,13*S*,14*S*,17*S*)-17-Acetyl-10,13-dimethylhexadecahydro-1*H*-cyclopenta[*a*]phenanthren-3yl)oxy)-4-oxobutan-1-aminium 3-carboxypropanoate (44a)**

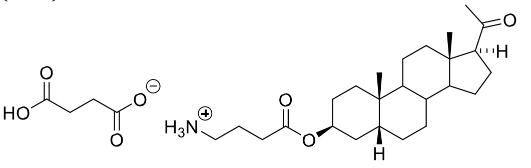

Starting from **44** (0.193 mmol, 78 mg), salt form **44a** was synthesized according to general procedure. The product **44a** was isolated as white crystalline solid (54 mg, 54%).

$[\alpha]_D$ +61.5, c 0.267, MeOH. Selected signals of **44a** in $^1$H NMR (400 MHz, DMSO-*d*$_6$): $\delta$ 5.02-4.94 (m, 1H, H-3), 2.812.73 (m, 2H, $CH_2$), 2.56 (t, $J$ = 8.9 Hz, 1H, H-17), 2.40 (t, $J$ = 7.5 Hz, 2H, $CH_2$), 2.25 (s, 4H, $CH_2$-$CH_2$ succinate), 2.05 (s, 3H, H-21), 0.93 (s, 3H, H-19), 0.51 (s, 3H, H-18). $^{13}$C NMR (101 MHz, DMSO-*d*$_6$): $\delta$ 208.7, 175.3 (2C), 171.8, 70.3,
63.0, 56.1, 43.8, 38.8, 38.5, 37.2, 35.3, 34.7, 32.4 (2C, succinate), 31.4, 31.1, 30.7, 30.2, 26.2, 25.9, 24.6, 24.1, 23.8, 23.7, 23.5, 22.5, 20.8, 13.3. IR (KBr, $\nu$(cm$^{-1}$)): 3137 ($NH_3^+$), 2936 ($CH_2$), 2874 ($CH_2$), 1728 (C=O), 1706 (C=O), 1641 ($NH_3^+$), 1202 (C-O). MS ESI: m/z 404.3 (100%, M + H). HR-MS (ESI) m/z: for $C_{25}H_{42}O_3N$ [M + H] calcd, 404.3159; found, 404.3157.

**4-(((3*R*,5*S*,8*R*,9*S*,10*S*,13*S*,14*S*,17*S*)-17-Acetyl-10,13-dimethylhexadecahydro-1*H*-cyclopenta[*a*]phenanthren-3yl)oxy)-4-oxobutan-1-aminium 3-carboxypropanoate (45a)**

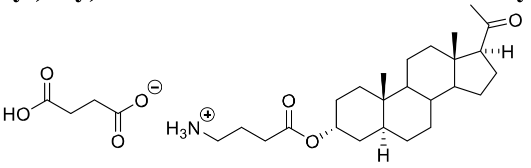

Starting from **45** (0.144 mmol, 58 mg), salt form **45a** was synthesized according to general procedure. The product **45a** was isolated as white crystalline solid (48 mg, 63%).

$[\alpha]_D$ +60.6, c 0.315, MeOH. Selected signals of **45a** in $^1$H NMR (400 MHz, DMSO-*d*$_6$): $\delta$ 4.98-4.88 (m, 1H, H-3), 2.852.74 (m, 2H, $CH_2$), 2.55 (t, $J$ = 8.9 Hz, 1H, H-17), 2.41 (t, $J$ = 7.5 Hz, 2H, $CH_2$), 2.27 (s, 4H, $CH_2$-$CH_2$ succinate), 2.05 (s, 3H, H-21), 0.77 (s, 3H, H-19), 0.51 (s, 3H, H-18). $^{13}$C NMR (101 MHz, DMSO-*d*$_6$): 208.7, 175.1 (2C), 171.8, 69.7,
62.9, 56.1, 53.8, 43.7, 38.6, 38.3, 35.6, 35.1, 32.7, 32.5 (2C, succinate), 32.0, 31.7, 31.4, 31.0, 28.0, 25.7, 24.1, 23.2, 22.4, 20.5, 13.3, 11.2. 1 carbon could not be detected. IR (KBr, $\nu$(cm$^{-1}$)): 3178 ($NH_3^+$), 2930 ($CH_2$), 2874 ($CH_2$), 1731 (C=O), 1706 (C=O), 1636 ($NH_3^+$), 1208 (C-O). MS ESI: m/z 404.3 (100%, M + H). HR-MS (ESI) m/z: for $C_{25}H_{42}O_3N$ [M + H] calcd, 404.3159; found, 404.3158.

**4-(((3*S*,5*S*,8*R*,9*S*,10*S*,13*S*,14*S*,17*S*)-17-Acetyl-10,13-dimethylhexadecahydro-1*H*-cyclopenta[*a*]phenanthren-3yl)oxy)-4-oxobutan-1-aminium succinate (46a)**

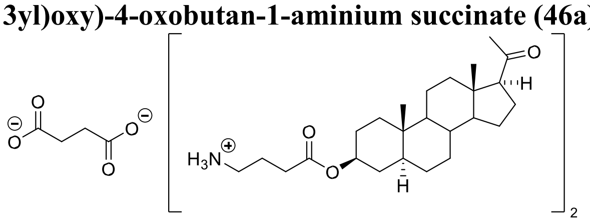

Starting from **46** (0.193 mmol, 78 mg), salt form **46a** was synthesized according to general procedure. The starting material showed limited solubility, therefore double volume of the solvent was used for the salt formation. The obtained crude material was partially soluble in methanol, therefore, the undissolved material was filtered and the solvent was evaporated. After evaporation, the obtained residue was further subjected to crystallization. The product **46a** was isolated as white crystalline solid (38 mg, 38%).

$[\alpha]_D$ +59.2, c 0.311, MeOH. Selected signals of **46a** in $^1$H NMR (400 MHz, DMSO-*d*$_6$): $\delta$ 4.60 (tt, $J$ = 11.2 Hz, 4.8 Hz, 1H, H-3), 2.73-2.63 (m, 2H, $CH_2$), 2.56 (t, $J$ = 8.9 Hz, 1H, H-17), 2.33 (t, $J$ = 7.4 Hz, 2H, $CH_2$), 2.23 (s, 4H, $CH_2$-$CH_2$, succinate counterion in ratio of 0.5), 2.05 (s, 3H, H-21), 0.78 (s, 3H, H-19), 0.51 (s, 3H, H-18). $^{13}$C NMR (150.9 MHz, DMSO-$d_6$): $\delta$ 208.8, 175.5 (2C), 172.1, 73.1, 62.9, 56.0, 53.5, 44.1, 43.7, 39.3, 38.3, 36.3, 35.2, 35.1, 33.9, 32.9 (2C, succinate), 31.7, 31.4, 31.1, 28.2, 27.3, 24.8, 24.1, 22.4, 20.9, 13.3, 12.1. IR (KBr, $\nu$(cm$^{-1}$)): 3098 ($NH_3^+$), 2936 ($CH_2$), 2847 ($CH_2$), 1732 (C=O), 1706 (C=O), 1630 ($NH_3^+$), 1200 (C-O). MS ESI: m/z 404.3 (100%, M + H). HR-MS (ESI) m/z: for $C_{25}H_{42}O_3N$ [M + H] calcd, 404.3159; found, 404.3157.

**4-(((3*S*,8*S*,9*S*,10*R*,13*S*,14*S*,17*S*)-17-Acetyl-10,13-dimethyl-2,3,4,7,8,9,10,11,12,13,14,15,16,17-tetradecahydro1*H*-cyclopenta[*a*]phenanthren-3-yl)oxy)-4-oxobutan-1-aminium chloride (49a)**

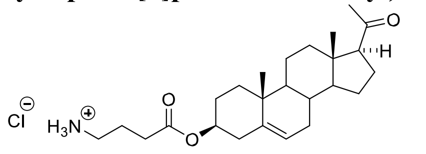

The Boc deprotection of **48** (0.2 mmol, 100 mg) was performed in the same manner as described in the general procedure for Boc deprotection in the presence of TFA. Modification of the workup: the crude material was dissolved in dichloromethane (40 ml) and washed with saturated $NaHCO_3$ solution (15 ml), with brine (15 ml), dried over $MgSO_4$, and filtered. For characterization, the obtained material (**49**) was further transformed to a hydrochloride salt: the amine was dissolved in diethyl ether (7 ml), and the solution was cooled in an ice-bath. It was followed by the addition of 1M HCl in diethyl ether (1 ml) and white precipitate appeared. The precipitate was collected and dried under a vacuum. The amine hydrochloride **49a** was obtained as a white solid (80 mg, 91%).
$[\alpha]_D$ +11.6, c 0.277, $CHCl_3$. Selected signals of **49a** in $^1$H NMR (400 MHz, $CD_3OD$): $\delta$ 5.45-5.37 (m, 1H, H-6), 4.644.50 (m, 1H, H-3), 3.03-2.93 (m, 2H, $CH_2$), 2.65 (t, $J$ = 8.9 Hz, 1H, H-17), 2.46 (t, $J$ = 7.1 Hz, 2H, $CH_2$), 2.13 (s, 3H, H-21), 1.05 (s, 3H, H-19), 0.64 (s, 3H, H-18). $^{13}$C NMR (101 MHz, MeOD): $\delta$ 212.3, 173.5, 141.0, 123.6, 75.8, 64.6, 58.0, 51.4, 45.1, 40.1, 39.8, 39.1, 38.2, 37.8, 33.2, 32.9, 31.9, 31.6, 28.8, 25.5, 23.8, 23.8, 22.2, 19.7, 13.6. IR (KBr, $\nu$(cm$^{-1}$)): 3032 ($NH_3^+$), 2942 ($CH_2$), 2849 ($CH_2$), 1730 (C=O), 1706 (C=O), 1608 ($NH_3^+$), 1193 (C-O). MS ESI: m/z 402.3 (100%, M + H). HR-MS (ESI) m/z: for $C_{25}H_{40}O_3N$ [M + H] calcd, 402.3003; found, 402.2999.

**4-(((3*S*,8*R*,9*S*,10*R*,13*S*,14*S*)-10,13-Dimethyl-17-oxo-2,3,4,7,8,9,10,11,12,13,14,15,16,17-tetradecahydro-1*H*cyclopenta[*a*]phenanthren-3-yl)oxy)-4-oxobutan-1-aminium chloride (52a)**

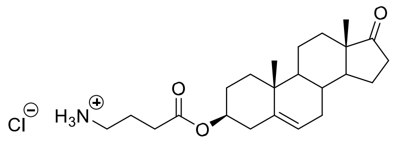

Steroid ester **51** (1.0 equiv, 2.69 mmol, 1.27 g, mixture with *tert*-butyl 2-oxopyrrolidine-1-carboxylate) was dissolved in 30 ml dichloromethane and under stirring, 2.7 ml of trifluorocaetic acid was added dropwise at room temperature. The reaction mixture was stirred at room temperature for 1.5 hours. The dichloromethane was evaporated, and the residue was further dried under vacuum. After the addition of $Et_2O$ to the crude material, white precipitate was formed and filtered off (1.34 g) (**52**). For characterization, part of the obtained material was converted to a hydrochloride salt: 0.499 g of the crude material was dissolved in 2 ml of EtOAc and 6 ml of EtOH. Concentrated HCl acid (0.5 ml) was added under stirring at room temperature. The reaction mixture was stirred for 1 hour at room temperature, the formed white precipitate was filtered, washed with diethyl ether and dried. The amine hydrochloride **52a** was obtained as a white solid (222 mg). Because of the presence of *tert*-butyl 2-oxopyrrolidine-1-carboxylate in the starting material, yield was not calculated.
$[\alpha]_D$ +2.1, c 0.263, $CHCl_3$. Selected signals of **52a** in $^1$H NMR (400 MHz, $CDCl_3$): $\delta$ 8.48-8.08 (brs, 3H, $NH_3^+$), 5.475.29 (m, 1H, 6-H), 4.69-4.47 (m, 1H, 3-H), 3.20-2.94 (m, 2H, $CH_2$), 1.02 (s, 3H, H-19), 0.87 (s, 3H, H-18). $^{13}$C NMR (101 MHz, $CDCl_3$): $\delta$ 172.2, 139.9, 122.2, 74.5, 51.8, 50.2, 47.6, 38.1, 37.0, 36.8, 35.9, 31.6, 31.5, 31.5, 30.9, 27.8, 22.0, 21.9, 20.4, 19.4, 13.6. C=O and another carbon could not be detected. IR (KBr, $\nu$(cm$^{-1}$)): 3226 ($NH_3^+$), 2975 ($CH_3$), 2940 ($CH_2$), 2887 ($CH_3$), 1733 (C=O), 1670 (C=C), 1193 (C-O). MS ESI: m/z 374.3 (100%, M + H). HR-MS (ESI) m/z: for $C_{23}H_{36}NO_3$ [M + H] calcd, 374.2690; found, 374.2687.

**(3*S*,5*R*,8*S*,9*S*,10*S*,13*R*,14*S*,17*S*)-10,13,17-Trimethylhexadecahydro-1*H*-cyclopenta[*a*]phenanthren-3-yl 4aminobutanoate (56)**

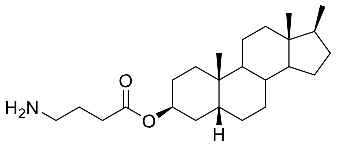

The Boc deprotection of **55** (0.29 mmol, 140 mg) was performed in the same manner as described in the general procedure for Boc deprotection in the presence of TFA. The product **56** was isolated as an off white solid (100 mg, 90%). Selected signals of **56** in $^1$H NMR (400 MHz, $CDCl_3$): $\delta$ 5.08 (p, $J$ = 3.0 Hz, 1H), 2.84-2.66 (m, 2H, $CH_2$), 2.36 (t, $J$ = 7.4 Hz, 2H, $CH_2$), 0.97 (s, 3H, H-19), 0.81 (d, $J$ = 6.8 Hz, 3H, H-20), 0.52 (s, 3H, H-18). $^{13}$C NMR (101 MHz, $CDCl_3$): $\delta$ 173.2, 70.9, 56.0, 45.3, 42.3, 41.6, 40.4, 37.9, 37.7, 36.0, 35.2, 32.4, 31.1, 30.8, 30.4, 28.9, 26.7, 26.5, 25.2, 24.9, 24.1, 21.0, 14.0, 12.2. IR ($CHCl_3$, $\nu$(cm$^{-1}$)): 3455 (NH), 2960 ($CH_3$), 1725 (C=O). MS ESI: m/z 376.3 (100%, M + H). HRMS (ESI) m/z: for $C_{24}H_{42}NO_2$ [M + H] calcd, 376.3210; found, 376.3209.

##### 2.3.2.5 Synthesis of steroid-TQ1 conjugates via a GABA linker

###### 2.3.2.5.1 General procedure for the coupling with TQ1

Steroid ester (0.056 mmol, 25 mg) was placed in a round-bottom flask equipped with a septum inlet and covered by aluminium foil. It was evacuated and refilled with argon two times. Freshly distilled dichloromethane (2 ml) was added upon stirring. It was followed by the addition of DIPEA (2 eq., 0.111 mmol, 19 µl) and Tide Quencher succinimidyl ester (1 eq., 0.056 mmol, 22 mg) was added under argon stream. The reaction mixture was stirred at room temperature overnight, in dark. After completion of the reaction, the solvent was evaporated in vacuo.

**(3*R*,5*R*,8*R*,9*S*,10*S*,13*S*,14*S*,17*S*)-17-Acetyl-10,13-dimethylhexadecahydro-1*H*-cyclopenta[*a*]phenanthren-3-yl 4(2-(2-((*E*)-(4-(dimethylamino)phenyl)diazenyl)thiazol-4-yl)acetamido)butanoate (7)**

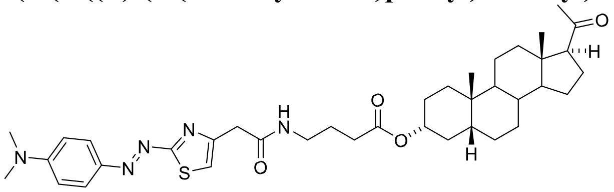

Starting from **43** (0.056 mmol, 25 mg), compound **7** was synthesized according to general procedure. The crude material was purified by flash chromatography on silica gel using a gradient dichloromethane/acetone (0% to 20% in 20 column volumes). The product **7** (33 mg, 87%) was isolated as red solid.
The $^1$H and $^{13}$C NMR chemical shifts and corresponding signal assignments are reported in **Table S4** and **Table S5**, respectively. MS ESI: m/z 676.4 (29%, M + H), 698.4 (100%, M + Na). HR-MS (ESI) m/z: for $C_{38}H_{54}O_4N_5S$ [M + H] calcd, 676.3891; found, 676.3888.

**(3*S*,5*R*,8*R*,9*S*,10*S*,13*S*,14*S*,17*S*)-17-Acetyl-10,13-dimethylhexadecahydro-1*H*-cyclopenta[*a*]phenanthren-3-yl 4(2-(2-((*E*)-(4-(dimethylamino)phenyl)diazenyl)thiazol-4-yl)acetamido)butanoate (8)**

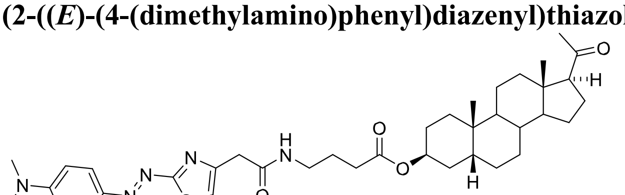

Starting from **44** (0.0847 mmol, 34 mg), compound **8** was synthesized according to general procedure. The crude material was purified by flash chromatography on silica gel using a gradient dichloromethane/acetone (0% to 17% in 20 column volumes). The product **8** (48 mg, 84%) was isolated as red solid.
The $^1$H and $^{13}$C NMR chemical shifts and corresponding signal assignments are reported in **Table S4** and **Table S5**, respectively. MS ESI: m/z 676.4 (23%, M + H), 698.4 (100%, M + Na). HR-MS (ESI) m/z: for $C_{38}H_{54}O_4N_5S$ [M + H] calcd, 676.3891; found, 676.3890. **(3*R*,5*S*,8*R*,9*S*,10*S*,13*S*,14*S*,17*S*)-17-Acetyl-10,13-dimethylhexadecahydro-1*H*-cyclopenta[*a*]phenanthren-3-yl 4-**

**(2-(2-((*E*)-(4-(dimethylamino)phenyl)diazenyl)thiazol-4-yl)acetamido)butanoate (9)**

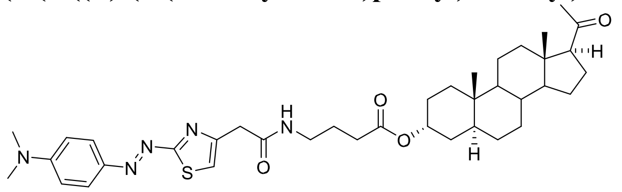

Starting from **45** (0.0847 mmol, 34 mg), compound **9** was synthesized according to general procedure. The crude material was purified by flash chromatography on silica gel using a gradient dichloromethane/acetone (0% to 17% in 20 column volumes). The product **9** (44 mg, 77%) was isolated as red solid.
The $^{1}$H and $^{13}$C NMR chemical shifts and corresponding signal assignments are reported in **Table S4** and **Table S5**, respectively. MS ESI: m/z 676.4 (6%, M + H), 698.4 (100%, M + Na). HR-MS (ESI) m/z: for $C_{38}H_{54}O_4N_5S$ [M + H] calcd, 676.3891; found, 676.3893.

**(3*S*,5*S*,8*R*,9*S*,10*S*,13*S*,14*S*,17*S*)-17-Acetyl-10,13-dimethylhexadecahydro-1*H*-cyclopenta[*a*]phenanthren-3-yl 4(2-(2-((*E*)-(4-(dimethylamino)phenyl)diazenyl)thiazol-4-yl)acetamido)butanoate (10)**

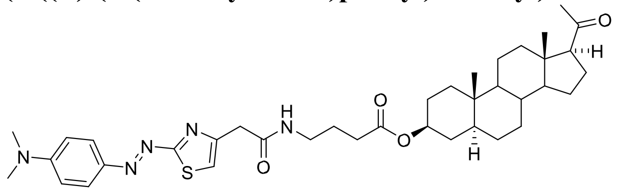

Starting from **46** (0.0847 mmol, 34 mg), compound **10** was synthesized according to general procedure. The crude material was purified by flash chromatography on silica gel using a gradient dichloromethane/acetone (0% to 17% in 20 column volumes). The product **10** (48 mg, 84%) was isolated as red solid.
The $^{1}$H and $^{13}$C NMR chemical shifts and corresponding signal assignments are reported in **Table S4** and **Table S5**, respectively. MS ESI: m/z 676.4 (13%, M + H), 698.4 (100%, M + Na). HR-MS (ESI) m/z: for $C_{38}H_{54}O_4N_5S$ [M + H] calcd, 676.3891; found, 676.3895.

**(3*S*,8*S*,9*S*,10*R*,13*S*,14*S*,17*S*)-17-Acetyl-10,13-dimethyl-2,3,4,7,8,9,10,11,12,13,14,15,16,17-tetradecahydro-1*H*cyclopenta[*a*]phenanthren-3-yl 4-(2-(2-((*E*)-(4-(dimethylamino)phenyl)diazenyl)thiazol-4yl)acetamido)butanoate (15)**

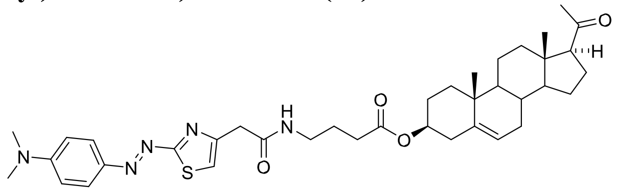

Starting from **49** (0.127 mmol, 51 mg), compound **15** was synthesized according to general procedure. The crude material was purified by flash chromatography on silica gel using a gradient dichloromethane/acetone (0% to 15% in 20 column volumes, then isocratic conditions for 4 column volumes). The product **15** (72 mg, 84%) was isolated as red solid.
The $^{1}$H and $^{13}$C NMR chemical shifts and corresponding signal assignments are reported in **Table S4** and **Table S5**, respectively. MS ESI: m/z 674.4 (100%, M + H), 696.4 (26%, M + Na). HR-MS (ESI) m/z: for $C_{38}H_{52}O_4N_5S$ [M + H] calcd, 674.3735; found, 674.3731. **(3*S*,8*R*,9*S*,10*R*,13*S*,14*S*)-10,13-Dimethyl-17-oxo-2,3,4,7,8,9,10,11,12,13,14,15,16,17-tetradecahydro-1*H*cyclopenta[*a*]phenanthren-3-yl 4-(2-(2-((*E*)-(4-(dimethylamino)phenyl)diazenyl)thiazol-4yl)acetamido)butanoate (16)**

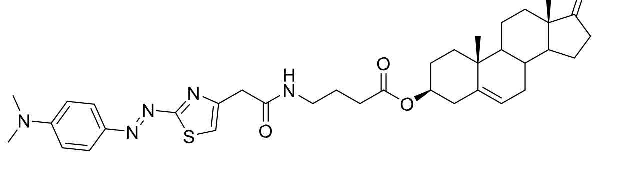

Starting from **52** (0.0857 mmol, 32 mg), compound **16** was synthesized according to general procedure. The crude material was purified by flash chromatography on silica gel using a gradient dichloromethane/acetone (0% to 17% in 20 column volumes). The product **16** (37 mg, 67%) was isolated as red solid.
The $^{1}$H and $^{13}$C NMR chemical shifts and corresponding signal assignments are reported in **Table S4** and **Table S5**, respectively. MS ESI: m/z 646.3 (31%, M + H), 668.3 (100%, M + Na). HR-MS (ESI) m/z: for $C_{36}H_{48}O_4N_5S$ [M + H] calcd, 646.3422; found, 646.3418.

**(3*S*,5*R*,8*S*,9*S*,10*S*,13*R*,14*S*,17*S*)-10,13,17-Trimethylhexadecahydro-1*H*-cyclopenta[*a*]phenanthren-3-yl 4-(2-(2((*E*)-(4-(dimethylamino)phenyl)diazenyl)thiazol-4-yl)acetamido)butanoate (17)**

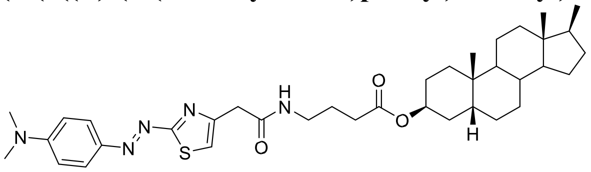

Starting from **56** (0.0799 mmol, 30 mg), compound **17** was synthesized according to general procedure. The crude material was purified by flash chromatography on silica gel using a gradient dichloromethane/acetone (0% to 17% in 20 column volumes). The product **17** (35 mg, 67%) was isolated as red solid.
The $^{1}$H and $^{13}$C NMR chemical shifts and corresponding signal assignments are reported in **Table S4** and **Table S5**, respectively. MS ESI: m/z 648.4 (27%, M + H), 670.4 (100%, M + Na). HR-MS (ESI) m/z: for $C_{37}H_{54}O_3N_5S$ [M + H] calcd, 648.3942; found, 648.3945.

#### 2.3.3 Synthesis of directly coupled steroid-TQ1 conjugates

##### 2.3.3.1 General procedure for the synthesis of directly coupled steroid-TQ1 conjugates

3-Hydroxysteroid (0.126 mmol, 40 mg) and DMAP (1 eq., 0.126 mmol, 15 mg) were placed in a round bottom flask equipped with a septum inlet. It was evacuated and refilled with argon two times. Freshly distilled dichloromethane (3.2 ml) was added upon stirring. It was followed by the addition of DIPEA (1.2 eq., 0.151 mmol, 26 µl) and EDCI.HCl (1.2 eq., 0.151 mmol, 29 mg). Finally, TQ1 acid (1.2 eq., 0.151 mmol, 44 mg) was added under an argon stream. The reaction mixture was stirred at room temperature overnight, in dark. The solvent was evaporated in vacuo.

**(3*R*,5*R*,8*R*,9*S*,10*S*,13*S*,14*S*,17*S*)-17-Acetyl-10,13-dimethylhexadecahydro-1*H*-cyclopenta[*a*]phenanthren-3-yl 2(2-((*E*)-(4-(dimethylamino)phenyl)diazenyl)thiazol-4-yl)acetate (11)**

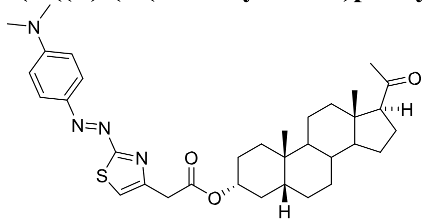

Starting from **3α5β-PRG** (**23**) (0.126 mmol, 40 mg), compound **11** was synthesized according to general procedure. The crude material was purified by flash chromatography on silica gel using a gradient petroleum ether/ethyl acetate (5% to 50% in 20 column volumes). The product **11** (39.6 mg, 54%) was isolated as deep red solid.
The $^{1}$H and $^{13}$C NMR chemical shifts and corresponding signal assignments are reported in **Table S6**. MS ESI: m/z 591.3 (100%, M + H). HR-MS (ESI) m/z: for $C_{34}H_{47}O_3N_4S$ [M + H] calcd, 591.3363; found, 591.3361.
**(3*S*,5*R*,8*R*,9*S*,10*S*,13*S*,14*S*,17*S*)-17-Acetyl-10,13-dimethylhexadecahydro-1*H*-cyclopenta[*a*]phenanthren-3-yl 2-(2-((*E*)-(4-(dimethylamino)phenyl)diazenyl)thiazol-4-yl)acetate (12)**

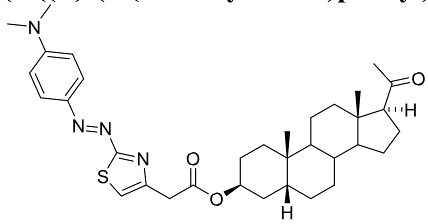

Starting from **3β5β-PRG** (**24**) (0.157 mmol, 50 mg), compound **12** was synthesized according to general procedure. The crude material was purified by flash chromatography on silica gel using a gradient petroleum ether/ethyl acetate (1% to 40% in 28 column volumes). The product **12** (23 mg, 25%) was isolated as deep red solid.
The $^{1}H$ and $^{13}C$ NMR chemical shifts and corresponding signal assignments are reported in **Table S6**. MS ESI: m/z 591.3 (93%, M + H), 613.3 (100%, M + Na). HR-MS (ESI) m/z: for $C_{34}H_{47}O_{3}N_{4}S$ [M + H] calcd, 591.3363; found, 591.3360.

**(3*R*,5*S*,8*R*,9*S*,10*S*,13*S*,14*S*,17*S*)-17-Acetyl-10,13-dimethylhexadecahydro-1*H*-cyclopenta[*a*]phenanthren-3-yl 2(2-((*E*)-(4-(dimethylamino)phenyl)diazenyl)thiazol-4-yl)acetate (13)**

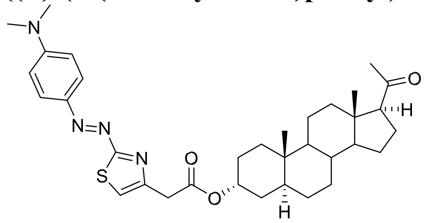

Starting from **3α5α-PRG** (**25**) (0.126 mmol, 40 mg), compound **13** was synthesized according to general procedure. The crude material was purified by flash chromatography on silica gel using a gradient dichloromethane/acetone (0% to 3% in 20 column volumes). The product **13** (50 mg, 68%) was isolated as deep red solid.
The $^{1}H$ and $^{13}C$ NMR chemical shifts and corresponding signal assignments are reported in **Table S6**. MS ESI: m/z 613.3 (100%, M + Na). HR-MS (ESI) m/z: for $C_{34}H_{47}O_{3}N_{4}S$ [M + H] calcd, 591.3363; found, 591.3365.

**(3*S*,5*S*,8*R*,9*S*,10*S*,13*S*,14*S*,17*S*)-17-Acetyl-10,13-dimethylhexadecahydro-1*H*-cyclopenta[*a*]phenanthren-3-yl 2(2-((*E*)-(4-(dimethylamino)phenyl)diazenyl)thiazol-4-yl)acetate (14)**

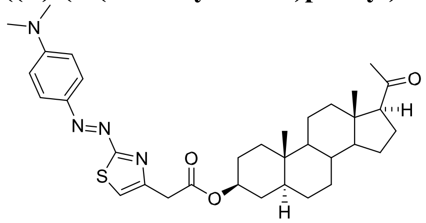

Starting from **3β5α-PRG** (**26**) (0.157 mmol, 50 mg), compound **14** was synthesized according to general procedure. The crude material was purified by flash chromatography on silica gel using a gradient petroleum ether/ethyl acetate (5% to 40% in 20 column volumes). The product **14** (23 mg, 25%) was isolated as deep red solid.
The $^{1}H$ and $^{13}C$ NMR chemical shifts and corresponding signal assignments are reported in **Table S6**. MS ESI: m/z 591.3 (100%, M + H), 613.3 (69%, M + Na). HR-MS (ESI) m/z: for $C_{34}H_{47}O_{3}N_{4}S$ [M + H] calcd, 591.3363; found, 591.3365.

#### 2.3.4 Synthesis of non-quenching analogues

##### 2.3.4.1 General procedure for the coupling with 2-(4-phenylpiperazino)-1,3-thiazole-4-carboxylic acid

Steroid amine (0.068 mmol, 30 mg) and DMAP (1 eq., 0.068 mmol, 8.3 mg) were placed in a round bottom flask equipped with a septum inlet. It was evacuated and refilled with argon two times. Freshly distilled dichloromethane (2 ml) was added upon stirring. It was followed by the addition of DIPEA (3.5 eq., 0.238 mmol, 41 μl) and EDCI.HCl (2.5 eq., 0.17 mmol, 33 mg). Finally, 2-(4-phenylpiperazino)-1,3-thiazole-4-carboxylic acid (2.5 eq., 0.17 mmol, 49 mg) was added under an argon stream. The reaction mixture was stirred at room temperature overnight, then the solvent was evaporated in vacuo. The obtained crude material was dissolved in 25 ml dichloromethane and was washed with water (2 x 10 ml). The collected organic phase was dried on $Na_2SO_4$, filtered and the solvent was evaporated.

**(3*R*,5*R*,8*R*,9*S*,10*S*,13*S*,14*S*,17*S*)-17-Acetyl-10,13-dimethylhexadecahydro-1*H*-cyclopenta[*a*]phenanthren-3-yl 4(2-(4-phenylpiperazin-1-yl)thiazole-4-carboxamido)butanoate (18)**

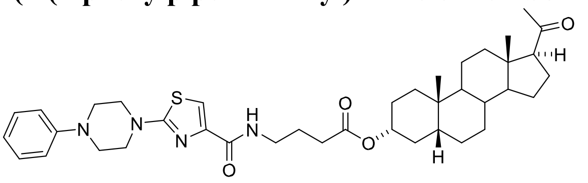

Starting from **43** (0.068 mmol, 30 mg), compound **18** was synthesized according to general procedure. The crude material was purified by flash chromatography on silica gel using a gradient of dichloromethane/acetone (0% to 15% in 20 column volumes). The product **18** was obtained as a white amorphous solid (28 mg, 61%).
The $^{1}$H and $^{13}$C NMR chemical shifts and corresponding signal assignments are reported in **Table S6**. MS ESI: m/z 697.4 (100%, M + Na). HR-MS (ESI) m/z: for $C_{39}H_{55}O_4N_4S$ [M + H] calcd, 675.3939; found, 675.3945.

**(3*R*,5*S*,8*R*,9*S*,10*S*,13*S*,14*S*,17*S*)-17-Acetyl-10,13-dimethylhexadecahydro-1*H*-cyclopenta[*a*]phenanthren-3-yl 4(2-(4-phenylpiperazin-1-yl)thiazole-4-carboxamido)butanoate (19)**

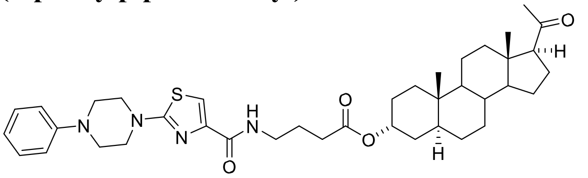

Starting from **45** (0.087 mmol, 35 mg), compound **19** was synthesized according to general procedure. The crude material was purified by flash chromatography on silica gel using a gradient of dichloromethane/acetone (0% to 15% in 20 column volumes). The product **19** was obtained as a white amorphous solid (32 mg, 54%).
The $^{1}$H and $^{13}$C NMR chemical shifts and corresponding signal assignments are reported in **Table S6**. MS ESI: m/z 675.4 (83%, M + H), 697.4 (100%, M + Na). HR-MS (ESI) m/z: for $C_{39}H_{55}O_4N_4S$ [M + H] calcd, 675.3939; found, 675.3937.

###### 2.3.4.2 Coupling of glutamate esters with Phe-Pip-Thiazo succinimidyl ester (57)

###### 2.3.4.2.1 Synthesis of Phe-Pip-Thiazo succinimidyl ester (57)

**2,5-Dioxopyrrolidin-1-yl 2-(4-phenylpiperazin-1-yl)thiazole-4-carboxylate (57)**

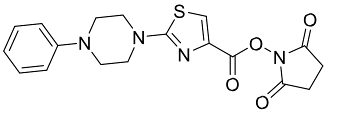

2-(4-Phenylpiperazino)-1,3-thiazole-4-carboxylic acid (0.52 mmol, 150 mg), *N*-hydroxysuccinimide (1.1 eq., 0.57 mmol, 66 mg) and DMAP (1.13 eq., 0.59 mmol, 72 mg) were placed in a round-bottom flask equipped with a septum inlet. It was evacuated and refilled with argon two times. Freshly distilled dichloromethane (6 ml) was added upon stirring. It was followed by the addition of EDCI.HCl (1.5 eq., 0.8 mmol, 153 mg) under an argon stream. The reaction mixture was stirred at room temperature overnight. After completion of the reaction, the solvent was evaporated in vacuo. Upon addition of dichloromethane (5 ml), precipitate was formed, which was filtered and further washed with dichloromethane (2x1 ml). The product **57** was obtained as off white solid (118 mg, 59%).
MS ESI: m/z 387.1 (35%, M + H), 409.1 (100%, M + Na). HR-MS (ESI) m/z: for $C_{18}H_{19}O_4N_4S$ [M + H] calcd, 387.1122; found, 387.1121.

###### 2.3.4.2.2 General procedure for the coupling of glutamate esters with Phe-Pip-Thiazo succinimidyl ester (57)

Glutamic acid ester (0.08 mmol, 35 mg) was placed in a round-bottom flask equipped with a septum inlet. It was evacuated and refilled with argon two times. Freshly distilled dichloromethane (2.8 ml) was added upon stirring. It was followed by the addition of DIPEA (1 eq., 0.08 mmol, 14 µl) and Phe-Pip-Thiazo succinimidyl ester (**57**) (1 eq., 0.08 mmol, 30 mg) was added under an argon stream. The reaction mixture was stirred at room temperature overnight. The solvent was evaporated in vacuo.

**(*S*)-5-(((3*S*,5*R*,8*R*,9*S*,10*S*,13*S*,14*S*,17*S*)-17-Acetyl-10,13-dimethylhexadecahydro-1*H*-cyclopenta[*a*]phenanthren3-yl)oxy)-5-oxo-4-(2-(4-phenylpiperazin-1-yl)thiazole-4-carboxamido)pentanoic acid (20)**

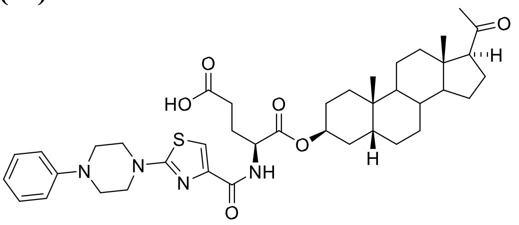

The crude material was purified by flash chromatography on silica gel using a gradient of dichloromethane/ethanol (1% to 6% in 20 column volumes). The product **20** was obtained as a white amorphous solid (51 mg, 91%).

**(*S*)-5-(((3*S*,5*S*,8*R*,9*S*,10*S*,13*S*,14*S*,17*S*)-17-Acetyl-10,13-dimethylhexadecahydro-1*H*-cyclopenta[*a*]phenanthren3-yl)oxy)-5-oxo-4-(2-(4-phenylpiperazin-1-yl)thiazole-4-carboxamido)pentanoic acid (21)**

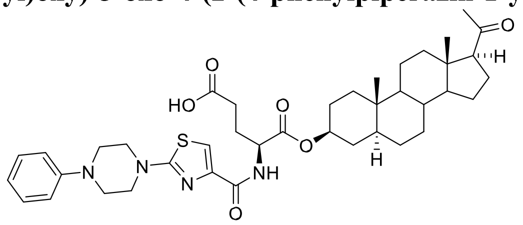

The crude material was purified by flash chromatography on silica gel using a gradient of dichloromethane/ethanol (1% to 10% in 20 column volumes). The product **21** was obtained as a white amorphous solid (49 mg, 88%).

**2.3.4.3 Coupling of compound 3α5α-PRG (25) with 2-(4-phenylpiperazino)-1,3-thiazole-4-carboxylic acid (3*R*,5*S*,8*R*,9*S*,10*S*,13*S*,14*S*,17*S*)-17-Acetyl-10,13-dimethylhexadecahydro-1*H*-cyclopenta[*a*]phenanthren-3-yl 2(4-phenylpiperazin-1-yl)thiazole-4-carboxylate (22)**

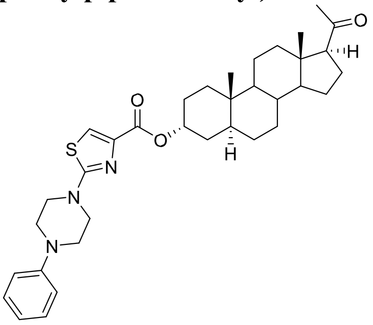

Steroid alcohol **3α5α-PRG** (**25**) (0.126 mmol, 40 mg), and DMAP (1 eq., 0.126 mmol, 15 mg) were placed in a round bottom flask equipped with a septum inlet. It was evacuated and refilled with argon two times. Freshly distilled dichloromethane (3.2 ml) was added upon stirring. It was followed by the addition of DIPEA (1.2 eq., 0.151 mmol, 26 µl) and EDCI.HCl (1.2 eq., 0.151 mmol, 29 mg). Finally, 2-(4-phenylpiperazino)-1,3-thiazole-4-carboxylic acid (1.2 eq., 0.151 mmol, 44 mg) was added under an argon stream. The reaction mixture was stirred at room temperature overnight, then the solvent was evaporated in vacuo. The obtained crude material was dissolved in 20 ml dichloromethane and was washed with water (2 x 5 ml). The collected organic phase was dried on $Na_2SO_4$, filtered and the solvent was evaporated. The crude material was purified by flash chromatography on silica gel using a gradient of dichloromethane/acetone (0% to 10% in 20 column volumes). The obtained material was further recrystallized from ethyl acetate/n-heptane 2/1 (v/v) mixture. The product **22** was obtained as a white crystalline material (20 mg, 27%). MS ESI: m/z 590.3 (54%, M + H), 612.3 (100%, M + Na). HR-MS (ESI) m/z: for $C_{35}H_{48}O_3N_3S$ [M + H] calcd, 590.3412; found, 590.3410.

#### 2.4 Proton and carbon chemical shifts (ppm) of steroid derivatives in DMSO-$d_6$ and determination of ring A stereochemistry

**Table S2. Proton chemical shifts of selected steroid derivatives in DMSO-$d_6$**

| Proton | 3α5β-PRG-Glu (35) [a] | 3α5βPRG-Glu-TQ1 (3) [b] | 3β5β-PRG-Glu (36) [c] | 3β5βPRG-Glu-TQ1 (4) [d] | 3β5α-PRG-Glu (38) [e] | 3β5αPRG-Glu-TQ1 (6) [f] | 3α5α-PRG-Glu (37) [g] | 3α5α-PRG-Glu-TQ1 (5) [h] | 3α5βPRG-GluDabcyl (1) [i] | 3β5βPRG-GluDabcyl (2) [j] |
|---|---|---|---|---|---|---|---|---|---|---|
| H-1□ | 1.78 | 1.74 | 1.51 | 1.04 | 1.02 | 0.945 | 1.17 | 1.35 | 1.08 | 1.04 |
| H-1□ | 1.03 | 1.00 | 1.27 | 1.34 | 1.69 | 1.60 | 1.47 | 1.46 | 1.38 | 1.34 |
| H-2□ | 1.38 | 1.31 | 1.51 | 1.50 | 1.46 | 1.36 | 1.61 | 1.58 | 1.36 | 1.52 |
| H-2□ | 1.62 | 1.58 | 1.56 | 1.50 | 1.74 | 1.67 | 1.65 | 1.62 | 1.61 | 1.52 |
| H-3 | 4.65 tt | 4.62 tt | 4.98 p | 4.96 p | 4.61 tt | 4.56 tt | 4.93 p | 4.90 p | 4.66 tt | 5.00 p |
| H-4□ | 1.81 | 1.74 | 1.98 | 1.95 | 1.52 | 1.39 | 1.37 | 1.09 | 1.80 | 1.95 |
| H-4□ | 1.48 | 1.42 | 1.34 | 1.32 | 1.30 | 1.19 | 1.50 | 1.39 | 1.51 | 1.32 |
| H-5 | 1.45 | 1.42 | 1.60 | 1.56 | 1.18 | 1.08 | 1.42 | 1.38 | 1.45 | 1.52 |
| H-6□ | 1.22 | 1.18 | 1.12 | 1.10 | 1.24 | 1.15 | 1.16 | 1.14 | 1.22 | 1.06 |
| H-6□ | 1.82 | 1.78 | 1.83 | 1.80 | 1.21 | 1.09 | 1.16 | 1.09 | 1.81 | 1.75 |
| H-7□ | 1.09 | 1.04 | 1.06 | 1.23 | 0.90 | 0.83 | 0.92 | 0.86 | 1.02 | 1.24 |
| H-7□ | 1.38 | 1.33 | 1.36 | 1.46 | 1.61 | 1.52 | 1.62 | 1.67 | 1.76 | 1.46 |
| H-8 | 1.36 | 1.33 | 1.36 | 1.33 | 1.32 | 1.19 | 1.33 | 1.28 | 1.35 | 1.31 |
| H-9 | 1.45 | 1.40 | 1.41 | 1.37 | 0.71 | 0.64 | 0.76 | 0.65 | 1.42 | 1.38 |
| H-11□ | 1.18 | 1.16 | 1.20 | 1.14 | 1.55 | 1.46 | 1.56 | 1.49 | 1.17 | 1.16 |
| H-11□ | 1.44 | 1.41 | 1.44 | 1.38 | 1.24 | 1.14 | 1.21 | 1.15 | 1.43 | 1.40 |
| H-12□ | 1.48 | 1.45 | 1.45 | 1.42 | 1.39 | 1.35 | 1.40 | 1.32 | 1.44 | 1.43 |
| H-12□ | 1.96 | 1.93 | 1.97 | 1.92 | 1.96 | 1.91 | 1.96 | 1.89 | 1.94 | 1.94 |
| H-14 | 1.26 | 1.23 | 1.23 | 1.20 | 1.12 | 1.08 | 1.15 | 1.06 | 1.23 | 1.20 |
| H-15□ | 1.59 | 1.56 | 1.59 | 1.58 | 1.59 | 1.55 | 1.59 | 1.53 | 1.51 | 1.47 |
| H-15□ | 1.11 | 1.08 | 1.11 | 1.09 | 1.12 | 1.07 | 1.12 | 1.06 | 1.10 | 1.01 |
| H-16□ | 1.56 | 1.54 | 1.56 | 1.54 | 1.54 | 1.54 | 1.54 | 1.46 | 1.54 | 1.54 |
| H-16□ | 2.01 | 2.00 | 2.02 | 2.00 | 2.01 | 1.99 | 2.01 | 1.96 | 2.00 | 2.00 |
| H-17 | 2.58 t | 2.55 t | 2.56 t | 2.55 t | 2.56 t | 2.54 t | 2.56 t | 2.47 t | 2.56 t | 2.54 t |
| H-18 | 0.50 s | 0.48 s | 0.50 s | 0.48 s | 0.51 s | 0.46 s | 0.51 s | 0.46 s | 0.49 s | 0.48 s |
| H-19 | 0.90 s | 0.88 s | 0.92 s | 0.88 s | 0.78 s | 0.68 s | 0.77 s | 0.73 s | 0.90 s | 0.76 s |
| H-21 | 2.05 s | 2.03 s | 2.05 s | 2.04 s | 2.05 s | 2.04 s | 2.05 s | 1.99 s | 2.03 s | 2.04 s |

**C(3)-substituents:**

[a] 3.32 dd (N-CH<); 1.82 m and 1.62 m (-CH2-); 2.31 m and 2.28 m (-CH2-).

[b] 4.23 dd (N−CH<); 1.97 m and 1.84 m (−$CH_2$−); 2.33 m (−$CH_2$−); 8.54 d (>NH−CO); 3.70 d and 3.66 d (−$CH_2$−); 7.36 t (S−CH=); 7.79 m (2x Ar−CH=); 6.87 m (2x Ar−CH=); 3.12 s (−$N(CH_3)_2$). [c] 3.37 dd (N−CH<); 1.83 m and 1.66 m (−$CH_2$−); 2.32 m and 2.26 m (−$CH_2$−);

[d] 4.22 dd (N−CH<); 1.96 m and 1.84 m (−$CH_2$−); 2.32 m (−$CH_2$−); 8.56 d (>NH−CO); 3.70 d and 3.66 d (−$CH_2$−); 7.36 t (S−CH=); 7.79 m (2x Ar−CH=); 6.87 m (2x Ar−CH=); 3.12 s (−$N(CH_3)_2$).

[e] 3.30 dd (N−CH<); 1.79 m and 1.61 m (−$CH_2$−); 2.37 m (−$CH_2$−).

[f] 4.20 dd (N−CH<); 1.94 m and 1.82 m (−$CH_2$−); 2.31 m (−$CH_2$−); 8.54 d (>NH−CO); 3.69 d and 3.62 d (−$CH_2$−); 7.36 t (S−CH=); 7.79 m (2x Ar−CH=); 6.87 m (2x Ar−CH=); 3.11 s (−$N(CH_3)_2$).

[g] 3.38 dd (N-CH<); 1.84 m and 1.66 m (-CH2-); 2.33 m and 2.28 m (-$CH_2$-).

[h] 4.27 ddd (N−CH<); 1.98 m and 1.86 m (−$CH_2$−); 2.32 m (−$CH_2$−); 8.58 d (>NH−CO); 3.71 s (−$CH_2$−); 7.39 t (S-CH=); 7.80 m (2x Ar−CH=);
6.87 m (2x Ar−CH=); 3.12 s (−$N(CH_3)_2$).

[i] 4.34 dd (N−CH<); 2.04 m and 1.38 m (−$CH_2$−); 1.24 m (−$CH_2$−); 9.71 br (>NH−CO); 8.05 m (2x Ar−CH=); 7.83 m (2x Ar−CH=); 7.82 m (2x Ar−CH=); 6.85 m (2x Ar−CH=); 3.08 s (−$N(CH_3)_2$).

[j] 4.33 ddd (N−CH<); 2.10 m and 2.01 m (−$CH_2$−); 2.40 m (−$CH_2$−); 8.70 bd (>NH−CO); 8.02 m (2x Ar−CH=); 7.83 m (4x Ar−CH=); 7.82 m (2x Ar−CH=); 6.86 m (2x Ar−CH=); 3.08 s (−$N(CH_3)_2$).

#### Table S3. Carbon-13 chemical shifts of selected steroid derivatives in DMSO-$d_6$

| Carbon | 3α5β-PRG-Glu (35) [a] | 3α5β-PRGGlu-TQ1 (3) [b] | 3β5β-PRG-Glu (36) [c] | 3β5βPRG-Glu-TQ1 (4) [d] | 3β5α-PRG-Glu (38) [e] | 3β5αPRG-Glu-TQ1 (6) [f] | 3α5α-PRG-Glu (37) [g] | 3α5α-PRG-Glu-TQ1 (5) [h] | 3α5βPRG-GluDabcyl (1) [i] | 3β5βPRG-GluDabcyl (2) [j] |
|---|---|---|---|---|---|---|---|---|---|---|
| **C-1** | 34.64 | 34.61 | 30.65 | 30.50 | 36.28 | 36.12 | 32.65 | 32.41 | 26.07 | 25.86 |
| **C-2** | 26.40 | 26.35 | 24.54 | 24.54 | 27.25 | 27.13 | 25.68 | 25.60 | 26.39 | 24.64 |
| **C-3** | 74.07 | 74.48 | 70.65 | 71.08 | 73.43 | 73.58 | 69.97 | 70.30 | 74.25 | 70.95 |
| **C-4** | 31.98 | 31.90 | 30.18 | 30.12 | 33.74 | 33.52 | 32.47 | 32.56 | 31.99 | 30.17 |
| **C-5** | 41.32 | 41.30 | 37.09 | 37.00 | 44.10 | 43.95 | 39.77 | 40.06 | 41.29 | 36.88 |
| **C-6** | 26.68 | 26.61 | 26.24 | 26.22 | 28.20 | 28.18 | 28.02 | 28.00 | 26.64 | 26.19 |
| **C-7** | 26.08 | 26.08 | 25.89 | 25.90 | 31.67 | 31.58 | 31.61 | 31.61 | 34.60 | 30.53 |
| **C-8** | 35.43 | 35.42 | 35.32 | 35.30 | 35.10 | 35.02 | 35.08 | 35.05 | 35.41 | 35.29 |
| **C-9** | 39.88 | 39.95 | 39.19 | 39.17 | 53.51 | 53.32 | 53.61 | 53.55 | 39.92 | 39.18 |
| **C-10** | 34.39 | 34.38 | 34.69 | 34.62 | 35.25 | 35.16 | 35.58 | 35.51 | 34.37 | 34.58 |
| **C-11** | 20.56 | 20.53 | 20.84 | 20.81 | 20.92 | 20.83 | 20.51 | 20.45 | 20.54 | 20.80 |
| **C-12** | 38.26 | 38.35 | 38.47 | 38.76 | 38.33 | 38.27 | 38.29 | 38.23 | 38.34 | 38.46 |
| **C-13** | 43.78 | 43.77 | 43.81 | 43.79 | 43.71 | 43.63 | 43.69 | 43.63 | 43.77 | 43.78 |
| **C-14** | 55.90 | 55.85 | 56.02 | 56.00 | 56.05 | 55.93 | 56.07 | 55.94 | 55.88 | 56.00 |
| **C-15** | 24.13 | 24.11 | 24.14 | 24.13 | 24.14 | 24.09 | 24.08 | 24.04 | 24.12 | 24.12 |
| **C-16** | 22.43 | 22.49 | 22.48 | 22.47 | 22.39 | 22.36 | 22.38 | 22.34 | 22.47 | 22.47 |
| **C-17** | 62.90 | 63.90 | 62.97 | 62.96 | 62.87 | 62.84 | 62.86 | 62.82 | 62.86 | 62.95 |
| **C-18** | 13.29 | 13.28 | 13.31 | 13.28 | 13.34 | 13.24 | 13.33 | 13.30 | 13.29 | 13.27 |
| **C-19** | 23.11 | 23.11 | 23.76 | 23.71 | 12.10 | 12.05 | 11.22 | 11.20 | 23.10 | 23.55 |
| **C-20** | 208.73 | 208.73 | 208.75 | 208.72 | 208.79 | 208.71 | 208.74 | 208.67 | 208.73 | 208.74 |
| **C-21** | 31.36 | 31.35 | 31.38 | 31.36 | 31.38 | 31.33 | 31.36 | 31.28 | 31.35 | 31.35 |

**C(3)-substituents:**

[a] 174.39 (O−CO−); 53.42 (>CH−N); 29.47 and 30.98 (2x −$CH_2$−); 174.62 (−COOH).

[b] 171.38 (O−CO−); 51.89 (N−CH<); 26.32 and 30.10 (2x −$CH_2$−); 173.85 (−COOH); 169.06 (NH−CO); 38.53 (−$CH_2$−); 150.88 (N−C=); 116.40 (S−CH=); 176.69 (N−C(=N)−S); 141.73 (Ar =C<); 126.34 (2x Ar−CH=); 112.13 (2x Ar−CH=); 153.97 (Ar =C<); 40.07 (−$N(CH_3)_2$).

[c] 174.46 (O−CO−); 53.58 (>CH−N); 29.52 and 31.06 (2x −$CH_2$−); 174.39 (−COOH).

[d] 171.46 (O−CO−); 52.06 (N−CH<); 29.34 and 30.23 (2x −$CH_2$−); 173.87 (−COOH); 169.02 (NH−CO−); 38.45 (−$CH_2$−); 150.80 (N−C=); 116.60 (S−CH=); 176.69 (N−(C=N)−S); 141.72 (Ar =C<); 126.30 (2x Ar−CH=); 112.13 (2x Ar−CH=); 153.96 (Ar =C<); 40.13 (−$N(CH_3)_2$).

[e] 174.73 (O−CO−); 53.39 (>CH−N); 29.58 and 31.11 (2x −$CH_2$−); 174.43 (−COOH).

[f] 171.29 (O−CO−); 52.05 (N−CH<); 26.27 and 30.95 (2x −$CH_2$−); 173.90 (−COOH); 169.03 (NH-CO−); 38.57 (−$CH_2$−); 150.87 (N−C=); 116.65 (S−CH=); 176.70 (N−C(=N)−S); 141.72 (Ar =C<); 126.39 (2x Ar−CH=); 112.07 (2x Ar−CH=); 153.92 (Ar =C<); 40.04 (−$N(CH_3)_2$).

[g] 174.37 (O−CO−); 53.52 (>CH−N); 29.56 and 31.06 (2x −$CH_2$−); 174.48 (−COOH).

[h] 171.23 (O−CO−); 52.06 (N−CH<); 26.44 and 30.28 (2x −$CH_2$−); 173.90 (−COOH); 169.05 (NH−CO−); 38.29 (−$CH_2$−); 150.71 (N−C−); 116.41 (S−CH=); 176.65 (N−C(=N)−S); 141.72 (Ar =C<); 126.32 (2x Ar−CH=); 112.12 (2x Ar−CH=); 153.95 (Ar =C<); 40.06 (−$N(CH_3)_2$). [i] 171.68 (O-CO−); 53.48 (N−CH<); 26.22 and 29.19 (2x −$CH_2$−); 174.00 (−COOH); 166.01 (NH-CO−); 134.27 (Ar =C<); 128.78 (2x Ar−CH=); 121.69 (2x Ar−CH=); 154.27 (Ar >C=); 142.82 (Ar >C=); 125.29 (2x Ar−CH=); 111.75 (2x Ar−CH=); 153.03 (Ar >C=); 40.04 (−$N(CH_3)_2$). [j] 171.26 (O-CO−); 52.84 (N−CH<); 25.83 and 30.53 (2x −$CH_2$−); 174.04 (−COOH); 166.38 (NH−CO−); 134.20 (Ar =C<); 128.80 (2x Ar−CH=); 121.62 (2x Ar−CH=); 154.33 (Ar >C=); 142.81 (Ar >C=); 125.31 (2x Ar−CH=); 111.77 (2x Ar−CH=); 153.06 (Ar >C=); 40.24
(−$N(CH_3)_2$).

#### Table S4. Proton chemical shifts of selected steroid derivatives in DMSO-$d_6$

| Proton | 3α5β-PRG-GABA (43) [a] | 3α5β-PRG-GABA-TQ1 (7) [b] | 3β5β-PRG-GABA-TQ1 (8) [c] | 3β5α-PRGGABA-TQ1 (10) [d] | 3α5α-PRGGABA-TQ1 (9) [e] | 3α5β-PRGGABA-PhePip-Thiazo (18) [f] | 3α5α-PRGGABA-PhePip-Thiazo (19) [g] | Pregnenolone-GABA-TQ1 (15) [h] | DHEA-GABA-TQ1 (16) [i] | 17β-Me-5βAND-3β-GABA-TQ1 (17) [j] |
|---|---|---|---|---|---|---|---|---|---|---|
| H-1□ | 1.08 | 1.06 | 1.025 | 0.95 | 1.14 | 1.06 | 1.15 | 1.03 | 1.03 | 1.02 |
| H-1□ | 1.38 | 1.35 | 1.33 | 1.61 | 1.43 | 1.35 | 1.44 | 1.78 | 1.78 | 1.34 |
| H-2□ | 1.38 | 1.35 | 1.46 | 1.68 | 1.61 | 1.35 | 1.62 | 1.73 | 1.73 | 1.47 |
| H-2□ | 1.63 | 1.60 | 1.50 | 1.36 | 1.61 | 1.60 | 1.62 | 1.49 | 1.50 | 1.47 |
| H-3 | 4.64 tt | 4.61 tt | 4.96 p | 4.54 tt | 4.90 p | 4.61 tt | 4.90 p | 4.43 m | 4.43 m | 4.95 p |
| H-4□ | 1.78 | 1.76 | 1.94 | 1.465 | 1.34 | 1.76 | 1.355 | 2.22 | 2.24 | 1.94 |
| H-4□ | 1.48 | 1.44 | 1.30 | 1.24 | 1.45 | 1.42 | 1.46 | 2.22 | 2.24 | 1.29 |
| H-5 | 1.44 | 1.41 | 1.56 | 1.10 | 1.38 | 1.41 | 1.39 | -- | -- | 1.56 |
| H-6□ | 1.21 | 1.18 | 1.09 | 1.18 | 1.15 | 1.19 | 1.14 | 5.30 m | 5.34 dt | 1.08 |
| H-6□ | 1.82 | 1.79 | 1.79 | 1.18 | 1.11 | 1.78 | 1.14 | -- | -- | 1.80 |
| H-7□ | 1.03 | 1.00 | 1.20 | 0.86 | 0.87 | 0.99 | 0.88 | 1.36 | 1.60 | 1.19 |
| H-7□ | 1.78 | 1.74 | 1.46 | 1.57 | 1.58 | 1.75 | 1.59 | 1.91 | 2.04 | 1.46 |
| H-8 | 1.36 | 1.34 | 1.32 | 1.27 | 1.30 | 1.34 | 1.30 | 1.53 | 1.58 | 1.34 |
| H-9 | 1.44 | 1.41 | 1.37 | 0.65 | 0.71 | 1.43 | 0.72 | 0.94 | 0.94 | 1.33 |
| H-11□ | 1.19 | 1.46 | 1.15 | 1.50 | 1.52 | 1.17 | 1.54 | 1.35 | 1.38 | 1.14 |
| H-11□ | 1.44 | 1.16 | 1.38 | 1.18 | 1.18 | 1.42 | 1.19 | 1.54 | 1.56 | 1.34 |
| H-12□ | 1.45 | 1.43 | 1.42 | 1.36 | 1.36 | 1.44 | 1.37 | 1.40 | 1.17 | 0.95 |
| H-12□ | 1.97 | 1.94 | 1.93 | 1.93 | 1.92 | 1.95 | 1.94 | 1.98 | 1.66 | 1.62 |
| H-14 | 1.23 | 1.21 | 1.19 | 1.09 | 1.09 | 1.23 | 1.10 | 1.12 | 1.25 | 0.99 |
| H-15□ | 1.59 | 1.57 | 1.57 | 1.57 | 1.55 | 1.57 | 1.56 | 1.59 | 1.48 | 1.53 |
| H-15□ | 1.11 | 1.09 | 1.08 | 1.09 | 1.09 | 1.09 | 1.10 | 1.12 | 1.84 | 1.03 |
| H-16□ | 1.56 | 1.54 | 1.54 | 1.53 | 1.51 | 1.56 | 1.52 | 1.55 | 2.00 | 1.15 |
| H-16□ | 2.01 | 2.00 | 2.00 | 2.00 | 1.99 | 2.00 | 1.99 | 2.03 | 2.40 | 1.74 |
| H-17 | 2.57 t | 2.54 t | 2.54 t | 2.54 t | 2.51 t | 2.56 t | 2.52 t | 2.55 t | -- | 1.35 m |
| H-18 | 0.50 s | 0.49 s | 0.48 s | 0.49 s | 0.49 s | 0.49 s | 0.50 s | 0.52 s | 0.79 s | 0.48 s |
| H-19 | 0.90 s | 0.88 s | 0.87 s | 0.72 s | 0.74 s | 0.88 s | 0.75 s | 0.92 d | 0.95 s | 0.89 s |
| H-21 | 2.06 s | 2.04 s | 2.05 s | 2.01 s | 2.03 s | 2.04 s | 2.03 s | 2.06 s | -- | 0.80 d |

**C(3)-substituents:**

[a] 2.39 m (−$CH_2$−); 1.79 m (−$CH_2$−); 2.80 m (−$CH_2$−); 7.86 br (−$NH_3$).

[b] 2.29 m (−$CH_2$−); 1.68 m (−$CH_2$−); 3.10 m (−$CH_2$−); 8.07 t (−NH−CO); 3.60 d (−$CH_2$−); 7.34 s (S−CH=); 7.79 m (2x Ar−CH=); 6.87 m (2x Ar−CH=); 3.12 s (−$N(CH_3)_2$).

[c] 2.30 m (−$CH_2$−); 1.68 m (−$CH_2$−); 3.10 m (−$CH_2$−); 8.06 t (−NH−CO); 3.59 d (−$CH_2$−); 7.34 t (S−CH=); 7.79 m (2x Ar−CH=); 6.87 m (2x Ar−CH=); 3.12 s (−$N(CH_3)_2$).

[d] 2.26 m (−$CH_2$−); 1.66 m (−$CH_2$−); 3.09 m (−$CH_2$−); 8.06 t (−NH-CO); 3.58 d (−$CH_2$−); 7.34 t (S−CH=); 7.79 m (2x Ar−CH=); 6.87 m (2x Ar−CH=); 3.12 s (−$N(CH_3)_2$).

[e] 2.32 m (−$CH_2$−); 1.68 m (−$CH_2$−); 3.12 m (−$CH_2$−); 8.09 t (−NH−CO); 3.60 d (−$CH_2$−); 7.34 t (S−CH=); 7.79 m (2x Ar−CH=); 6.87 m (2x Ar−CH=); 3.12 s (−$N(CH_3)_2$).

[f] 2.28 m (−$CH_2$−); 1.76 m (−$CH_2$−); 3.24 m (−$CH_2$−); 8.10 t (−NH−CO); 7.42 s (S−CH=); 3.61 m and 3.28 m (2x >N−$CH_2CH_2$-N<); 7.01 m (2x Ar−CH=); 7.25 (2x Ar−CH=); 6.83 m (Ar−CH=).

[g] 2.32 t (−$CH_2$−); 1.77 p (−$CH_2$-); 3.25 q (−$CH_2$−); 8.11 t (−NH−CO); 7.42 s (S−CH=); 3.61 m and 3.28 m (2x >N−$CH_2CH_2$−N<); 7.25 m (2x Ar−CH=); 7.01 (2x Ar−CH=); 6.83 m (Ar−CH=).

[h] 2.28 m (−$CH_2$−); 1.67 m (−$CH_2$−); 3.10 m (−$CH_2$−); 8.06 t (−NH−CO); 3.59 d (−$CH_2$−); 7.35 t (S−CH=); 7.79 m (2x Ar−CH=); 6.87 m (2x Ar−CH=); 3.12 s (−$N(CH_3)_2$).

[i] 2.28 m (−$CH_2$−); 1.67 m (−$CH_2$−); 3.10 m (−$CH_2$−); 8.06 t (−NH−CO); 3.59 d (−$CH_2$−); 7.35 t (S−CH=); 7.79 m (2x Ar−CH=); 6.87 m (2x Ar−CH=); 3.12 s (−$N(CH_3)_2$).

[j] 2.30 t (−$CH_2$−); 1.68 m (−$CH_2$−); 3.10 m (−$CH_2$−); 8.06 s (−NH−CO); 3.59 d (−$CH_2$−); 7.34 t (S−CH=); 7.79 m (2x Ar−CH=); 6.87 m (2x Ar−CH=); 3.12 s (−$N(CH_3)_2$).

**Table S5. Carbon-13 chemical shifts of steroid derivatives in DMSO-*d*$_6$**

| Carbon | 3α5β-PRG-GABA (43) [a] | 3α5β-PRGGABA-TQ1 (7) [b] | 3β5β-PRGGABA-TQ1 (8) [c] | 3β5α-PRGGABA-TQ1 (10) [d] | 3α5α-PRGGABA-TQ1 (9) [e] | 3α5β-PRGGABA-PhePip-Thiazo (18) [f] | 3α5α-PRGGABA-PhePip-Thiazo (19) [g] | Pregnenolone-GABA-TQ1 (15) [h] | DHEA-GABA-TQ1 (16) [i] | 17β-Me-5β-AND3β-GABA-TQ1 (17) [j] |
|---|---|---|---|---|---|---|---|---|---|---|
| **C-1** | 26.09 | 26.05 | 25.85 | 36.25 | 32.70 | 26.07 | 32.71 | 36.61 | 36.56 | 26.09 |
| **C-2** | 26.44 | 26.43 | 24.60 | 27.28 | 25.72 | 26.42 | 25.68 | 27.50 | 27.51 | 24.61 |
| **C-3** | 73.90 | 73.82 | 69.99 | 72.96 | 69.36 | 73.68 | 69.40 | 73.24 | 73.20 | 70.04 |
| **C-4** | 32.03 | 32.00 | 30.91 | 33.79 | 32.53 | 31.99 | 32.50 | 37.81 | 37.82 | 30.26 |
| **C-5** | 41.30 | 41.32 | 37.12 | 44.08 | 39.76 | 41.37 | 39.82 | 139.64 | 139.79 | 37.24 |
| **C-6** | 26.66 | 26.66 | 26.17 | 28.19 | 28.03 | 26.67 | 27.99 | 122.06 | 121.83 | 26.26 |
| **C-7** | 34.67 | 34.70 | 30.64 | 31.62 | 31.61 | 34.74 | 31.62 | 31.38 | 30.35 | 30.74 |
| **C-8** | 35.42 | 35.42 | 35.28 | 35.07 | 35.09 | 35.43 | 35.06 | 31.37 | 31.05 | 35.56 |
| **C-9** | 39.89 | 39.80 | 39.16 | 53.47 | 53.61 | 39.15 | 53.67 | 49.43 | 49.70 | 39.80 |
| **C-10** | 34.39 | 34.37 | 34.63 | 35.19 | 35.58 | 34.38 | 35.57 | 36.24 | 36.36 | 34.75 |
| **C-11** | 20.55 | 20.53 | 20.79 | 20.89 | 20.51 | 20.55 | 20.50 | 20.70 | 20.04 | 20.55 |
| **C-12** | 38.41 | 38.57 | 38.45 | 38.31 | 38.29 | 38.38 | 38.29 | 38.03 | 31.30 | 37.27 |
| **C-13** | 43.77 | 43.75 | 43.77 | 43.69 | 43.69 | 43.76 | 43.67 | 43.41 | 46.98 | 41.93 |
| **C-14** | 55.96 | 55.87 | 55.98 | 56.00 | 56.01 | 55.90 | 56.03 | 56.14 | 50.94 | 55.31 |
| **C-15** | 24.14 | 24.12 | 24.11 | 24.13 | 24.10 | 24.16 | 24.07 | 24.17 | 21.59 | 24.51 |
| **C-16** | 22.49 | 22.47 | 22.46 | 22.38 | 22.37 | 22.49 | 22.37 | 22.39 | 35.47 | 29.96 |
| **C-17** | 62.91 | 62.89 | 62.95 | 62.87 | 62.87 | 62.90 | 62.86 | 62.72 | 219.84 | 44.71 |
| **C-18** | 13.29 | 13.27 | 13.28 | 13.32 | 13.34 | 13.28 | 13.31 | 13.08 | 13.35 | 12.03 |
| **C-19** | 23.12 | 23.12 | 22.76 | 12.05 | 11.24 | 23.13 | 11.22 | 19.08 | 19.11 | 23.86 |
| **C-20** | 208.72 | 208.72 | 208.72 | 208.78 | 208.78 | 208.75 | 208.72 | 200.69 | -- | -- |
| **C-21** | 31.36 | 31.35 | 31.36 | 31.37 | 31.37 | 31.36 | 31.35 | 31.45 | -- | 14.01 |

**C(3)-substituents:**

[a] 171.80 (O−CO−); 30.78, 22.68 and 38.30 (3x −$CH_2$−).

[b] 172.28 (O−CO−); 31.38, 24.73 and 38.13 (3x −$CH_2$−); 168.80 (NH−CO−); 38.94 (−$CH_2$−); 151.17 (N−C=); 116.59 (S−CH=); 176.72 (N−C(=N)-S); 141.71 (Ar =C<); 126.32 (2x Ar−CH=); 112.13 (2x Ar−CH=); 153.96 (Ar =C<); 40.09 (−$N(CH_3)_2$).

[c] 172.17 (O−CO−); 31.63, 24.81 and 38.22 (3x −$CH_2$−); 168.79 (NH−CO); 38.96 (−$CH_2$−); 151.18 (N−C=); 116.61 (S−CH=); 176.75 (N−C(=N) −S); 141.72 (Ar =C<); 126.35 (2x Ar−CH=); 112.12 (2x Ar−CH=); 153.94 (Ar =C<); 40.06 (−$N(CH_3)_2$).

[d] 172.32 (O−CO−); 31.43, 24.72 and 38.17 (3x −$CH_2$−); 168.84 (NH−CO−); 38.98 (−$CH_2$−); 151.17 (N−C=); 116.67 (S−CH=); 176.77 (N−C(=N) −S); 141.74 (Ar =C<); 126.38 (2x Ar−CH=); 112.15 (2x Ar−CH=); 153.96 (Ar =C<); 40.08 (−$N(CH_3)_2$).

[e] 172.24 (O−CO−); 31.51, 24.81 and 38.13 (3x −$CH_2$−); 168.81 (NH−CO−); 38.95 (−$CH_2$−); 151.24 (N−C=); 116.53 (S−CH=); 176.75 (N−C(=N) −S); 141.73 (Ar =C<); 126.41 (2x Ar−CH=); 113.15 (2x Ar−CH=); 153.98 (Ar =C<); 40.09 (−$N(CH_3)_2$).

[f] 172.39 (O−CO−); 31.71, 24.90 and 38.17 (3x −$CH_2$−); 170.20 (NH−CO−); 161.02 (N−C=); 112.22 (S−CH=); 146.60 (N−C(=N)−S); 48.03 (2x N−$CH_2$−); 48.00 (2x N−$CH_2$−); 150.92 (Ar =C<); 116.32 (2x Ar−CH=); 129.20 (2x Ar−CH=); 119.75 (Ar−CH=).

[g] 172.23 (O−CO−); 31.76, 24.93 and 38.11 (3x −$CH_2$−); 170.22 (NH−CO−); 161.01 (N−C=); 112.22 (S−CH=); 146.62 (N−C(=N)−S); 48.01 (2x N−$CH_2$-); 48.03 (2x N-$CH_2$-); 150.92 (Ar =C<); 116.31 (2x Ar-CH=); 129.19 (2x Ar-CH=); 119.75 (Ar =CH-). [h] 172.23 (O-CO−); 31.33, 24.67 and 38.11 (3x -$CH_2$-); 168.83 (NH-CO); 38.98 (-$CH_2$-); 151.16 (N-C=); 116.63 (S-CH=); 176.76 (N-C(=N)−S); 141.72 (Ar =C<); 126.36 (2x Ar-CH=); 112.12 (2x Ar-CH=); 153.93 (Ar =C<); 40.05 (-$N(CH_3)_2$).

[i] 172.24 (O-CO−); 31.33, 24.69 and 38.11 (3x -$CH_2$-); 168.84 (NH-CO); 38.99 (-$CH_2$-); 151.16 (N-C=); 116.65 (S-CH=); 176.76 (N-C(=N)−S); 141.72 (Ar =C<); 126.36 (2x Ar−CH=); 112.13 (2x Ar−CH=); 153.95 (Ar =C<); 40.11 (−$N(CH_3)_2$). [j] 172.17 (O−CO−); 31.64, 24.81 and 38.22 (3x −$CH_2$−); 168.80 (NH−CO−); 38.96 (−$CH_2$−); 151.18 (N−C=); 116.61 (S−CH=); 176.75 (N−C(=N)−S); 141.72 (Ar =C<); 126.33 (2x Ar−CH=); 112.12 (2x Ar−CH=); 153.95 (Ar =C<); 40.06 (−$N(CH_3)_2$).

**Table S6. Proton and carbon-13 chemical shifts of steroid derivatives in DMSO-$d_6$**

| Proton | 3α5β-PRG-TQ1 (11) [a] | 3β5β-PRG-TQ1 (12) [b] | 3β5α-PRG-TQ1 (14) [c] | 3α5α-PRG-TQ1 (13) [d] | Carbon | 3α5β-PRG-TQ1 (11) [e] | 3β5β-PRG-TQ1 (12) [f] | 3β5α-PRG-TQ1 (14) [g] | 3α5α-PRG-TQ1 (13) [h] |
|---|---|---|---|---|---|---|---|---|---|
| H-1□ | 1.09 | 1.03 | 1.03 | 0.84 | C-1 | 26.08 | 25.86 | 36.29 | 32.21 |
| H-1□ | 1.38 | 1.33 | 1.70 | 1.24 | C-2 | 26.42 | 24.57 | 27.29 | 25.50 |
| H-2□ | 1.41 | 1.51 | 1.47 | 1.55 | C-3 | 74.44 | 70.86 | 73.82 | 70.22 |
| H-2□ | 1.66 | 1.51 | 1.78 | 1.55 | C-4 | 31.98 | 30.14 | 33.80 | 32.24 |
| H-3 | 4.69 tt | 5.00 p | 4.66 tt | 4.83 p | C-5 | 41.31 | 36.83 | 44.13 | 38.94 |
| H-4□ | 1.82 | 1.95 | 1.57 | 1.35 | C-6 | 26.65 | 26.16 | 28.21 | 27.91 |
| H-4□ | 1.52 | 1.34 | 1.33 | 1.30 | C-7 | 34.66 | 30.40 | 31.66 | 31.22 |
| H-5 | 1.45 | 1.49 | 1.18 | 1.04 | C-8 | 35.43 | 35.19 | 35.11 | 34.94 |
| H-6□ | 1.22 | 1.09 | 1.22 | 1.01 | C-9 | 39.87 | 39.17 | 53.51 | 53.32 |
| H-6□ | 1.81 | 1.75 | 1.22 | 1.01 | C-10 | 34.39 | 34.58 | 35.25 | 35.28 |
| H-7□ | 1.04 | 1.14 | 0.90 | 0.71 | C-11 | 20.55 | 20.80 | 20.93 | 20.29 |
| H-7□ | 1.78 | 1.44 | 1.61 | 1.41 | C-12 | 38.37 | 38.45 | 38.33 | 38.09 |
| H-8 | 1.36 | 1.31 | 1.31 | 1.10 | C-13 | 43.77 | 43.78 | 43.71 | 43.33 |
| H-9 | 1.44 | 1.38 | 0.70 | 0.33 | C-14 | 55.89 | 56.01 | 56.05 | 55.47 |
| H-11□ | 1.18 | 1.15 | 1.55 | 1.02 | C-15 | 24.13 | 24.11 | 24.14 | 23.79 |
| H-11□ | 1.44 | 1.40 | 1.30 | 1.32 | C-16 | 22.49 | 22.46 | 22.39 | 22.18 |
| H-12□ | 1.46 | 1.42 | 1.39 | 0.95 | C-17 | 62.89 | 62.94 | 62.87 | 62.55 |
| H-12□ | 1.96 | 1.94 | 1.96 | 1.73 | C-18 | 13.28 | 13.29 | 13.35 | 13.05 |
| H-14 | 1.24 | 1.20 | 1.12 | 0.63 | C-19 | 23.10 | 23.59 | 12.12 | 11.18 |
| H-15□ | 1.58 | 1.57 | 1.58 | 1.29 | C-20 | 208.74 | 208.74 | 208.80 | 208.56 |
| H-15□ | 1.10 | 1.08 | 1.12 | 0.90 | C-21 | 31.35 | 31.35 | 31.88 | 31.11 |
| H-16□ | 1.55 | 1.54 | 1.54 | 0.84 | | | | | |
| H-16□ | 2.01 | 2.00 | 2.00 | 1.82 | | | | | |
| H-17 | 2.56 t | 2.54 t | 2.56 t | 1.93 t | | | | | |
| H-18 | 0.50 s | 0.48 s | 0.50 s | 0.35 s | | | | | |
| H-19 | 0.90 s | 0.81 s | 0.78 s | 0.64 s | | | | | |
| H-21 | 2.05 s | 2.04 s | 2.05 s | 1.92 s | | | | | |

**C(3)-substituents:**

[a] 3.82 s (−$CH_2$−); 7.43 s (S−CH=); 7.80 m (2x Ar−CH=); 6.88 m (2x Ar−CH=); 3.12 s (−N($CH_3$)$_2$). [b] 3.83 d (−$CH_2$−); 7.43 t (S−CH=); 7.79 m (2x Ar−CH=); 6.88 m (2x Ar−CH=); 3.12 s (−N($CH_3$)$_2$). [c] 3.81 d (−$CH_2$−); 7.42 t (S−CH=); 7.80 m (2x Ar−CH=); 6.87 m (2x Ar−CH=); 3.12 s (−N($CH_3$)$_2$). [d] 3.85 d and 3.80 d (−$CH_2$−); 7.44 s (S−CH=); 7.84 m (2x Ar−CH=); 6.90 m (2x Ar−CH=); 3.08 s (−N($CH_3$)$_2$).

[e] 169.56 (O−CO−); 37.34 (−$CH_2$−); 149.32 (N−C=); 117.36 (S−CH=); 176.94 (N−C(=N)−S); 141.68 and 154.06 (2x Ar N−C=); 126.44 (2x Ar−CH=); 112.17 (2x Ar−CH=); 40.07 (−N($CH_3$)$_2$).

[f] 169.38 (O−CO−); 37.59 (−$CH_2$−); 149.65 (N−C=); 117.30 (S−CH=); 176.90 (N−C(=N)−S); 141.69 and 154.01 (2x Ar N−C=); 126.38 (2x Ar−CH=); 112.17 (2x Ar−CH=); 40.08 (−N($CH_3$)$_2$).

[g] 169.60 (O−CO−); 37.31 (−$CH_2$−); 149.34 (N−C=); 117.40 (S−CH=); 176.95 (N−C(=N)−S); 141.68 and 154.06 (2x Ar N−C=); 126.48 (2x Ar−CH=); 112.20 (2x Ar−CH=); 40.11 (−N($CH_3$)$_2$).

[h] 168.95 (O−CO−); 37.99 (−$CH_2$−); 150.02 (N−C=); 117.24 (S−CH=); 177.16 (N−C(=N)−S); 141.81 and 153.96 (2x Ar N−C=); 126.21 (2x Ar−CH=); 112.10 (2x Ar−CH=); 39.72 (−N($CH_3$)$_2$).

#### Determination of stereochemistry of ring A

According to the configuration at carbon C-3 and C-5 most of prepared steroids (except 5-en derivatives **15** and **16**) belong to one of four stereochemical groups. Combination of NMR parameters observed for H-3, H-19 and C-19 (see **Table S7**) allow to distinguish four groups and determine stereochemistry of ring A.

**Table S7. Values of selected NMR parameters observed for steroid derivatives**

| Parameter | 3α-OR, 5α-H | 3β-OR, 5α-H | 3α-OR, 5β-H | 3β-OR, 5β-H |
|---|---|---|---|---|
| H-3: chem.shift [ppm] | 4.83 – 4.93 | 4.54 – 4.72 | 4.60 – 4.69 | 4.91 – 5.00 |
| multiplicity | pentet | triplet of triplets | triplet of triplets | pentet |
| *J* (H.H) in Hz | 2.8 | 11.1 and 4.8 | 11.1 and 4.8 | 2.9 |
| H-19: chem.shift [ppm] | 0.64 – 0.77 | 0.46 – 0.51 | 0.88 – 0.90 | 0.76 – 0.92 |
| C-19: chem.shift [ppm] | 11.18 – 11.24 | 12.05 – 12.12 | 23.10 – 23.13 | 22.76 – 23.86 |

#### 2.5 $^{1}$H and $^{13}$C NMR spectra of synthesized compounds

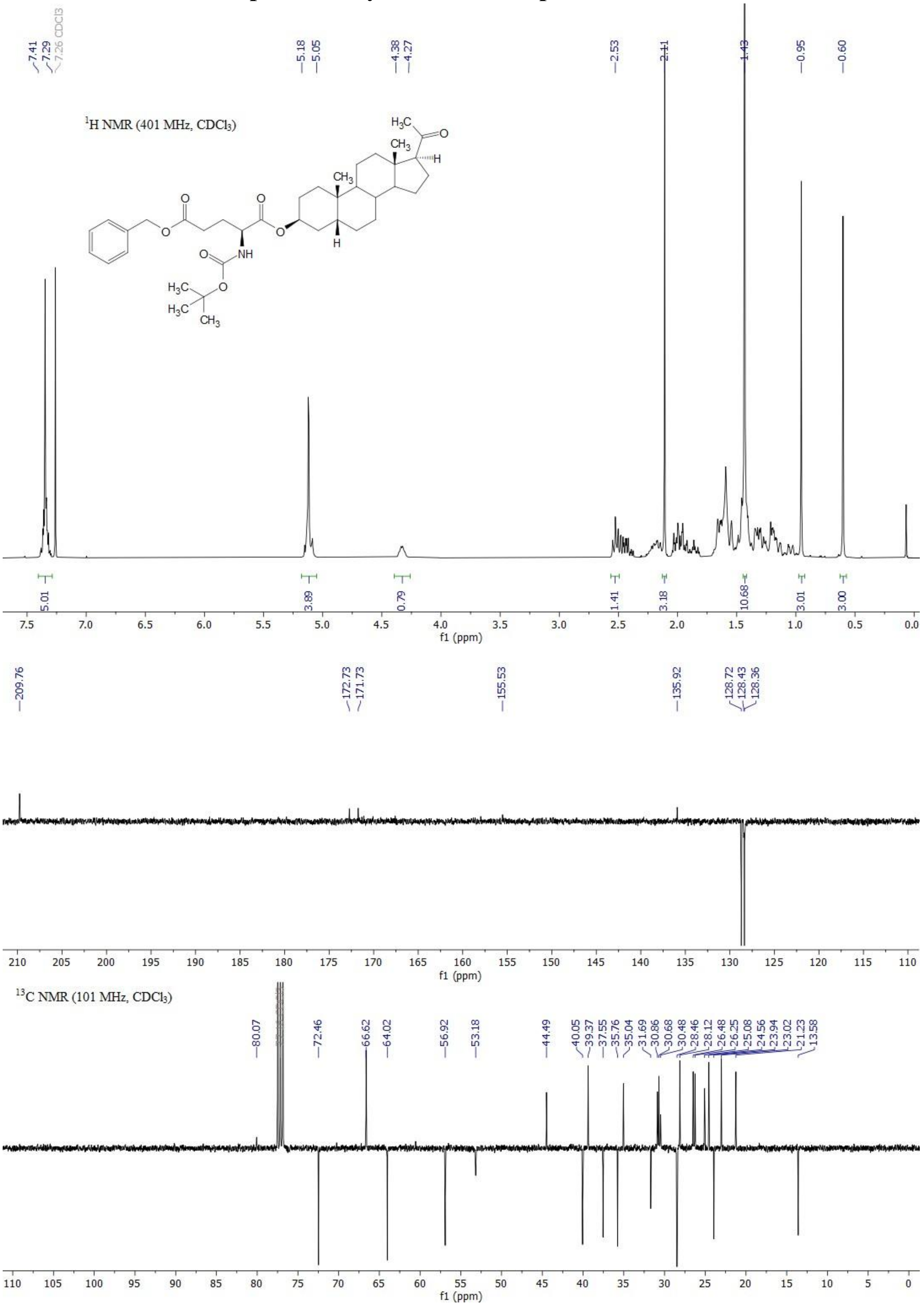

**Figure S4.** $^{1}$H NMR (top) and $^{13}$C APT NMR (bottom) spectra of compound **28**.

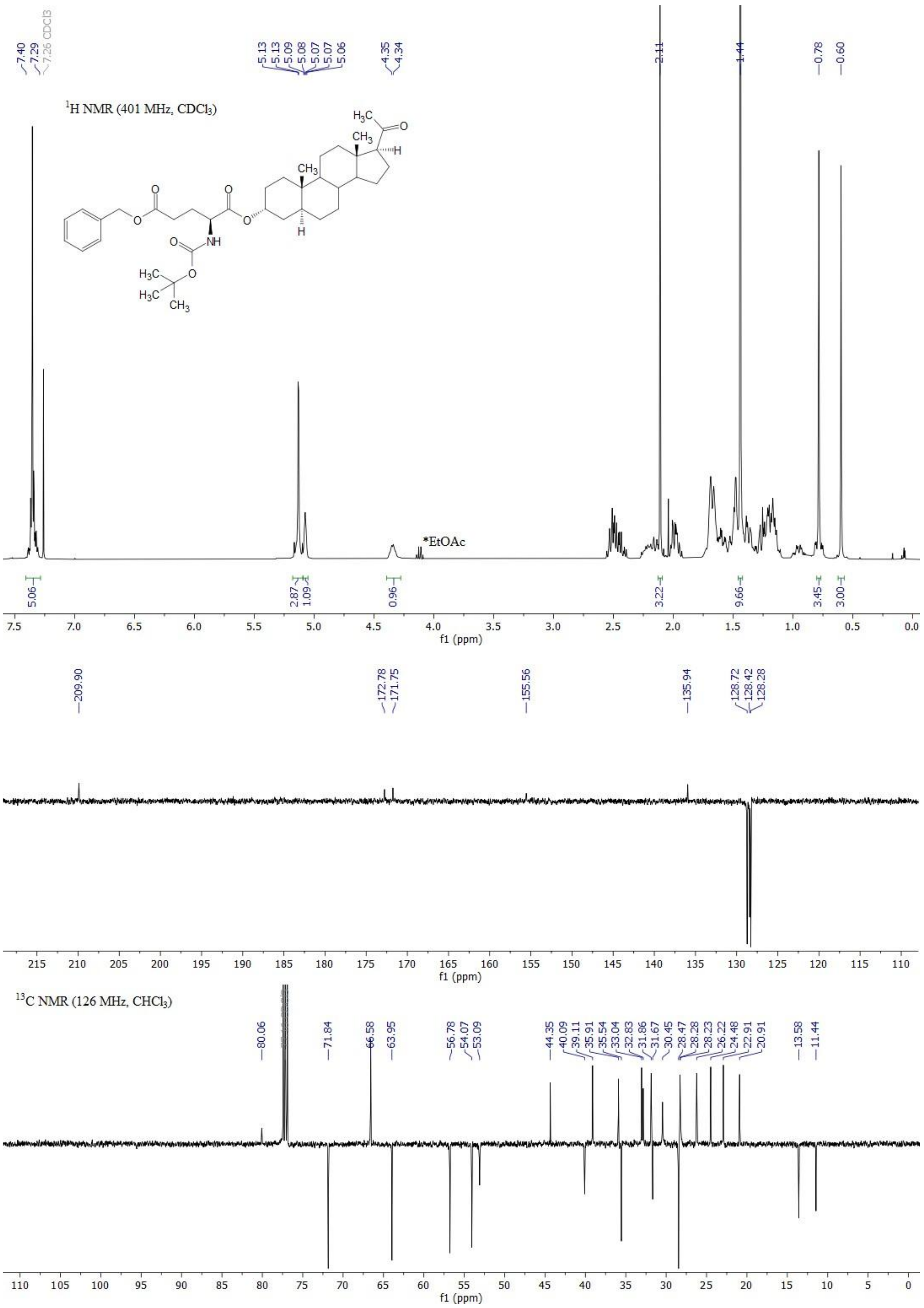

**Figure S5.** $^1$H NMR (top) and $^{13}$C APT NMR (bottom) spectra of compound **29**.

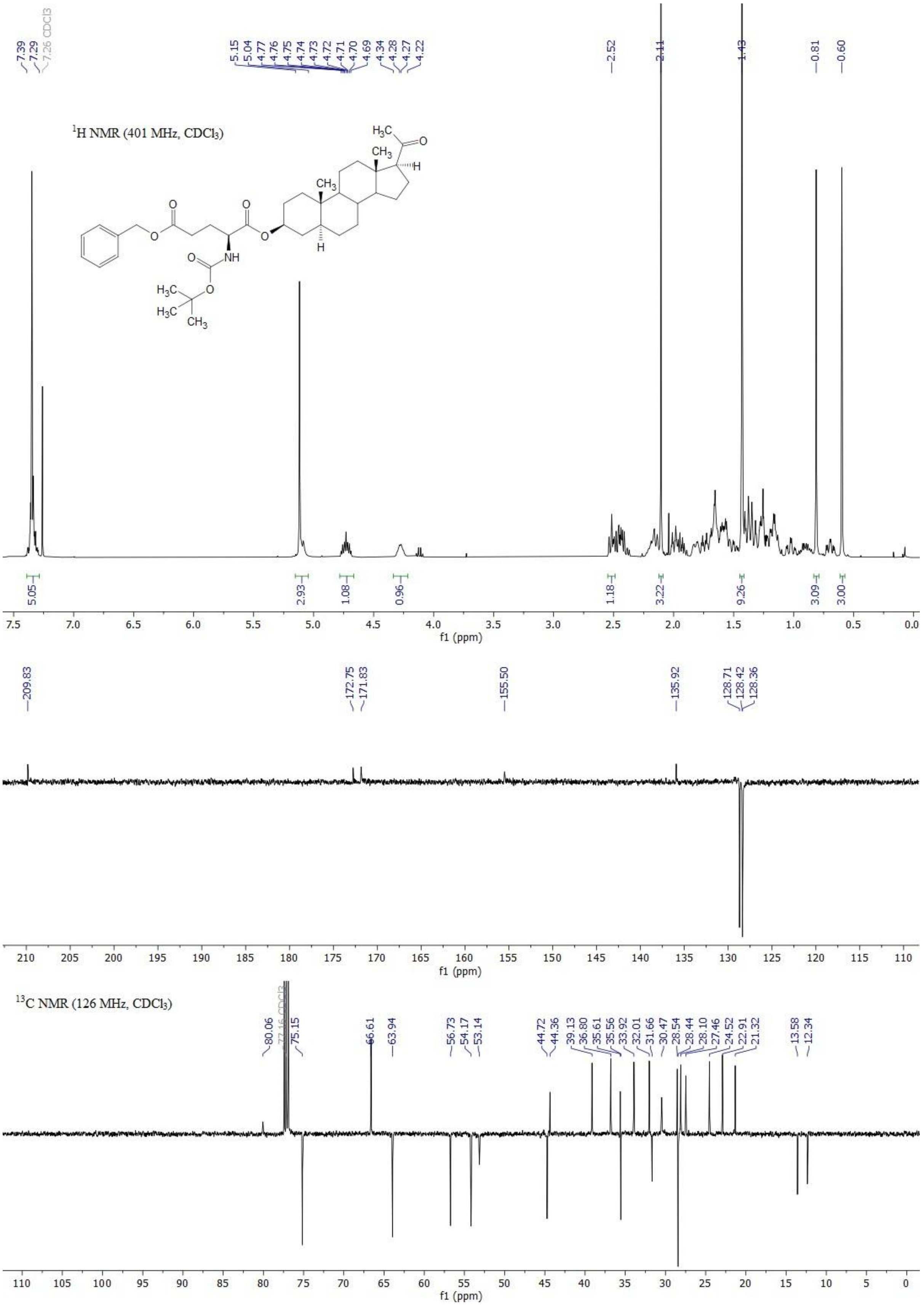

**Figure S6.** $^{1}H$ NMR (top) and $^{13}C$ APT NMR (bottom) spectra of compound **30**.

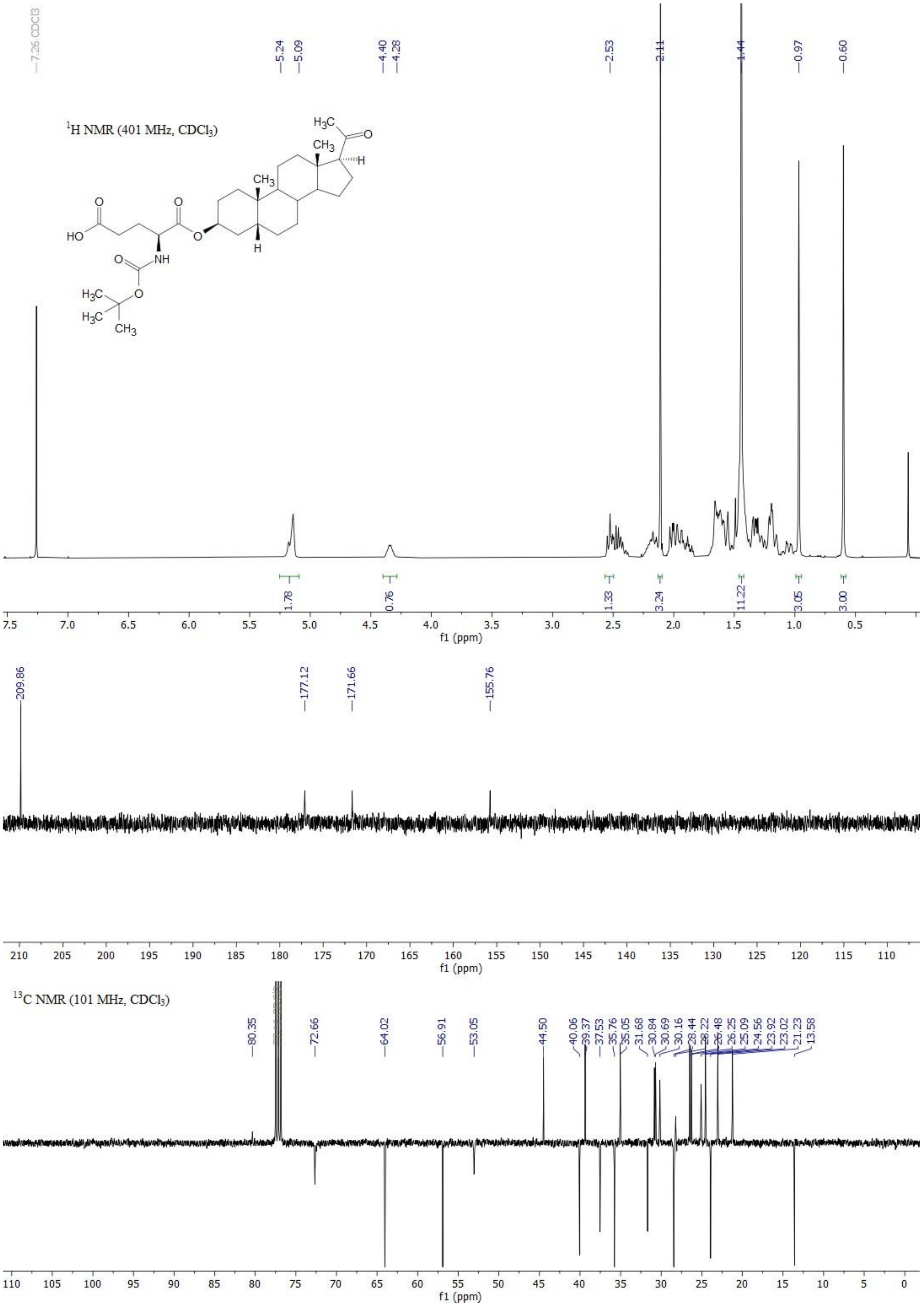

**Figure S7.** $^{1}H$ NMR (top) and $^{13}C$ APT NMR (bottom) spectra of compound **32**.

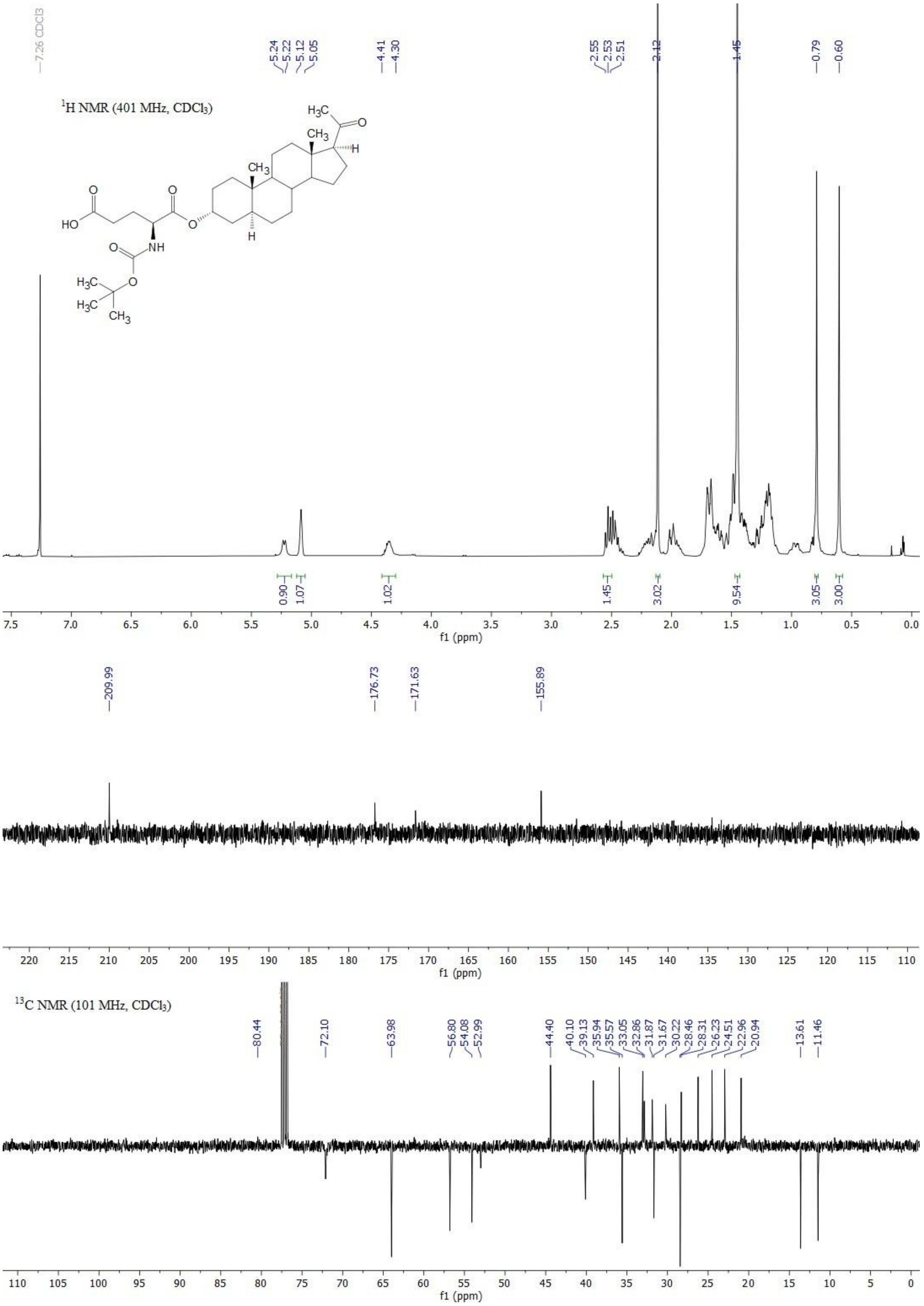

**Figure S8.** $^{1}H$ NMR (top) and $^{13}C$ APT NMR (bottom) spectra of compound **33**.

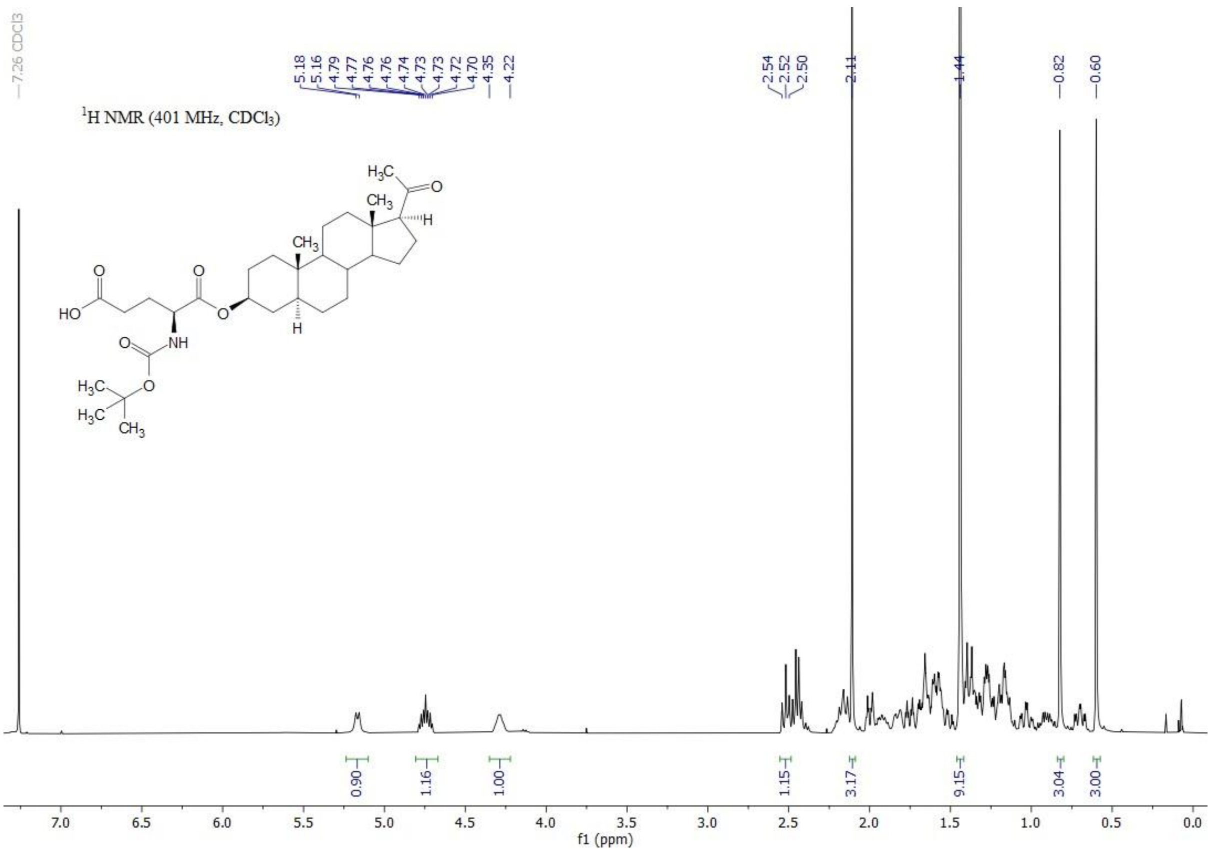

1H NMR (401 MHz, CDCl3)
7.26 CDCl3
5.18 5.16 4.79 4.77 4.76 4.76 4.74 4.73 4.73 4.72 4.70 4.35 4.22
2.54 2.52 2.50 2.11 1.44 0.82 0.60
0.90 1.16 1.00 1.15 3.17 9.15 3.04 3.00
f1 (ppm)

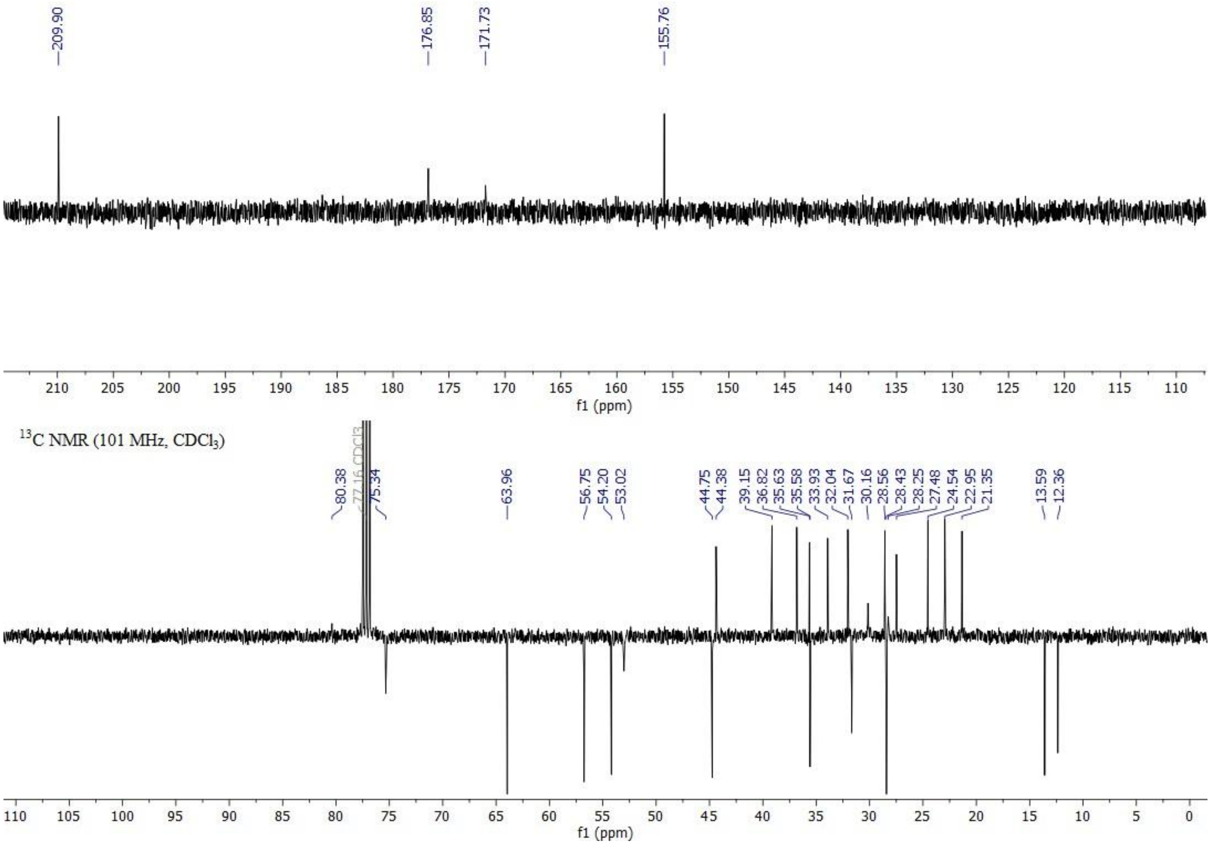

209.90 176.85 171.73 155.76
f1 (ppm)
13C NMR (101 MHz, CDCl3)
80.38 77.16 CDCl3 75.34 63.96 56.75 54.20 53.02 44.75 44.38 39.15 36.82 35.63 35.58 33.93 32.04 31.67 30.16 28.56 28.43 28.25 27.48 24.54 22.95 21.35 13.59 12.36
f1 (ppm)

**9.** $^{1}$H NMR (top) and $^{13}$C APT NMR (bottom) spectra of compound **34**.

**Figure S** $^{1}$H NMR (top) and $^{13}$

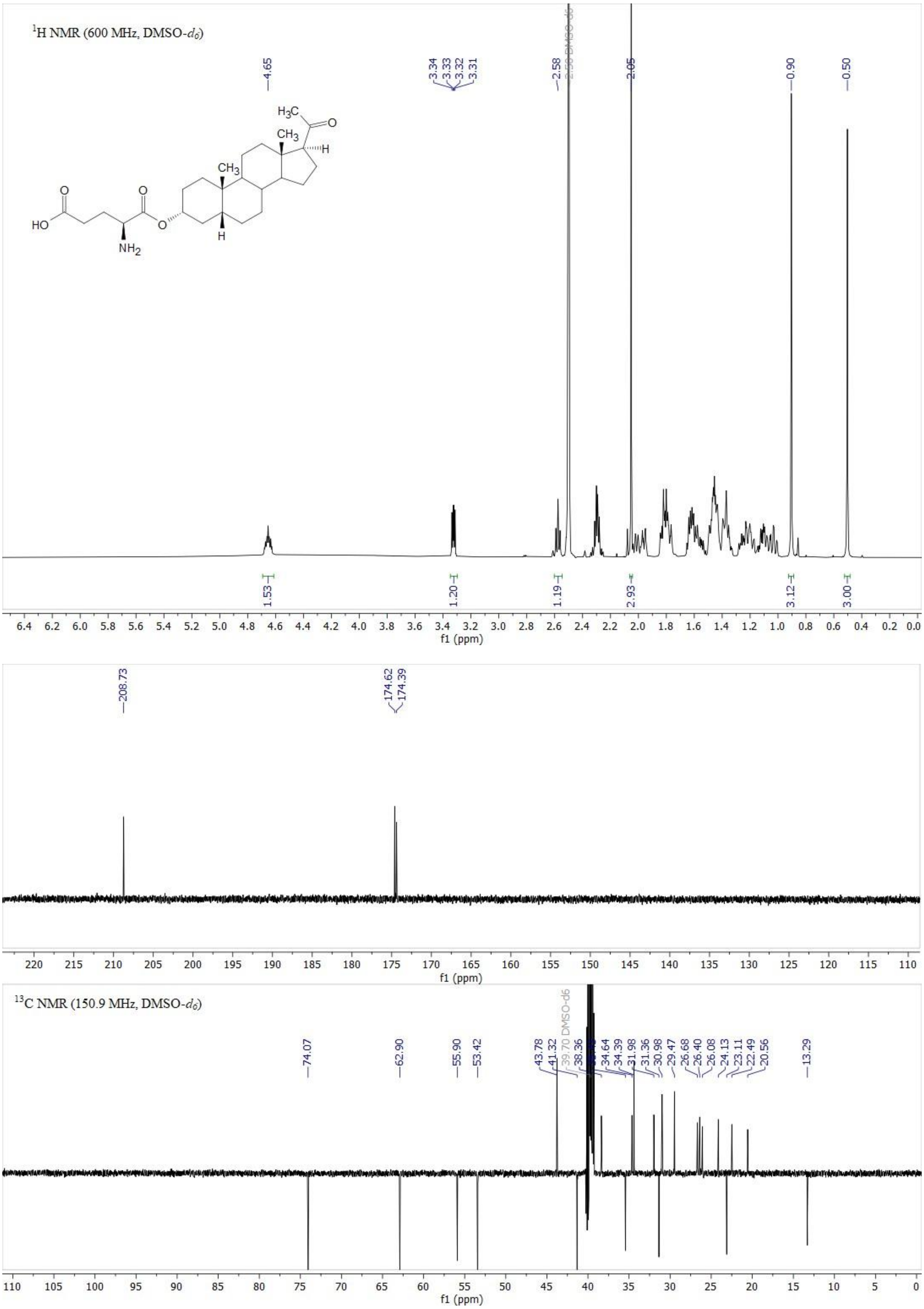

**Figure S10.** $^{1}H$ NMR (top) and $^{13}C$ APT NMR (bottom) spectra of compound **35**.

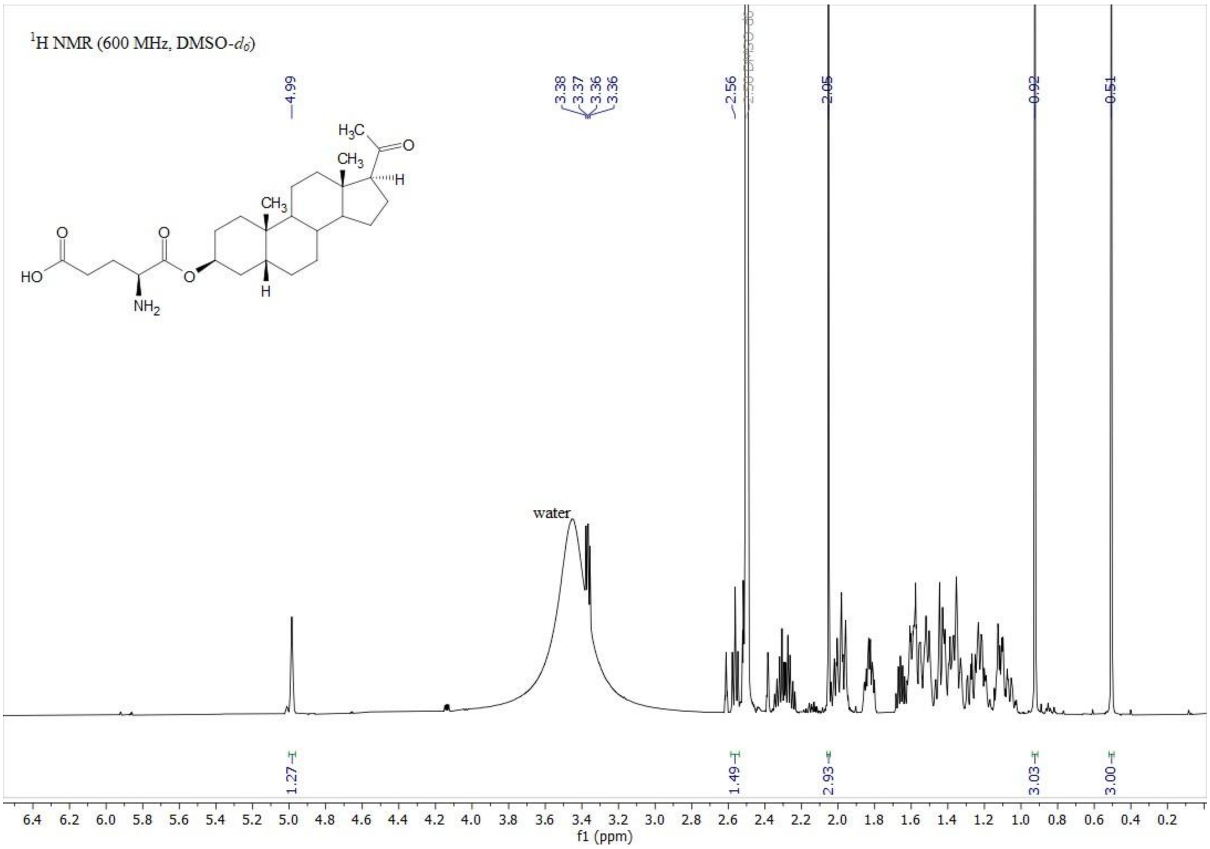

$^1$H NMR (600 MHz, DMSO-$d_6$)
water
f1 (ppm)

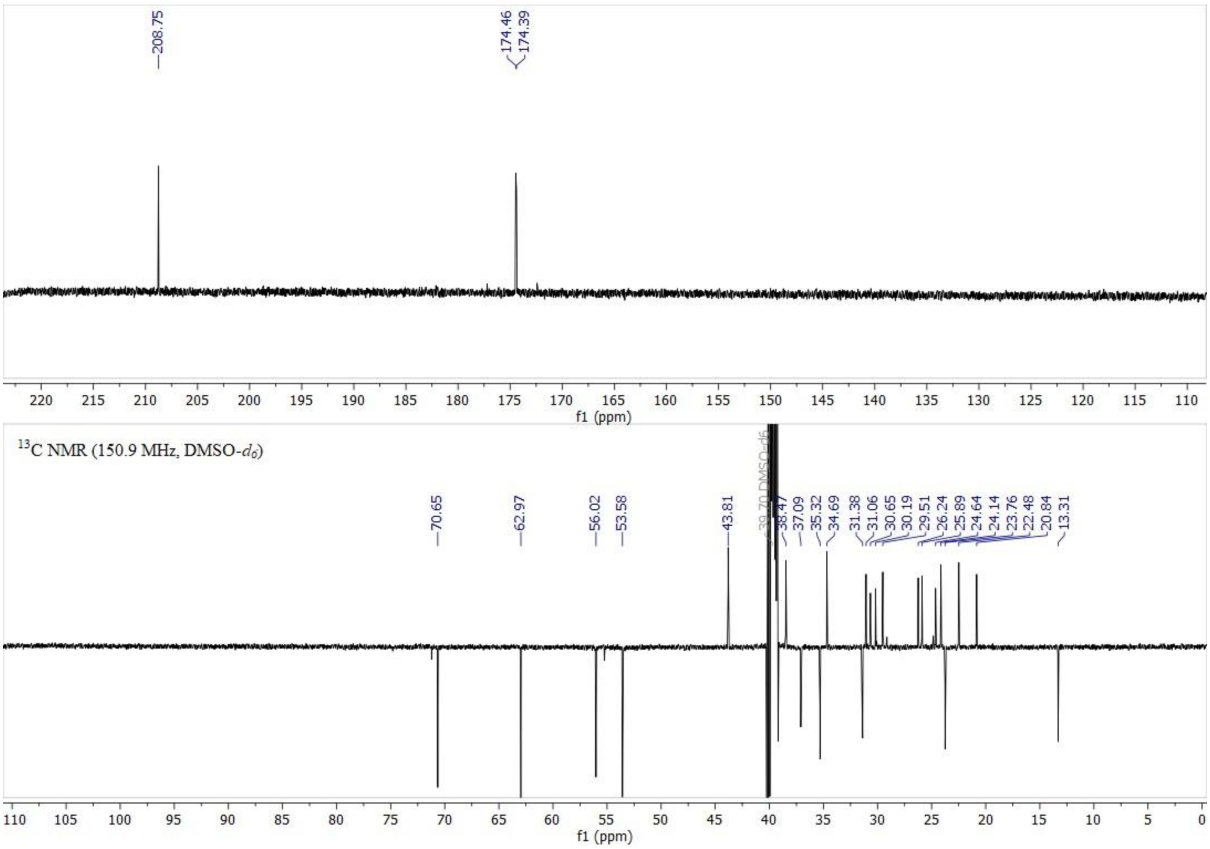

$^{13}$C NMR (150.9 MHz, DMSO-$d_6$)
f1 (ppm)

**11.** C APT NMR (bottom) spectra of compound **36**.

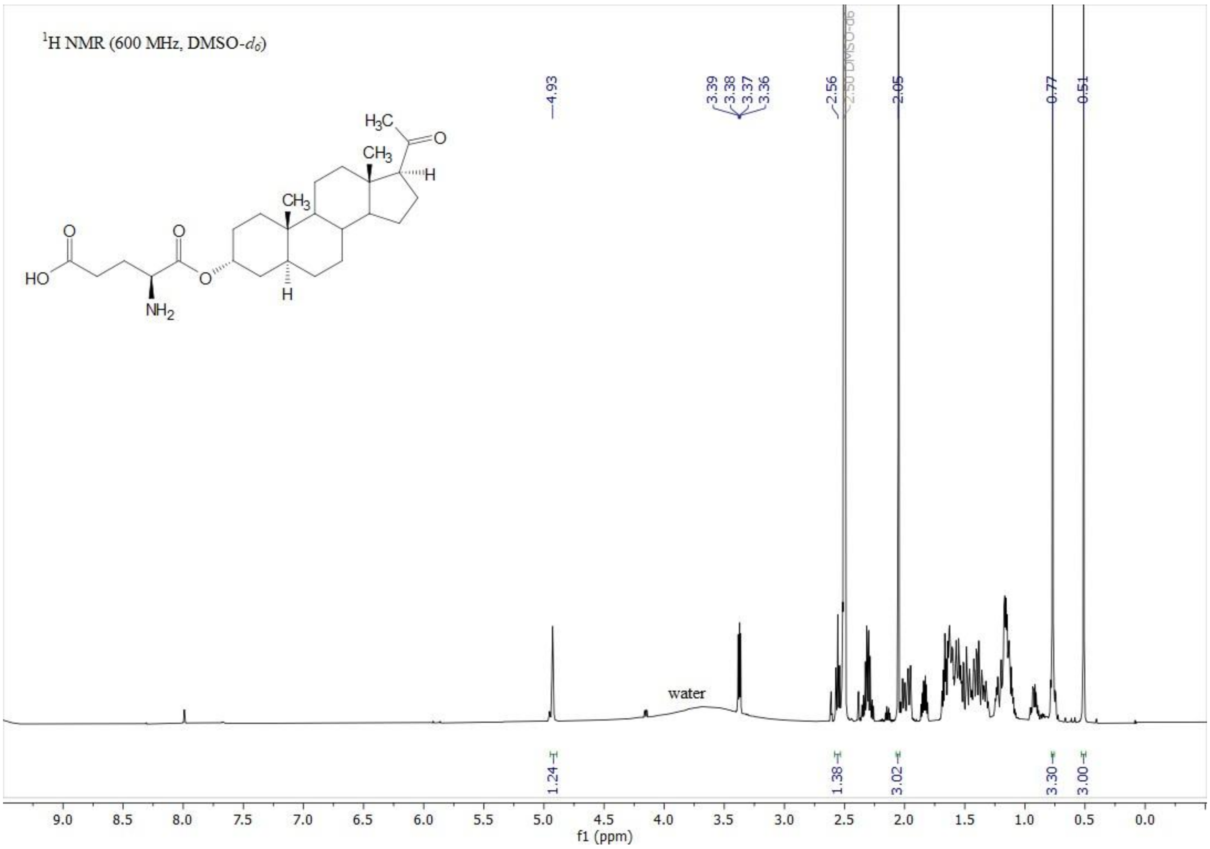

1H NMR (600 MHz, DMSO-d6)
H3C
CH3
H
CH3
O
O
O
HO
NH2
O
H
4.93
3.39
3.38
3.37
3.36
2.56
2.50 DMSO-d6
2.05
0.77
0.51
water
1.24
1.38
3.02
3.30
3.00
9.0
8.5
8.0
7.5
7.0
6.5
6.0
5.5
5.0
4.5
4.0
3.5
3.0
2.5
2.0
1.5
1.0
0.5
0.0
f1 (ppm)

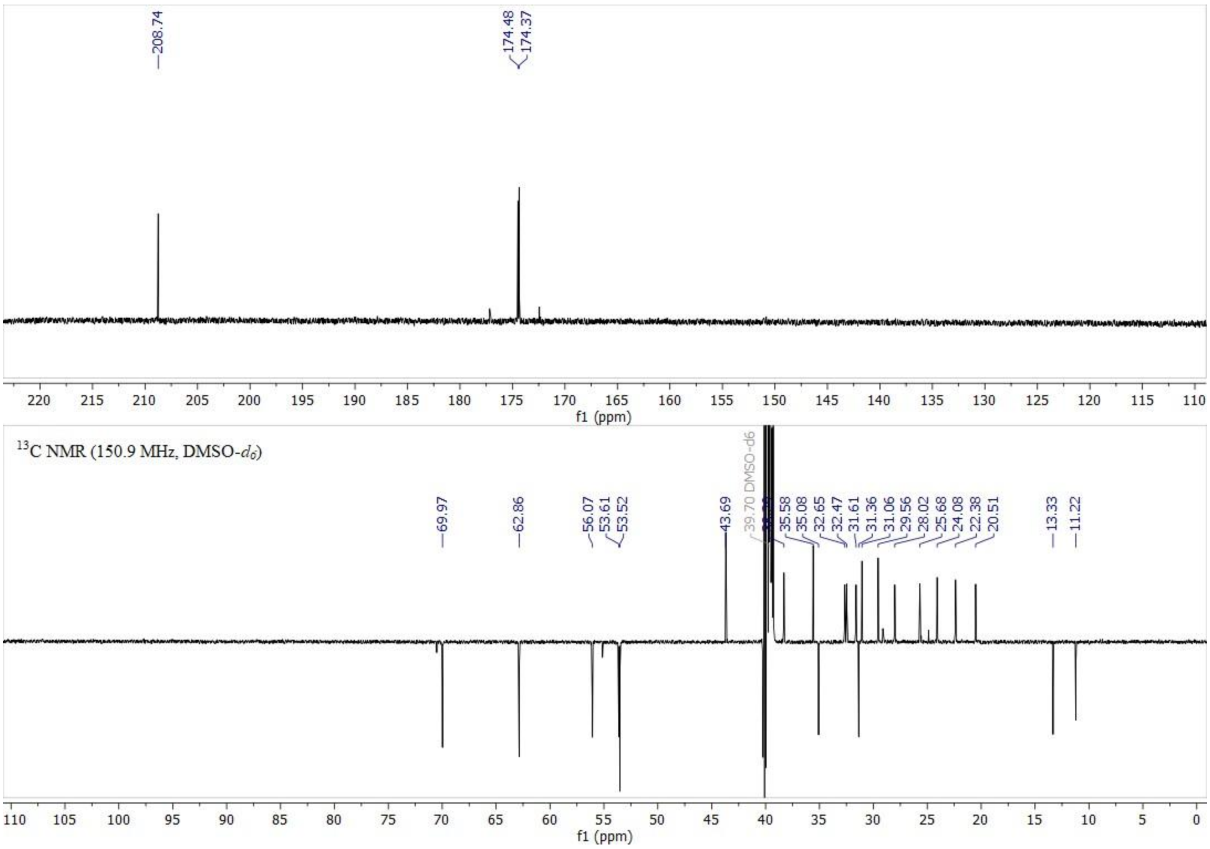

208.74
174.48
174.37
220
215
210
205
200
195
190
185
180
175
170
165
160
155
150
145
140
135
130
125
120
115
110
f1 (ppm)
13C NMR (150.9 MHz, DMSO-d6)
69.97
62.86
56.07
53.61
53.52
43.69
39.70 DMSO-d6
35.58
35.08
32.65
32.47
31.61
31.36
31.06
29.56
28.02
25.68
24.08
22.38
20.51
13.33
11.22
110
105
100
95
90
85
80
75
70
65
60
55
50
45
40
35
30
25
20
15
10
5
0
f1 (ppm)

**12.** C APT NMR (bottom) spectra of compound **37**.

**Figure S** $^{1}$H NMR (top) and $^{13}$

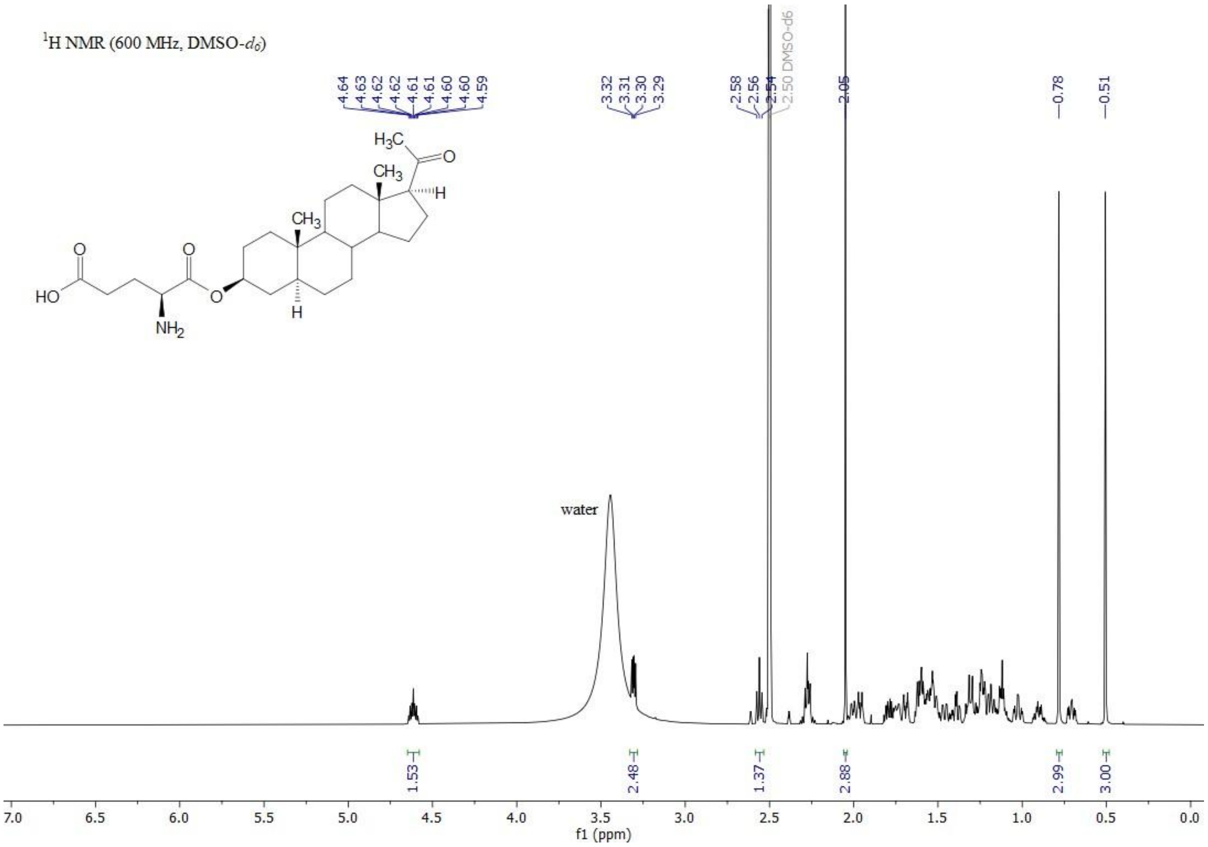

$^1$H NMR (600 MHz, DMSO-$d_6$)
4.64 4.63 4.62 4.62 4.61 4.61 4.60 4.60 4.59
3.32 3.31 3.30 3.29
2.58 2.56 2.54
2.50 DMSO-d6
2.05
0.78
0.51
H3C
CH3
H
CH3
O
HO
NH2
water
1.53
2.48
1.37
2.88
2.99
3.00
f1 (ppm)

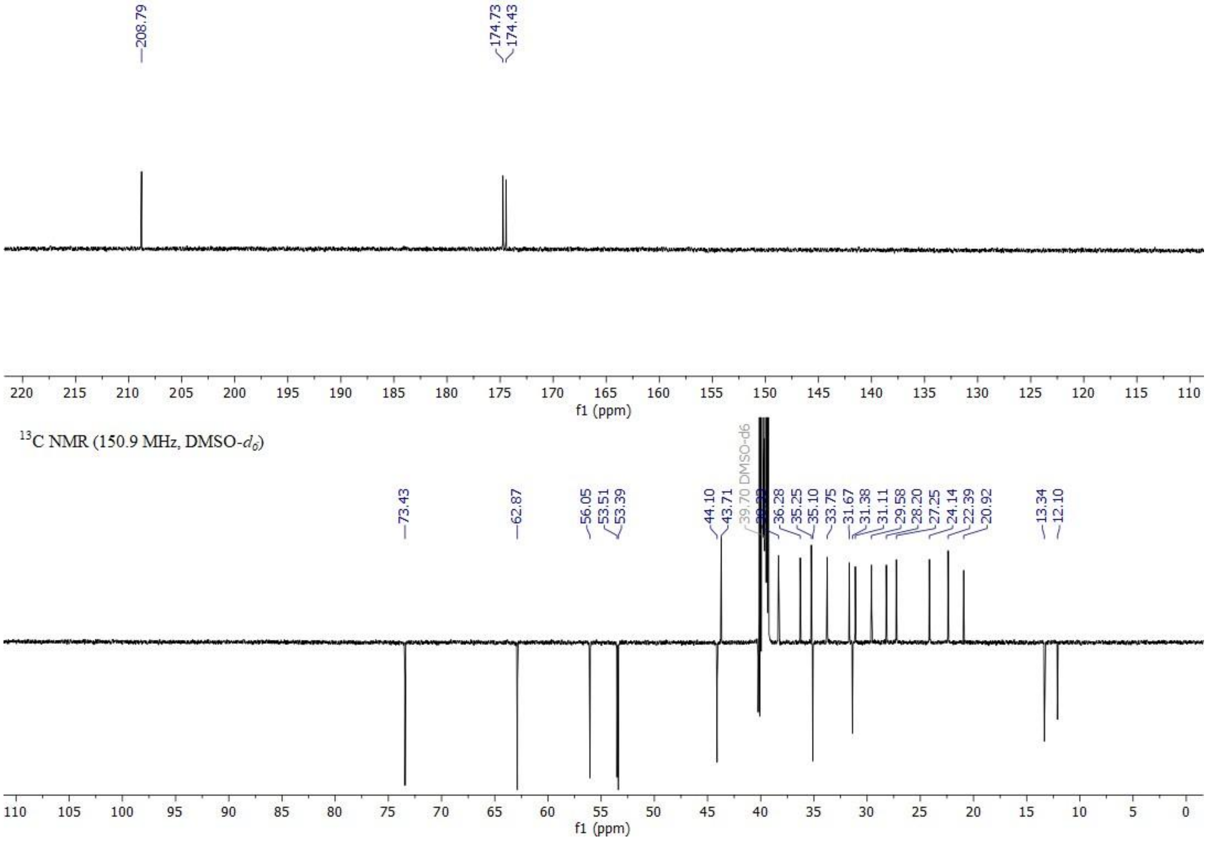

208.79
174.73 174.43
f1 (ppm)
$^{13}$C NMR (150.9 MHz, DMSO-$d_6$)
73.43
62.87
56.05 53.51 53.39
44.10 43.71
39.70 DMSO-d6
36.28 35.25 35.10 33.75 31.67 31.38 31.11 29.58 28.20 27.25 24.14 22.39 20.92
13.34 12.10
f1 (ppm)

**13.** C APT NMR (bottom) spectra of compound **38**.

**Figure S** $^{1}$H NMR (top) and $^{13}$

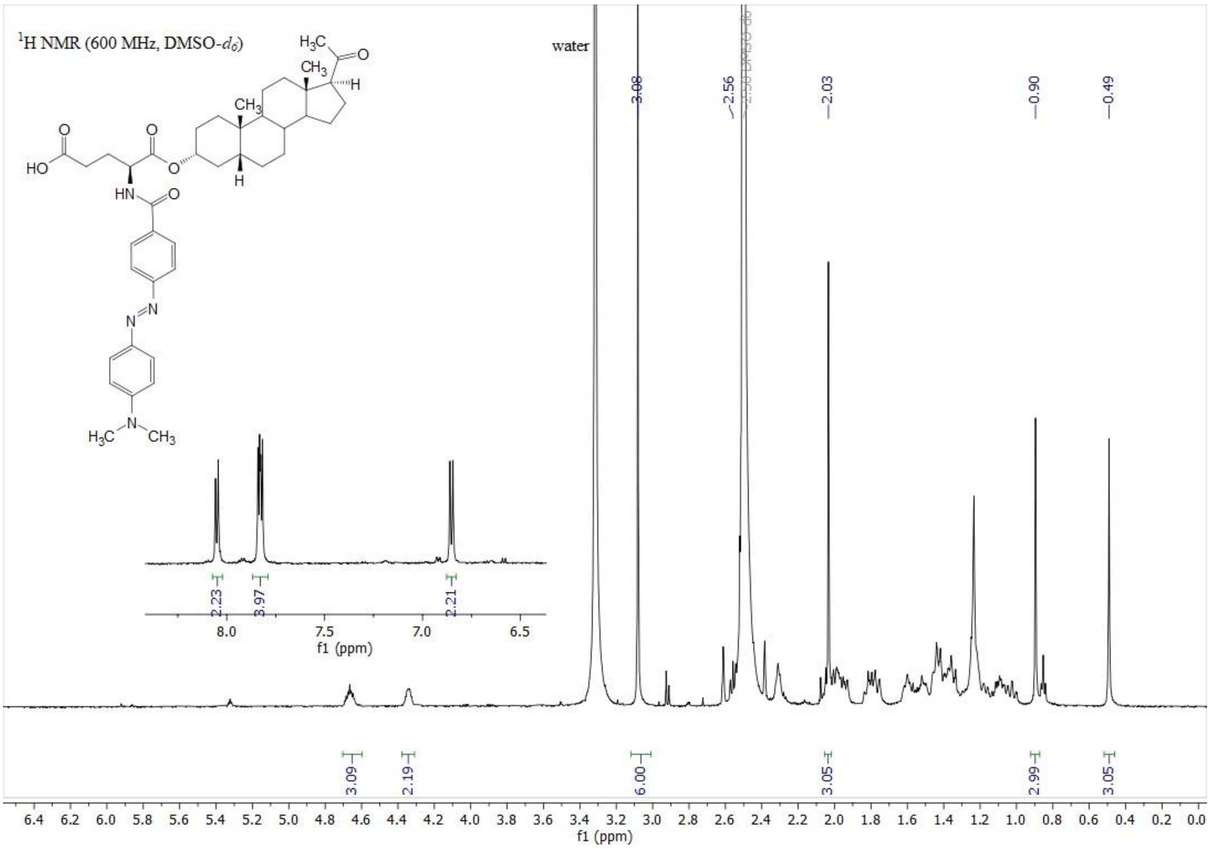
1H NMR (600 MHz, DMSO-d6)
water
3.08
2.56
2.50 DMSO-d6
2.03
0.90
0.49
2.23
3.97
2.21
8.0
7.5
7.0
6.5
f1 (ppm)
3.09
2.19
6.00
3.05
2.99
3.05
6.4 6.2 6.0 5.8 5.6 5.4 5.2 5.0 4.8 4.6 4.4 4.2 4.0 3.8 3.6 3.4 3.2 3.0 2.8 2.6 2.4 2.2 2.0 1.8 1.6 1.4 1.2 1.0 0.8 0.6 0.4 0.2 0.0
f1 (ppm)

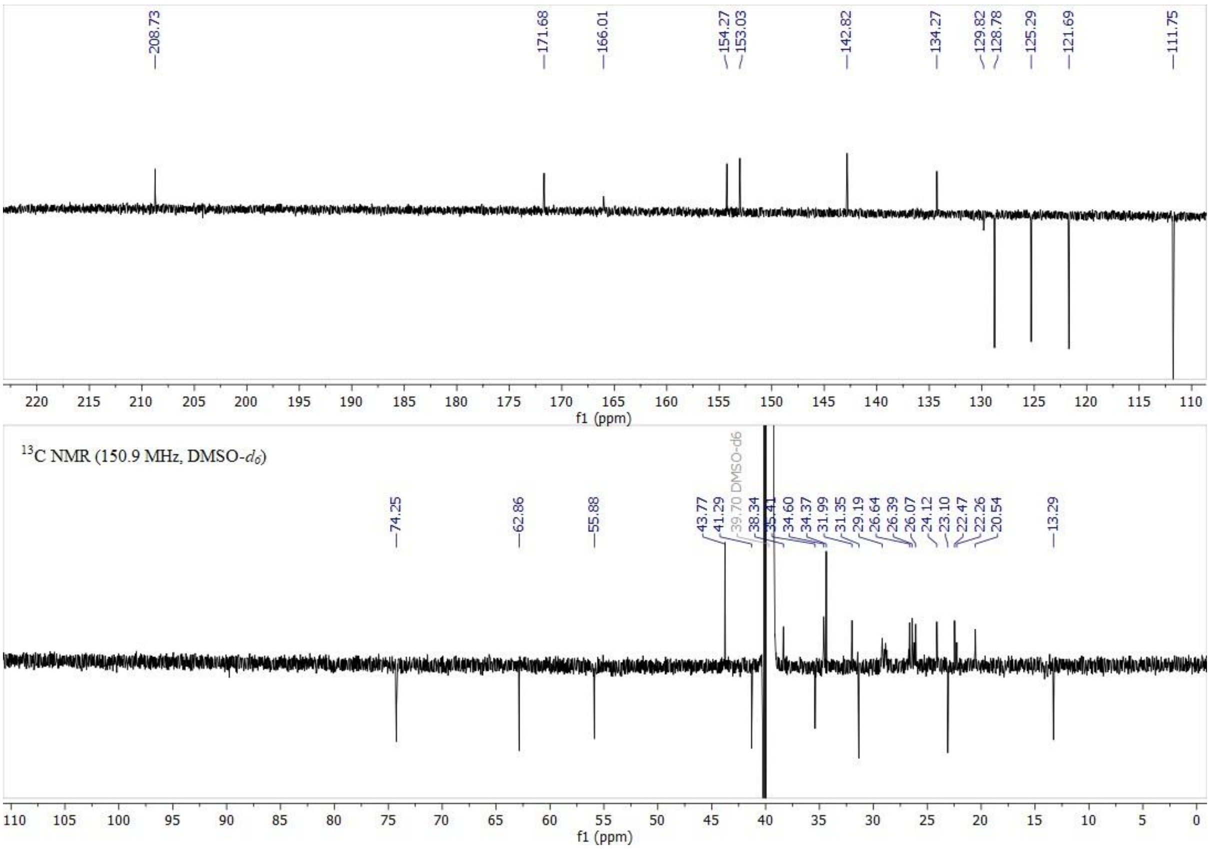
208.73
171.68
166.01
154.27
153.03
142.82
134.27
129.82
128.78
125.29
121.69
111.75
220 215 210 205 200 195 190 185 180 175 170 165 160 155 150 145 140 135 130 125 120 115 110
f1 (ppm)
13C NMR (150.9 MHz, DMSO-d6)
74.25
62.86
55.88
43.77
41.29
39.70 DMSO-d6
38.34
35.41
34.60
34.37
31.99
31.35
29.19
26.64
26.39
26.07
24.12
23.10
22.47
22.26
20.54
13.29
110 105 100 95 90 85 80 75 70 65 60 55 50 45 40 35 30 25 20 15 10 5 0
f1 (ppm)

**14.** C APT NMR (bottom) spectra of compound **1**.

**Figure S** $^{1}$H NMR (top) and $^{13}$

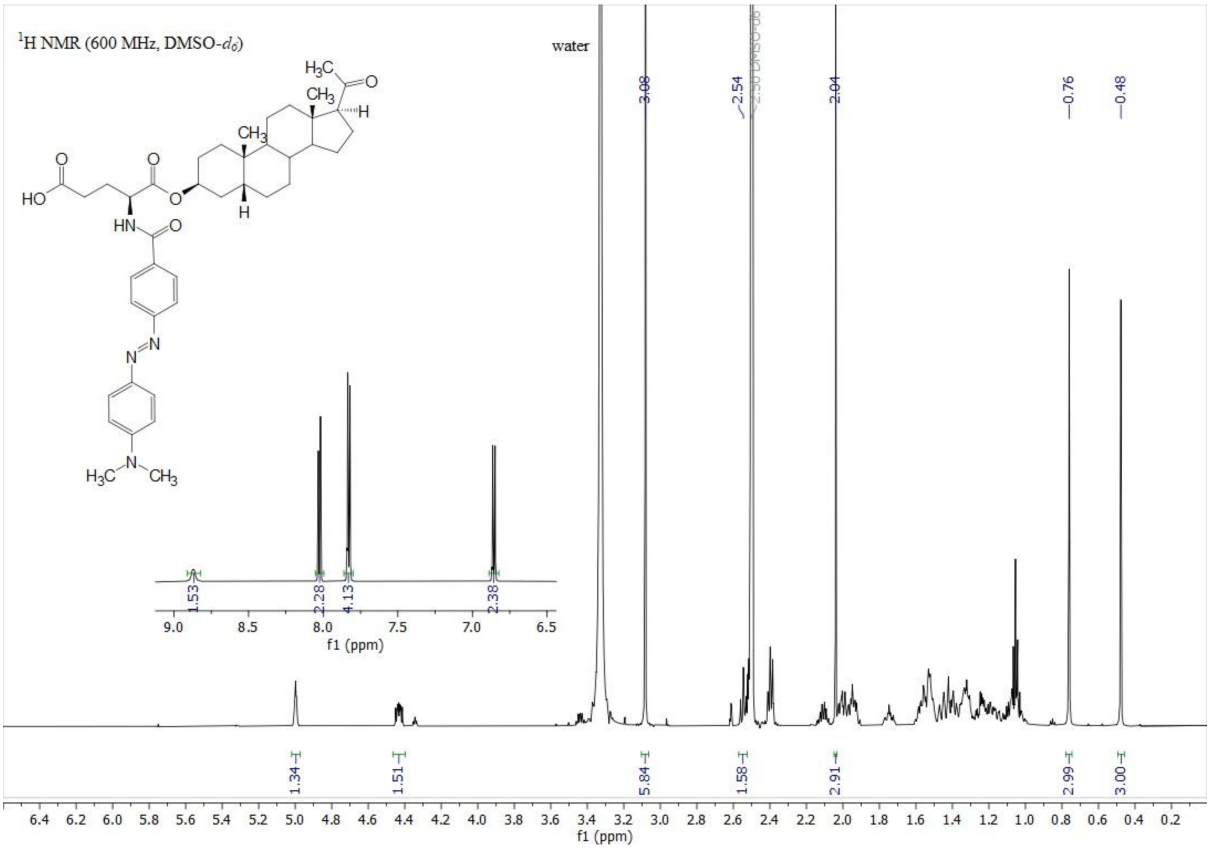

¹H NMR (600 MHz, DMSO-d6)
water
3.08
2.54
2.50 DMSO-d6
2.04
0.76
0.48
1.53
2.28
4.13
2.38
1.34
1.51
5.84
1.58
2.91
2.99
3.00
f1 (ppm)

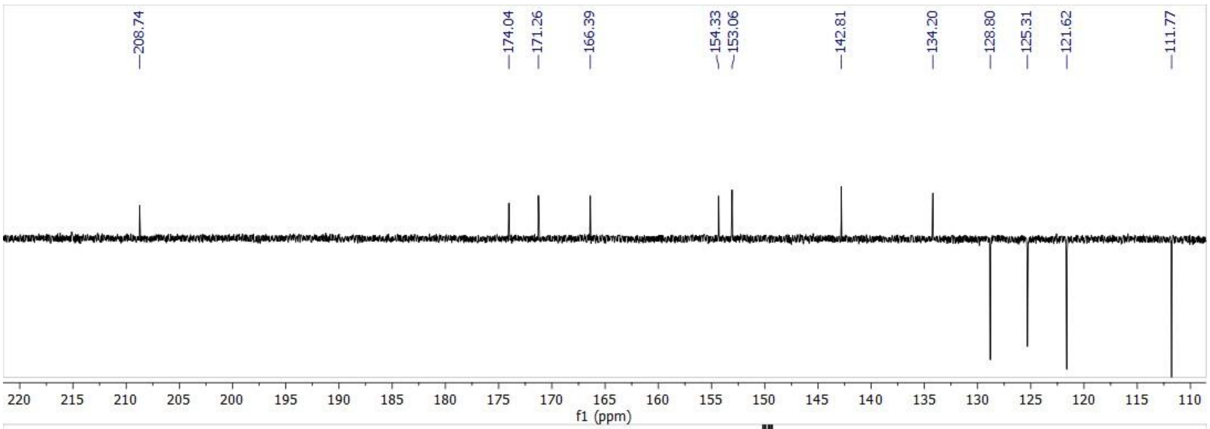

208.74
174.04
171.26
166.39
154.33
153.06
142.81
134.20
128.80
125.31
121.62
111.77
f1 (ppm)

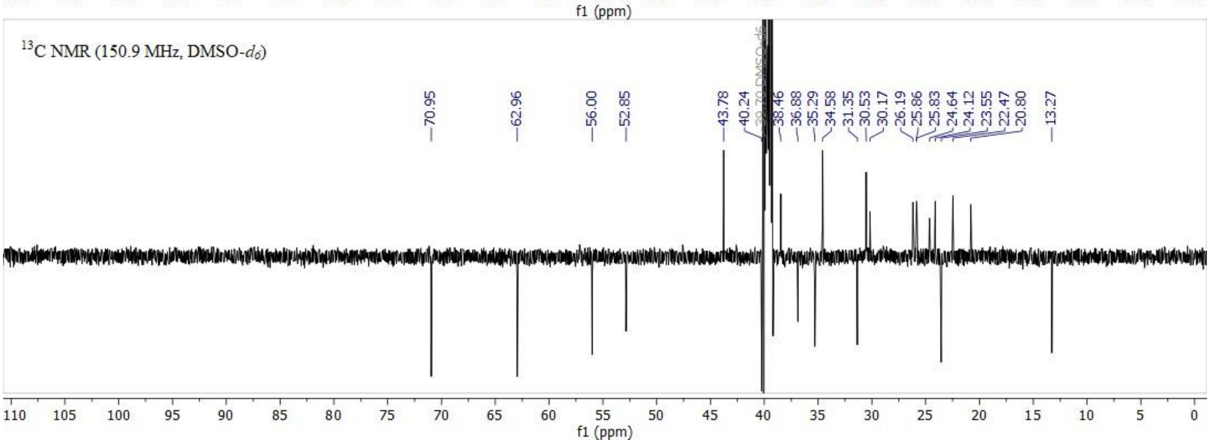

¹³C NMR (150.9 MHz, DMSO-d6)
70.95
62.96
56.00
52.85
43.78
40.24
38.46
36.88
35.29
34.58
31.35
30.53
30.17
26.19
25.86
25.83
24.64
24.12
23.55
22.47
20.80
13.27
f1 (ppm)

**15.** C APT NMR (bottom) spectra of compound **2**.

**Figure S** $^{1}$H NMR (top) and $^{13}$

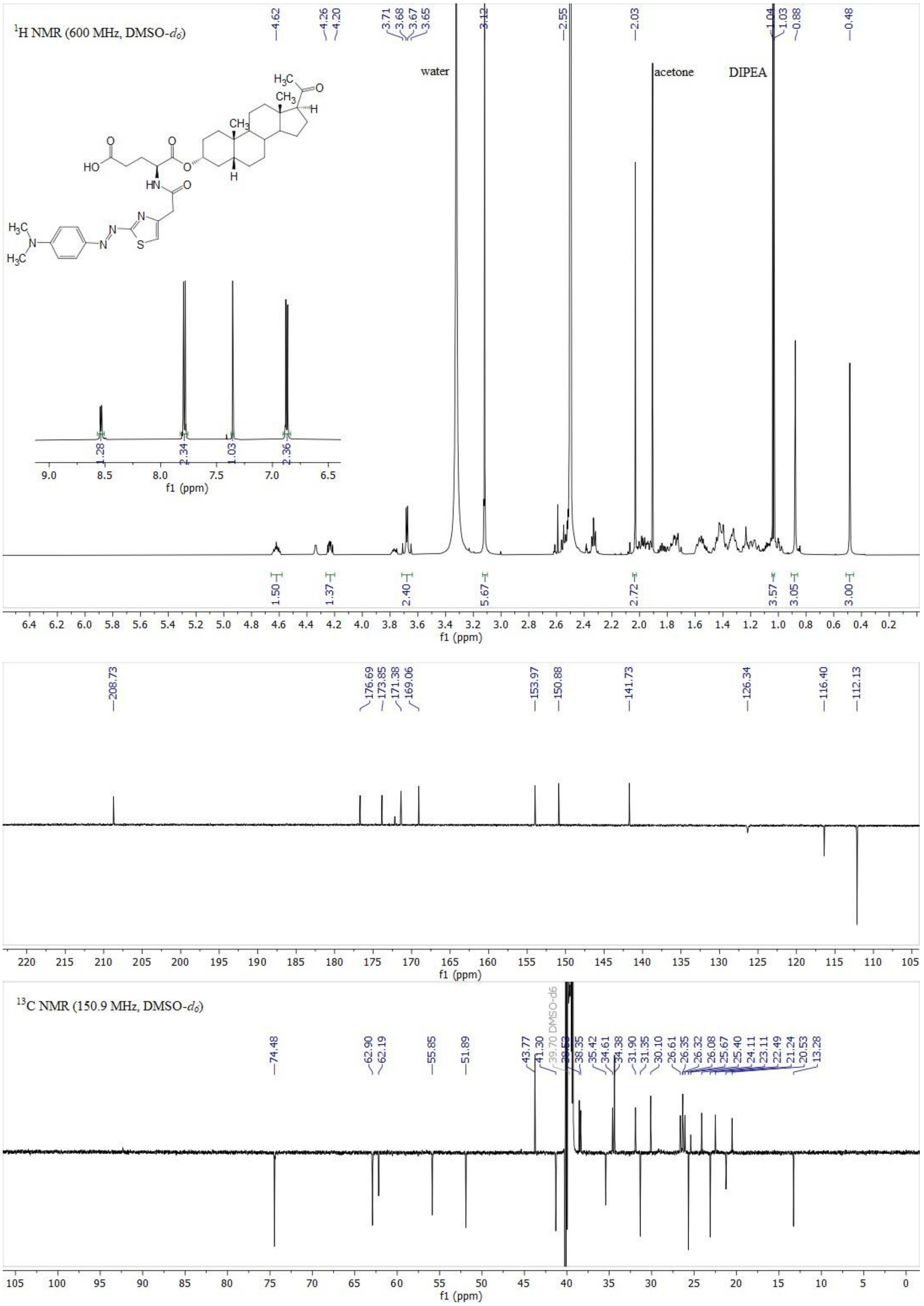

1H NMR (600 MHz, DMSO-d6)
4.62
4.26
4.20
3.71
3.68
3.67
3.65
3.12
2.55
2.03
1.04
1.03
0.88
0.48
water
acetone
DIPEA
1.28
2.34
1.03
2.36
1.50
1.37
2.40
5.67
2.72
3.57
3.05
3.00
f1 (ppm)
208.73
176.69
173.85
171.38
169.06
153.97
150.88
141.73
126.34
116.40
112.13
13C NMR (150.9 MHz, DMSO-d6)
74.48
62.90
62.19
55.85
51.89
43.77
41.30
39.70 DMSO-d6
38.35
35.42
34.61
34.38
31.90
31.35
30.10
26.61
26.35
26.32
26.08
25.67
25.40
24.11
23.11
22.49
21.24
20.53
13.28

**16.** C APT NMR (bottom) spectra of compound **3**.

**Figure S** $^{1}$H NMR (top) and $^{13}$

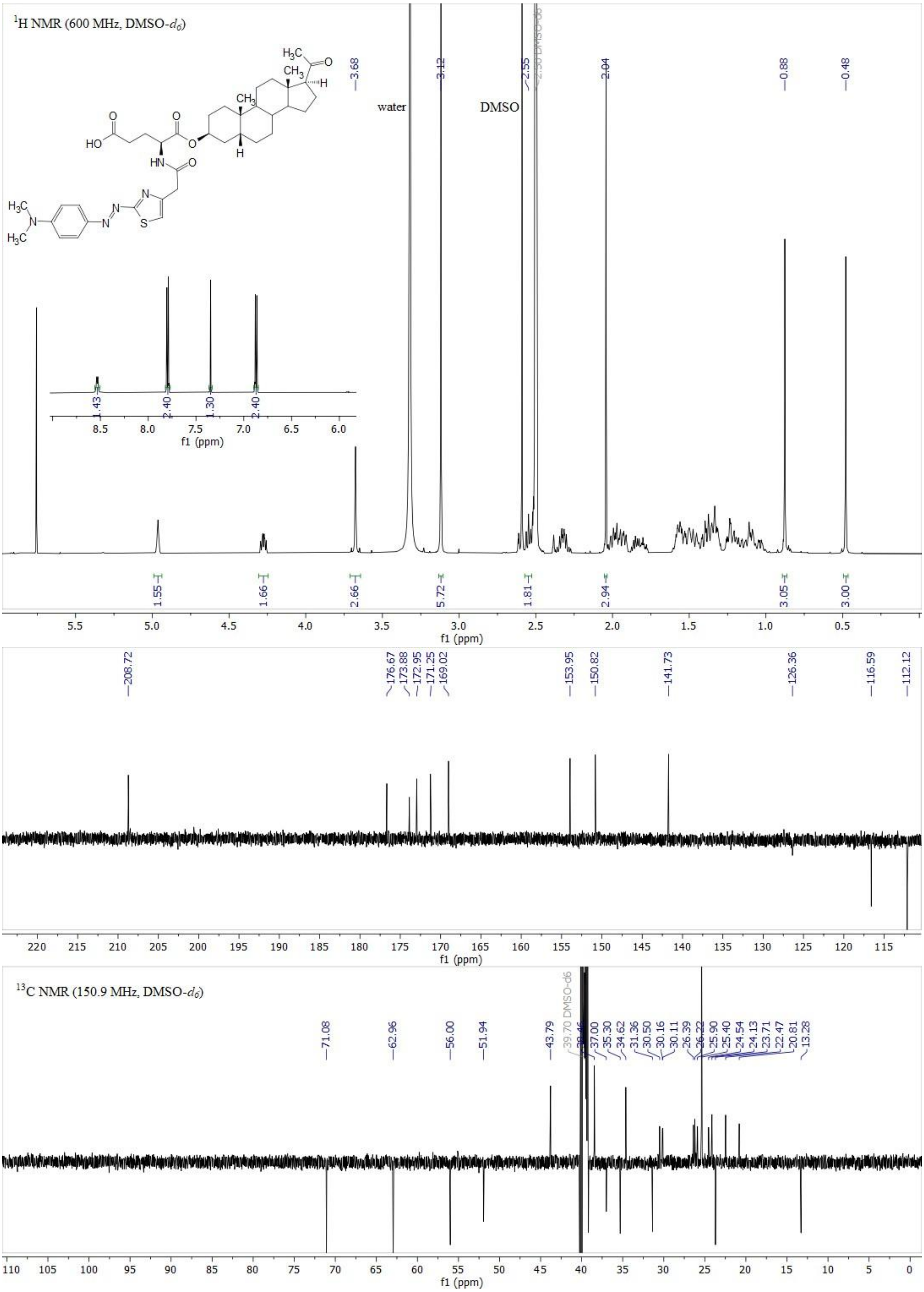

**Figure S17.** $^{1}H$ NMR (top) and $^{13}C$ APT NMR (bottom) spectra of compound **4**.

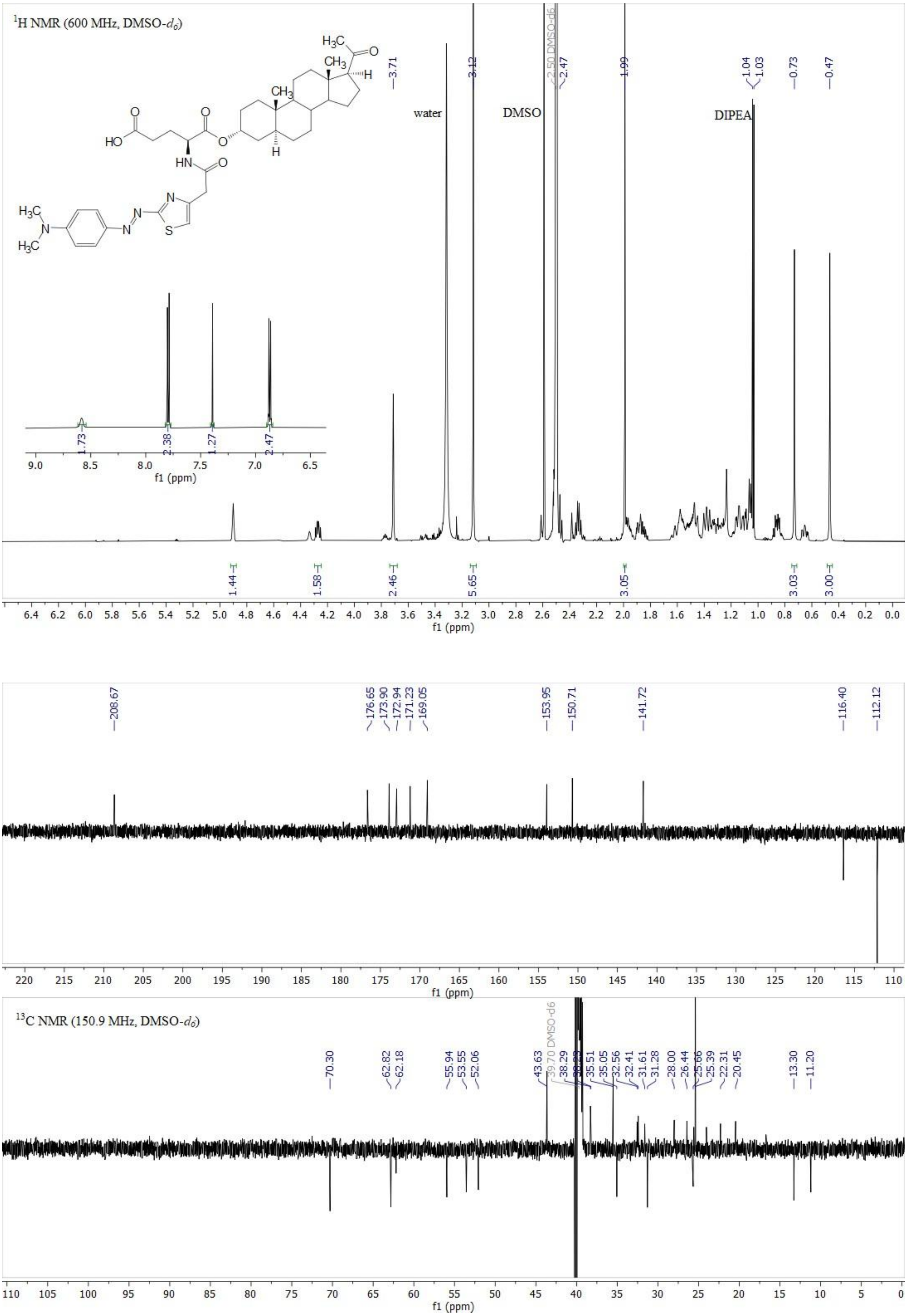

**Figure S18.** $^{1}H$ NMR (top) and $^{13}C$ APT NMR (bottom) spectra of compound **5**.

**Figure S** $^{1}H$ NMR (top) and $^{13}$

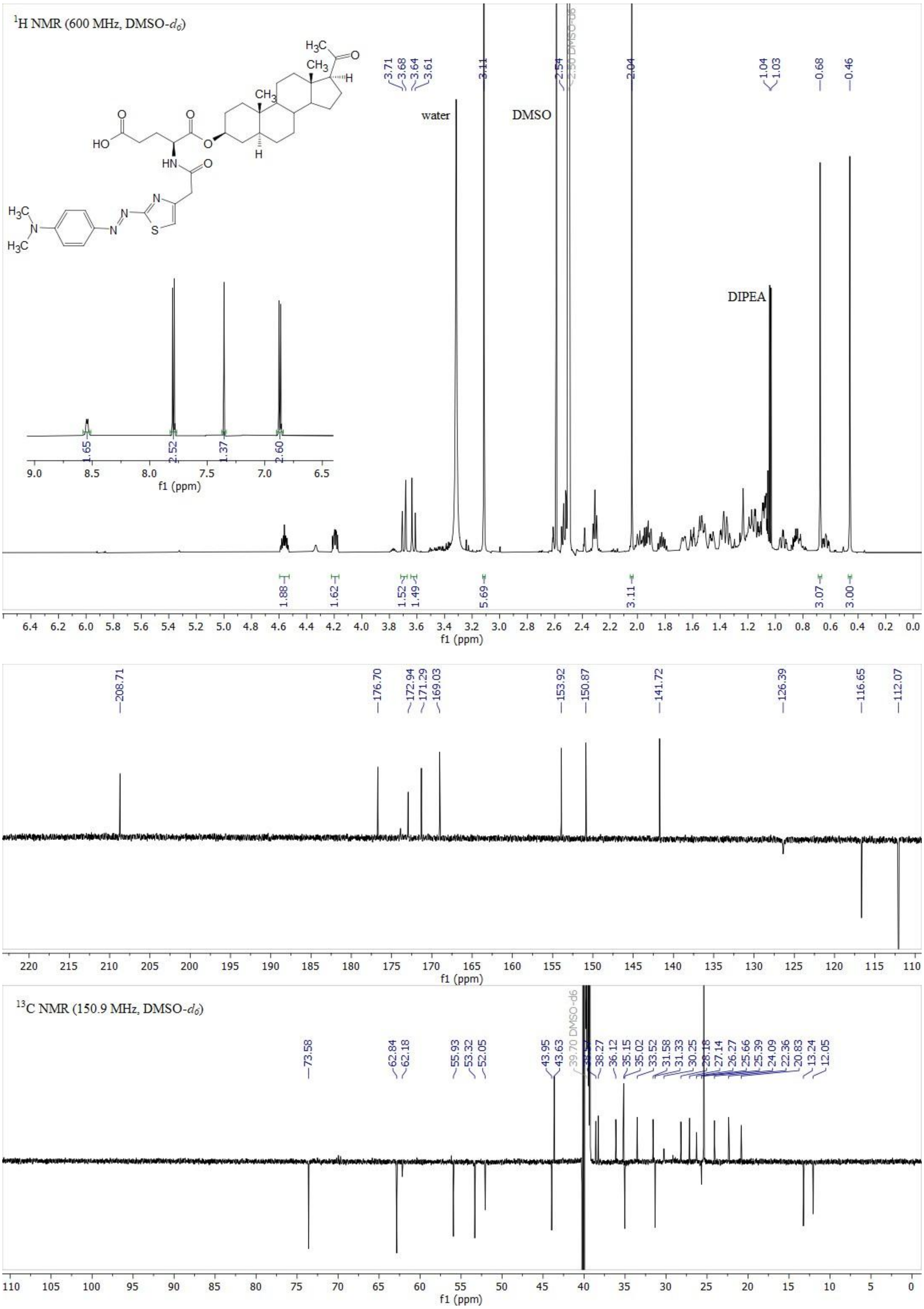

**19.** C APT NMR (bottom) spectra of compound **6**.

**Figure S** [1]H NMR (top) and [13]

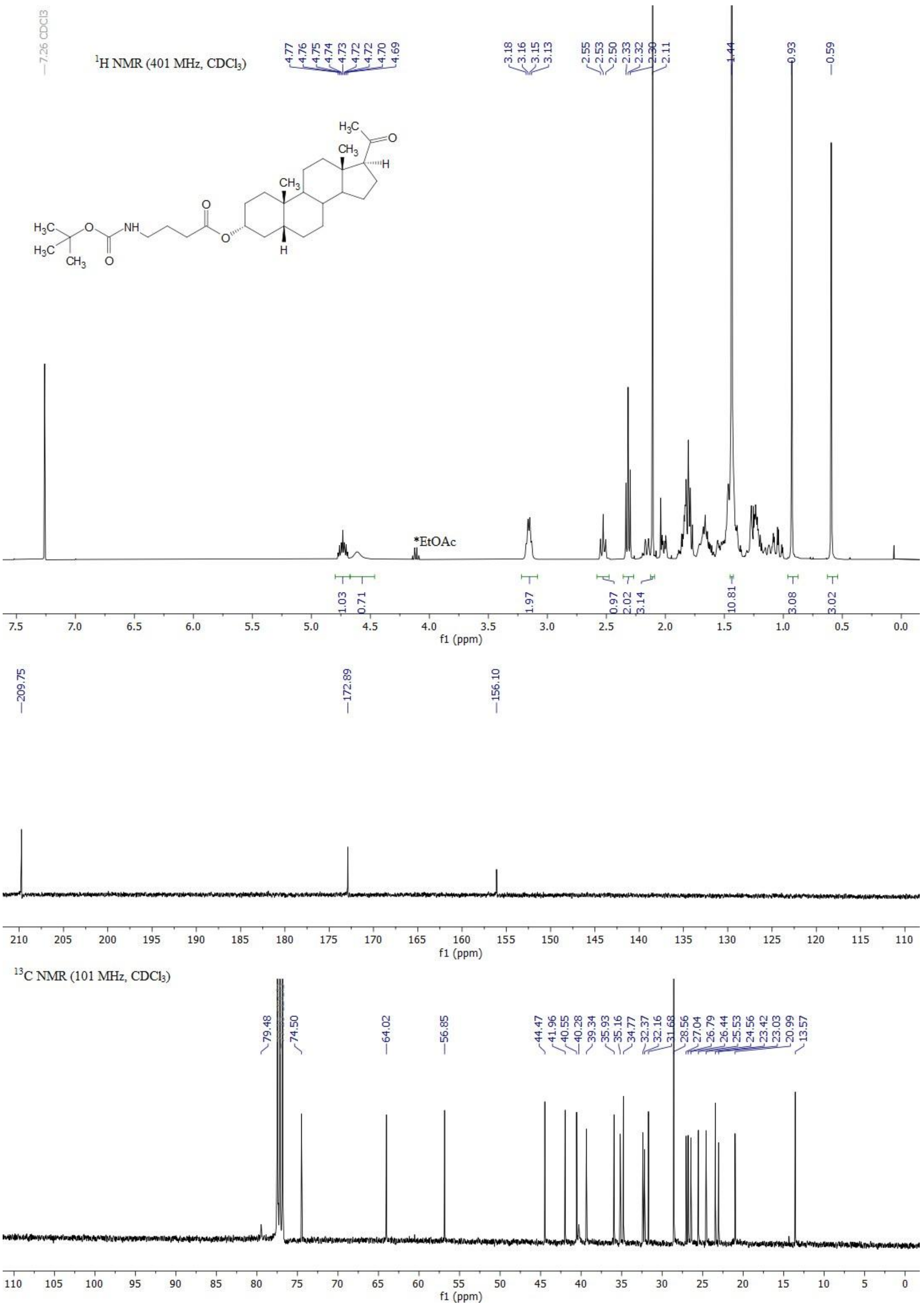

**Figure S20.** $^{1}H$ NMR (top) and $^{13}C$ NMR (bottom) spectra of compound **39**.

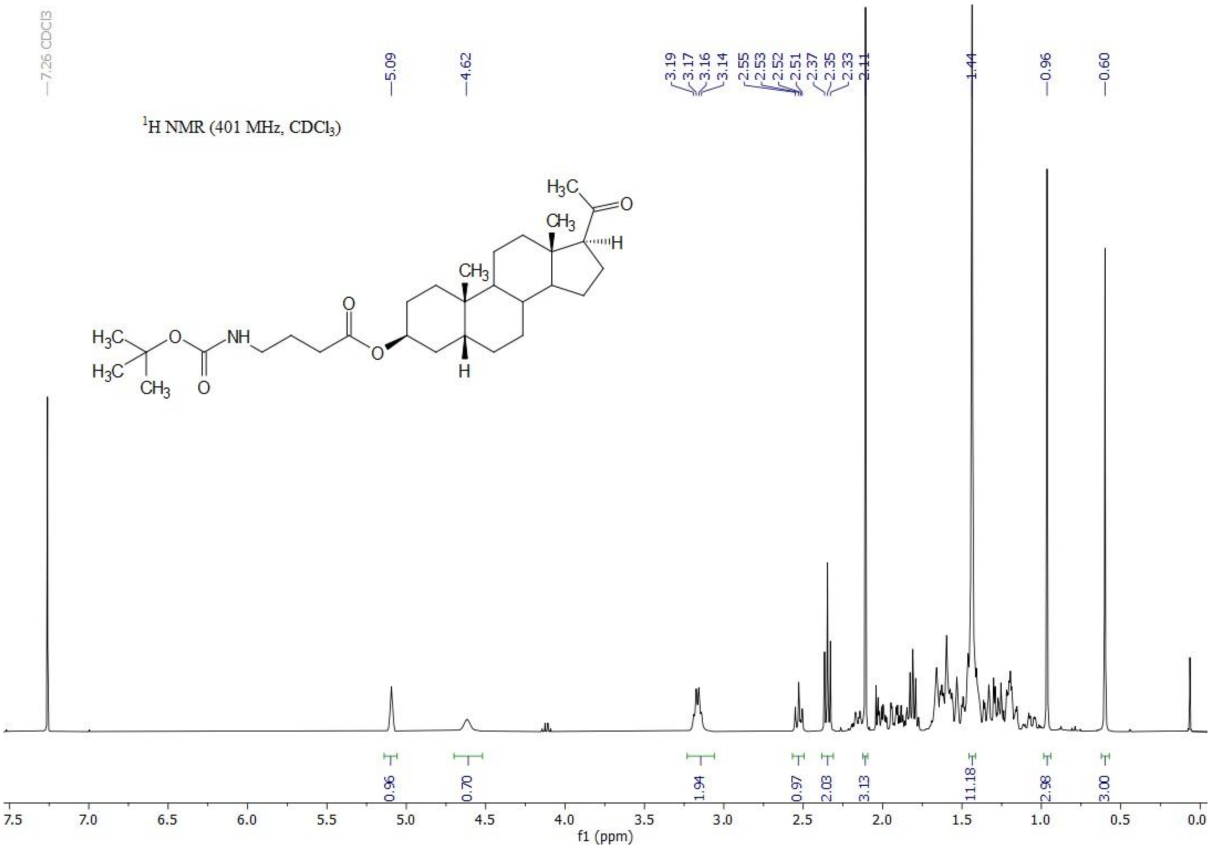

$^1$H NMR (401 MHz, CDCl$_3$)
f1 (ppm)

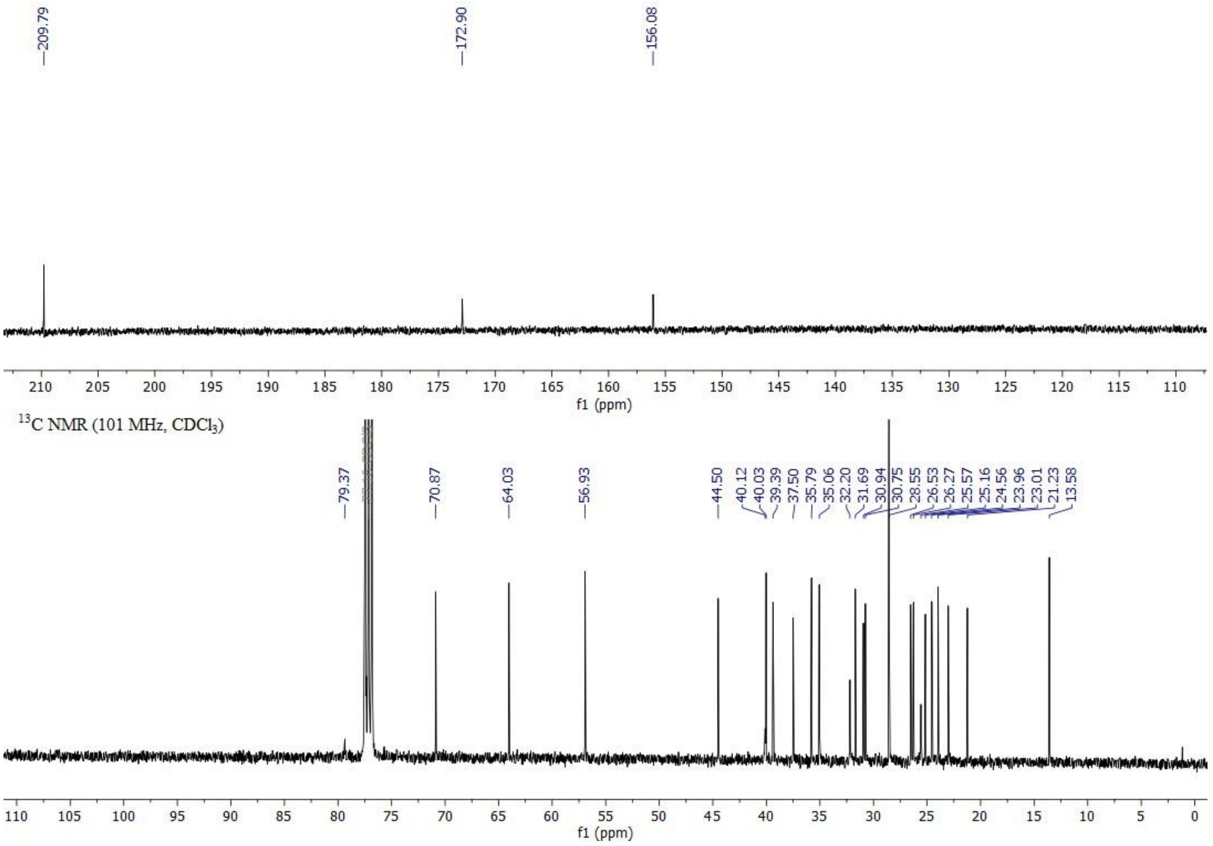

$^{13}$C NMR (101 MHz, CDCl$_3$)
f1 (ppm)

**21.** **40**.

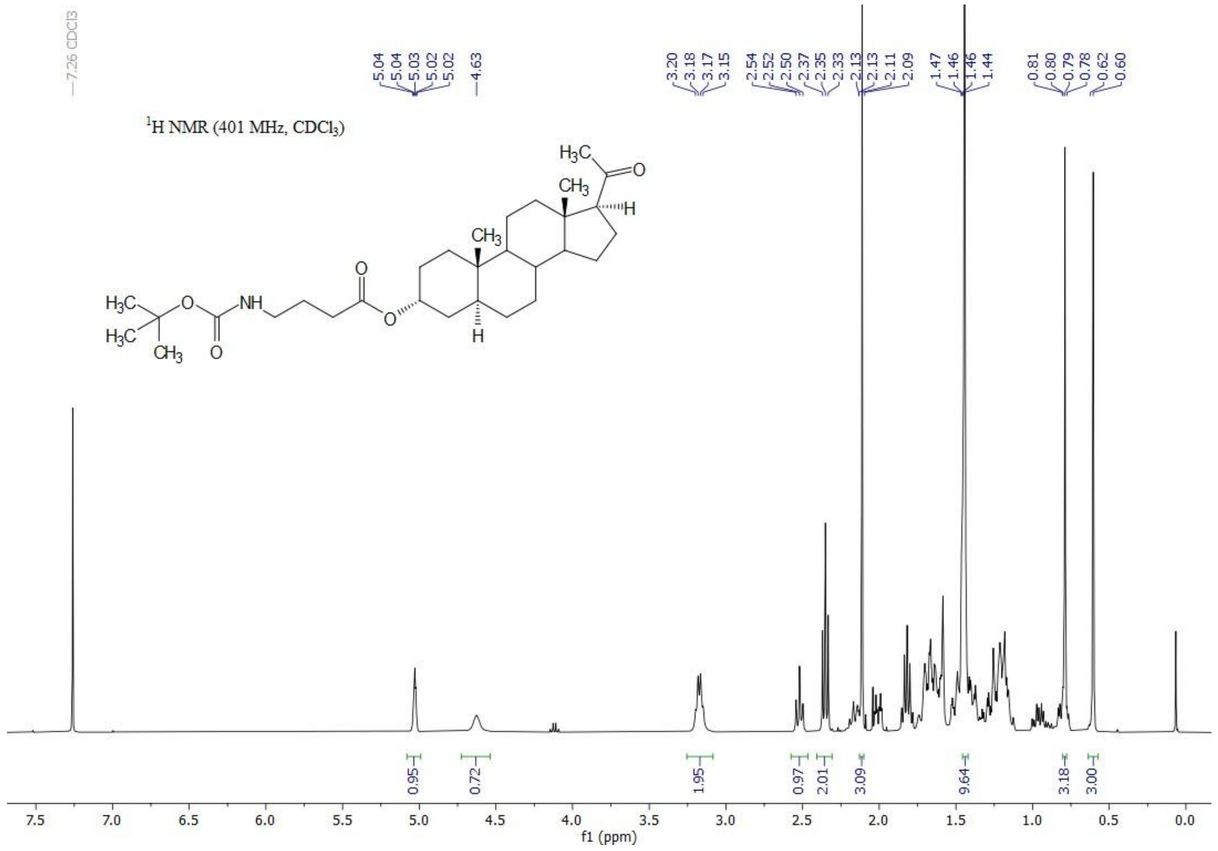

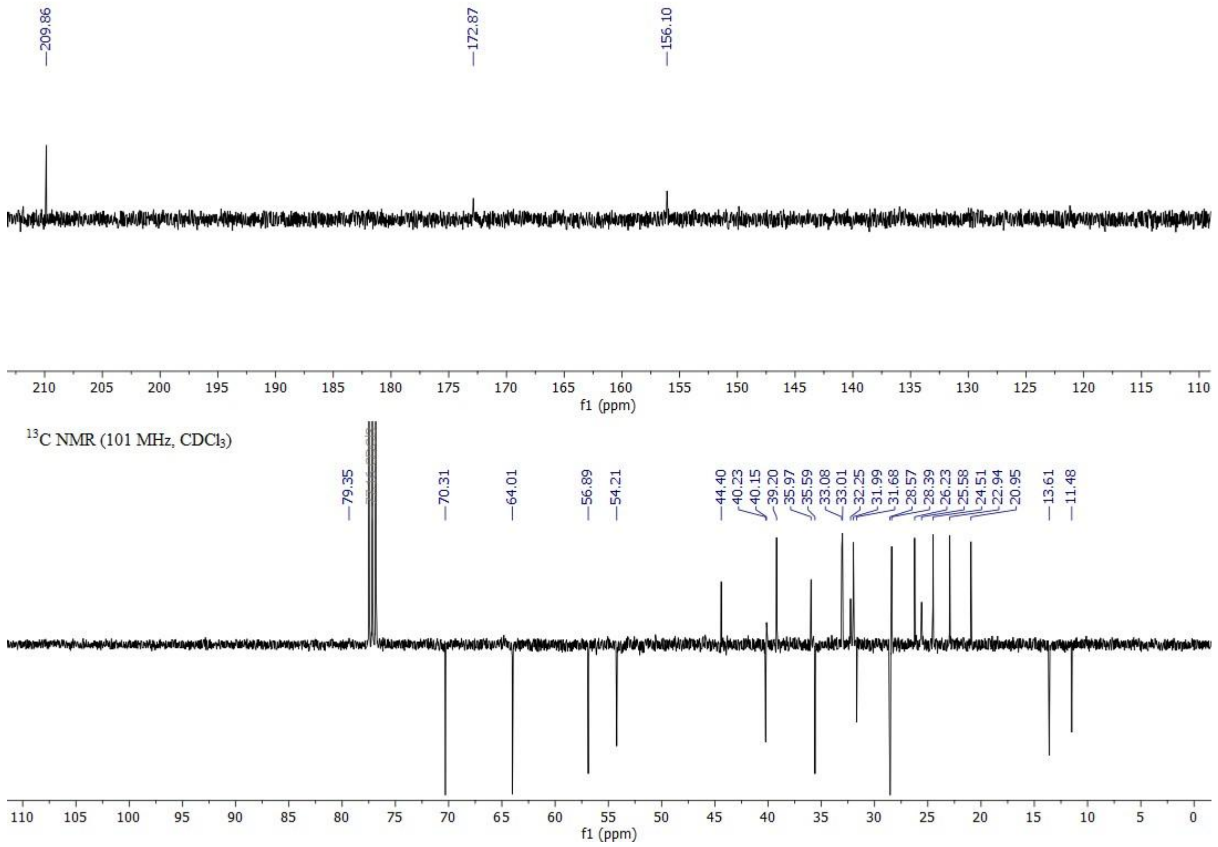

**Figure S22.** $^{1}$H NMR (top) and $^{13}$C NMR (bottom) spectra of compound **41**.

**Figure S** $^{1}H$ NMR (top) and $^{13}C$ NMR (bottom) spectra of compound

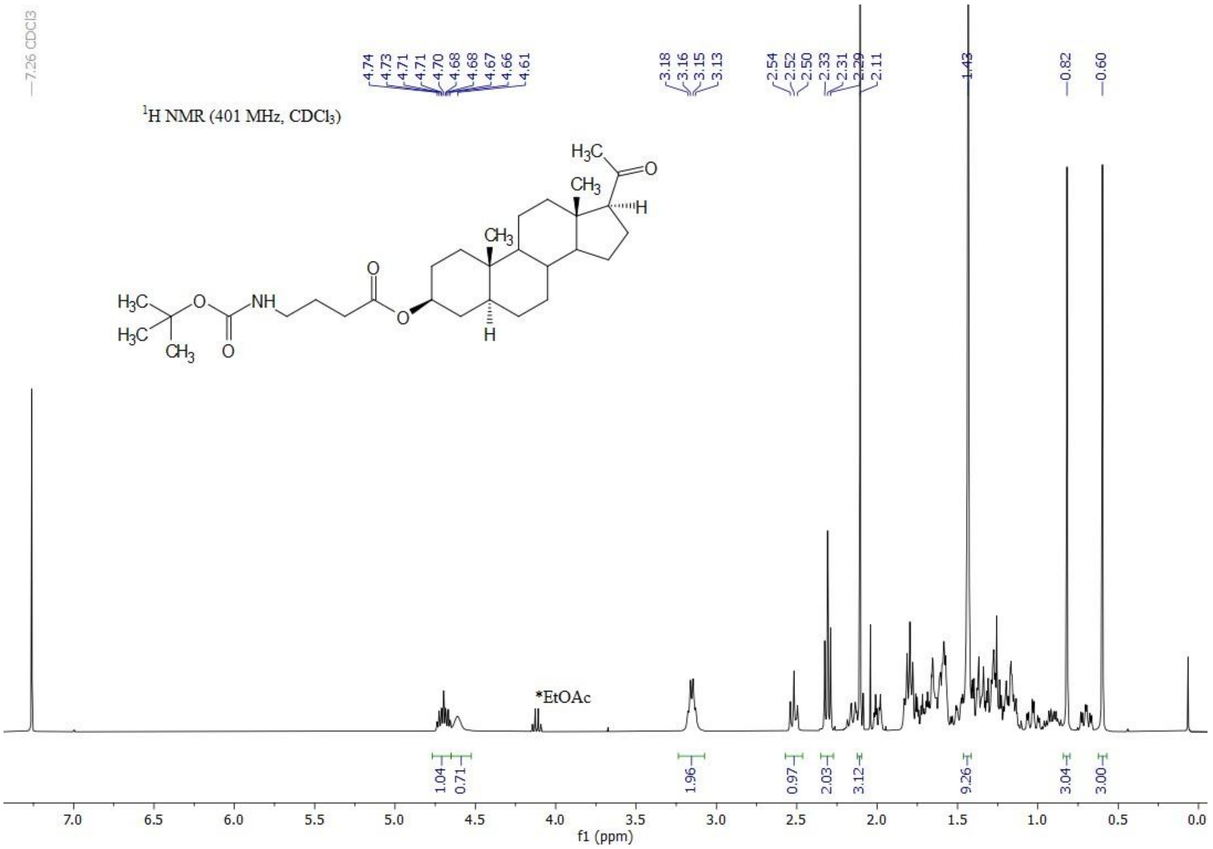

7.26 CDCl3
4.74 4.73 4.71 4.71 4.70 4.68 4.68 4.67 4.66 4.61
3.18 3.16 3.15 3.13
2.54 2.52 2.50 2.33 2.31 2.29 2.11
1.43
0.82
0.60
1H NMR (401 MHz, CDCl3)
*EtOAc
1.04 0.71 1.96 0.97 2.03 3.12 9.26 3.04 3.00
f1 (ppm)

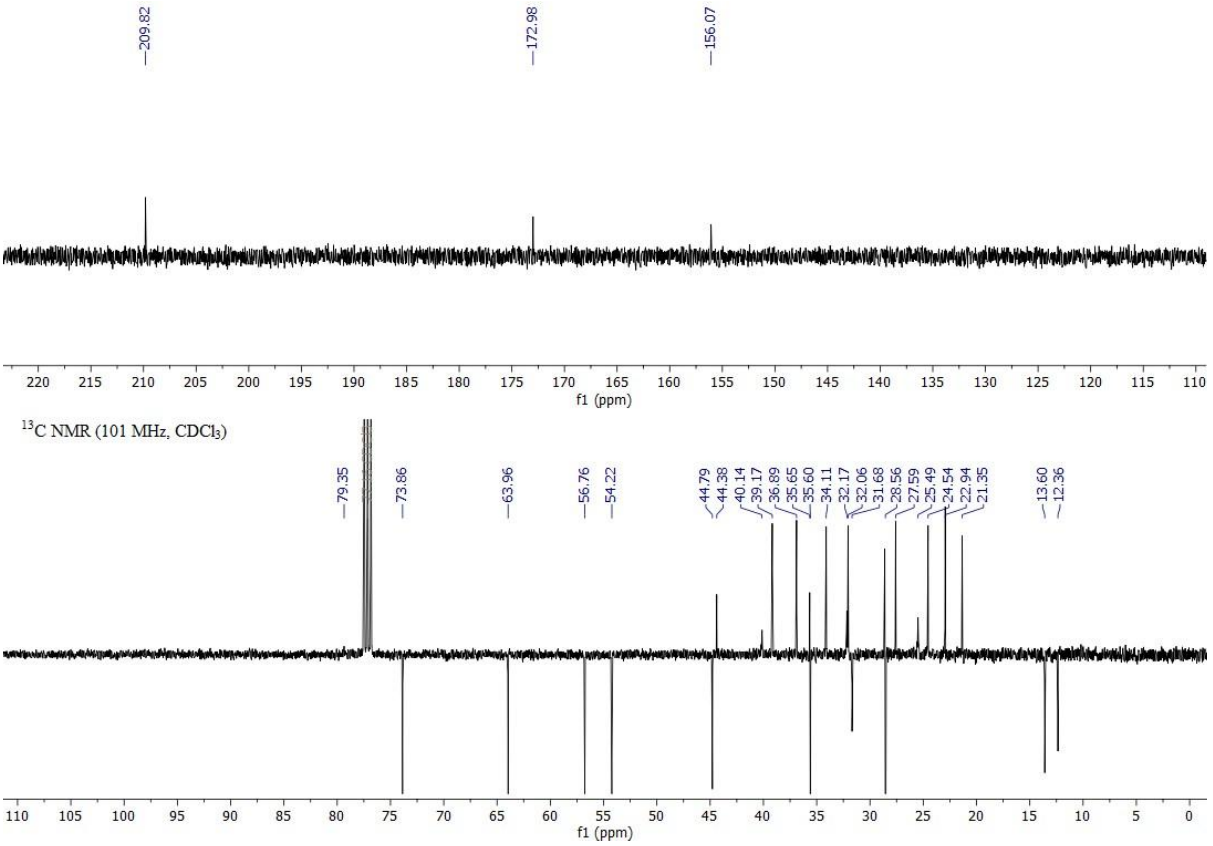

209.82
172.98
156.07
f1 (ppm)
13C NMR (101 MHz, CDCl3)
79.35
73.86
63.96
56.76
54.22
44.79 44.38 40.14 39.17 36.89 35.65 35.60 34.11 32.17 32.06 31.68 28.56 27.59 25.49 24.54 22.94 21.35
13.60 12.36
f1 (ppm)

**23.** **42**.

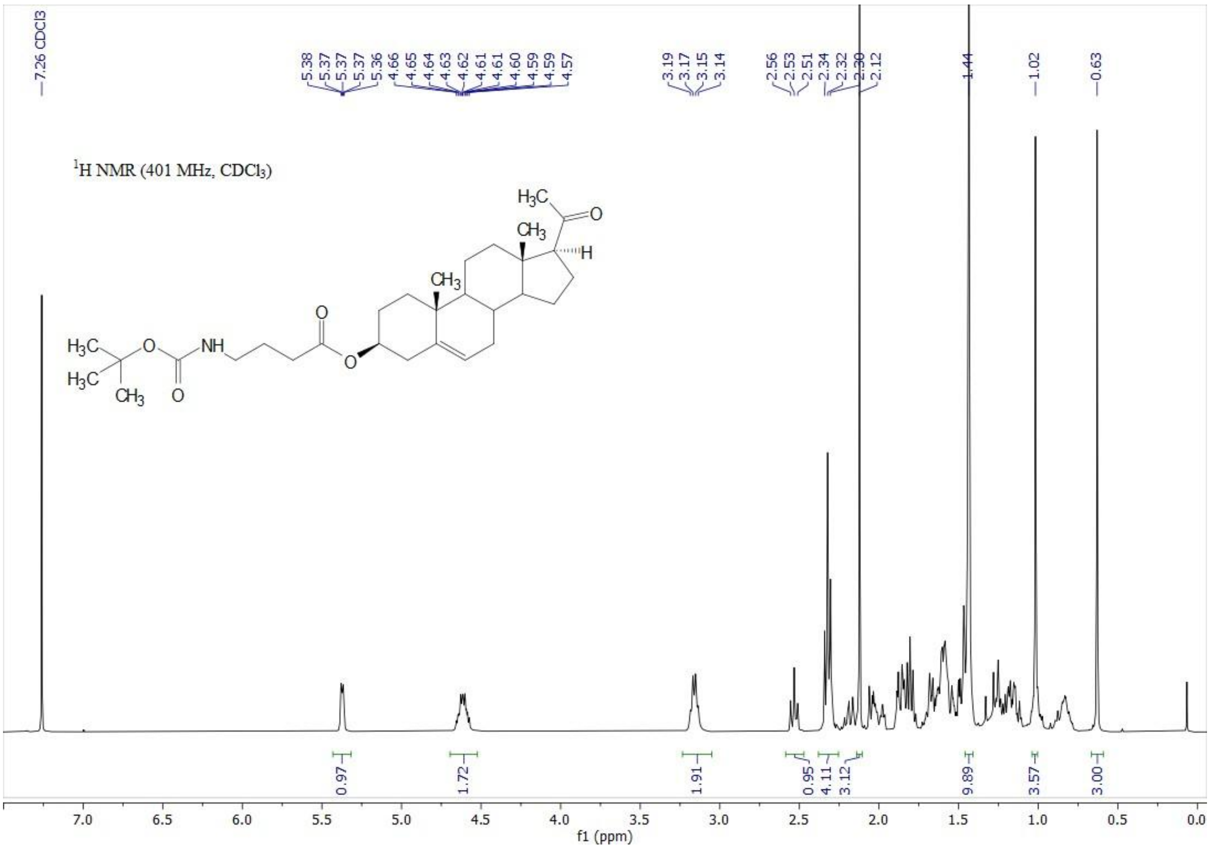
7.26 CDCl3
5.38
5.37
5.37
5.37
5.36
4.66
4.65
4.64
4.63
4.62
4.61
4.61
4.60
4.59
4.59
4.57
3.19
3.17
3.15
3.14
2.56
2.53
2.51
2.34
2.32
2.30
2.12
1.44
1.02
0.63
1H NMR (401 MHz, CDCl3)
0.97
1.72
1.91
0.95
4.11
3.12
9.89
3.57
3.00
f1 (ppm)

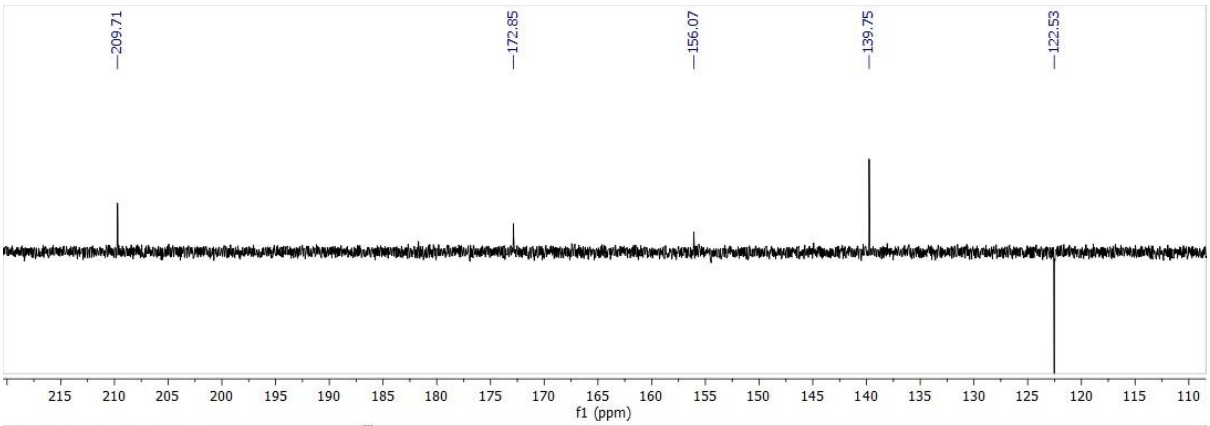
209.71
172.85
156.07
139.75
122.53
f1 (ppm)

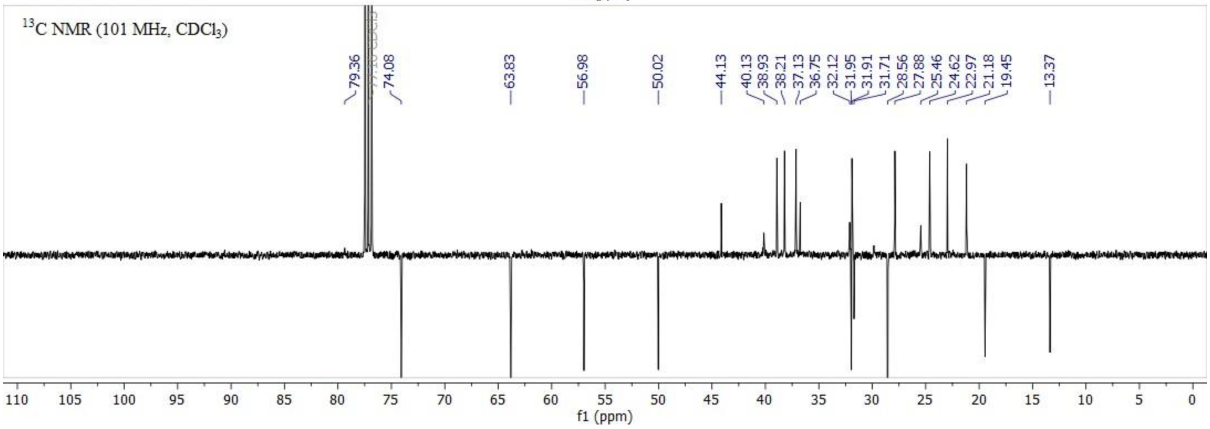
13C NMR (101 MHz, CDCl3)
79.36
74.08
63.83
56.98
50.02
44.13
40.13
38.93
38.21
37.13
36.75
32.12
31.95
31.91
31.71
28.56
27.88
25.46
24.62
22.97
21.18
19.45
13.37
f1 (ppm)

**24.** **48**.

**Figure S** $^{1}H$ NMR (top) and $^{13}C$ NMR (bottom) spectra of compound

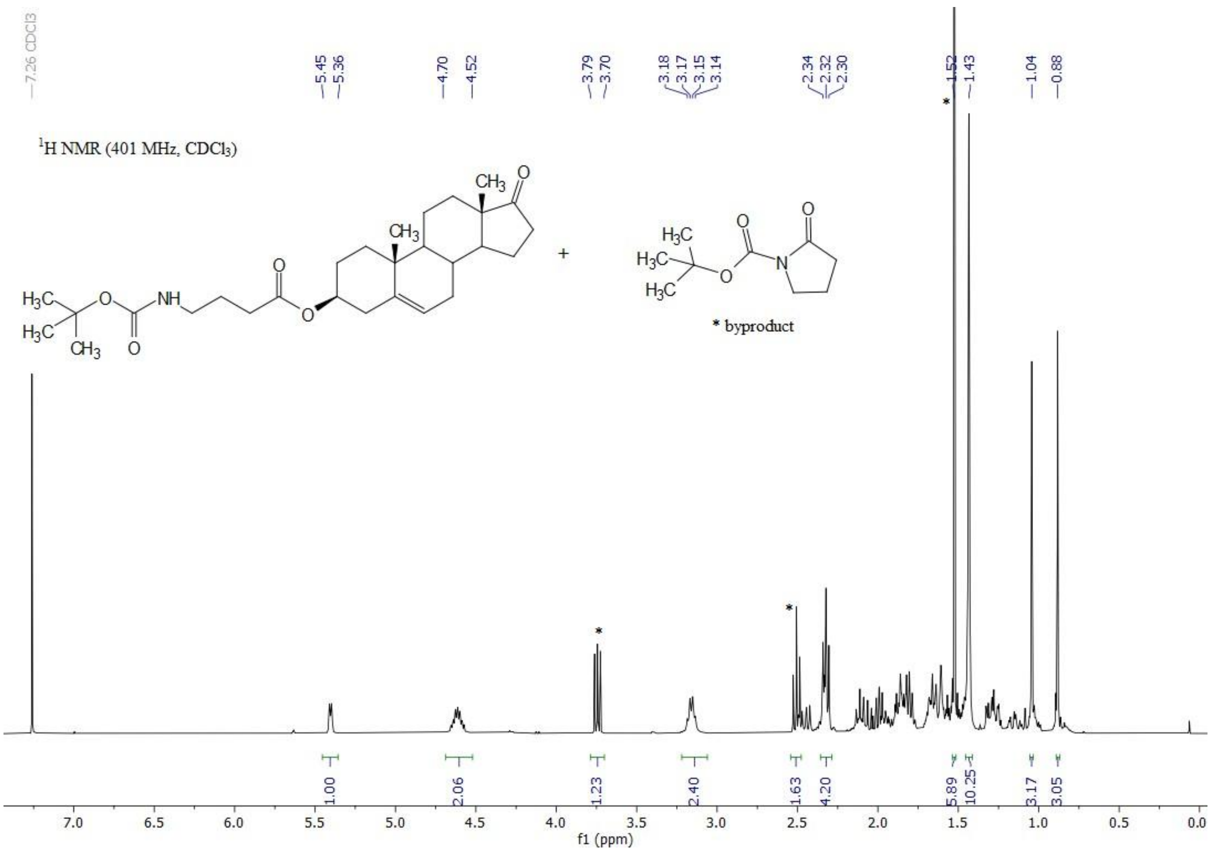
7.26 CDCl3
5.45
5.36
4.70
4.52
3.79
3.70
3.18
3.17
3.15
3.14
2.34
2.32
2.30
1.52
1.43
1.04
0.88
$^{1}$H NMR (401 MHz, CDCl$_3$)
* byproduct
1.00
2.06
1.23
2.40
1.63
4.20
5.89
10.25
3.17
3.05
f1 (ppm)

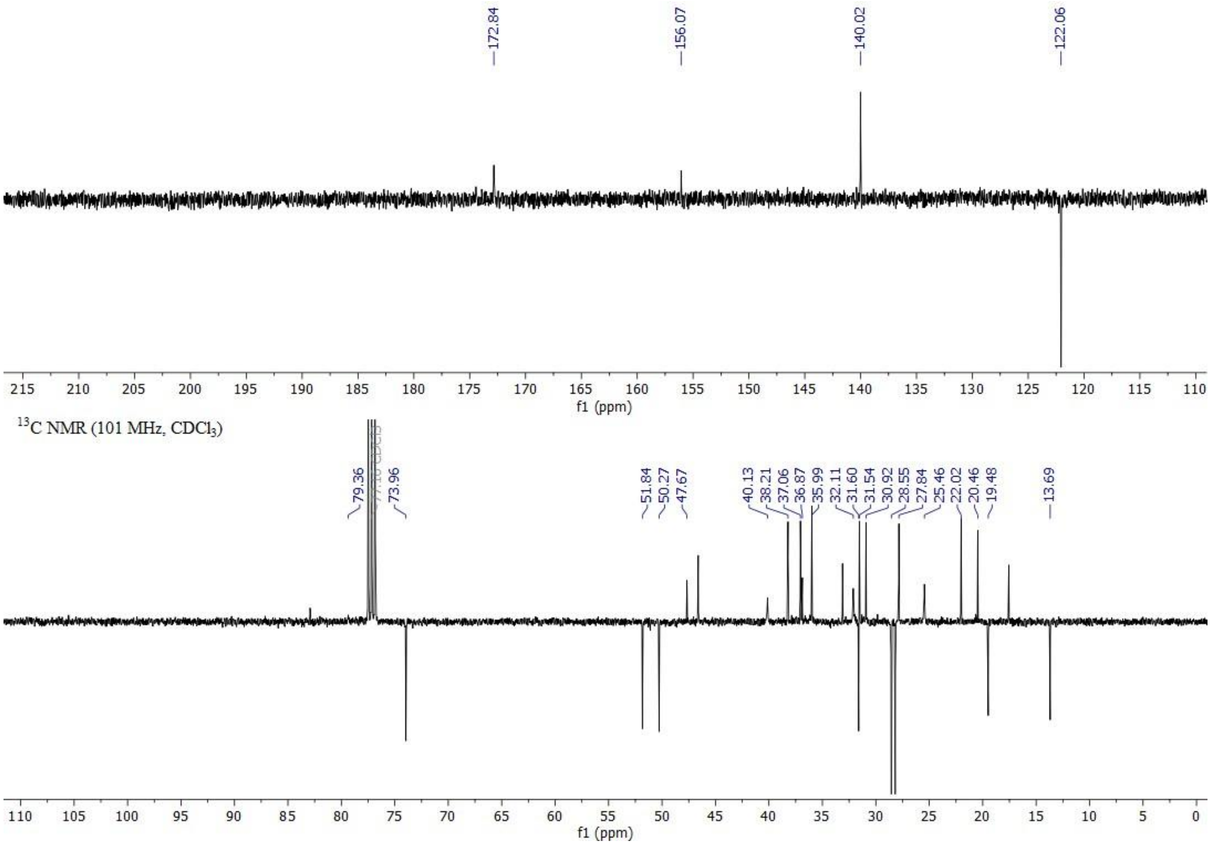
172.84
156.07
140.02
122.06
f1 (ppm)
$^{13}$C NMR (101 MHz, CDCl$_3$)
79.36
73.96
51.84
50.27
47.67
40.13
38.21
37.06
36.87
35.99
32.11
31.60
31.54
30.92
28.55
27.84
25.46
22.02
20.46
19.48
13.69
f1 (ppm)

**Figure S25.** [1] **51**. In the $^{13}C$ NMR spectrum, the signals of the steroid are selected.

**Figure S** $^{1}H$ NMR (top) and $^{13}C$ NMR (bottom) spectra of compound

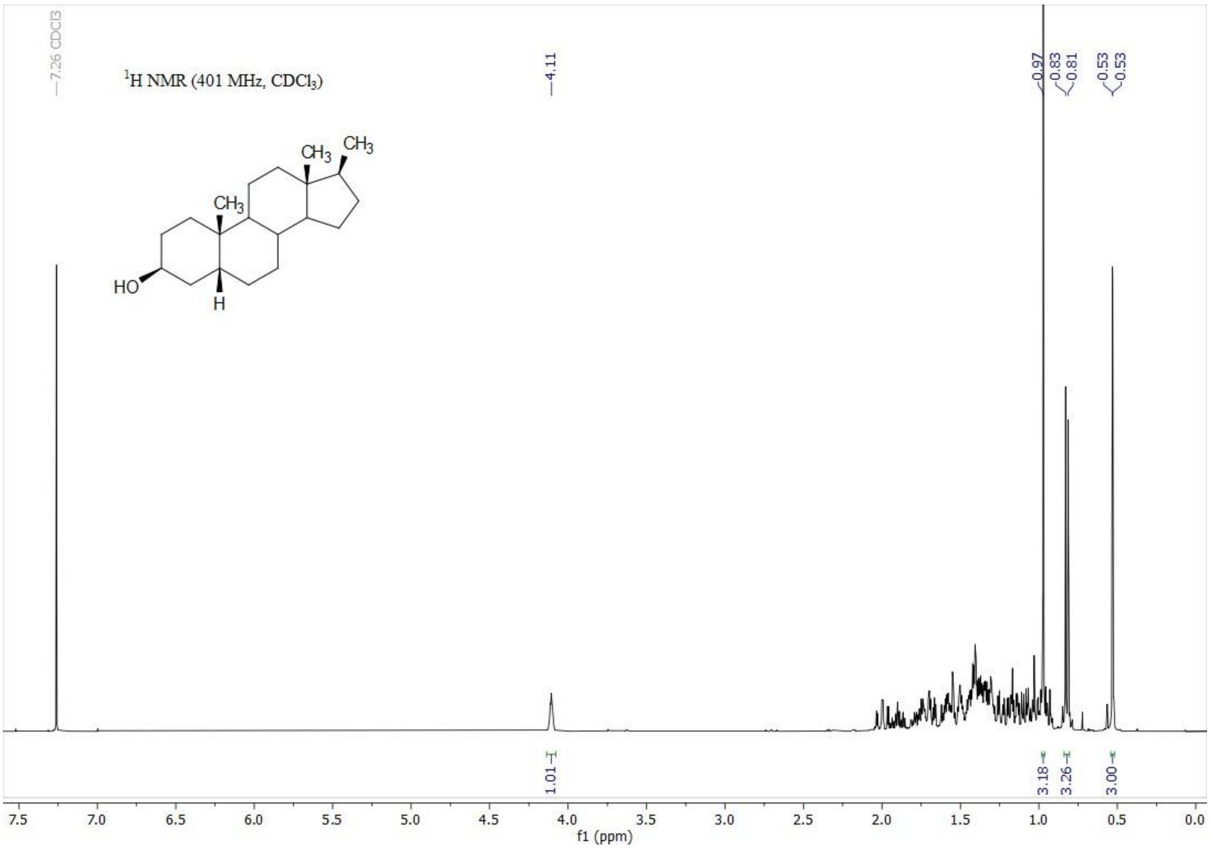

7.26 CDCl3
1H NMR (401 MHz, CDCl3)
4.11
0.97
0.83
0.81
0.53
0.53
HO
H
CH3
CH3
CH3
1.01
3.18
3.26
3.00
7.5 7.0 6.5 6.0 5.5 5.0 4.5 4.0 3.5 3.0 2.5 2.0 1.5 1.0 0.5 0.0
f1 (ppm)

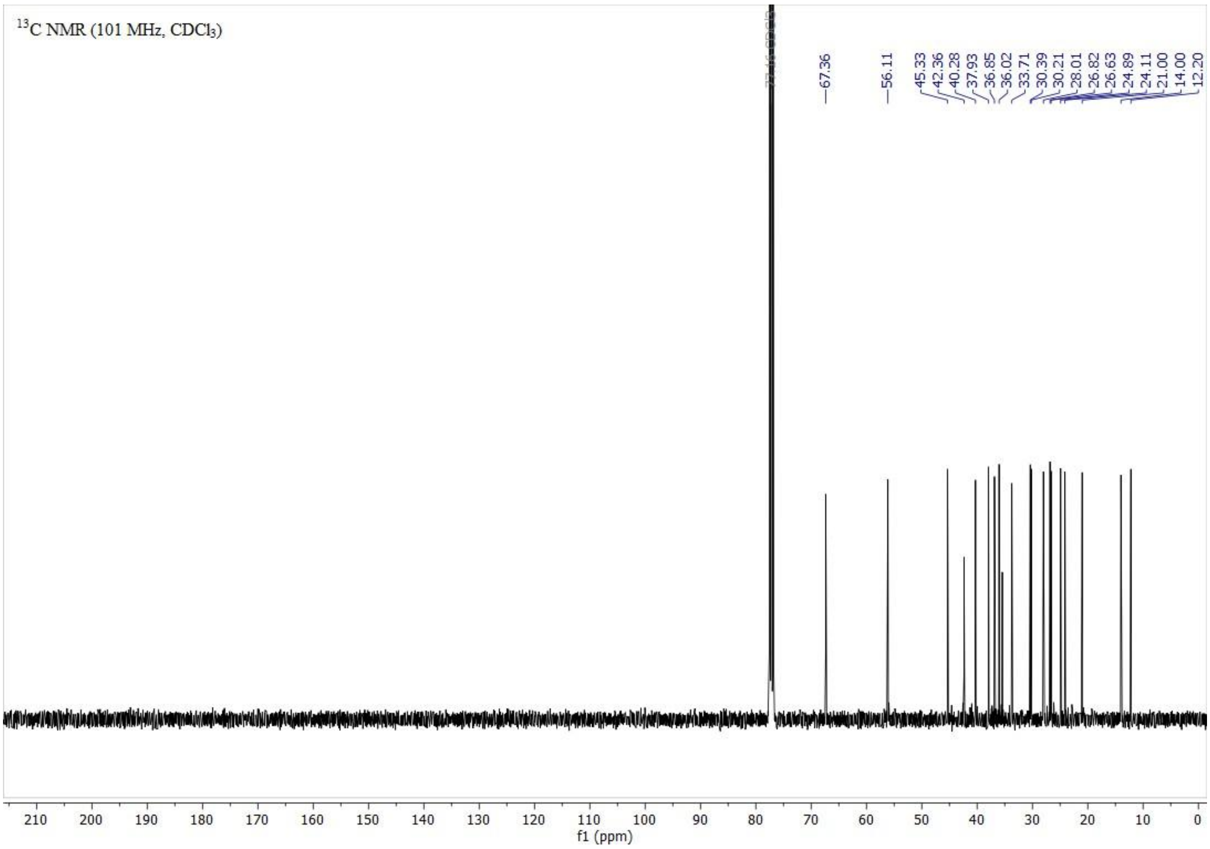

13C NMR (101 MHz, CDCl3)
67.36
56.11
45.33
42.36
40.28
37.93
36.85
36.02
33.71
30.39
30.21
28.01
26.82
26.63
24.89
24.11
21.00
14.00
12.20
210 200 190 180 170 160 150 140 130 120 110 100 90 80 70 60 50 40 30 20 10 0
f1 (ppm)

**26.** **54**.

**Figure S** $^{1}H$ NMR (top) and $^{13}C$ NMR (bottom) spectra of compound

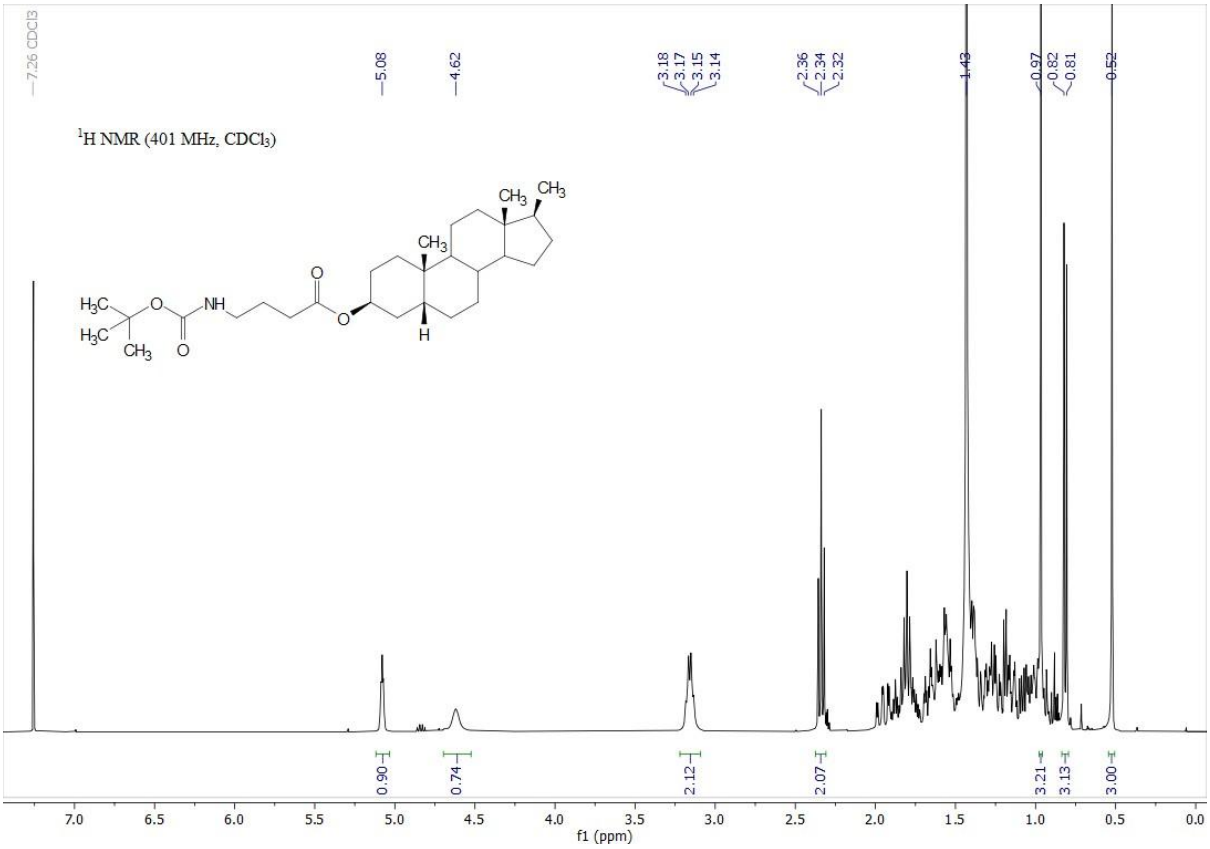
7.26 CDCl3
5.08
4.62
3.18
3.17
3.15
3.14
2.36
2.34
2.32
1.43
0.97
0.82
0.81
0.52
1H NMR (401 MHz, CDCl3)
0.90
0.74
2.12
2.07
3.21
3.13
3.00
f1 (ppm)

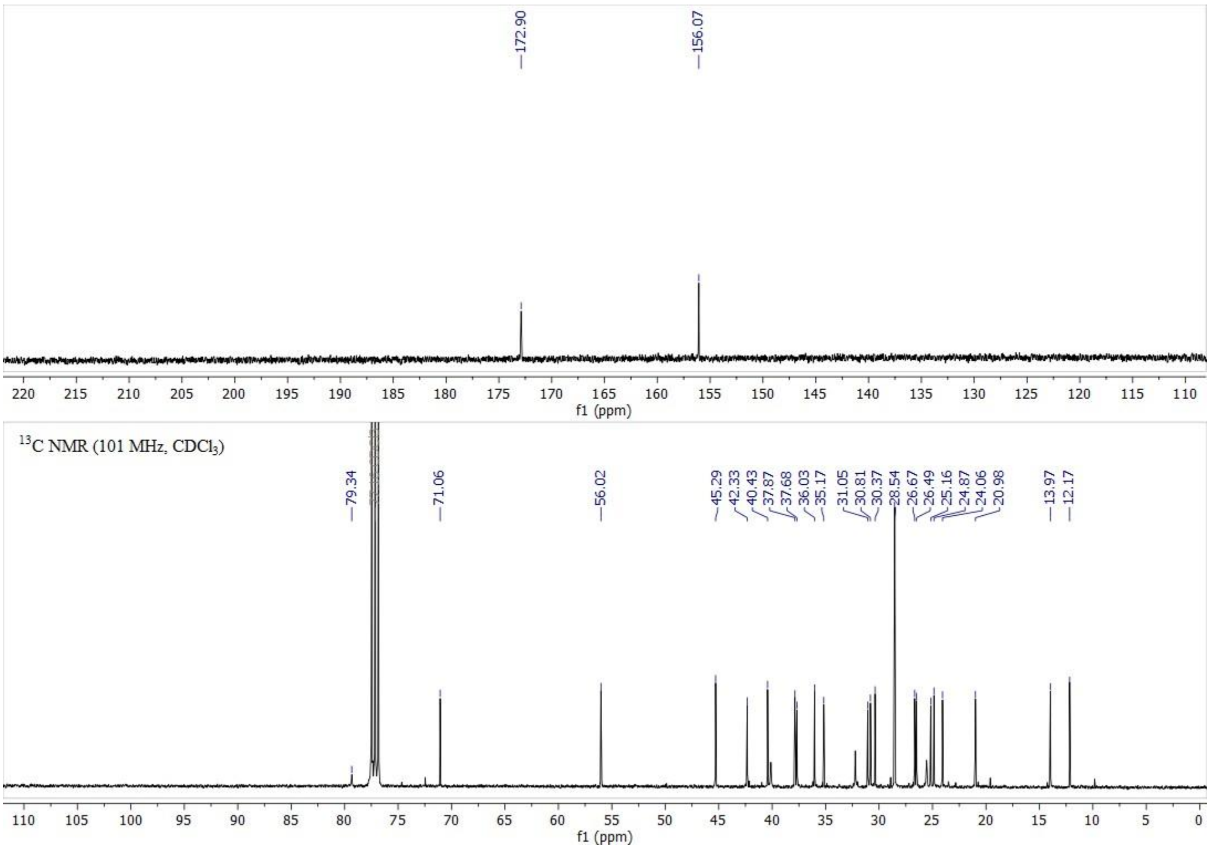
172.90
156.07
f1 (ppm)
13C NMR (101 MHz, CDCl3)
79.34
71.06
56.02
45.29
42.33
40.43
37.87
37.68
36.03
35.17
31.05
30.81
30.37
28.54
26.67
26.49
25.16
24.87
24.06
20.98
13.97
12.17
f1 (ppm)

**27.** **55**.

**Figure S** $^{1}H$ NMR (top) and $^{13}C$ NMR (bottom) spectra of compound

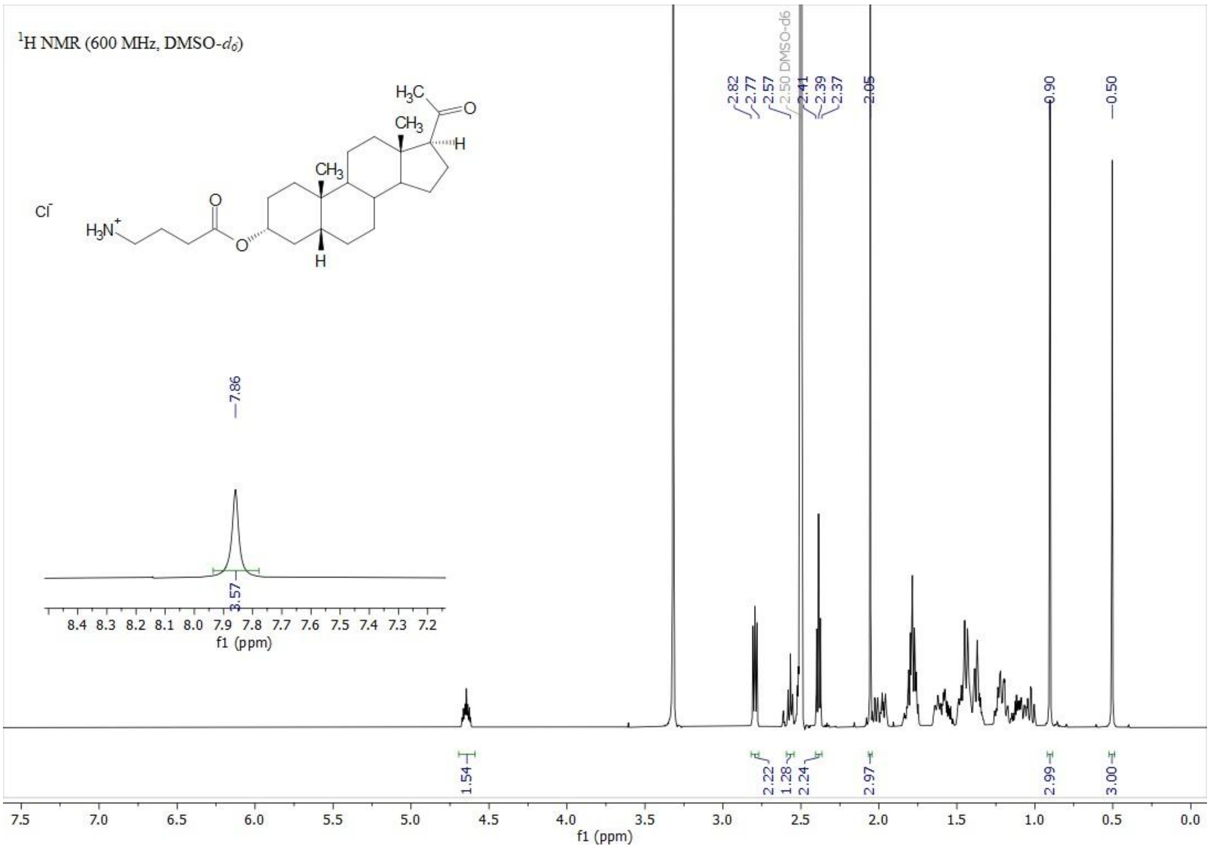
1H NMR (600 MHz, DMSO-d6)
Cl-
H3N+
H3C
CH3
CH3
H
H
O
2.82
2.77
2.57
2.50 DMSO-d6
2.41
2.39
2.37
2.05
0.90
0.50
7.86
3.57
8.4 8.3 8.2 8.1 8.0 7.9 7.8 7.7 7.6 7.5 7.4 7.3 7.2
f1 (ppm)
1.54
2.22
1.28
2.24
2.97
2.99
3.00
7.5 7.0 6.5 6.0 5.5 5.0 4.5 4.0 3.5 3.0 2.5 2.0 1.5 1.0 0.5 0.0
f1 (ppm)

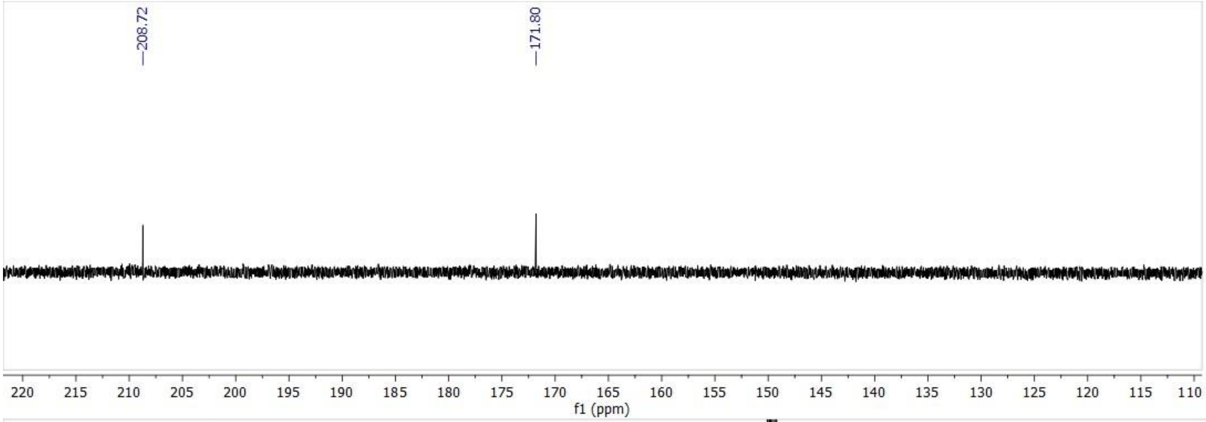
208.72
171.80
220 215 210 205 200 195 190 185 180 175 170 165 160 155 150 145 140 135 130 125 120 115 110
f1 (ppm)

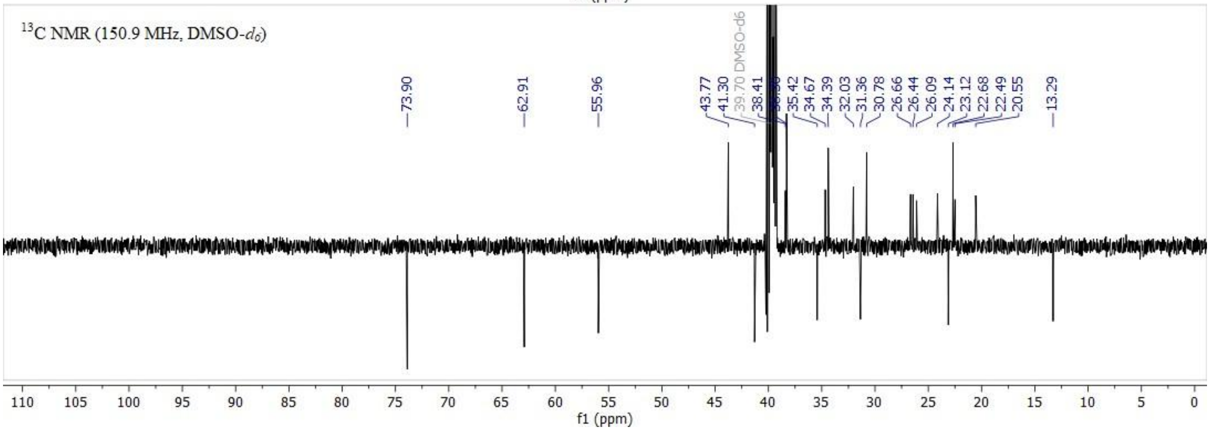
13C NMR (150.9 MHz, DMSO-d6)
73.90
62.91
55.96
43.77
41.30
39.70 DMSO-d6
38.41
35.42
34.67
34.39
32.03
31.36
30.78
26.66
26.44
26.09
24.14
23.12
22.68
22.49
20.55
13.29
110 105 100 95 90 85 80 75 70 65 60 55 50 45 40 35 30 25 20 15 10 5 0
f1 (ppm)

**28.** **43**.

**Figure S** $^{1}H$ NMR (top) and $^{13}C$ NMR (bottom) spectra of compound

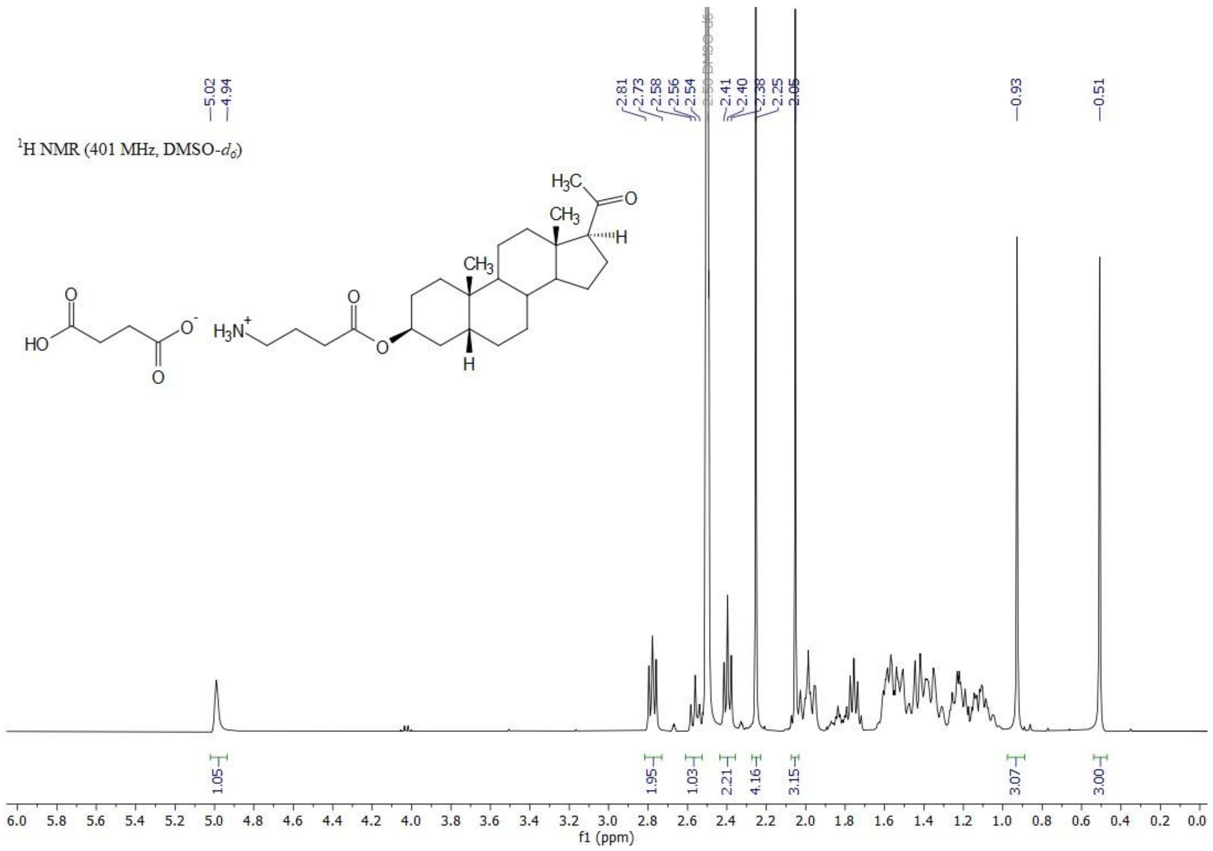

1H NMR (401 MHz, DMSO-d6)
5.02
4.94
2.81
2.73
2.58
2.56
2.54
2.50 DMSO-d6
2.41
2.40
2.38
2.25
2.05
0.93
0.51
1.05
1.95
1.03
2.21
4.16
3.15
3.07
3.00
f1 (ppm)
H3C
CH3
CH3
H
HO
O
O-
H3N+

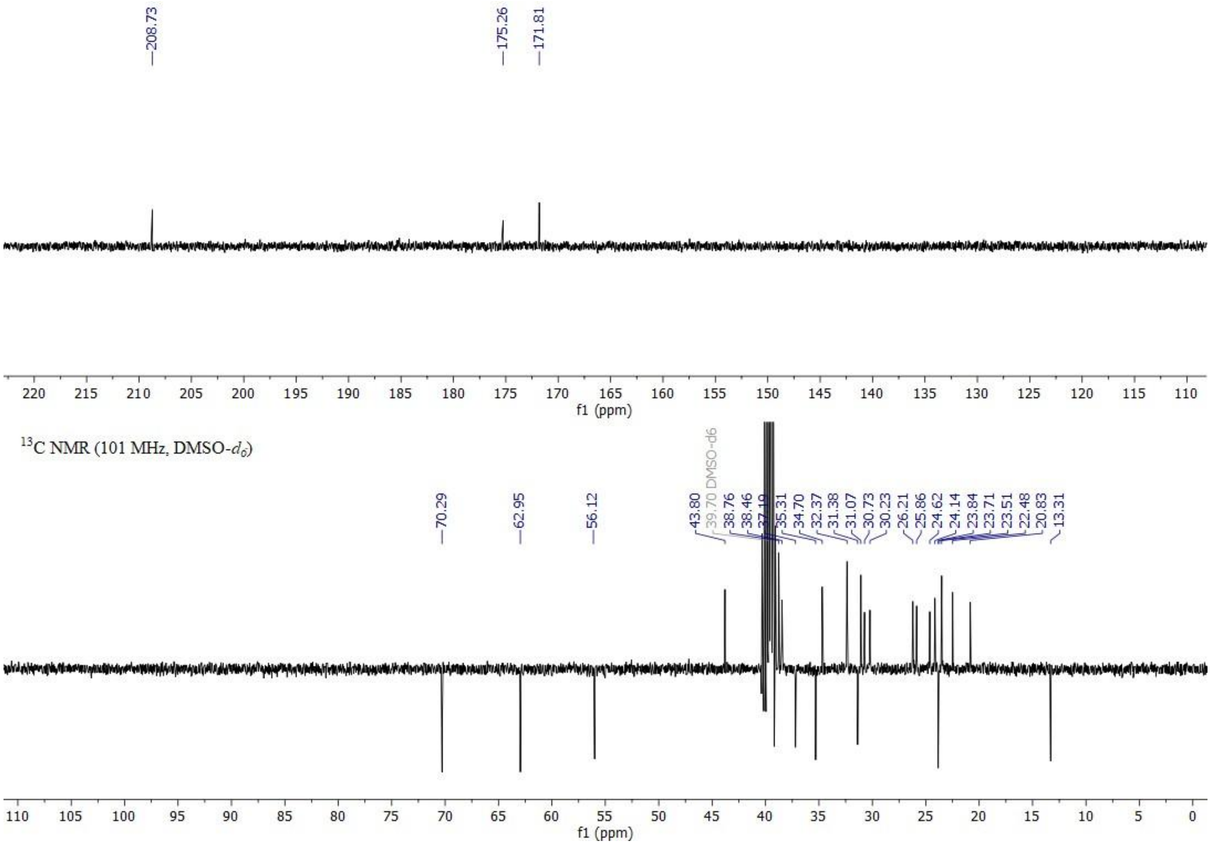

208.73
175.26
171.81
f1 (ppm)
13C NMR (101 MHz, DMSO-d6)
70.29
62.95
56.12
43.80
39.70 DMSO-d6
38.76
38.46
35.31
34.70
32.37
31.38
31.07
30.73
30.23
26.21
25.86
24.62
24.14
23.84
23.71
23.51
22.48
20.83
13.31
f1 (ppm)

**29.** **44a**.

**Figure S** $^{1}H$ NMR (top) and $^{13}C$ NMR (bottom) spectra of compound

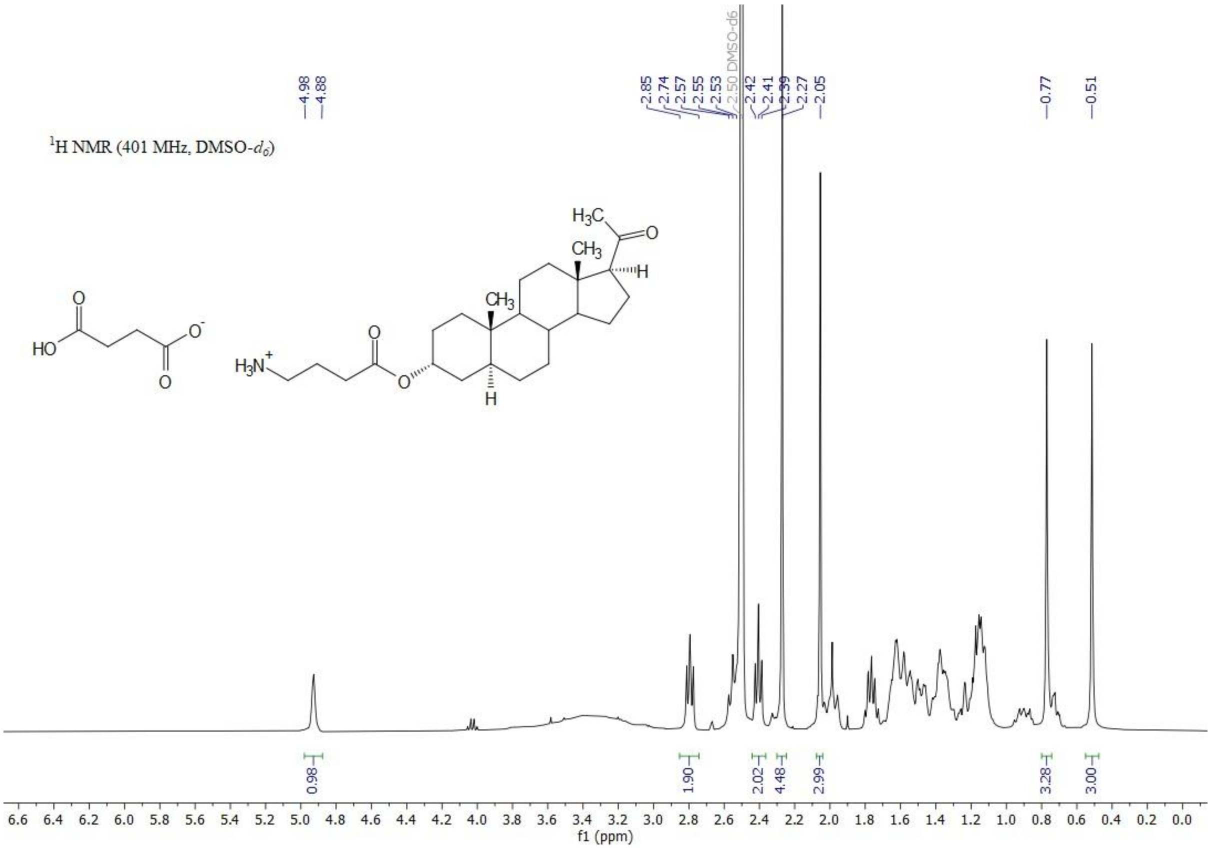
$^{1}$H NMR (401 MHz, DMSO-$d_6$)
4.98
4.88
2.85
2.74
2.57
2.55
2.53
2.50 DMSO-d6
2.42
2.41
2.39
2.27
2.05
0.77
0.51
0.98
1.90
2.02
4.48
2.99
3.28
3.00
f1 (ppm)

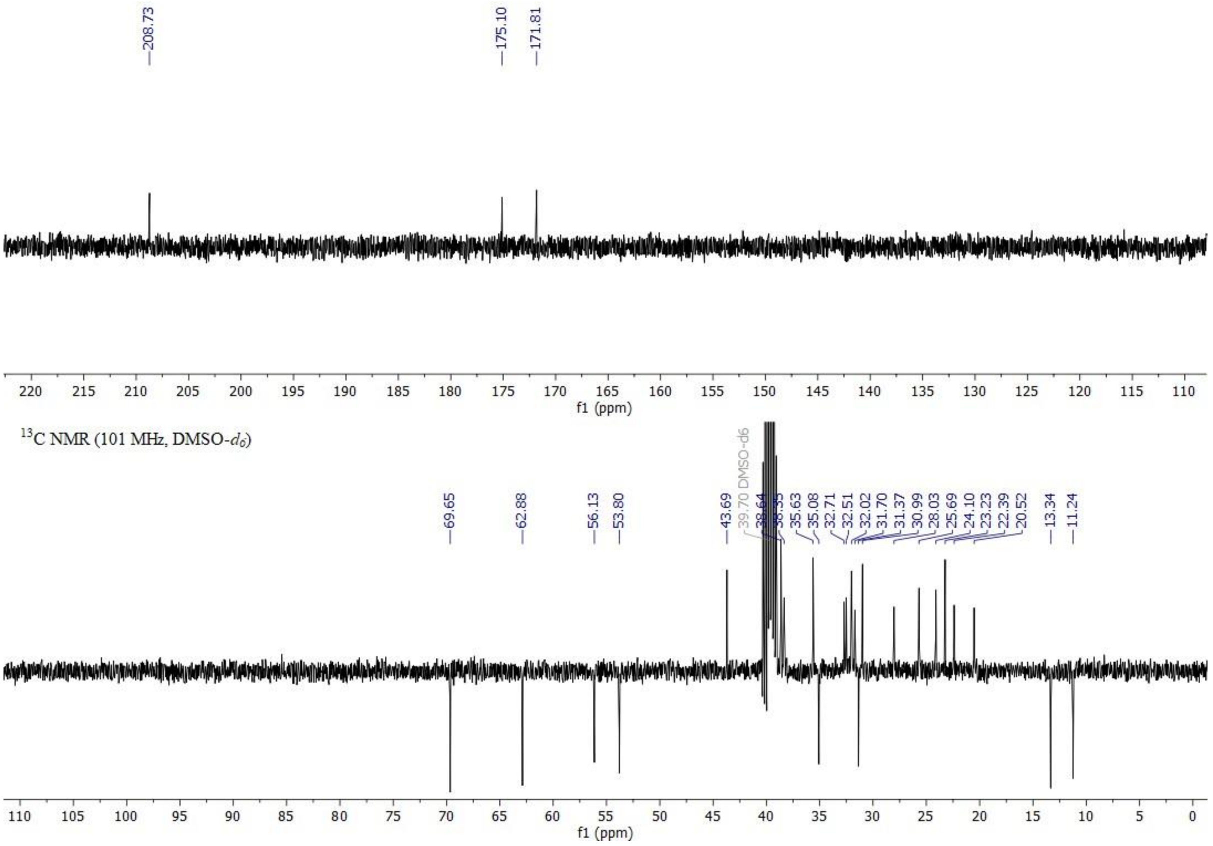
208.73
175.10
171.81
f1 (ppm)
$^{13}$C NMR (101 MHz, DMSO-$d_6$)
69.65
62.88
56.13
53.80
43.69
39.70 DMSO-d6
35.63
35.08
32.71
32.51
32.02
31.70
31.37
30.99
28.03
25.69
24.10
23.23
22.39
20.52
13.34
11.24
f1 (ppm)

**30.** **45a**.

**Figure S** [1]H NMR (top) and [13]C NMR (bottom) spectra of compound

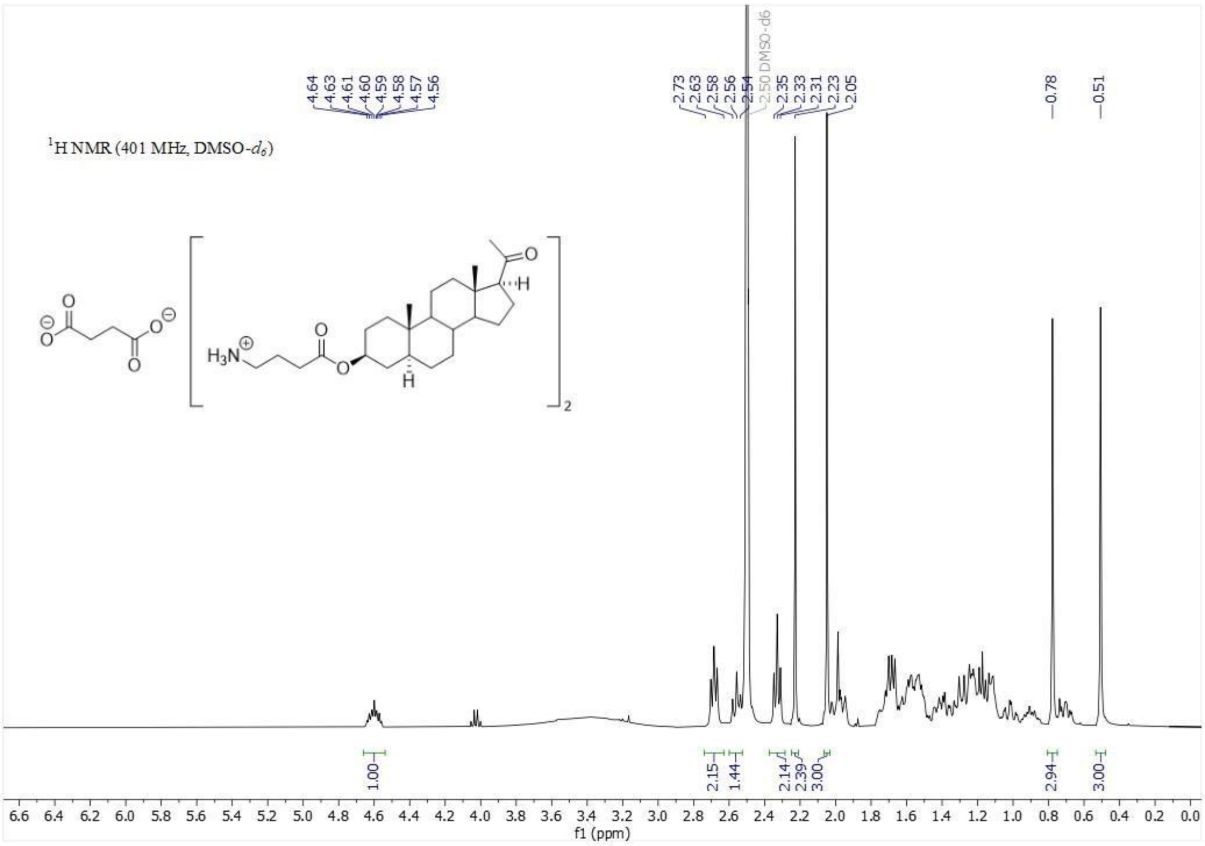

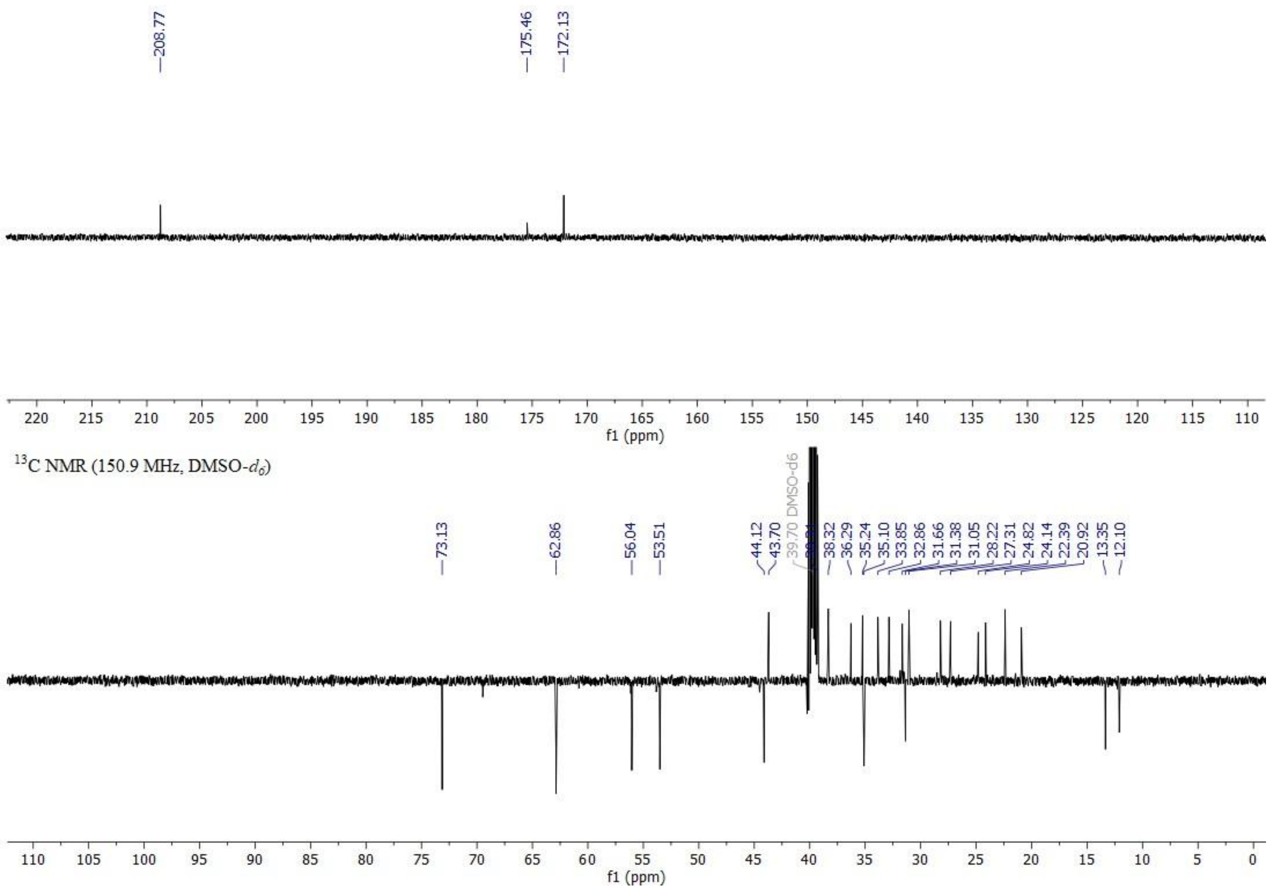

**Figure S31.** $^1$H NMR (top) and $^{13}$C NMR (bottom) spectra of compound **46a**.

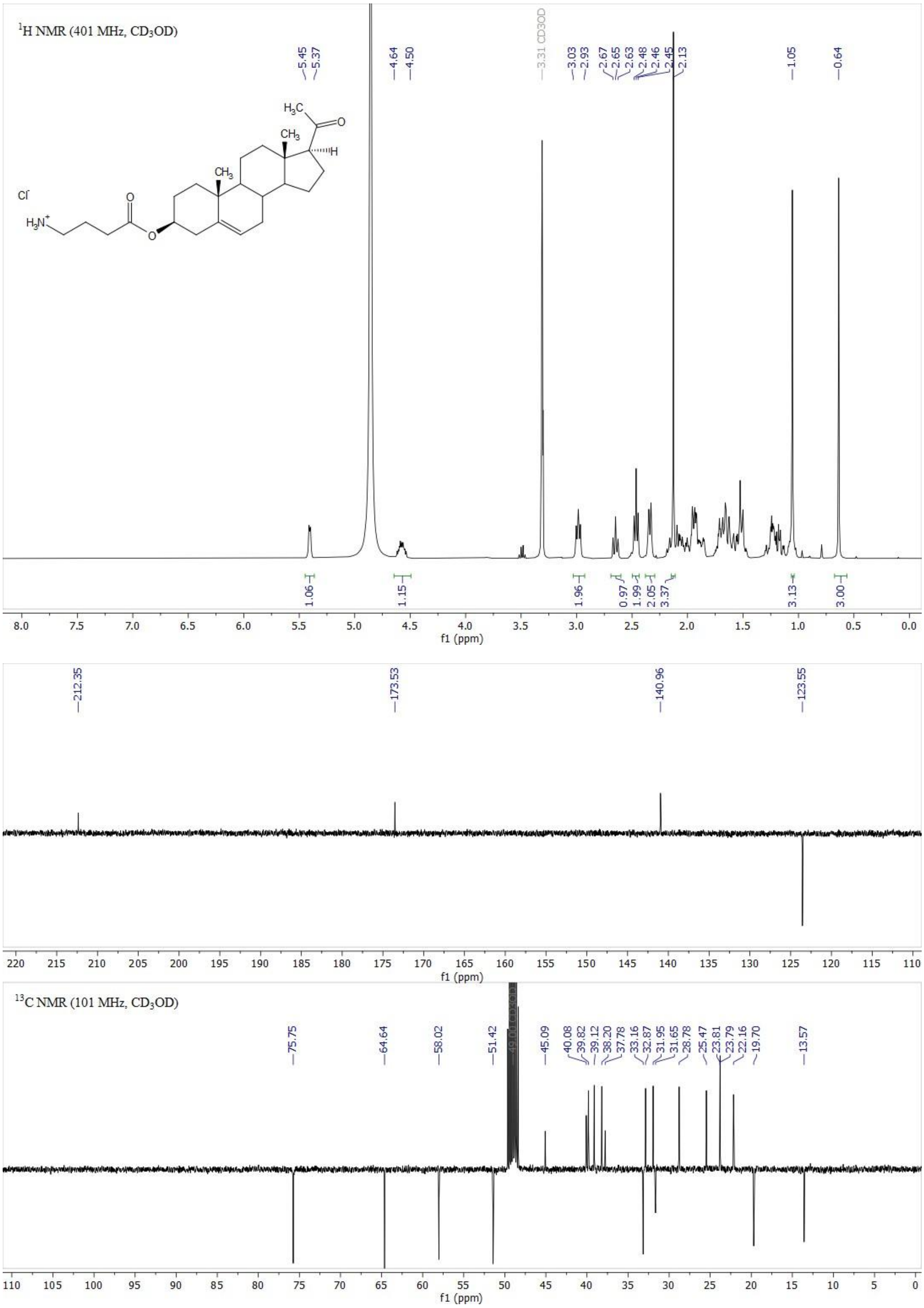

**Figure S32.** $^{1}$H NMR (top) and $^{13}$C NMR (bottom) spectra of compound **49**.

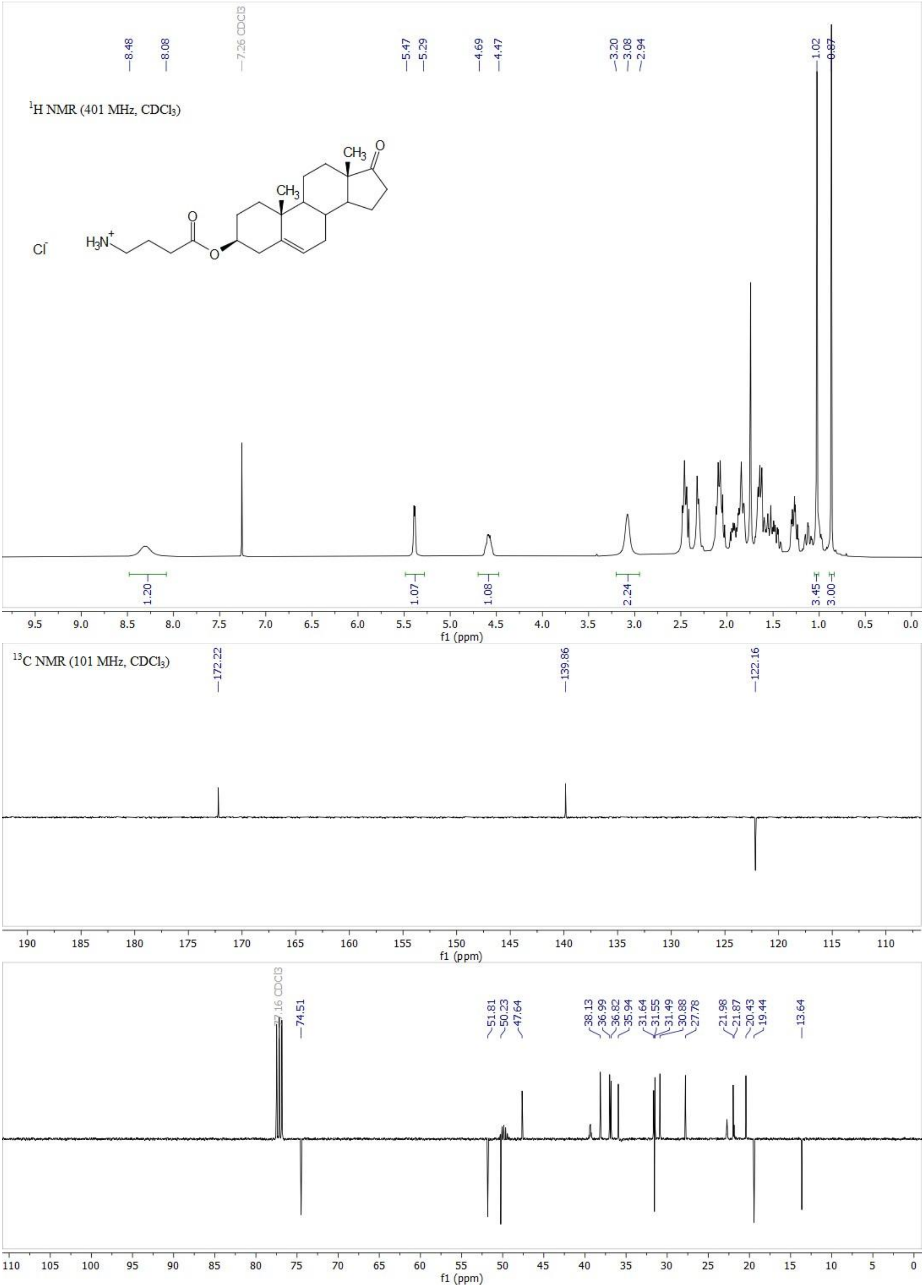

**Figure S33.** $^{1}H$ NMR (top) and $^{13}C$ NMR (bottom) spectra of compound **52a**.

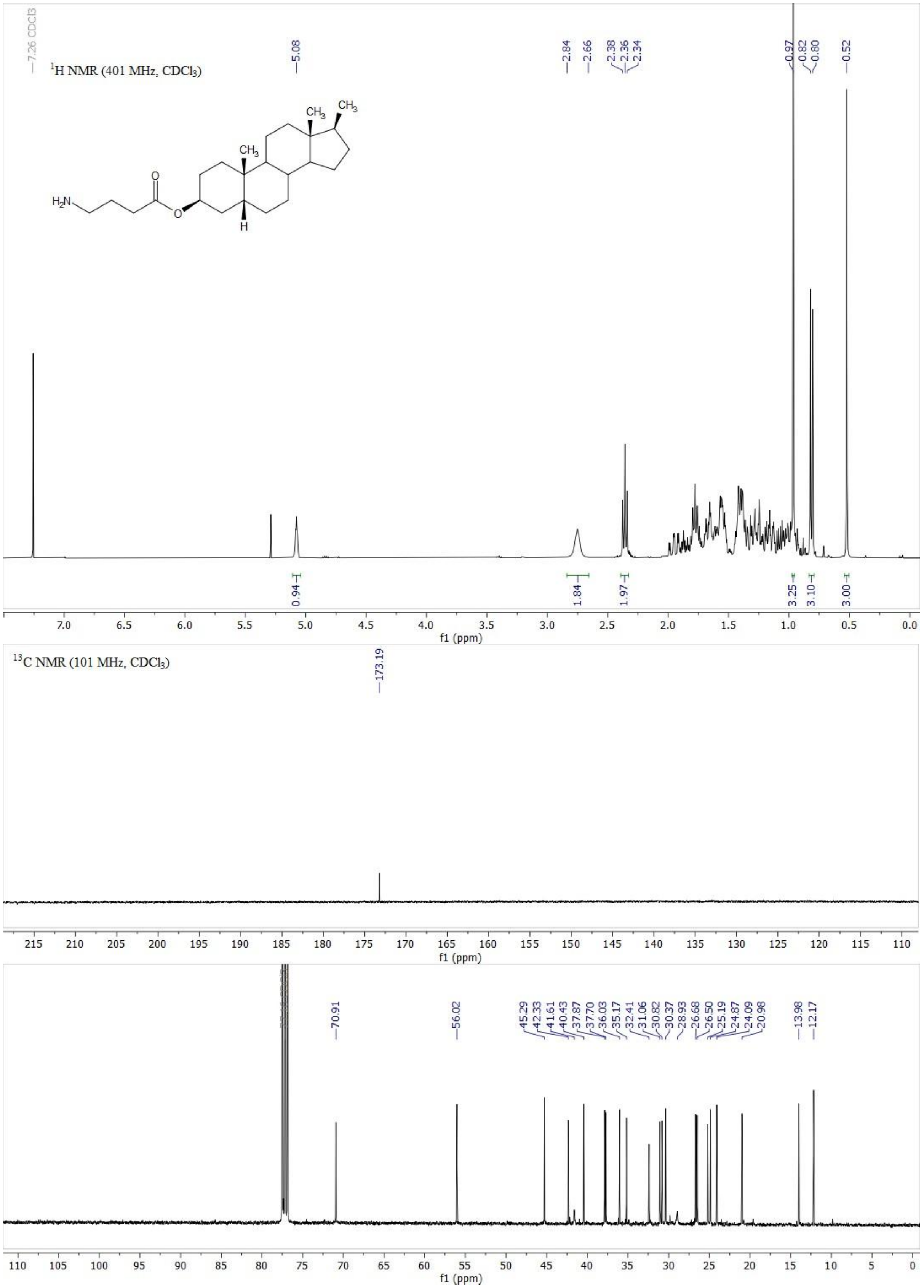

**Figure S34.** $^{1}H$ NMR (top) and $^{13}C$ NMR (bottom) spectra of compound **56**.

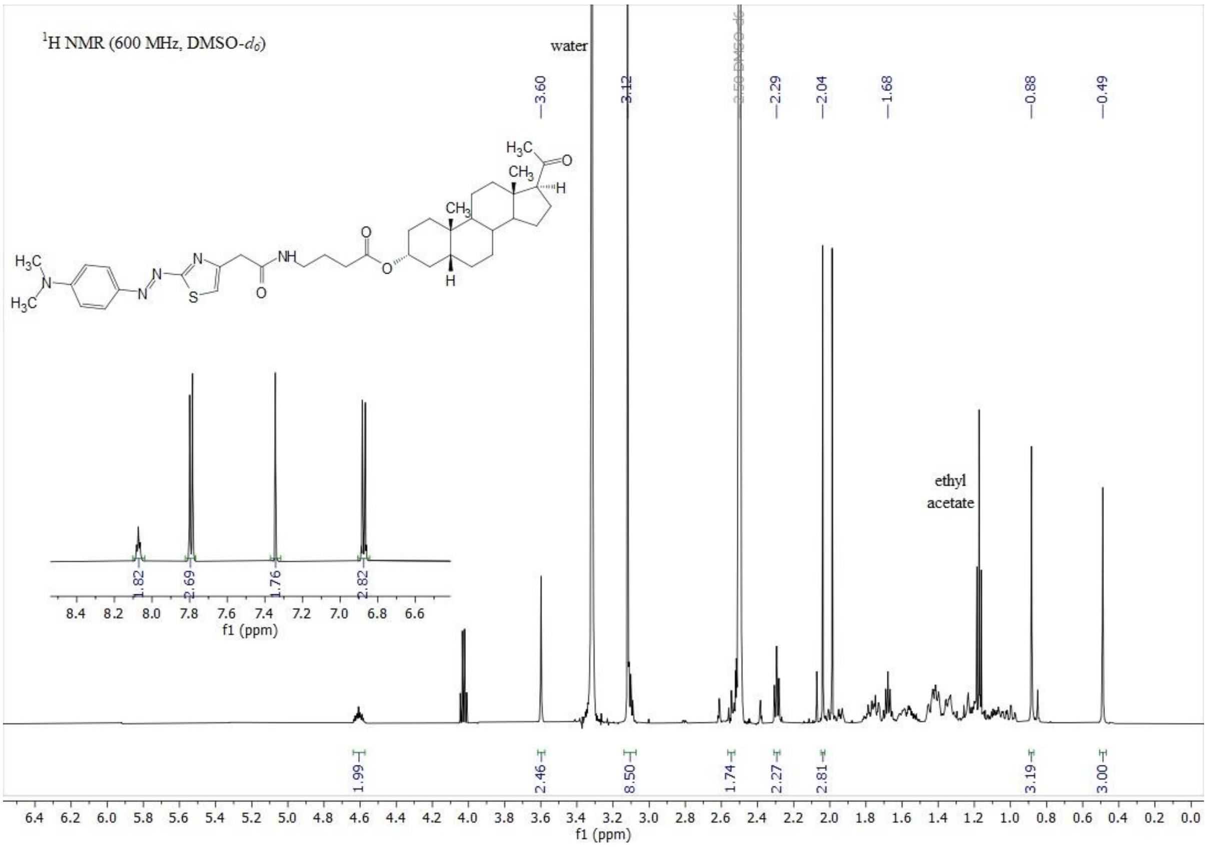

$^{1}$H NMR (600 MHz, DMSO-$d_6$)
water
3.60
3.12
2.50 DMSO-d6
2.29
2.04
1.68
0.88
0.49
ethyl acetate
1.82
2.69
1.76
2.82
1.99
2.46
8.50
1.74
2.27
2.81
3.19
3.00
f1 (ppm)

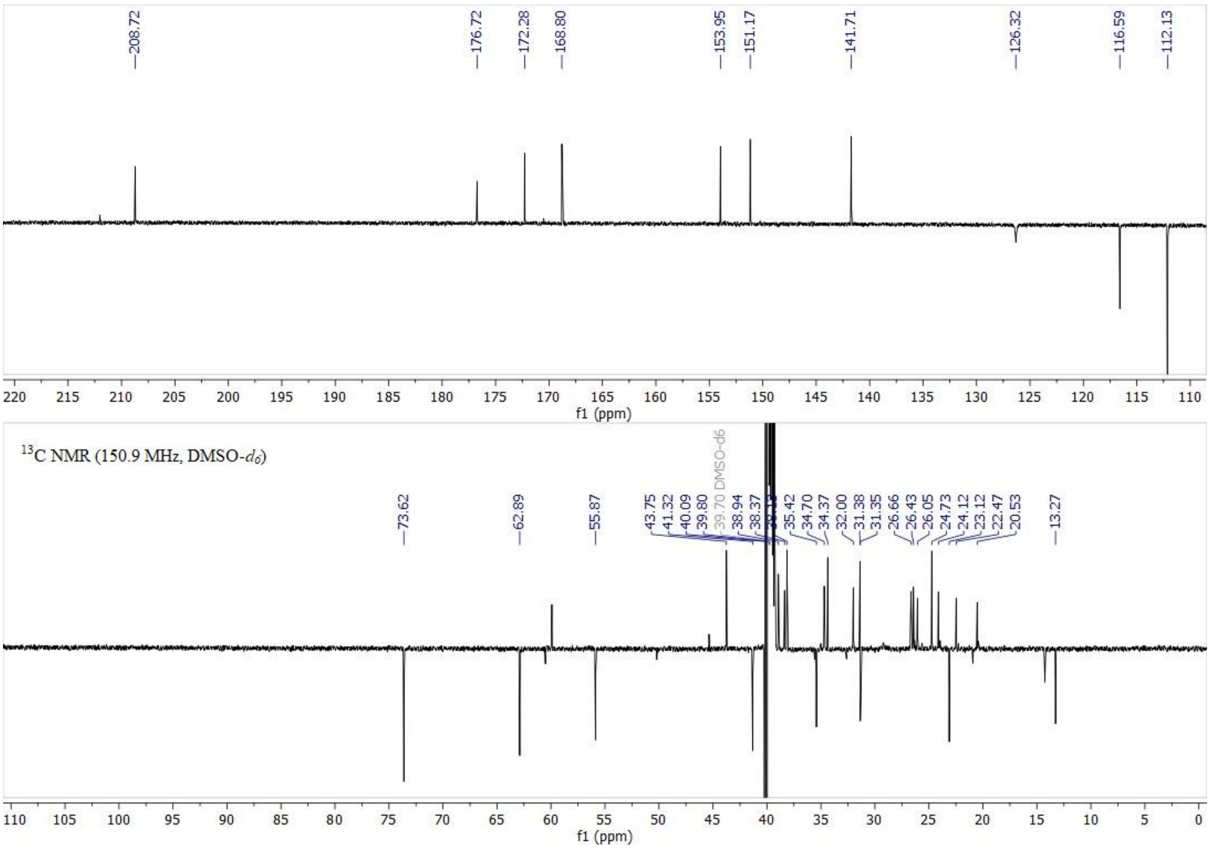

208.72
176.72
172.28
168.80
153.95
151.17
141.71
126.32
116.59
112.13
f1 (ppm)
$^{13}$C NMR (150.9 MHz, DMSO-$d_6$)
73.62
62.89
55.87
43.75
41.32
40.09
39.80
39.70 DMSO-d6
38.94
38.37
35.42
34.70
34.37
32.00
31.38
31.35
26.66
26.43
26.05
24.73
24.12
23.12
22.47
20.53
13.27
f1 (ppm)

**35.** C APT NMR (bottom) spectra of compound **7**.

**Figure S** $^{1}$H NMR (top) and $^{13}$

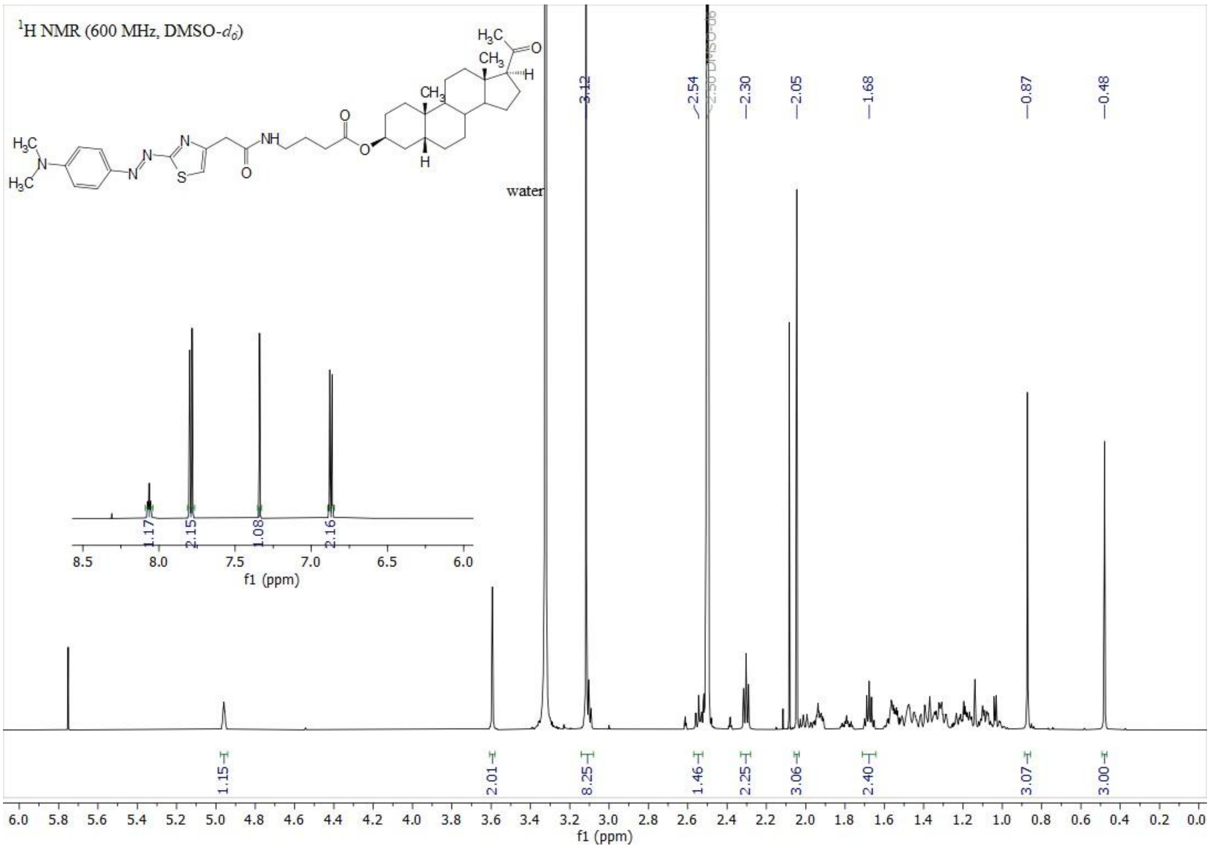

$^1$H NMR (600 MHz, DMSO-$d_6$)
water
3.12
2.54
2.50 DMSO-d6
2.30
2.05
1.68
0.87
0.48
1.17
2.15
1.08
2.16
1.15
2.01
8.25
1.46
2.25
3.06
2.40
3.07
3.00
f1 (ppm)

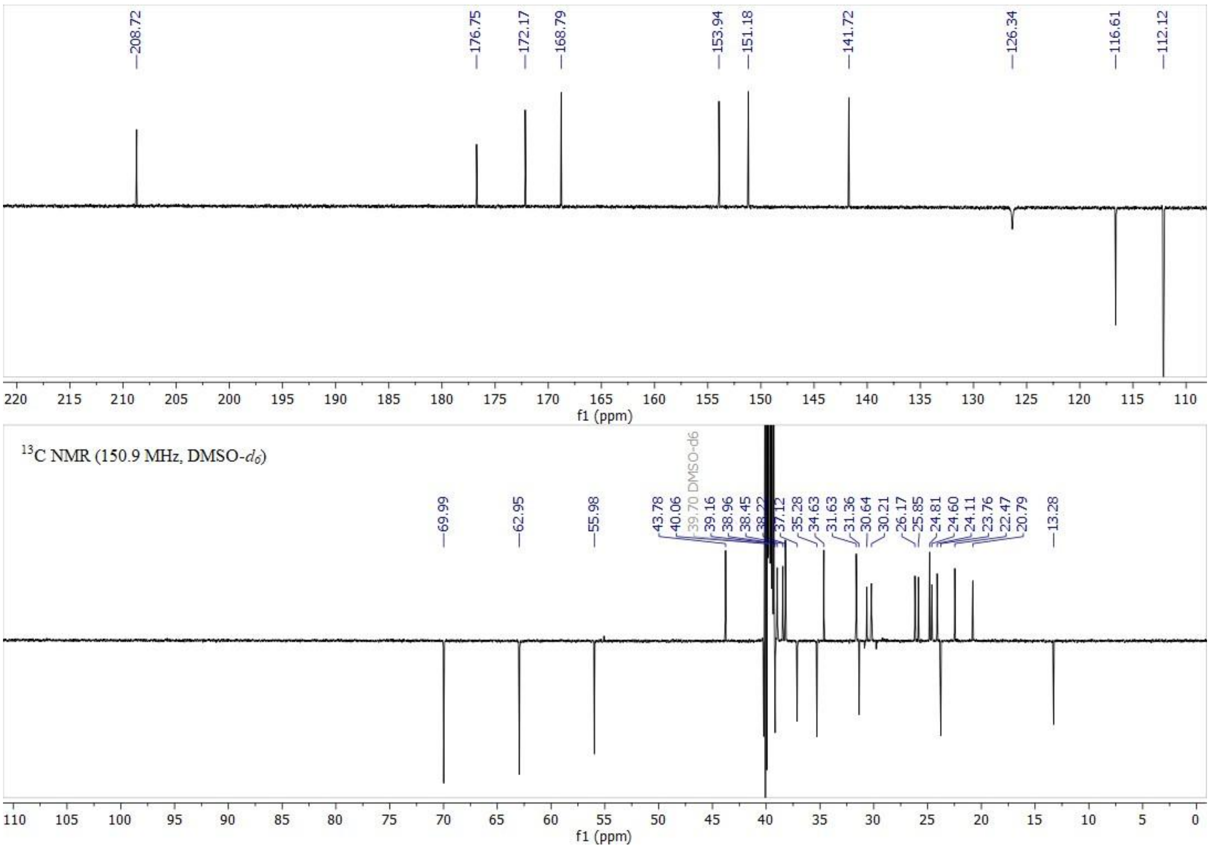

208.72
176.75
172.17
168.79
153.94
151.18
141.72
126.34
116.61
112.12
$^{13}$C NMR (150.9 MHz, DMSO-$d_6$)
69.99
62.95
55.98
43.78
40.06
39.70 DMSO-d6
39.16
38.96
38.45
38.22
37.12
35.28
34.63
31.63
31.36
30.64
30.21
26.17
25.85
24.81
24.60
24.11
23.76
22.47
20.79
13.28
f1 (ppm)

**36.** C APT NMR (bottom) spectra of compound **8**.

**Figure S** $^{1}$H NMR (top) and $^{13}$

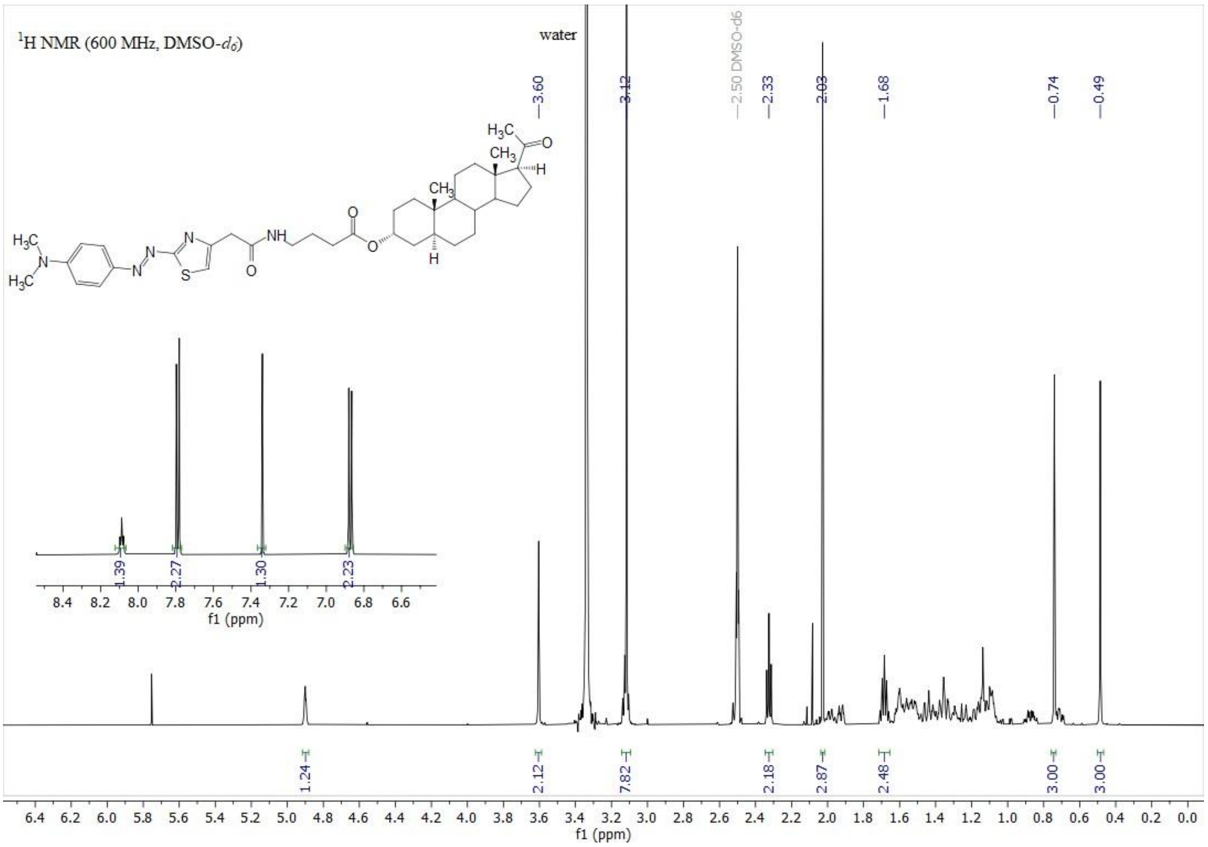

[1]H NMR (600 MHz, DMSO-$d_6$)
water
3.60
3.12
2.50 DMSO-d6
2.33
2.03
1.68
0.74
0.49
1.39
2.27
1.30
2.23
1.24
2.12
7.82
2.18
2.87
2.48
3.00
3.00
f1 (ppm)

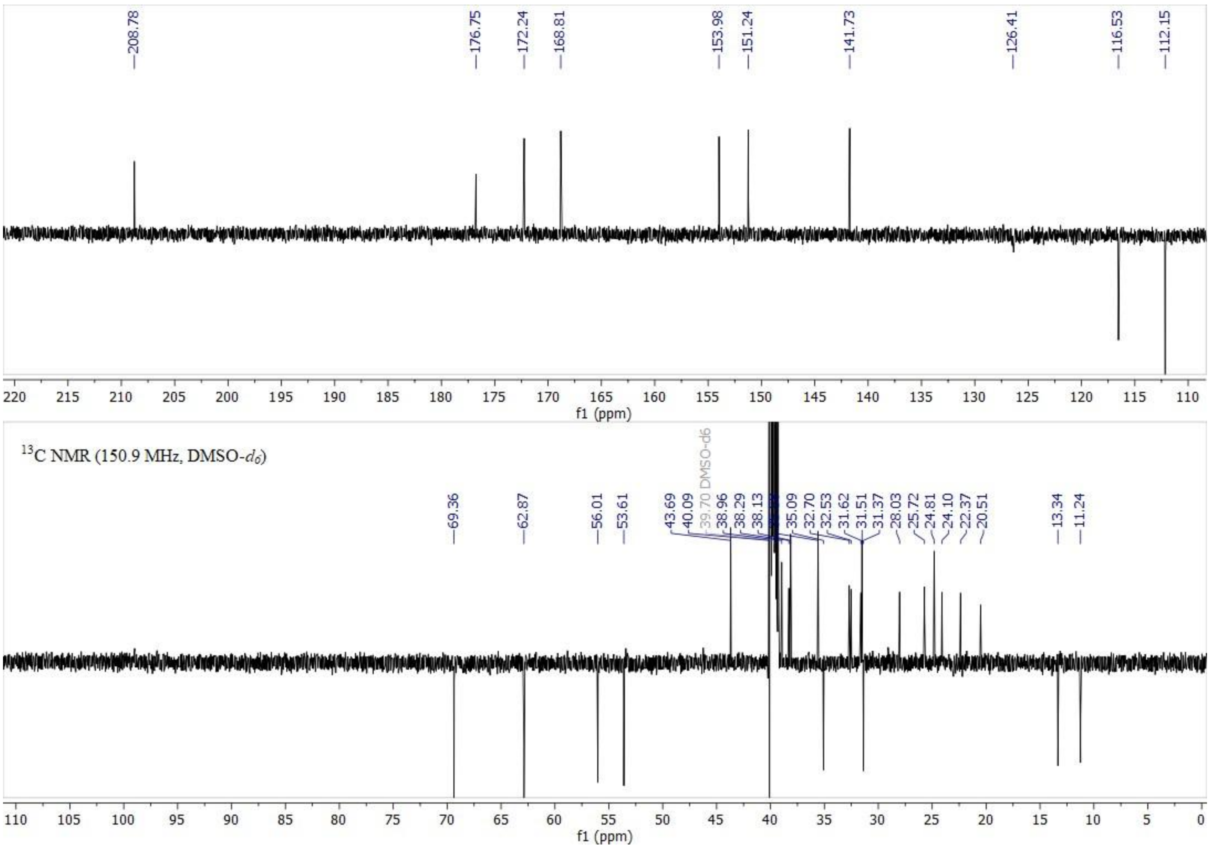

208.78
176.75
172.24
168.81
153.98
151.24
141.73
126.41
116.53
112.15
f1 (ppm)
[13]C NMR (150.9 MHz, DMSO-$d_6$)
69.36
62.87
56.01
53.61
43.69
40.09
39.70 DMSO-d6
38.96
38.29
38.13
35.09
32.70
32.53
31.62
31.51
31.37
28.03
25.72
24.81
24.10
22.37
20.51
13.34
11.24
f1 (ppm)

**37.** C APT NMR (bottom) spectra of compound **9**.

**Figure S** $^{1}$H NMR (top) and $^{13}$

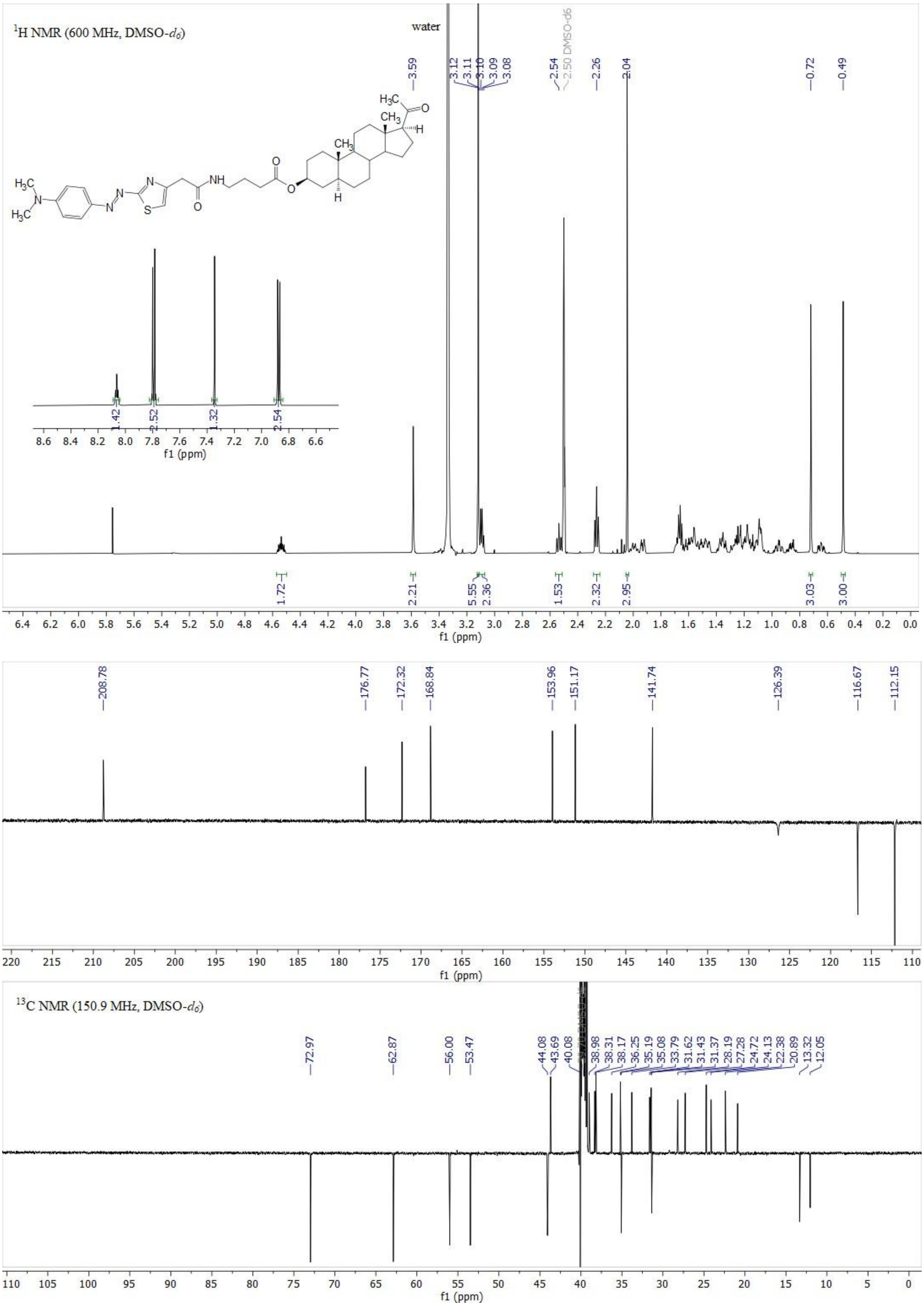

1H NMR (600 MHz, DMSO-d6)
water
3.59
3.12
3.11
3.10
3.09
3.08
2.54
2.50 DMSO-d6
2.26
2.04
0.72
0.49
1.42
2.52
1.32
2.54
f1 (ppm)
1.72
2.21
5.55
2.36
1.53
2.32
2.95
3.03
3.00
f1 (ppm)
208.78
176.77
172.32
168.84
153.96
151.17
141.74
126.39
116.67
112.15
f1 (ppm)
13C NMR (150.9 MHz, DMSO-d6)
72.97
62.87
56.00
53.47
44.08
43.69
40.08
38.98
38.31
38.17
36.25
35.19
35.08
33.79
31.62
31.43
31.37
28.19
27.28
24.72
24.13
22.38
20.89
13.32
12.05
f1 (ppm)

**Figure S38.** $^{1}$H NMR (top) and $^{13}$C APT NMR (bottom) spectra of compound **10**.

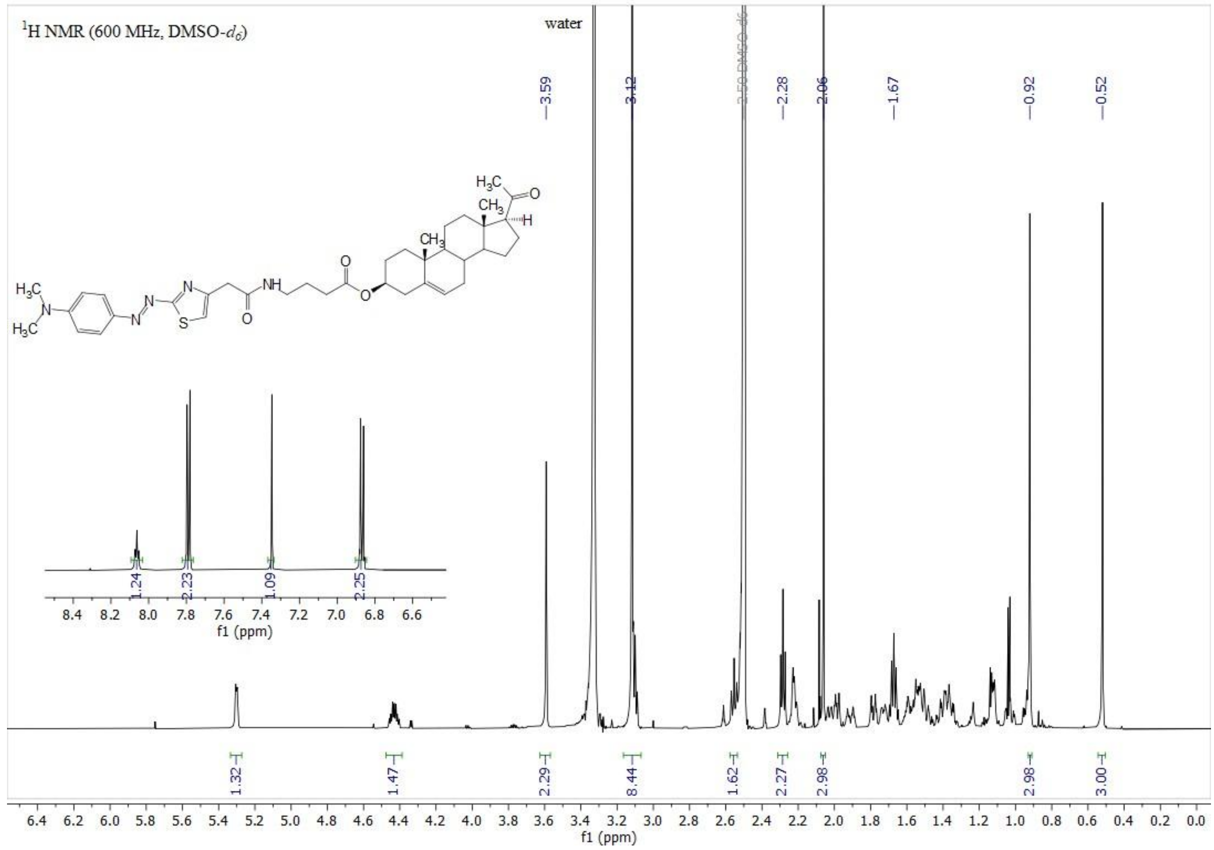

**Figure S** $^1$H NMR (top) and $^{13}$

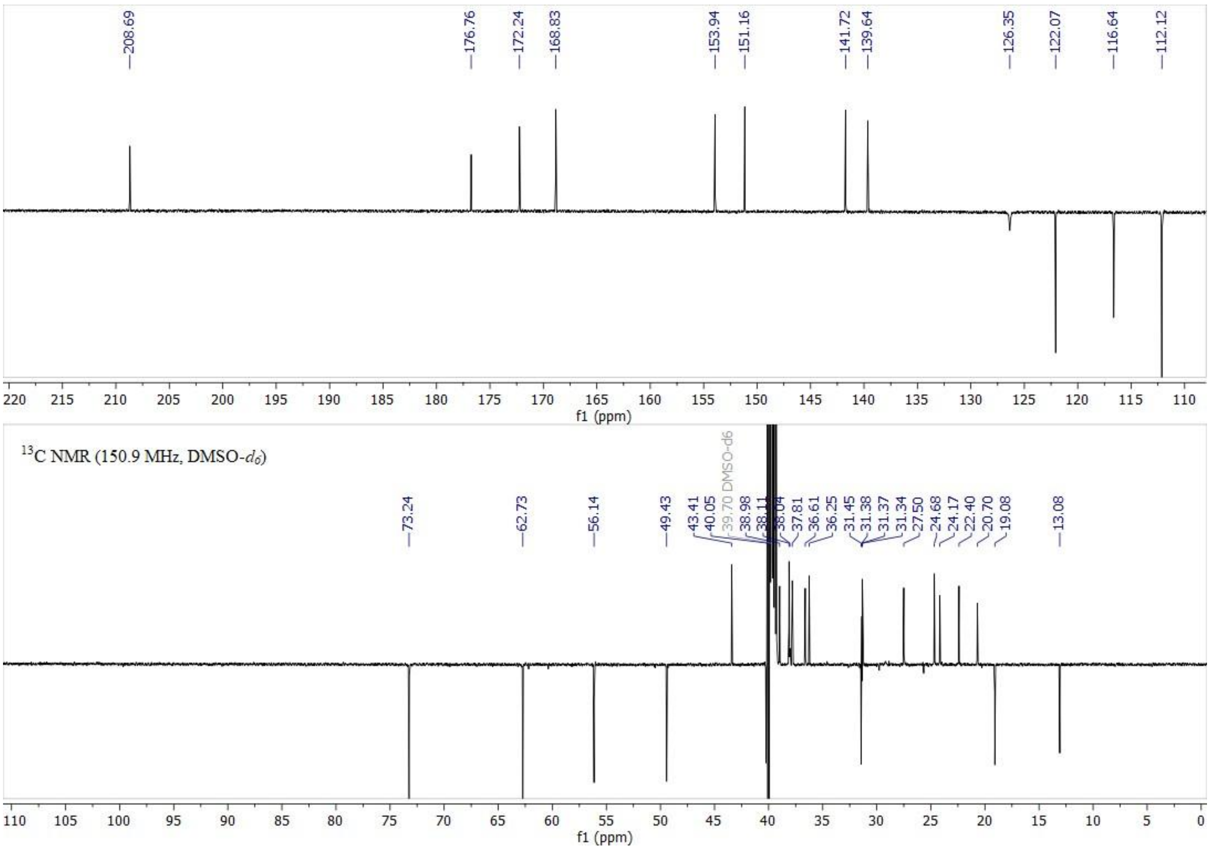

**Figure S39.** $^{1}H$ NMR (top) and $^{13}C$ APT NMR (bottom) spectra of compound **15**.

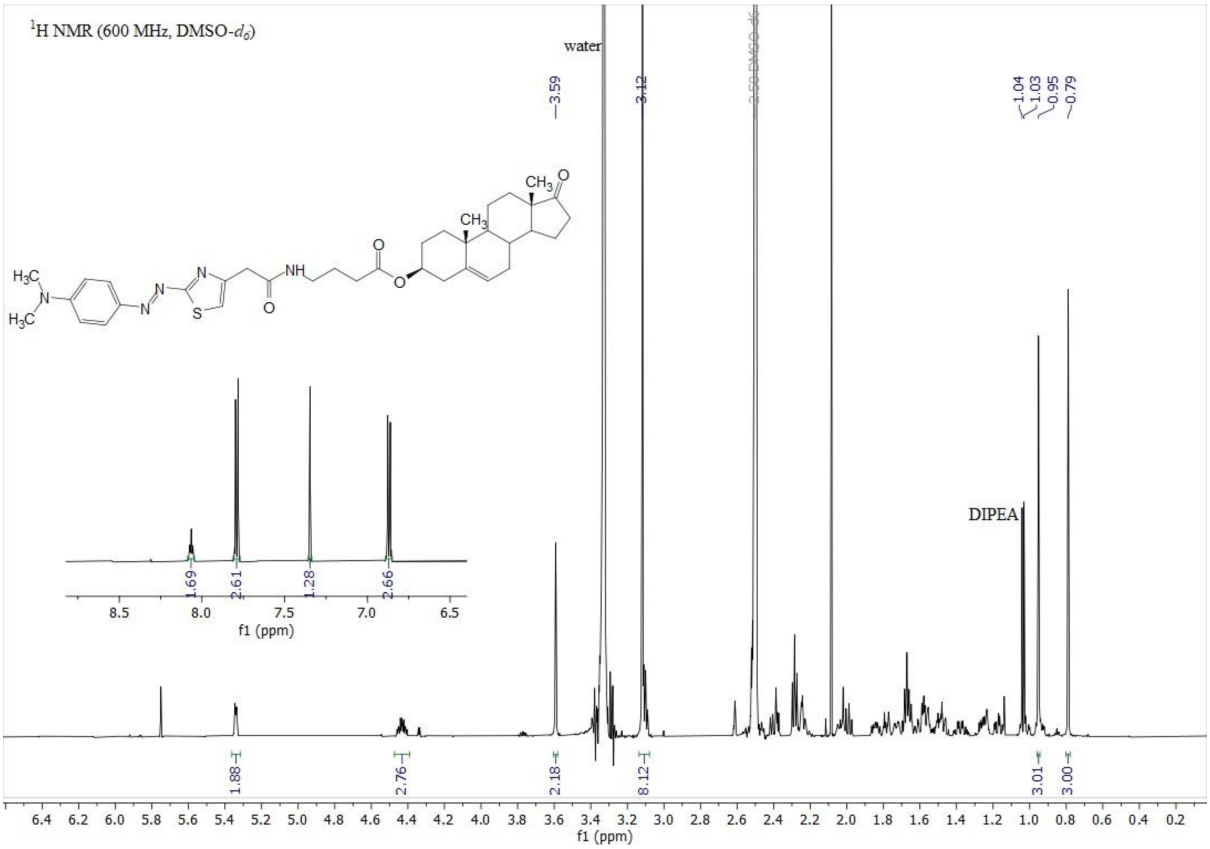

$^{1}$H NMR (600 MHz, DMSO-$d_6$)
water
3.59
3.12
2.50 DMSO-d6
1.04
1.03
0.95
0.79
DIPEA
1.69
2.61
1.28
2.66
8.5
8.0
7.5
7.0
6.5
f1 (ppm)
1.88
2.76
2.18
8.12
3.01
3.00
6.4 6.2 6.0 5.8 5.6 5.4 5.2 5.0 4.8 4.6 4.4 4.2 4.0 3.8 3.6 3.4 3.2 3.0 2.8 2.6 2.4 2.2 2.0 1.8 1.6 1.4 1.2 1.0 0.8 0.6 0.4 0.2
f1 (ppm)

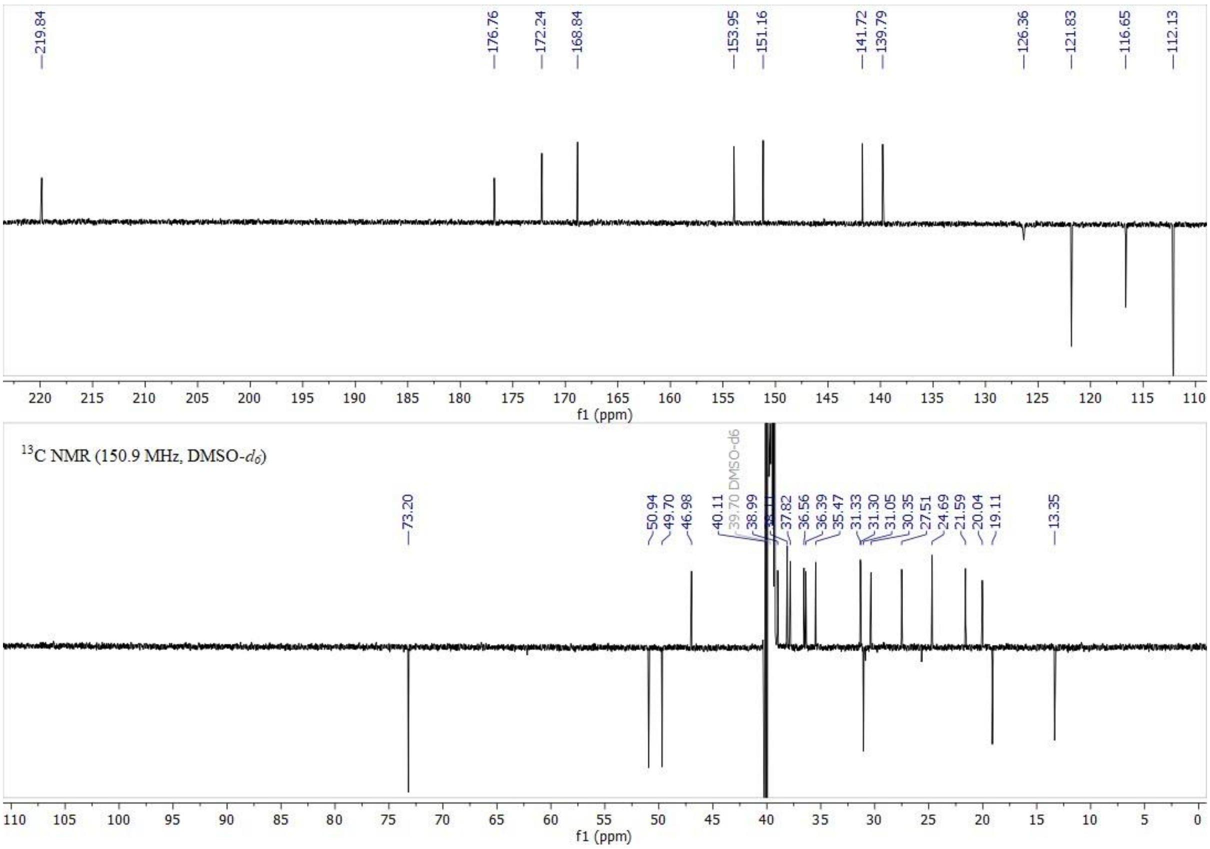

219.84
176.76
172.24
168.84
153.95
151.16
141.72
139.79
126.36
121.83
116.65
112.13
220 215 210 205 200 195 190 185 180 175 170 165 160 155 150 145 140 135 130 125 120 115 110
f1 (ppm)
$^{13}$C NMR (150.9 MHz, DMSO-$d_6$)
73.20
50.94
49.70
46.98
40.11
39.70 DMSO-d6
38.99
37.82
36.56
36.39
35.47
31.33
31.30
31.05
30.35
27.51
24.69
21.59
20.04
19.11
13.35
110 105 100 95 90 85 80 75 70 65 60 55 50 45 40 35 30 25 20 15 10 5 0
f1 (ppm)

**40.** C APT NMR (bottom) spectra of compound **16**.

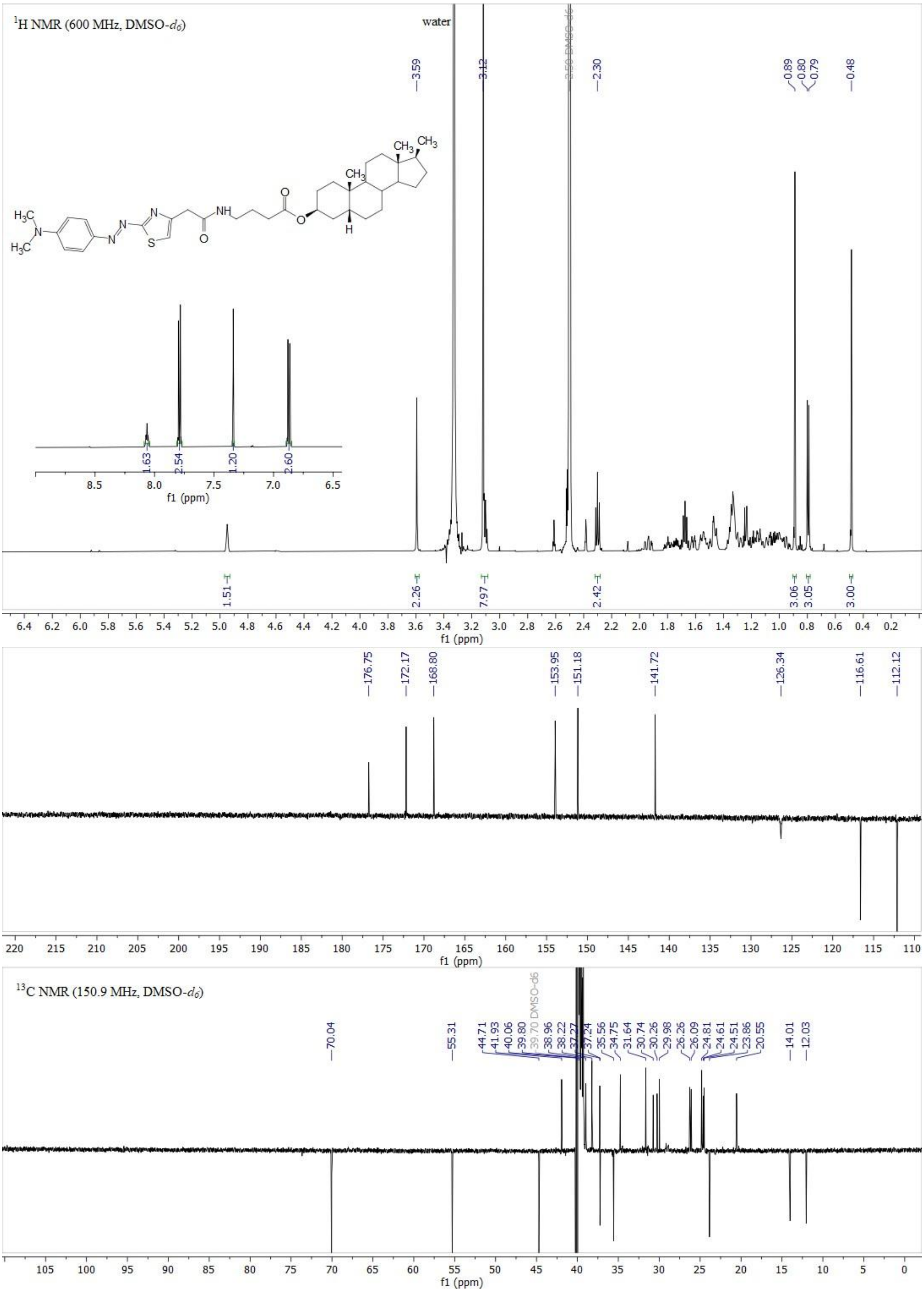

**Figure S41.** $^1$H NMR (top) and $^{13}$C APT NMR (bottom) spectra of compound **17**.

**Figure S** $^1$H NMR (top) and $^{13}$

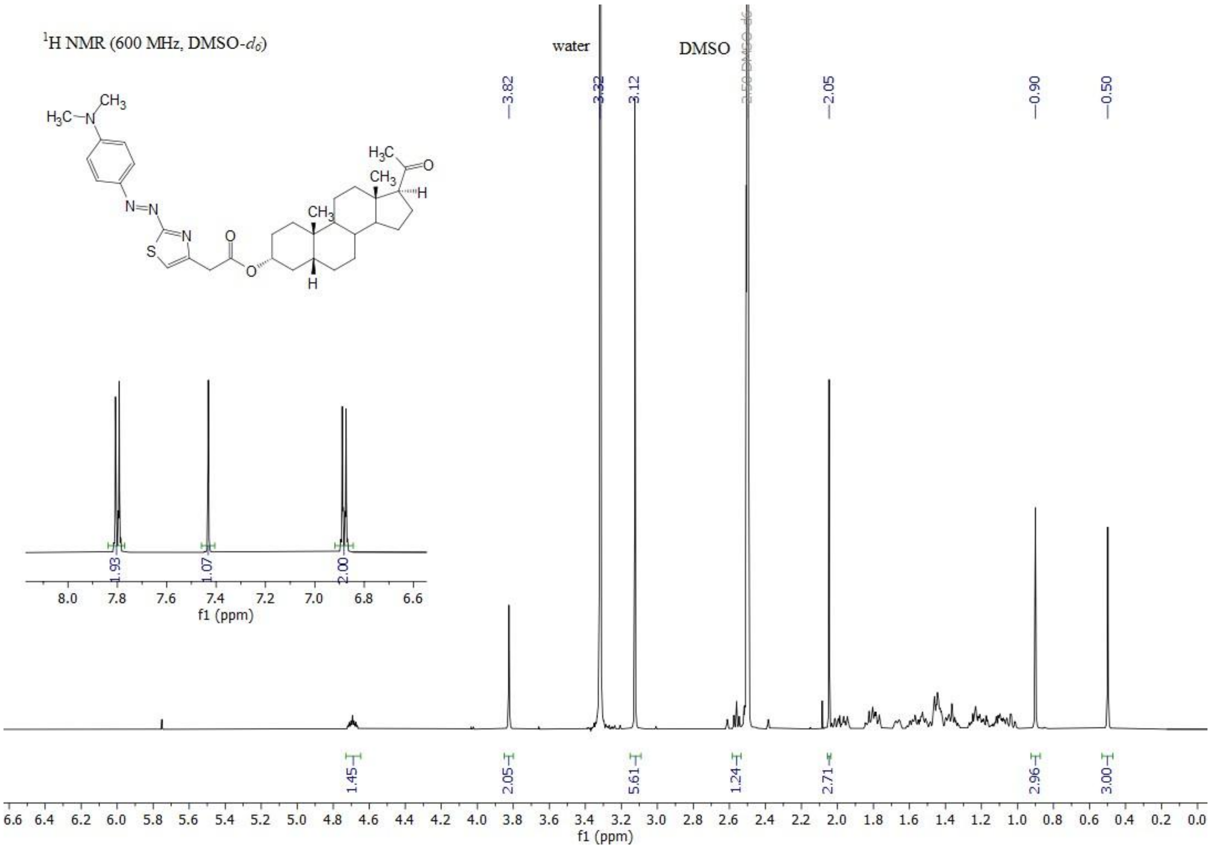

1H NMR (600 MHz, DMSO-d6)
water
DMSO
3.82
3.32
3.12
2.05
0.90
0.50
1.93
1.07
2.00
1.45
2.05
5.61
1.24
2.71
2.96
3.00
f1 (ppm)

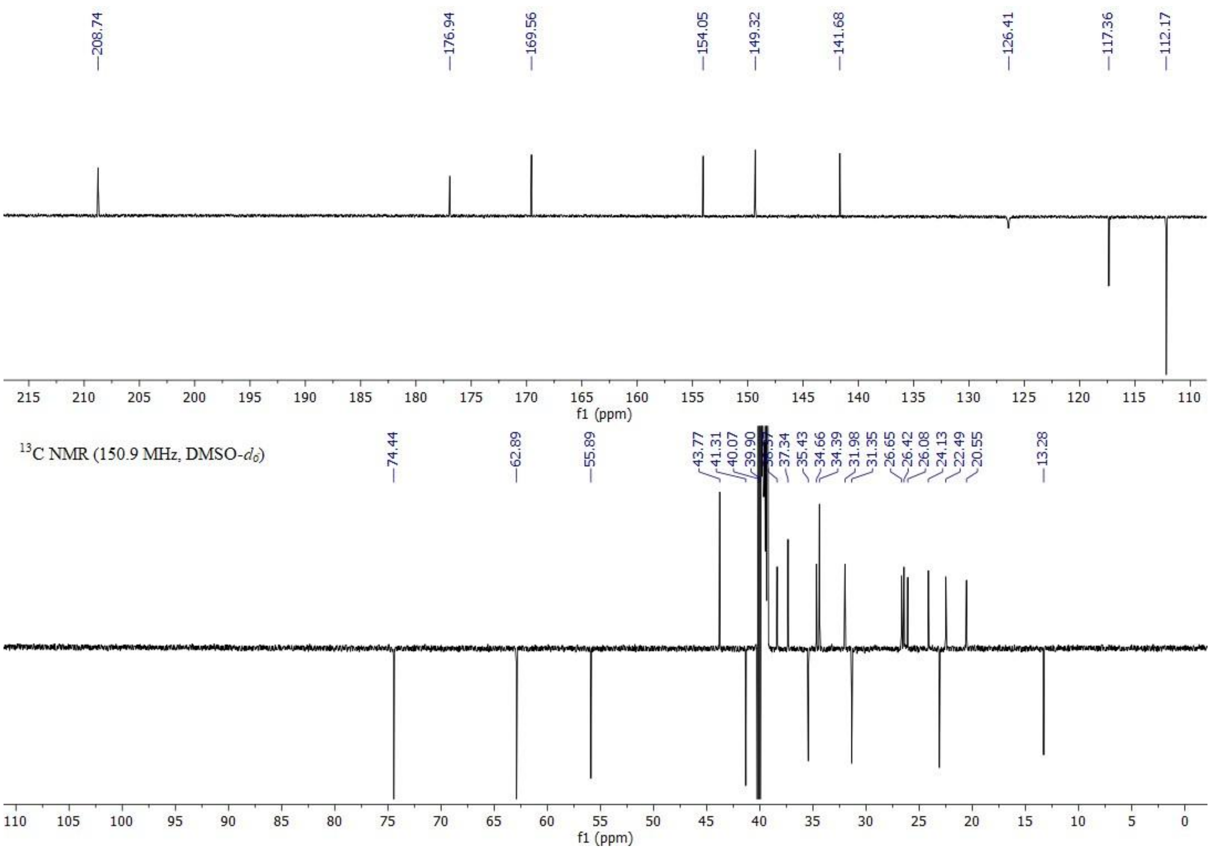

208.74
176.94
169.56
154.05
149.32
141.68
126.41
117.36
112.17
f1 (ppm)
13C NMR (150.9 MHz, DMSO-d6)
74.44
62.89
55.89
43.77
41.31
40.07
39.90
37.34
35.43
34.66
34.39
31.98
31.35
26.65
26.42
26.08
24.13
22.49
20.55
13.28
f1 (ppm)

**42.** **11**.

**Figure S** $^{1}$H NMR (top) and $^{13}$C APT NMR (bottom) spectra of compound

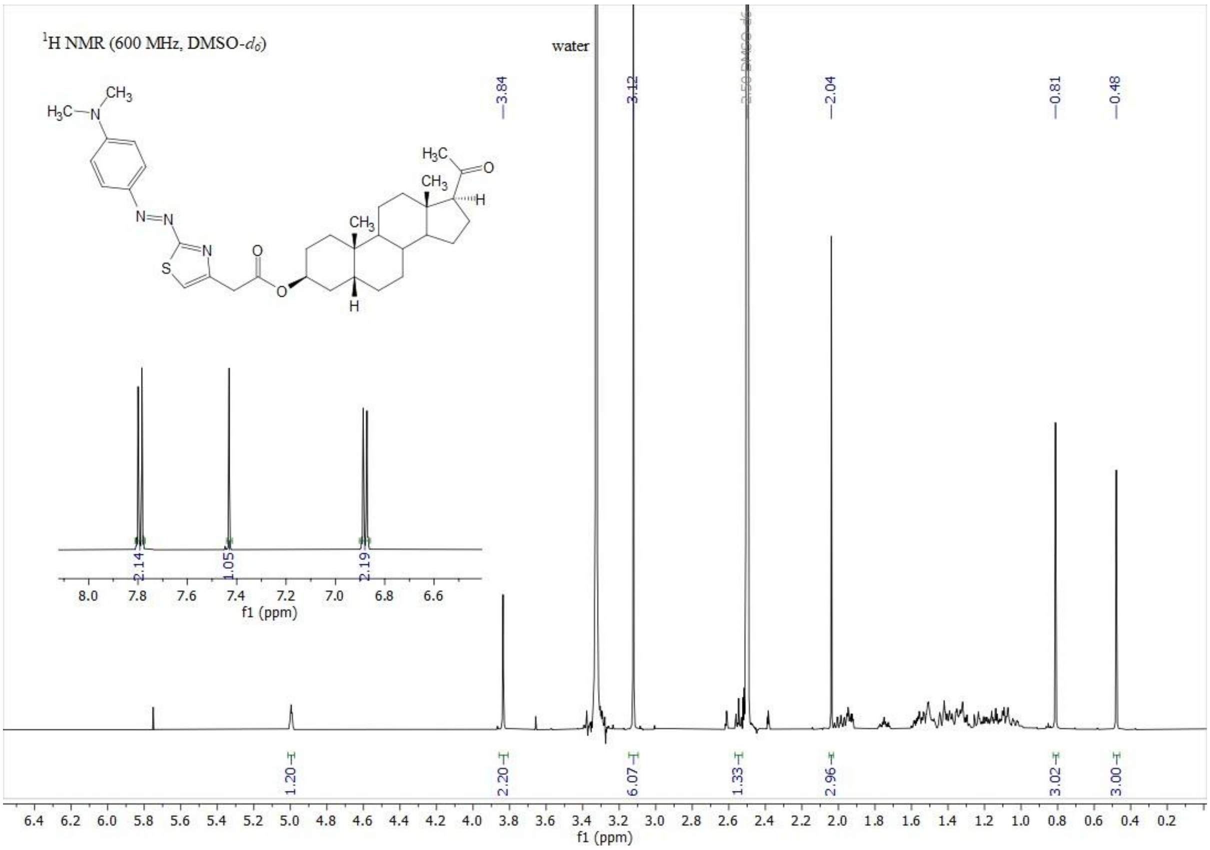

1H NMR (600 MHz, DMSO-d6)
water
3.84
3.12
3.50 DMSO-d6
2.04
0.81
0.48
2.14
1.05
2.19
1.20
2.20
6.07
1.33
2.96
3.02
3.00
f1 (ppm)

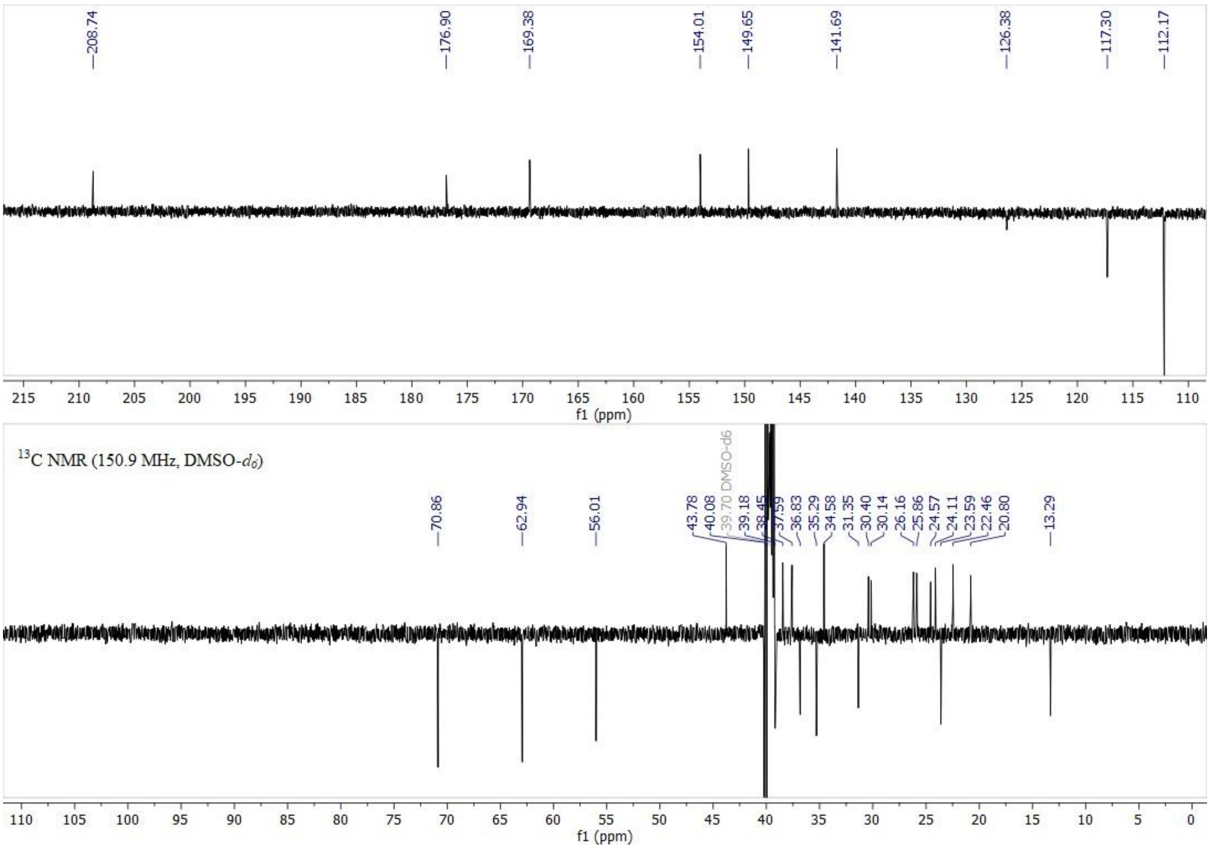

208.74
176.90
169.38
154.01
149.65
141.69
126.38
117.30
112.17
13C NMR (150.9 MHz, DMSO-d6)
70.86
62.94
56.01
43.78
40.08
39.70 DMSO-d6
39.18
38.45
37.59
36.83
35.29
34.58
31.35
30.40
30.14
26.16
25.86
24.57
24.11
23.59
22.46
20.80
13.29
f1 (ppm)

**43.** **12**.

**Figure S** $^{1}H$ NMR (top) and $^{13}C$ APT NMR (bottom) spectra of compound

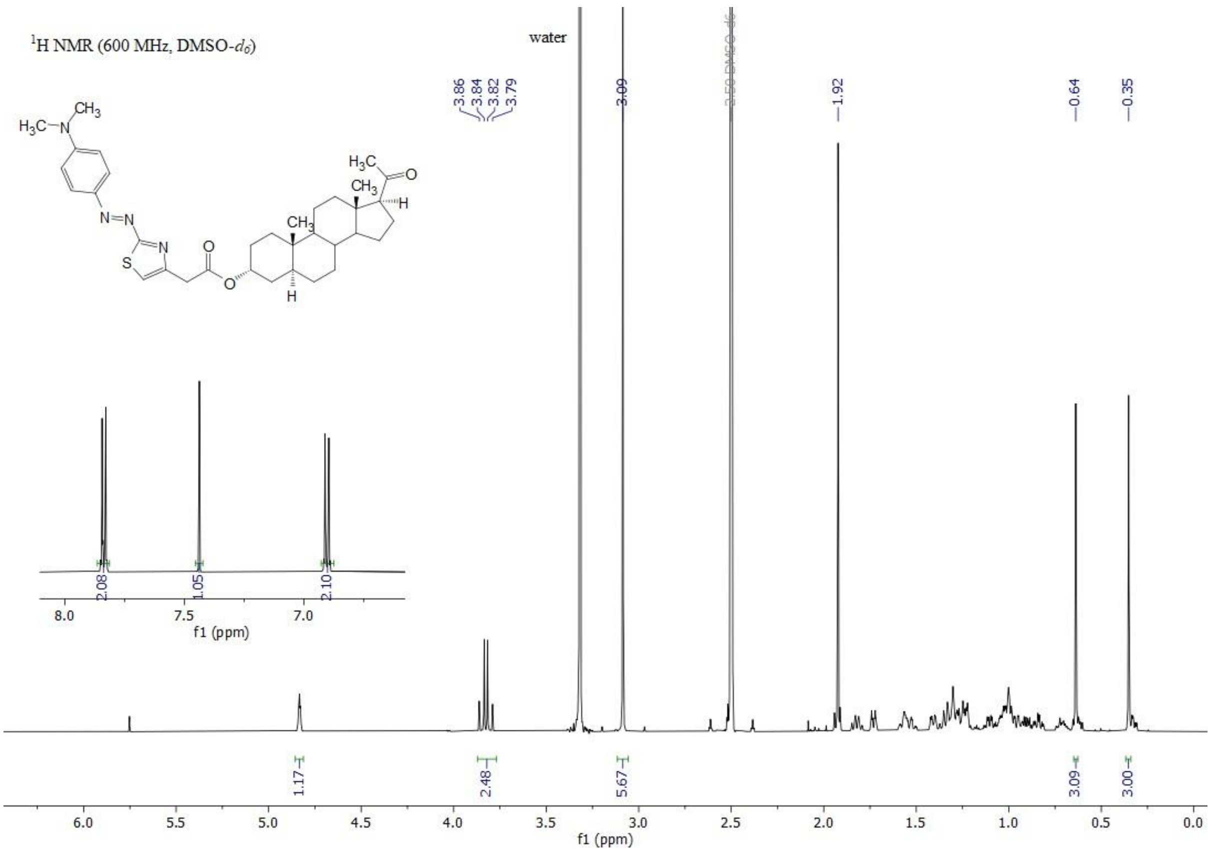

¹H NMR (600 MHz, DMSO-d6)
water
3.86 3.84 3.82 3.79
3.09
1.92
0.64
0.35
2.08
1.05
2.10
1.17
2.48
5.67
3.09
3.00
f1 (ppm)

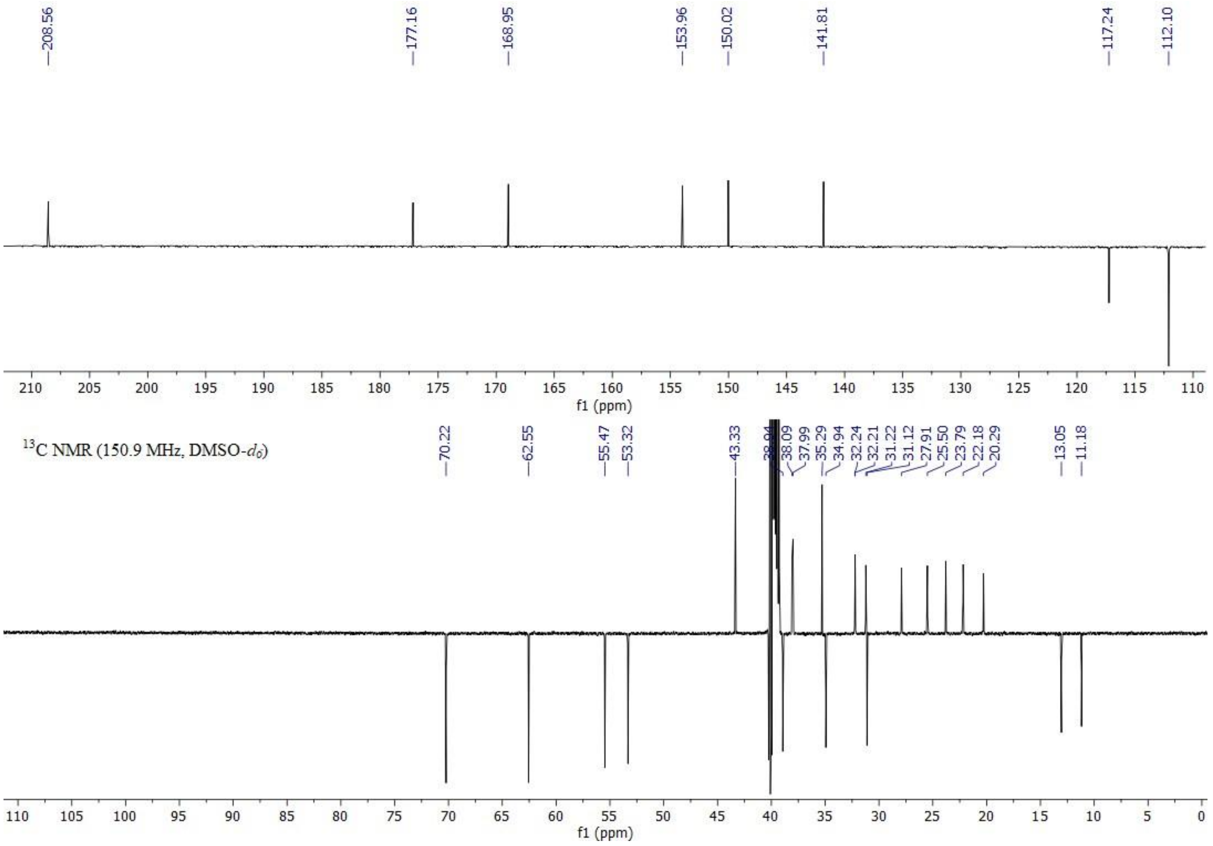

208.56
177.16
168.95
153.96
150.02
141.81
117.24
112.10
f1 (ppm)
¹³C NMR (150.9 MHz, DMSO-d6)
70.22
62.55
55.47
53.32
43.33
38.09
37.99
35.29
34.94
32.24
32.21
31.22
31.12
27.91
25.50
23.79
22.18
20.29
13.05
11.18
f1 (ppm)

**44.** **13**.

**Figure S** $^{1}H$ NMR (top) and $^{13}C$ APT NMR (bottom) spectra of compound

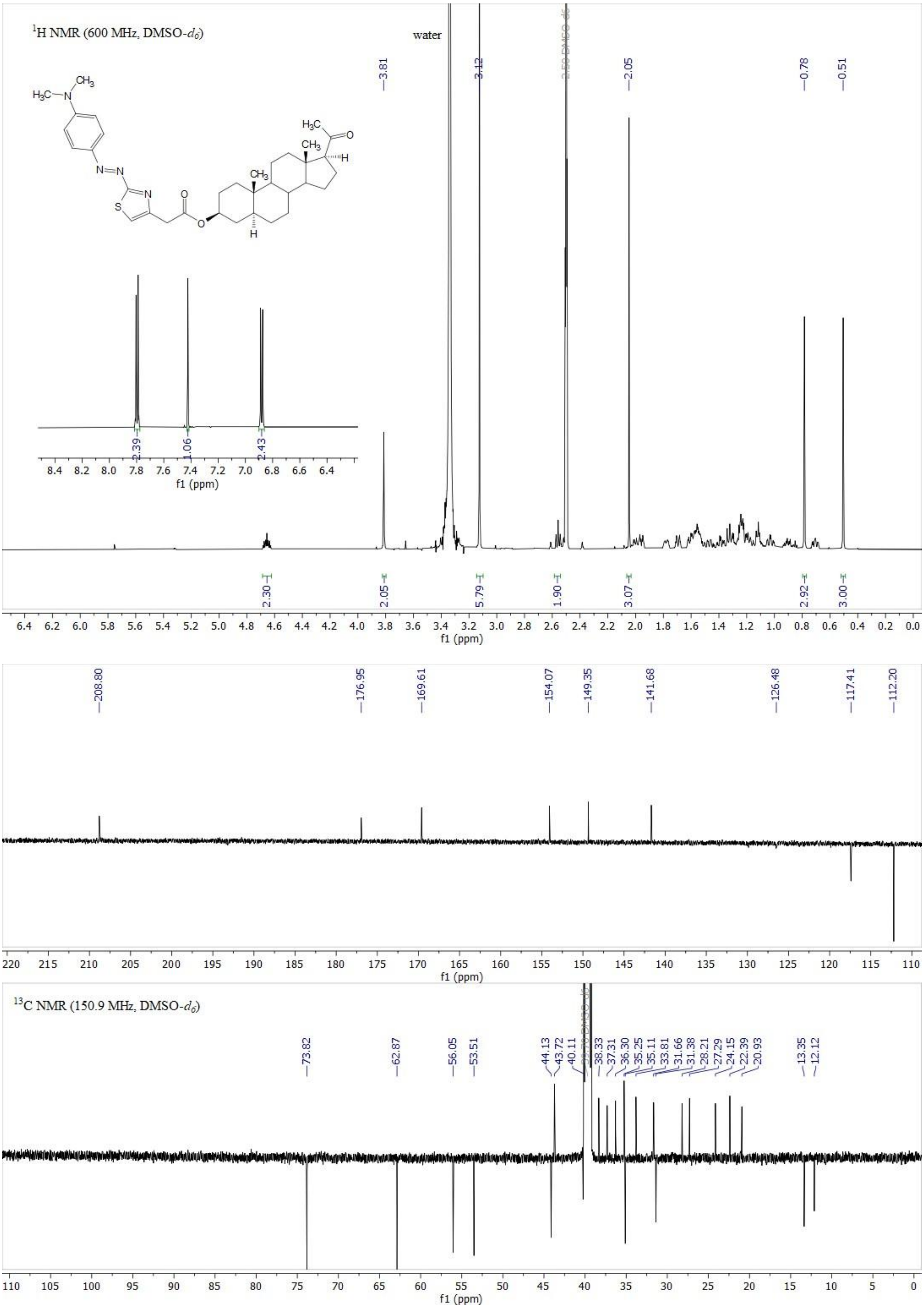

1H NMR (600 MHz, DMSO-d6)
water
3.81
3.12
2.50 DMSO-d6
2.05
0.78
0.51
2.39
1.06
2.43
2.30
2.05
5.79
1.90
3.07
2.92
3.00
f1 (ppm)
208.80
176.95
169.61
154.07
149.35
141.68
126.48
117.41
112.20
13C NMR (150.9 MHz, DMSO-d6)
73.82
62.87
56.05
53.51
44.13
43.72
40.11
38.33
37.31
36.30
35.25
35.11
33.81
31.66
31.38
28.21
27.29
24.15
22.39
20.93
13.35
12.12

**45.** **14**.

**Figure S** [1]H NMR (top) and [13]C APT NMR (bottom) spectra of compound

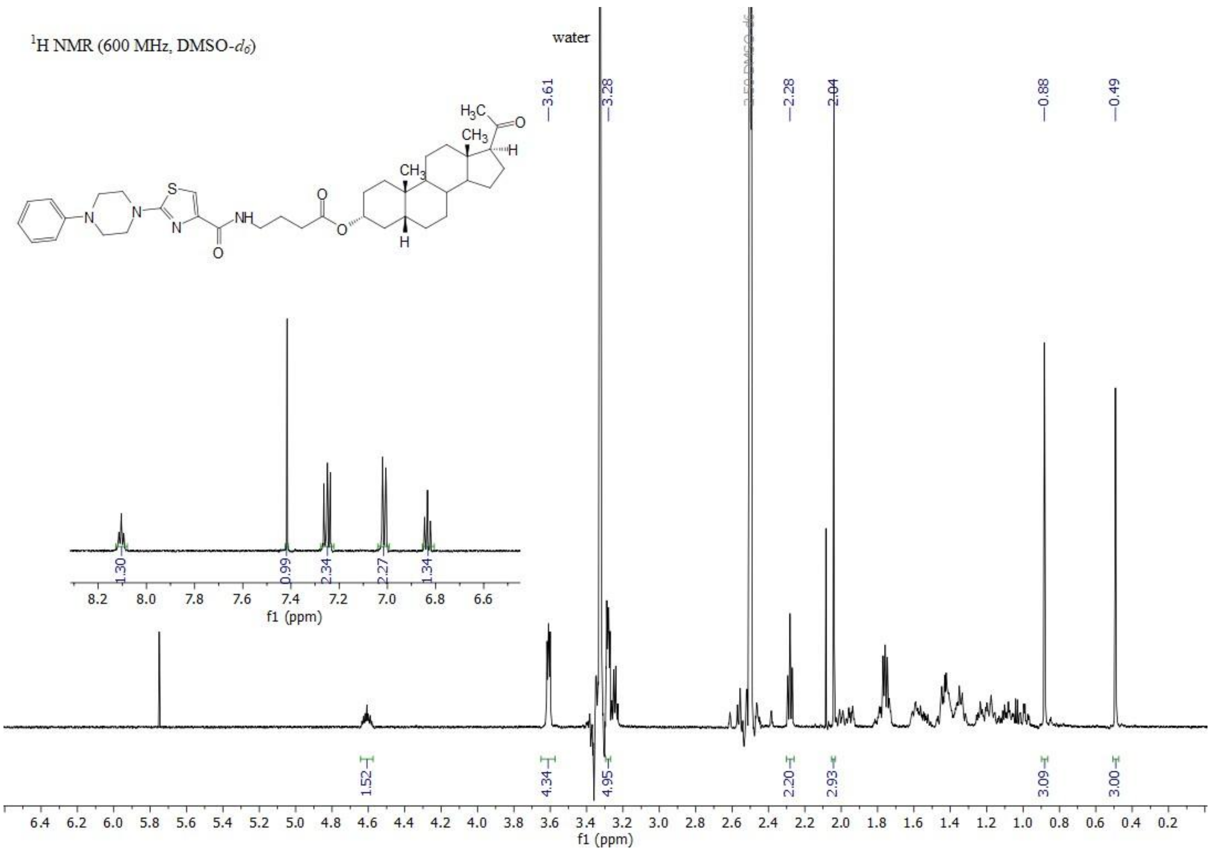

$^1$H NMR (600 MHz, DMSO-$d_6$)
water
3.61
3.28
2.50 DMSO-$d_6$
2.28
2.04
0.88
0.49
1.30
0.99
2.34
2.27
1.34
1.52
4.34
4.95
2.20
2.93
3.09
3.00
f1 (ppm)

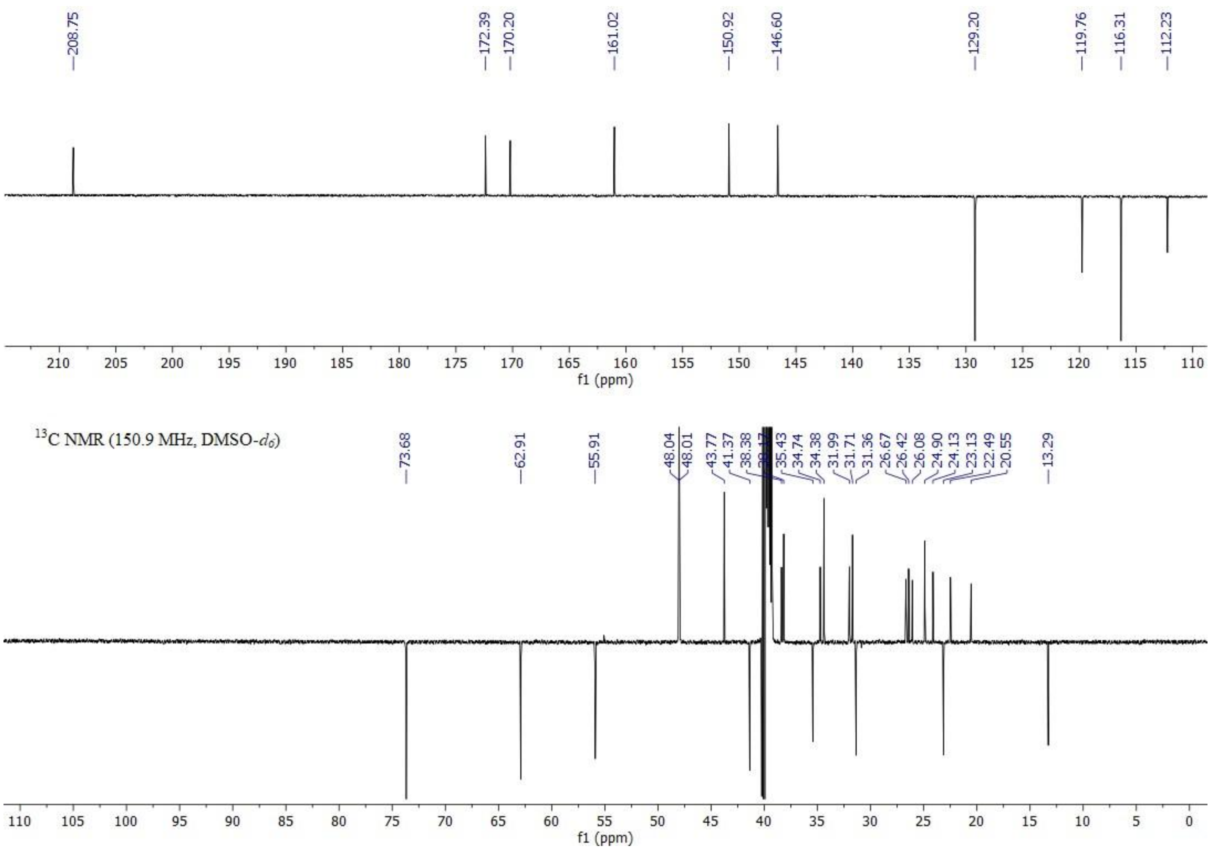

208.75
172.39
170.20
161.02
150.92
146.60
129.20
119.76
116.31
112.23
f1 (ppm)
$^{13}$C NMR (150.9 MHz, DMSO-$d_6$)
73.68
62.91
55.91
48.04
48.01
43.77
41.37
38.38
35.43
34.74
34.38
31.99
31.71
31.36
26.67
26.42
26.08
24.90
24.13
23.13
22.49
20.55
13.29
f1 (ppm)

**46.** **18**.

**Figure S** $^{1}H$ NMR (top) and $^{13}C$ APT NMR (bottom) spectra of compound

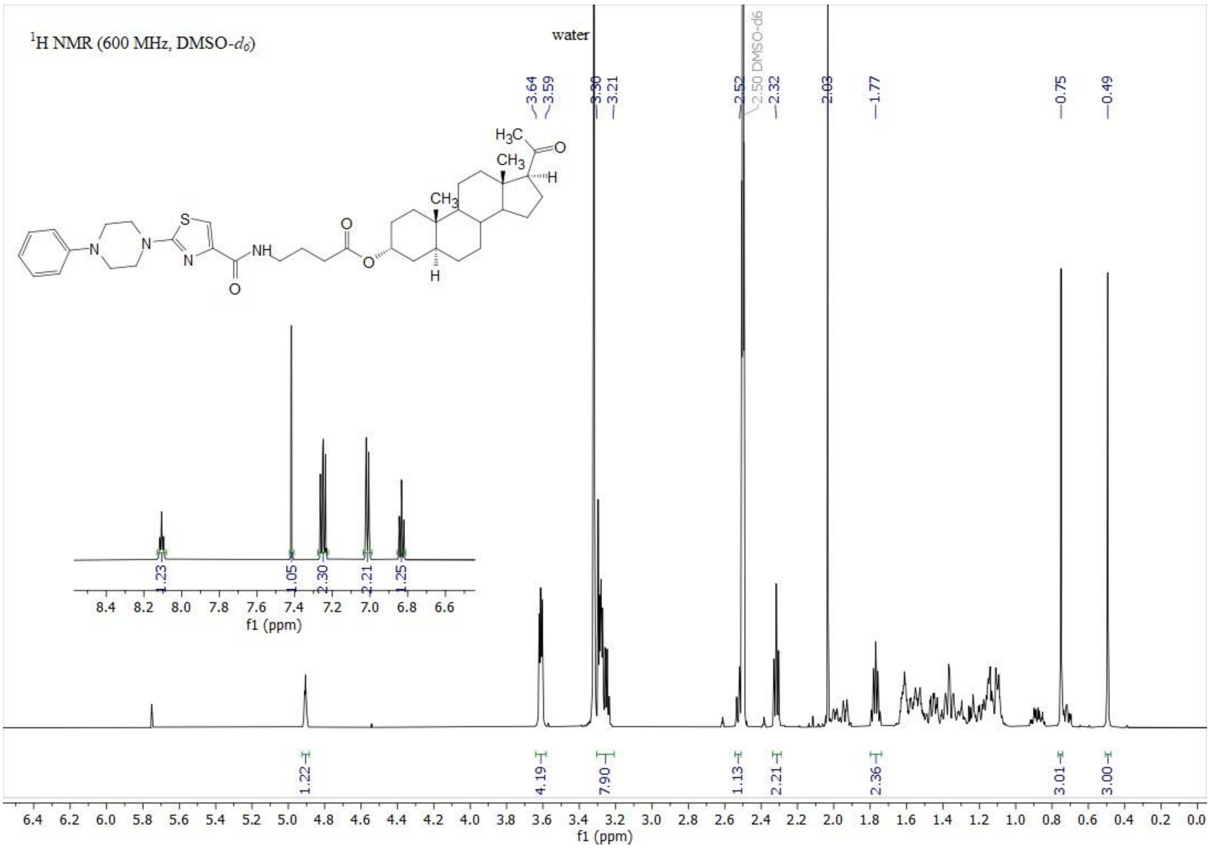

¹H NMR (600 MHz, DMSO-d6)
water
3.64
3.59
3.30
3.21
2.52
2.50 DMSO-d6
2.32
2.03
1.77
0.75
0.49
1.23
1.05
2.30
2.21
1.25
1.22
4.19
7.90
1.13
2.21
2.36
3.01
3.00
f1 (ppm)

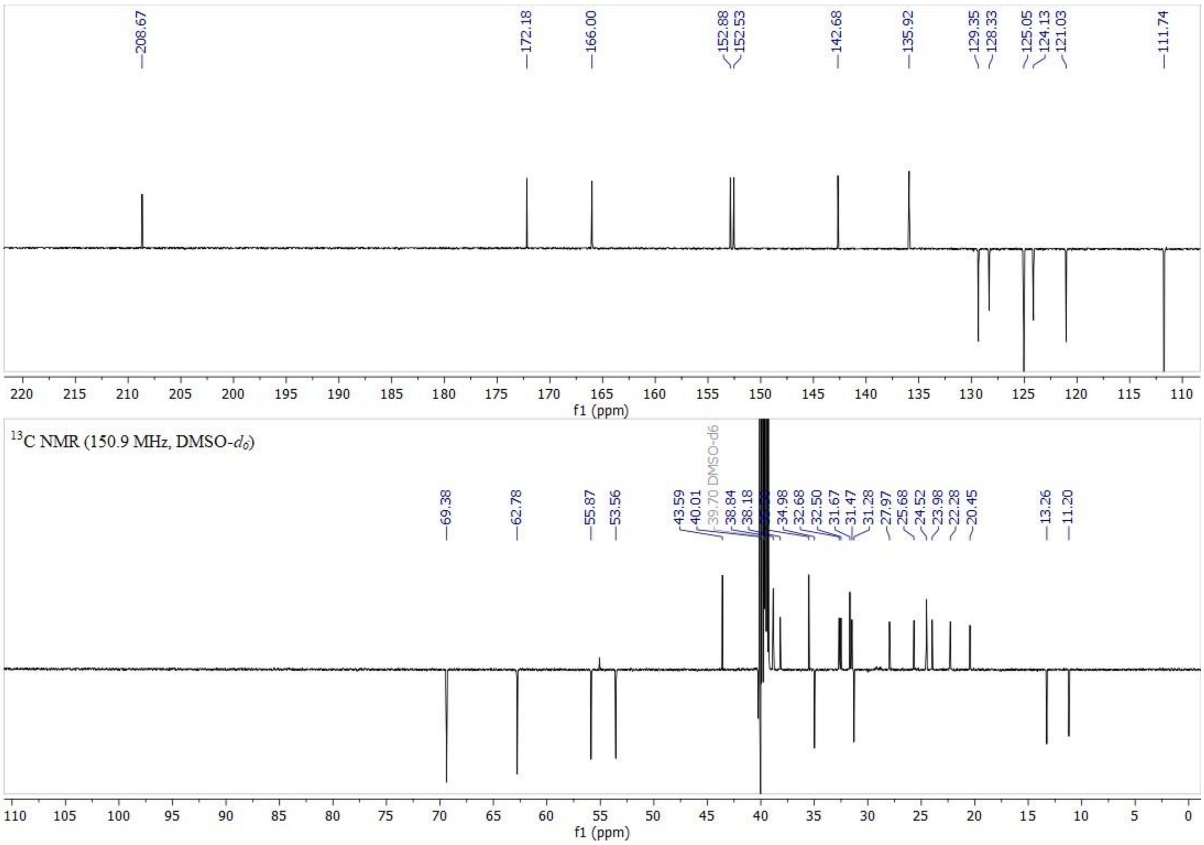

208.67
172.18
166.00
152.88
152.53
142.68
135.92
129.35
128.33
125.05
124.13
121.03
111.74
f1 (ppm)
¹³C NMR (150.9 MHz, DMSO-d6)
69.38
62.78
55.87
53.56
43.59
40.01
39.70 DMSO-d6
38.84
38.18
34.98
32.68
32.50
31.67
31.47
31.28
27.97
25.68
24.52
23.98
22.28
20.45
13.26
11.20
f1 (ppm)

**47.** **19**.

**Figure S** $^{1}H$ NMR (top) and $^{13}C$ APT NMR (bottom) spectra of compound

#### 2.6 LC-MS analysis of compounds 1-22

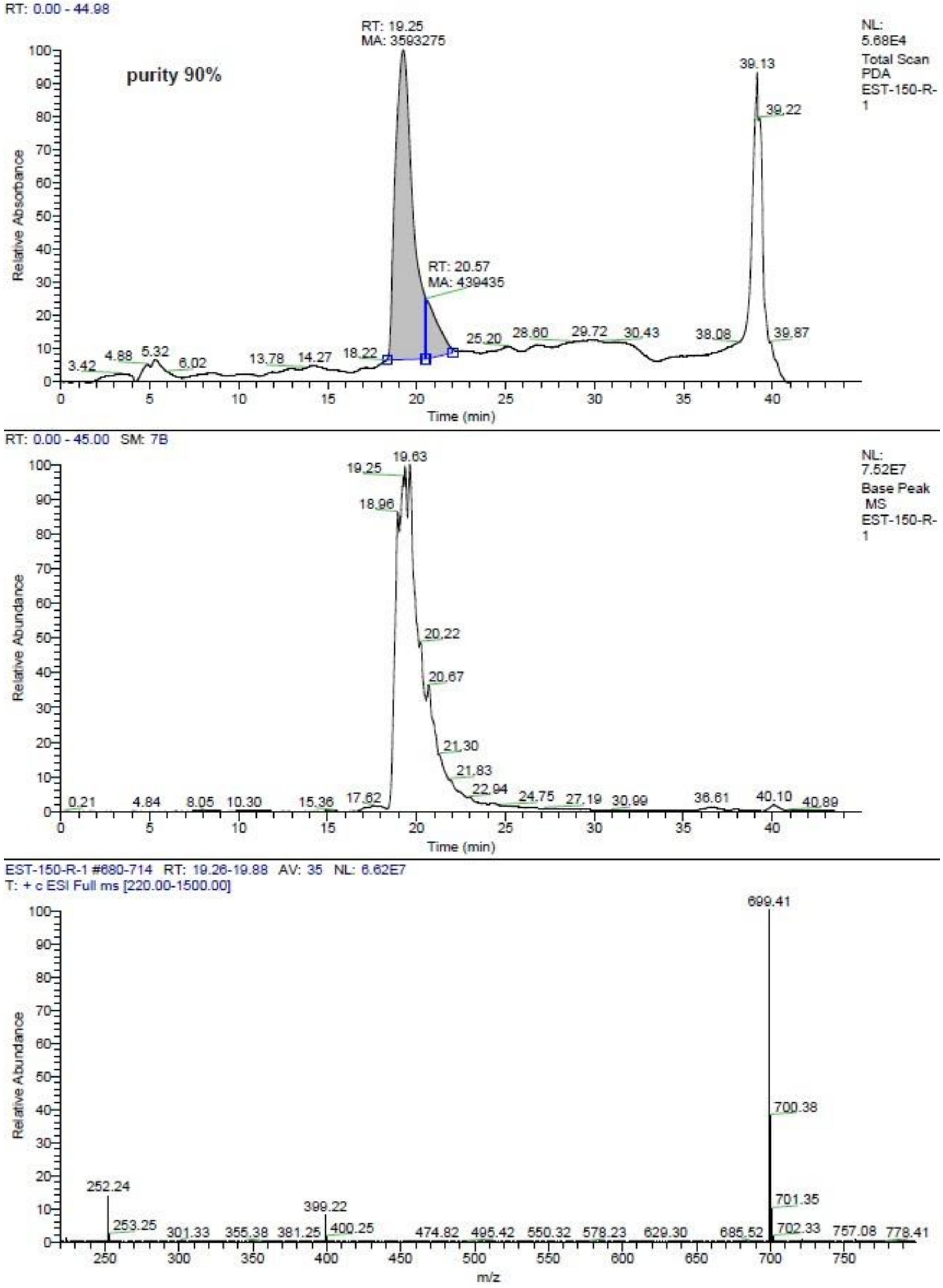

**Figure S52**. LC-MS analysis of compound **1**.

. LC-MS analysis of compound

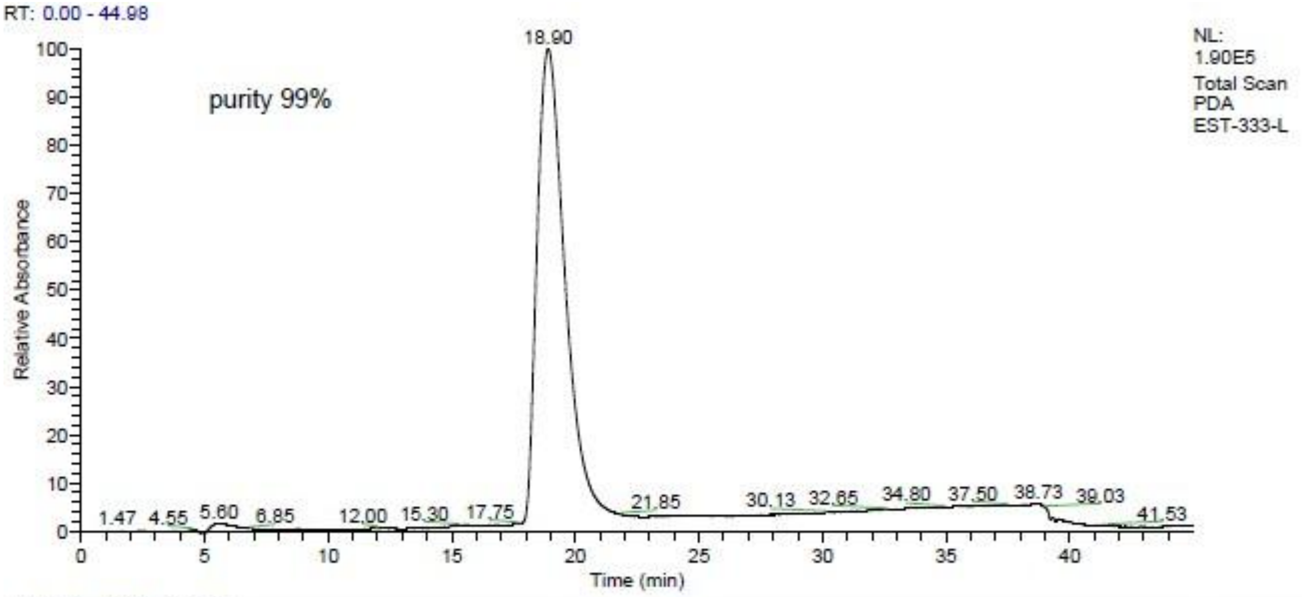
RT: 0.00 - 44.98
purity 99%
NL: 1.90E5
Total Scan PDA
EST-333-L
18.90
1.47 4.55 5.60 6.85 12.00 15.30 17.75 21.85 30.13 32.65 34.80 37.50 38.73 39.03 41.53
Relative Absorbance
Time (min)

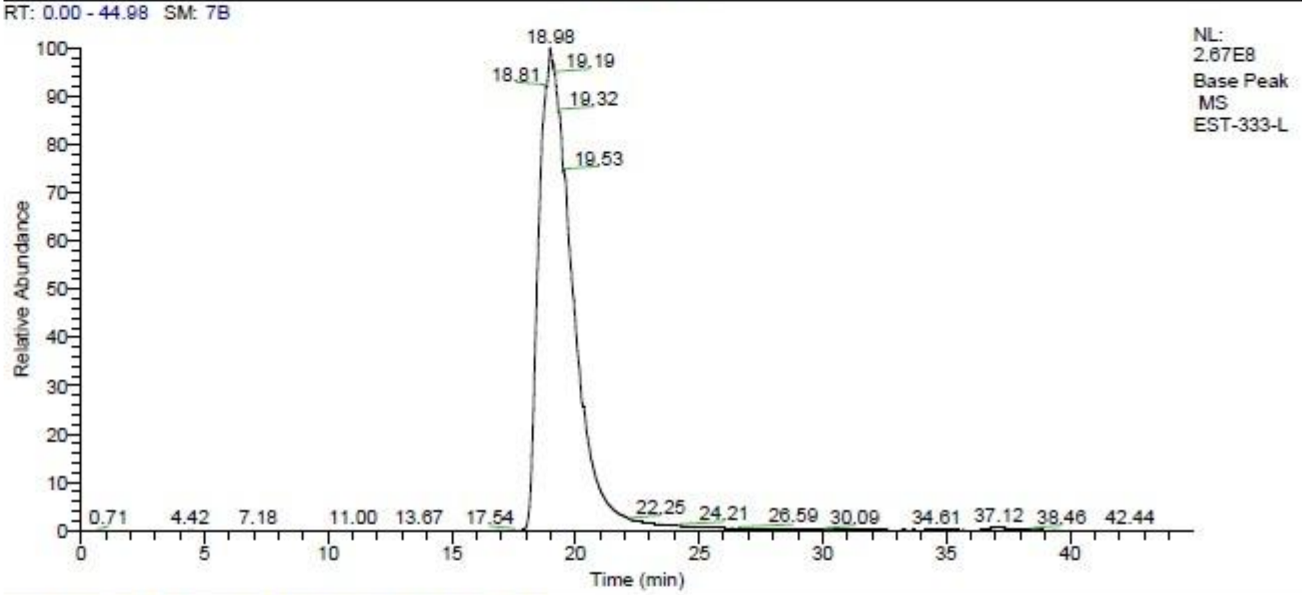
RT: 0.00 - 44.98 SM: 7B
NL: 2.67E8
Base Peak MS
EST-333-L
18.98
18.81 19.19 19.32 19.53
0.71 4.42 7.18 11.00 13.67 17.54 22.25 24.21 26.59 30.09 34.61 37.12 38.46 42.44
Relative Abundance
Time (min)

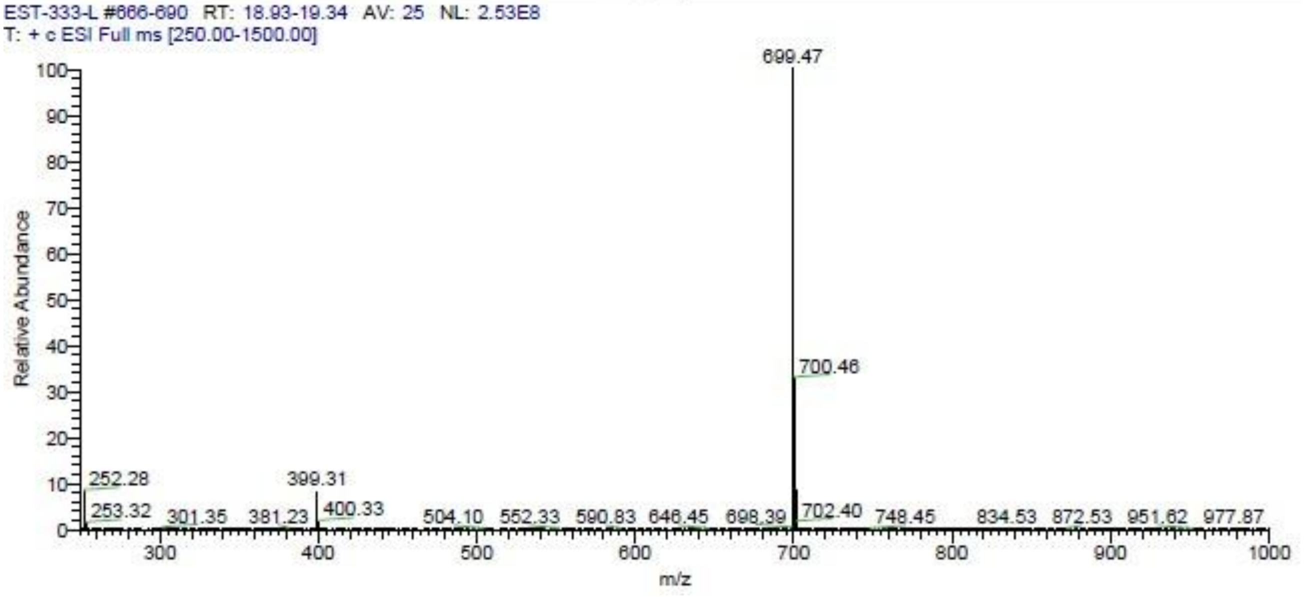
EST-333-L #666-690 RT: 18.93-19.34 AV: 25 NL: 2.53E8
T: + c ESI Full ms [250.00-1500.00]
699.47
700.46
252.28 399.31
253.32 301.35 381.23 400.33 504.10 552.33 590.83 646.45 698.39 702.40 748.45 834.53 872.53 951.62 977.87
Relative Abundance
m/z

**53**. LC-MS analysis of compound **2**.

. LC-MS analysis of compound

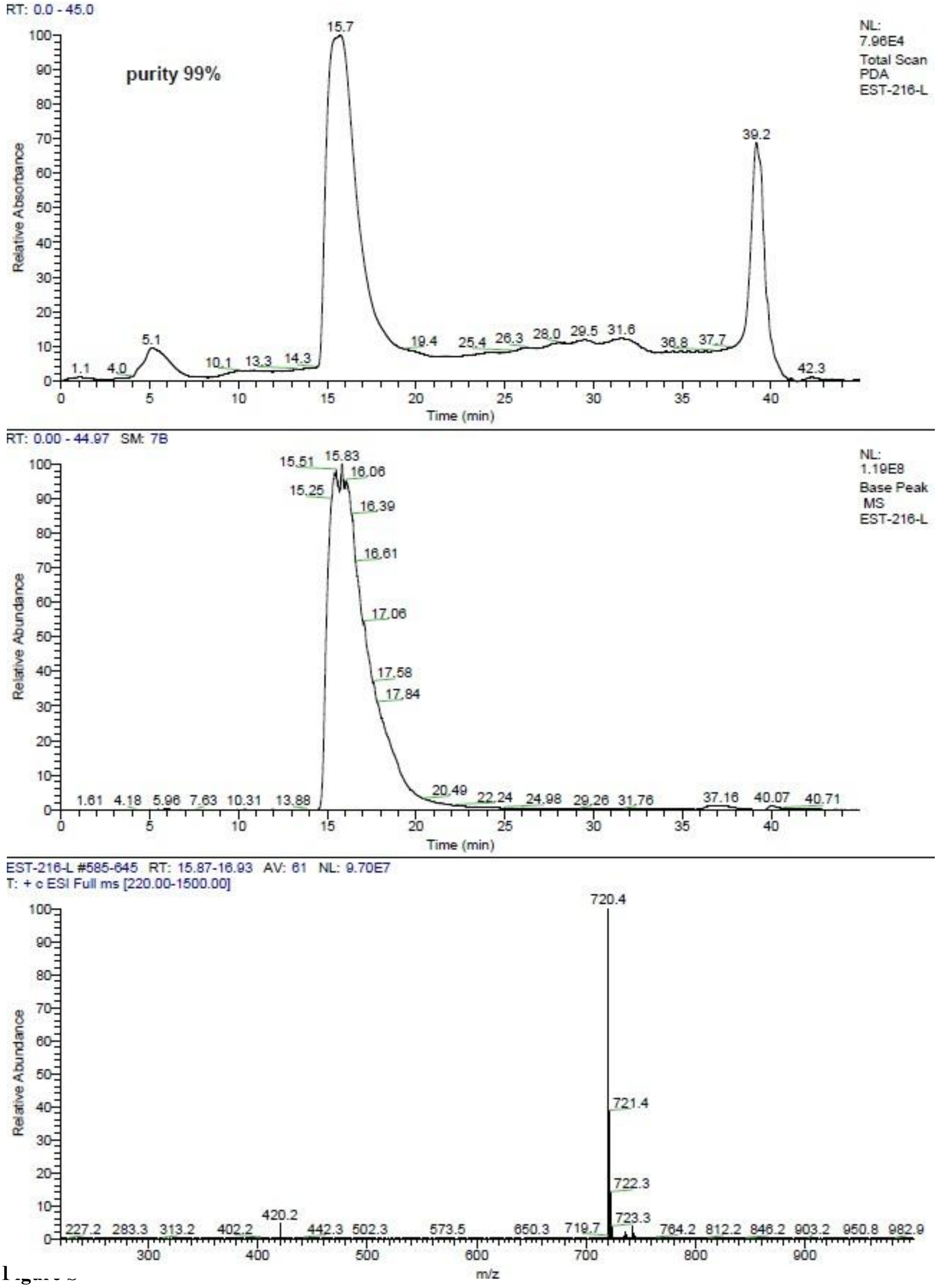
RT: 0.0 - 45.0
purity 99%
NL: 7.96E4
Total Scan
PDA
EST-216-L
Relative Absorbance
Time (min)
15.7
39.2
5.1
1.1 4.0 10.1 13.3 14.3 19.4 25.4 26.3 28.0 29.5 31.6 36.8 37.7 42.3
RT: 0.00 - 44.97 SM: 7B
NL: 1.19E8
Base Peak
MS
EST-216-L
Relative Abundance
Time (min)
15.51 15.83 16.06 15.25 16.39 16.61 17.06 17.58 17.84
1.61 4.18 5.96 7.63 10.31 13.88 20.49 22.24 24.98 29.26 31.76 37.16 40.07 40.71
EST-216-L #585-645 RT: 15.87-16.93 AV: 61 NL: 9.70E7
T: + c ESI Full ms [220.00-1500.00]
Relative Abundance
m/z
720.4
721.4
722.3
723.3
420.2
227.2 283.3 313.2 402.2 442.3 502.3 573.5 650.3 719.7 764.2 812.2 846.2 903.2 950.8 982.9

Figure [illegible]

**Figure S54** **3**.

. LC-MS analysis of compound

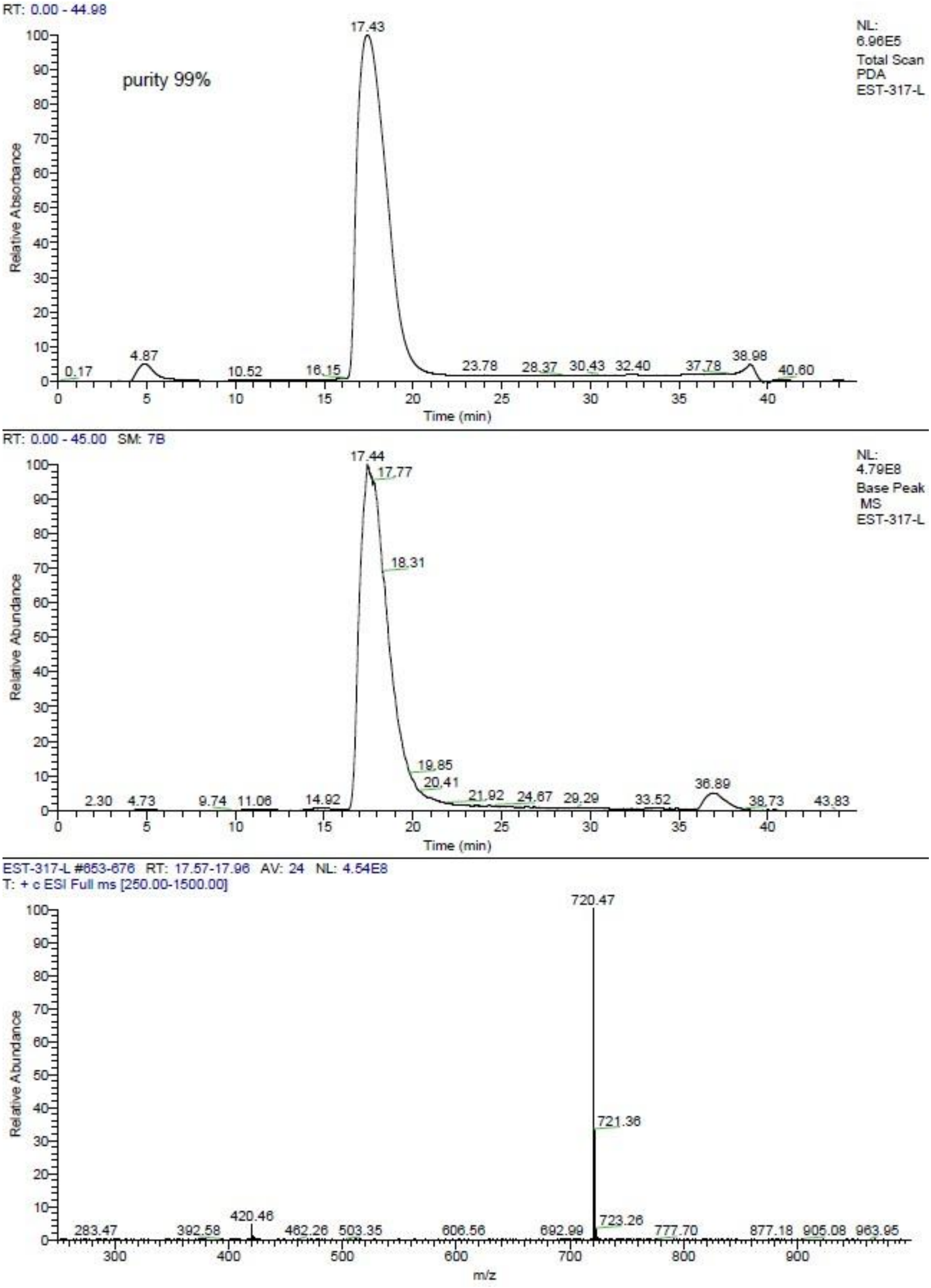
RT: 0.00 - 44.98
purity 99%
17.43
NL:
6.96E5
Total Scan
PDA
EST-317-L
Relative Absorbance
0.17
4.87
10.52
16.15
23.78
28.37
30.43
32.40
37.78
38.98
40.60
Time (min)
RT: 0.00 - 45.00 SM: 7B
17.44
17.77
18.31
19.85
20.41
NL:
4.79E8
Base Peak
MS
EST-317-L
Relative Abundance
2.30
4.73
9.74
11.06
14.92
21.92
24.67
29.29
33.52
36.89
38.73
43.83
Time (min)
EST-317-L #653-676 RT: 17.57-17.96 AV: 24 NL: 4.54E8
T: + c ESI Full ms [250.00-1500.00]
720.47
721.36
723.26
420.46
283.47
392.58
462.26
503.35
606.56
692.99
777.70
877.18
905.08
963.95
Relative Abundance
m/z

**55**. LC-MS analysis of compound **4**.

. LC-MS analysis of compound

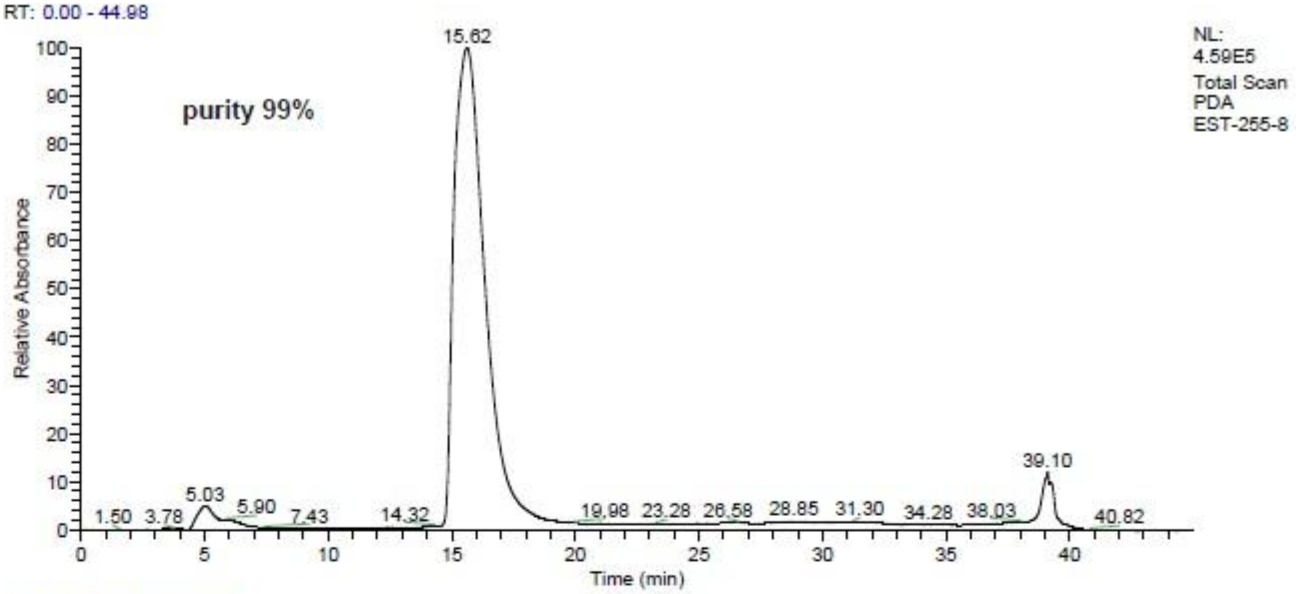
RT: 0.00 - 44.98
purity 99%
NL: 4.59E5
Total Scan
PDA
EST-255-8
Relative Absorbance
Time (min)
15.62
39.10

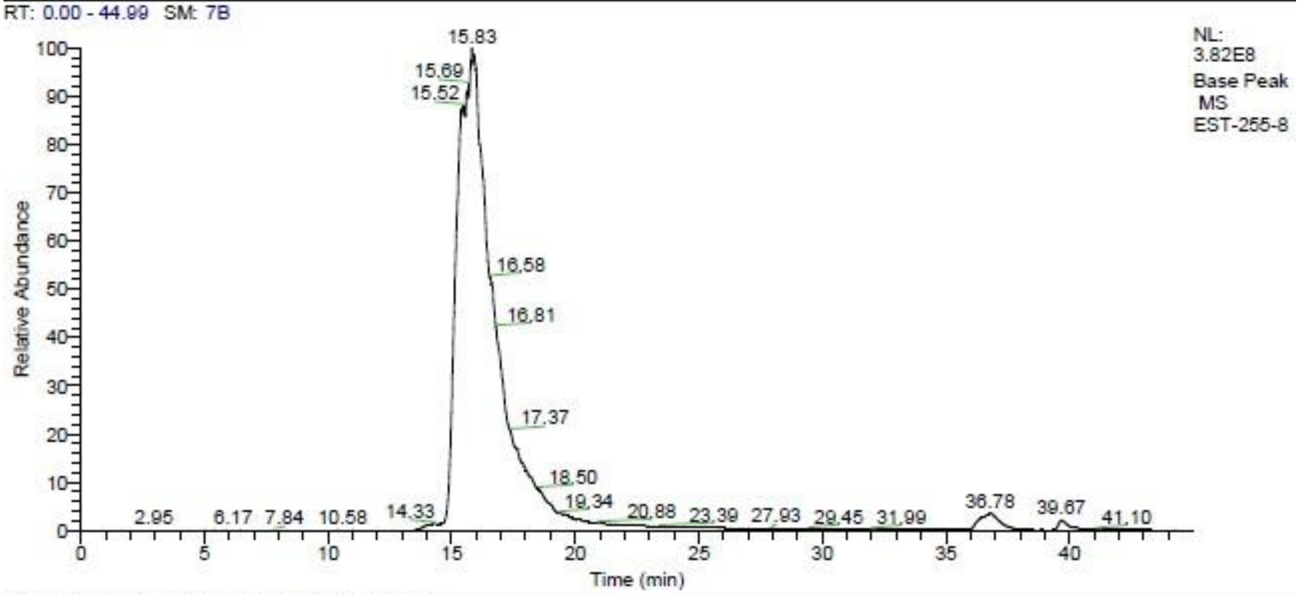
RT: 0.00 - 44.99 SM: 7B
NL: 3.82E8
Base Peak
MS
EST-255-8
Relative Abundance
Time (min)
15.83

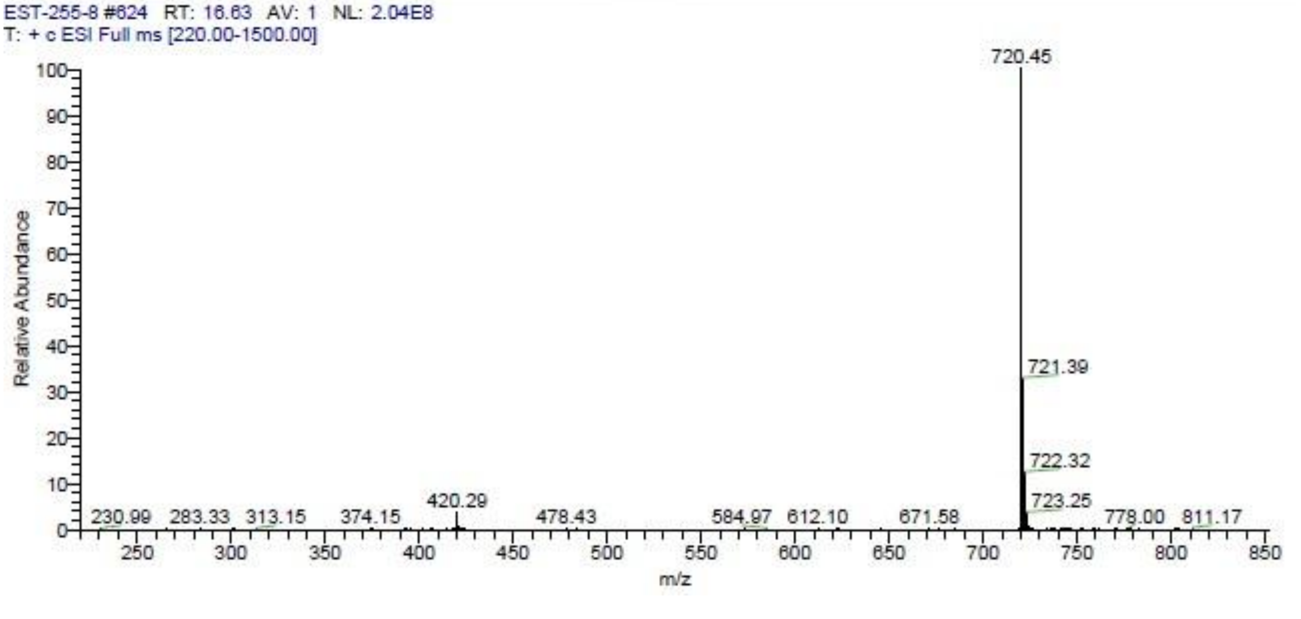
EST-255-8 #624 RT: 16.63 AV: 1 NL: 2.04E8
T: + c ESI Full ms [220.00-1500.00]
Relative Abundance
m/z
720.45
721.39
722.32
723.25
420.29

**56** **5**.

**Figure S** . LC-MS analysis of compound

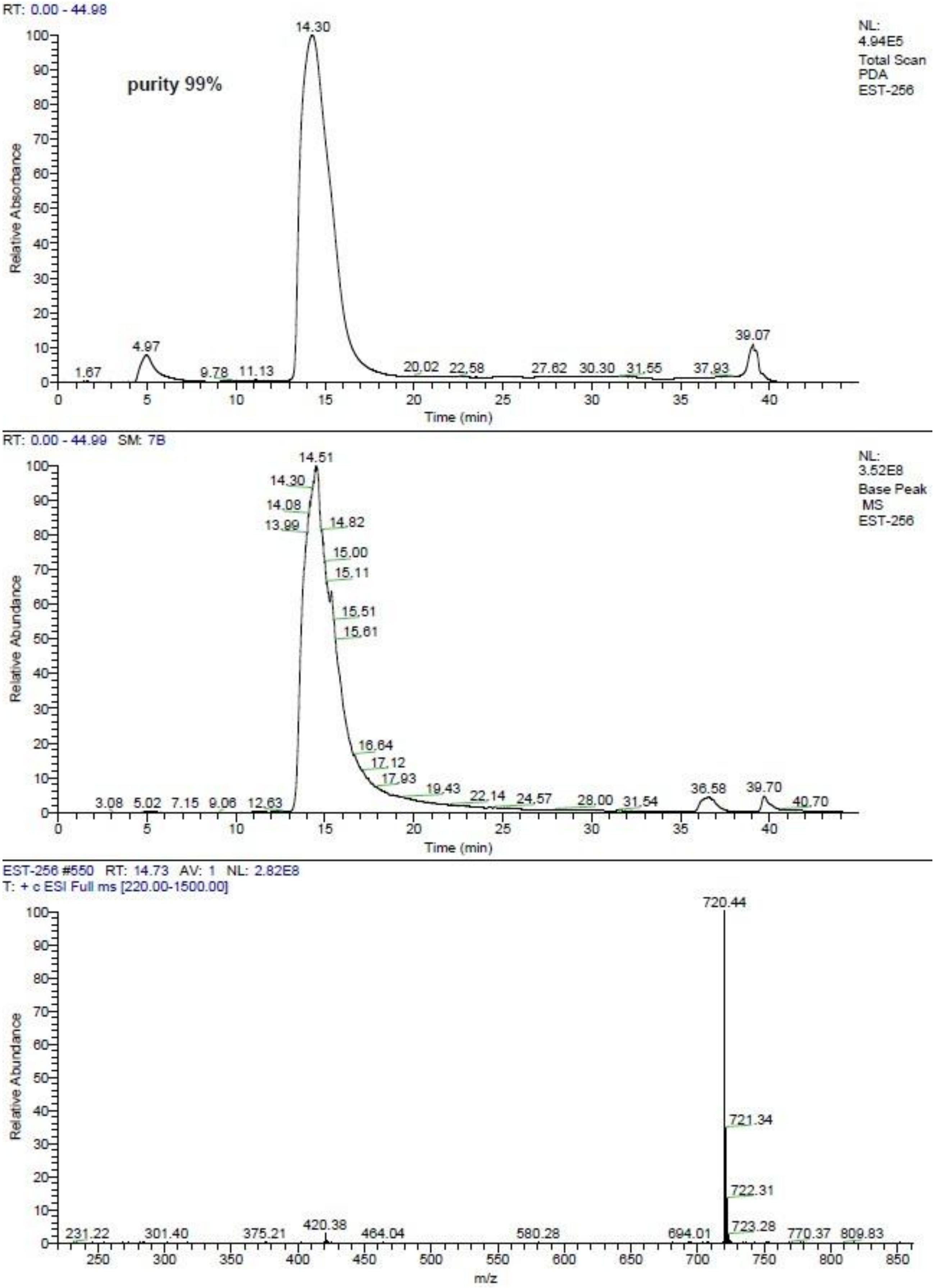
RT: 0.00 - 44.98
purity 99%
NL:
4.94E5
Total Scan
PDA
EST-256
Relative Absorbance
Time (min)
RT: 0.00 - 44.99 SM: 7B
NL:
3.52E8
Base Peak
MS
EST-256
Relative Abundance
Time (min)
EST-256 #550 RT: 14.73 AV: 1 NL: 2.82E8
T: + c ESI Full ms [220.00-1500.00]
720.44
721.34
722.31
723.28
Relative Abundance
m/z

**57** **6**.

**Figure S** . LC-MS analysis of compound

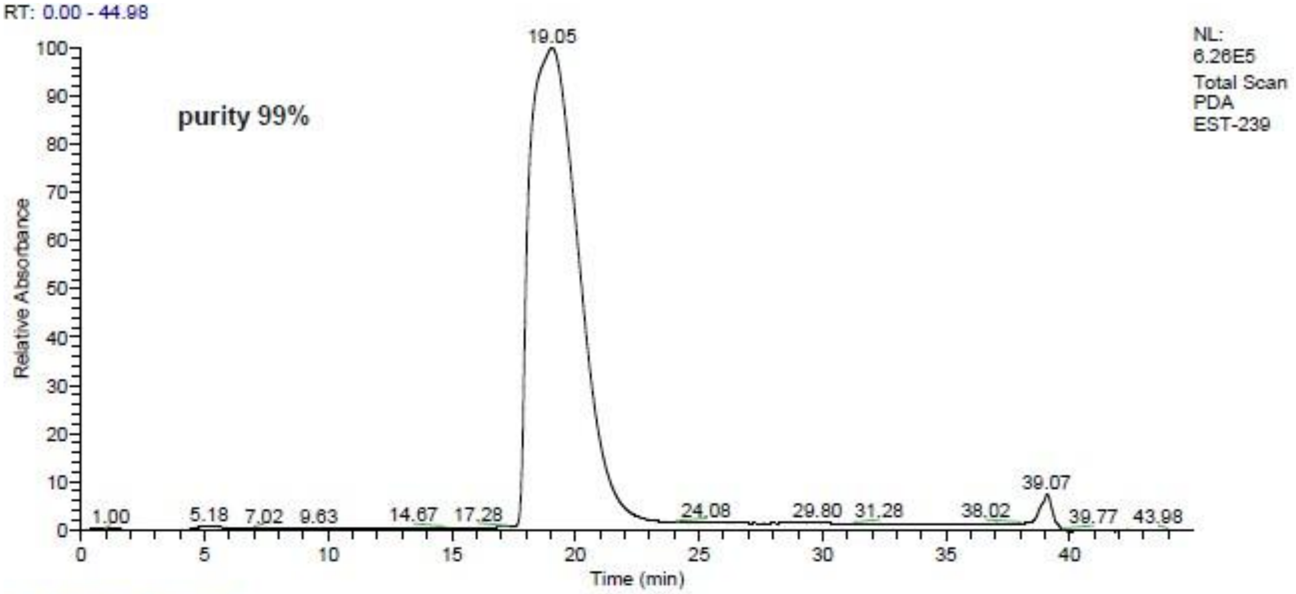
RT: 0.00 - 44.98
purity 99%
NL: 6.26E5 Total Scan PDA EST-239
19.05
1.00
5.18
7.02
9.63
14.67
17.28
24.08
29.80
31.28
38.02
39.07
39.77
43.98
Relative Absorbance
Time (min)

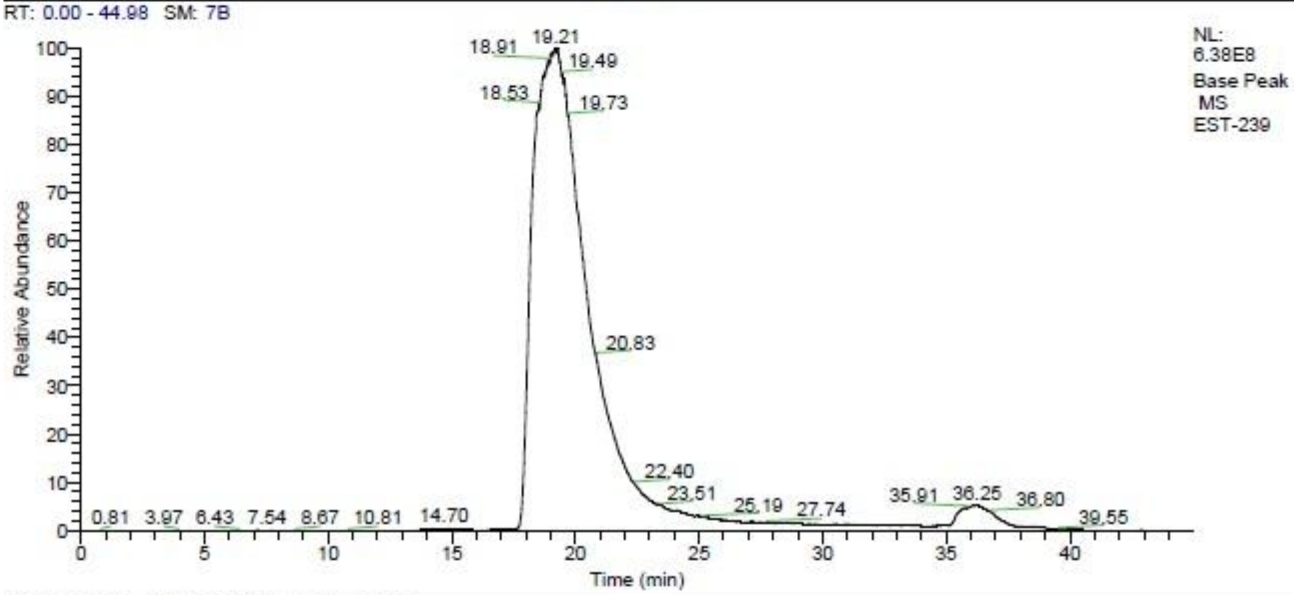
RT: 0.00 - 44.98 SM: 7B
NL: 6.38E8 Base Peak MS EST-239
18.91
19.21
19.49
18.53
19.73
20.83
22.40
23.51
25.19
27.74
35.91
36.25
36.80
39.55
0.81
3.97
6.43
7.54
8.67
10.81
14.70
Relative Abundance
Time (min)

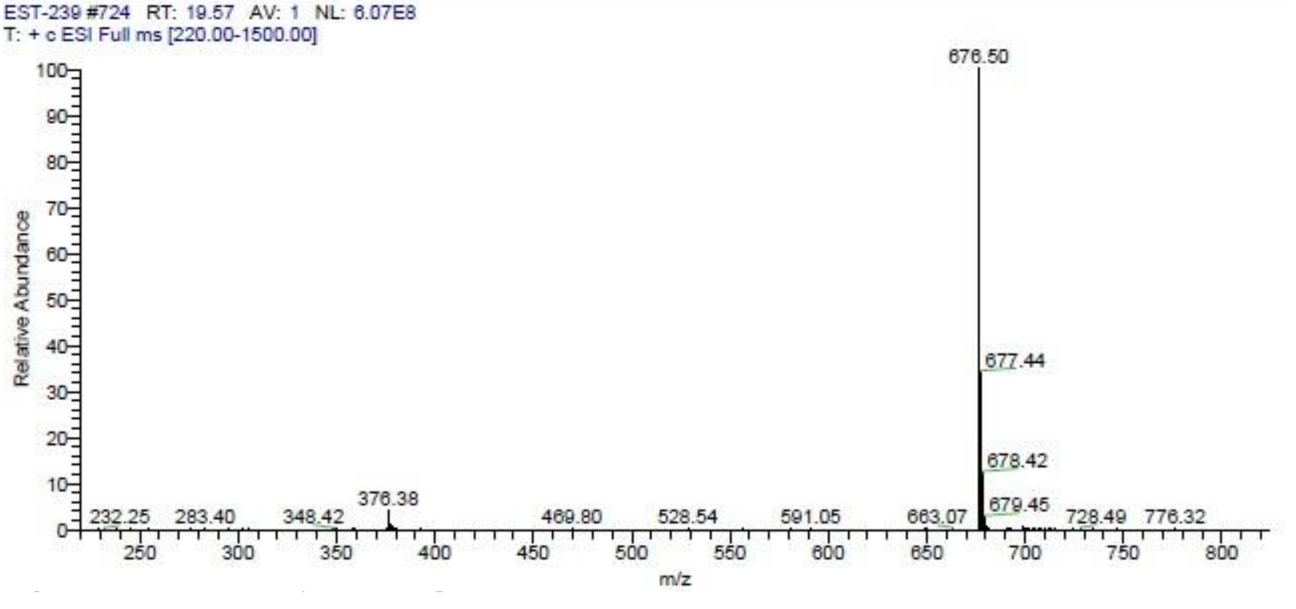
EST-239 #724 RT: 19.57 AV: 1 NL: 6.07E8
T: + c ESI Full ms [220.00-1500.00]
676.50
677.44
678.42
679.45
232.25
283.40
348.42
376.38
469.80
528.54
591.05
663.07
728.49
776.32
Relative Abundance
m/z

**58** **7**.

**Figure S** . LC-MS analysis of compound

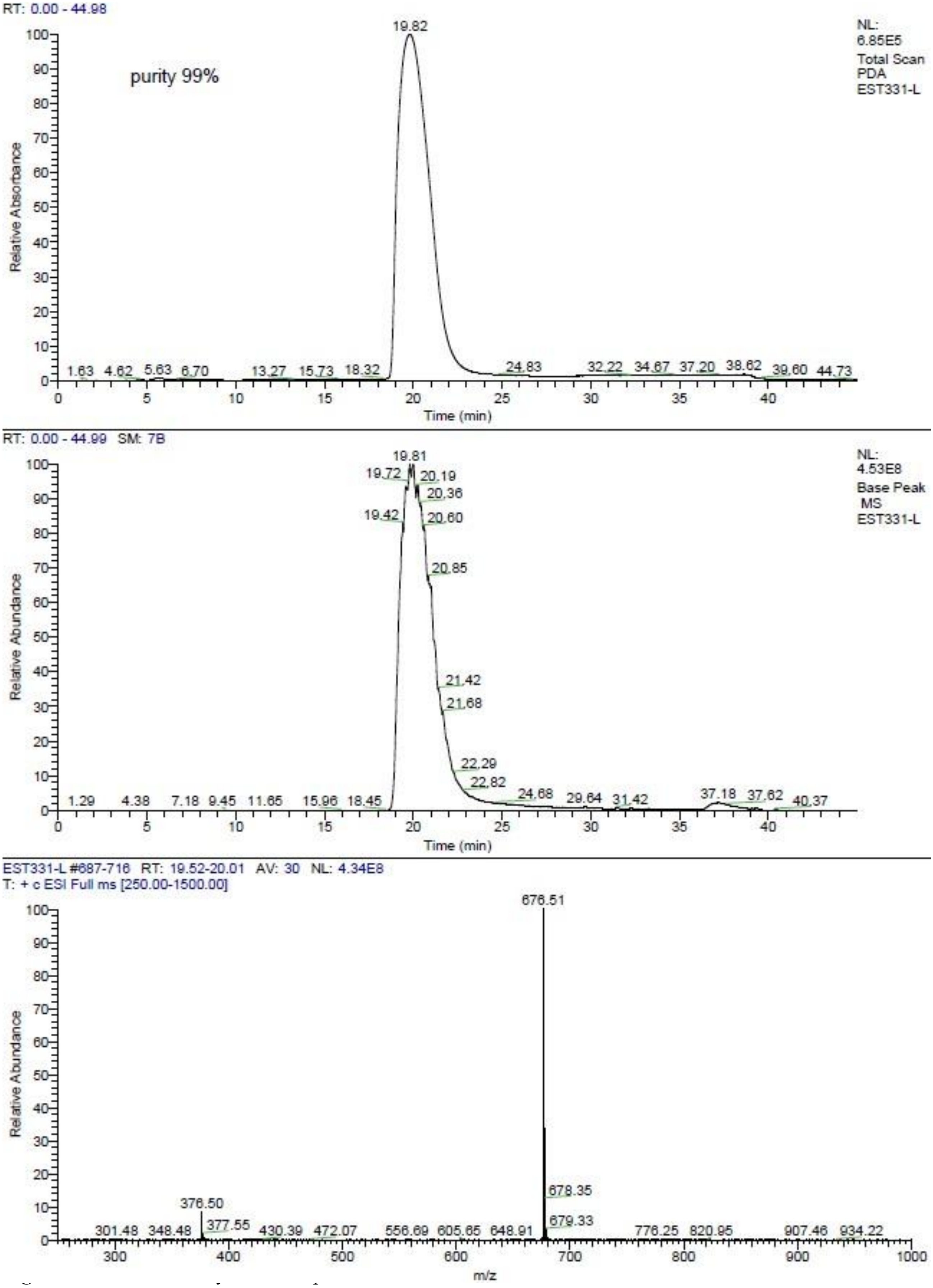

RT: 0.00 - 44.98
purity 99%
NL: 6.85E5
Total Scan PDA
EST331-L
19.82
1.63 4.62 5.63 6.70 13.27 15.73 18.32 24.83 32.22 34.67 37.20 38.62 39.60 44.73
Relative Absorbance
Time (min)
RT: 0.00 - 44.99 SM: 7B
NL: 4.53E8
Base Peak MS
EST331-L
19.81
19.72 20.19 20.36 19.42 20.60 20.85 21.42 21.68 22.29 22.82 24.68 29.64 31.42 37.18 37.62 40.37
1.29 4.38 7.18 9.45 11.65 15.96 18.45
Relative Abundance
Time (min)
EST331-L #687-716 RT: 19.52-20.01 AV: 30 NL: 4.34E8
T: + c ESI Full ms [250.00-1500.00]
676.51
678.35
679.33
376.50
377.55
301.48 348.48 430.39 472.07 556.69 605.65 648.91 776.25 820.95 907.46 934.22
Relative Abundance
m/z

**59** **8**.

**Figure S** . LC-MS analysis of compound

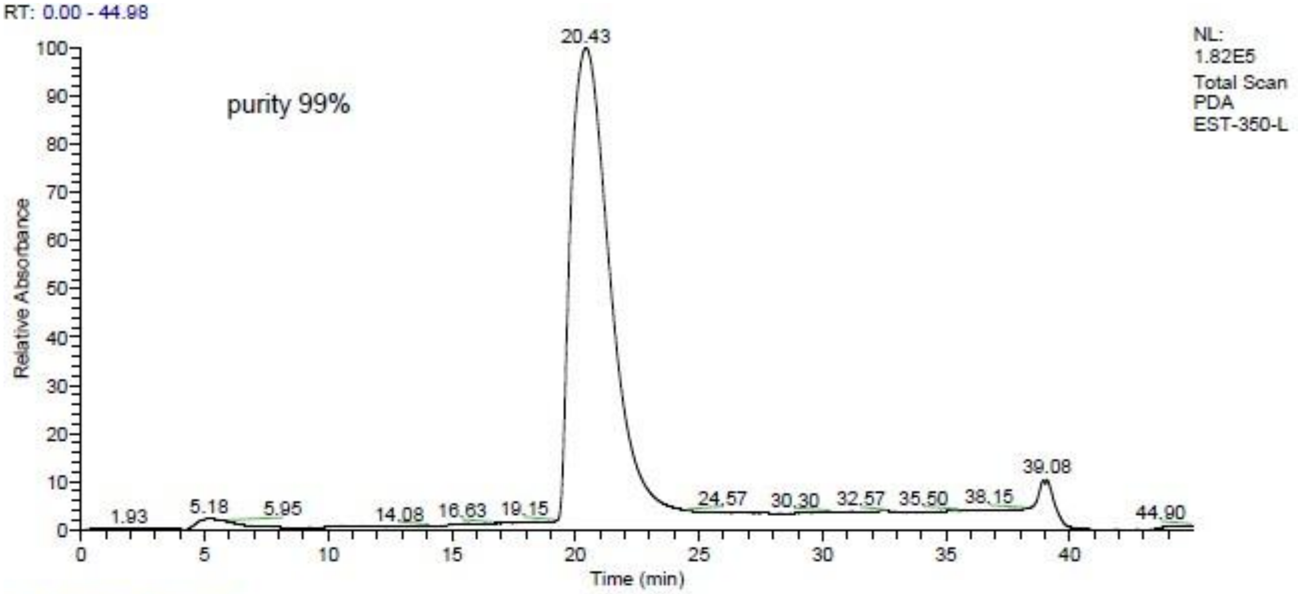
RT: 0.00 - 44.98
NL:
1.82E5
Total Scan
PDA
EST-350-L
purity 99%
Relative Absorbance
Time (min)
20.43
1.93
5.18
5.95
14.08
16.63
19.15
24.57
30.30
32.57
35.50
38.15
39.08
44.90

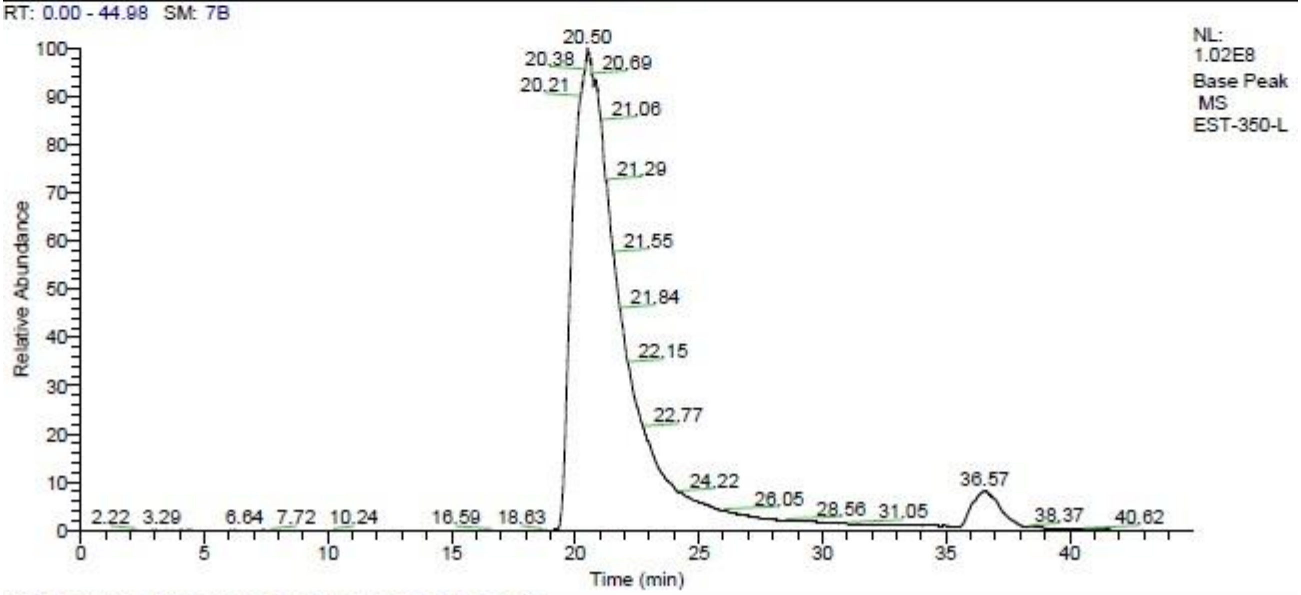
RT: 0.00 - 44.98 SM: 7B
NL:
1.02E8
Base Peak
MS
EST-350-L
Relative Abundance
Time (min)
20.50
20.38
20.69
20.21
21.06
21.29
21.55
21.84
22.15
22.77
24.22
26.05
28.56
31.05
36.57
38.37
40.62
2.22
3.29
6.64
7.72
10.24
16.59
18.63

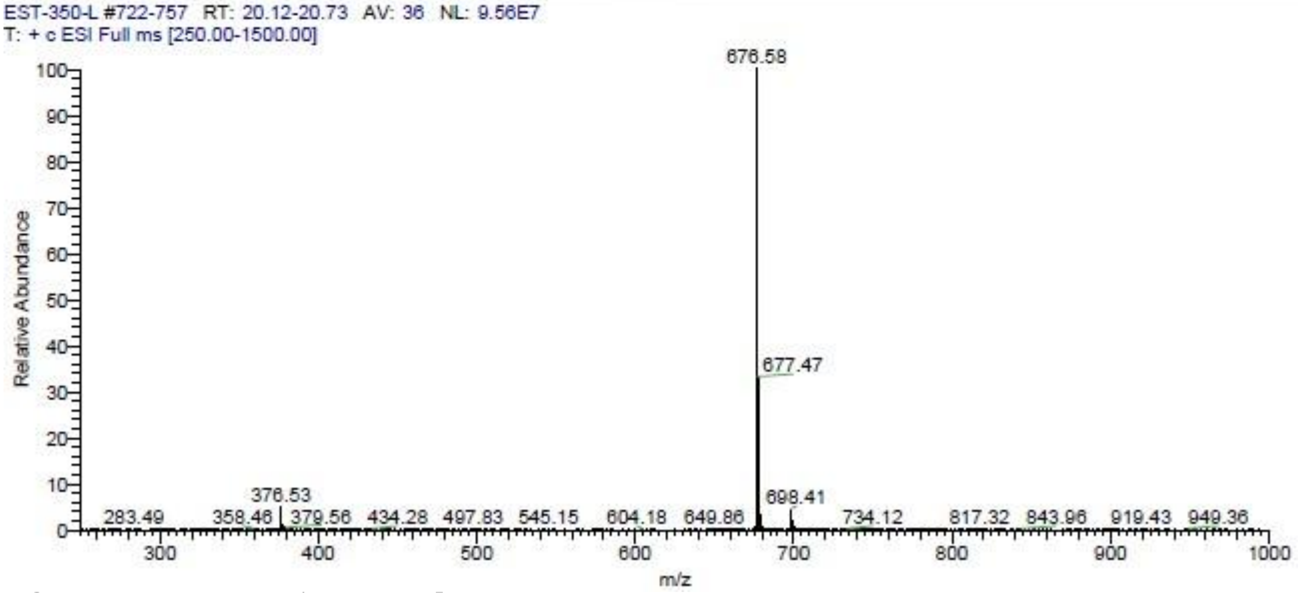
EST-350-L #722-757 RT: 20.12-20.73 AV: 36 NL: 9.56E7
T: + c ESI Full ms [250.00-1500.00]
Relative Abundance
m/z
676.58
677.47
376.53
698.41
283.49
358.46
379.56
434.28
497.83
545.15
604.18
649.86
734.12
817.32
843.96
919.43
949.36

**60** **9**.

**Figure S** . LC-MS analysis of compound

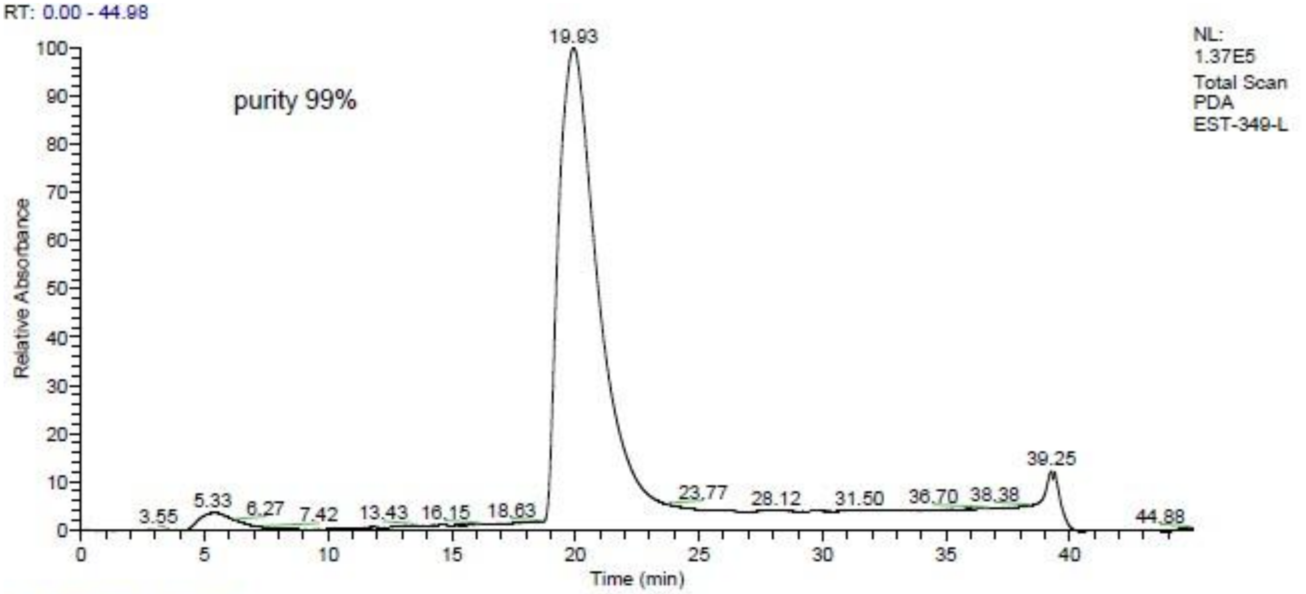
RT: 0.00 - 44.98
NL:
1.37E5
Total Scan
PDA
EST-349-L
purity 99%
Relative Absorbance
Time (min)
19.93
39.25
3.55
5.33
6.27
7.42
13.43
16.15
18.63
23.77
28.12
31.50
36.70
38.38
44.88

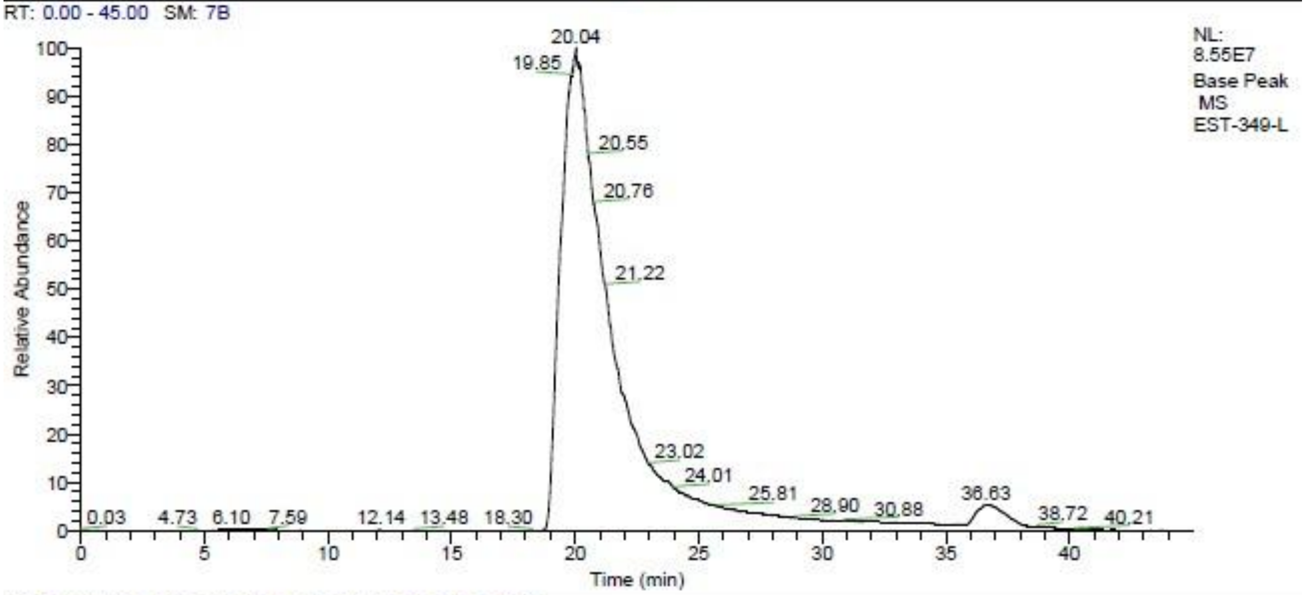
RT: 0.00 - 45.00 SM: 7B
NL:
8.55E7
Base Peak
MS
EST-349-L
Relative Abundance
Time (min)
20.04
19.85
20.55
20.76
21.22
23.02
24.01
25.81
28.90
30.88
36.63
38.72
40.21
0.03
4.73
6.10
7.59
12.14
13.48
18.30

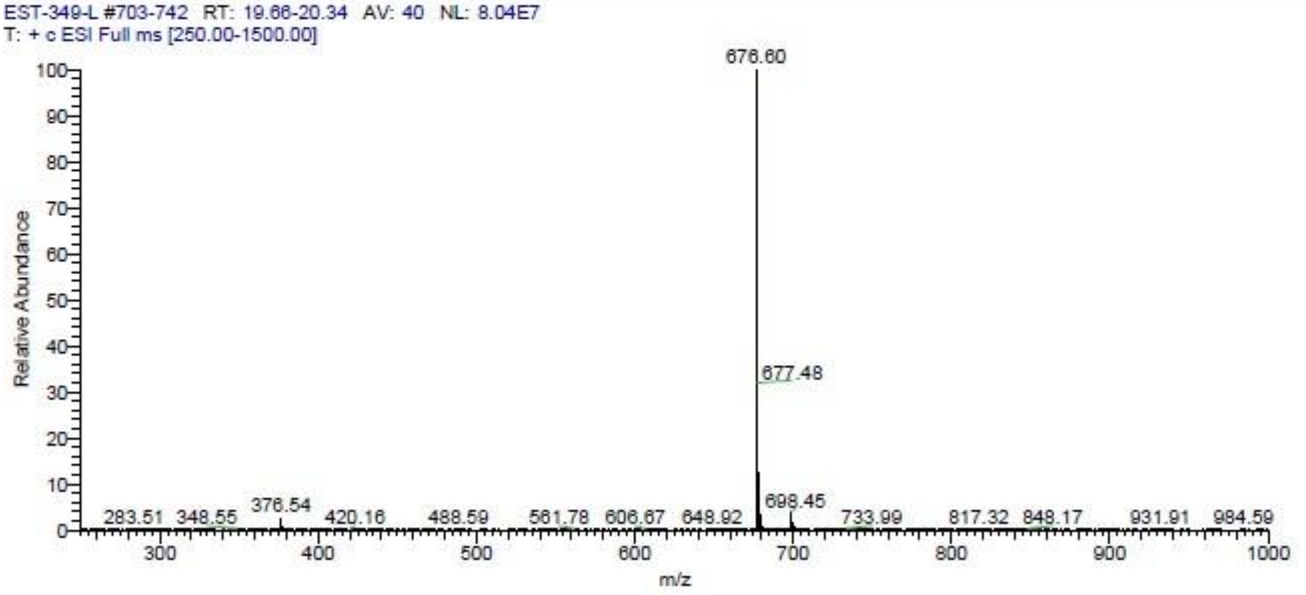
EST-349-L #703-742 RT: 19.66-20.34 AV: 40 NL: 8.04E7
T: + c ESI Full ms [250.00-1500.00]
Relative Abundance
m/z
676.60
677.48
698.45
283.51
348.55
376.54
420.16
488.59
561.78
606.67
648.92
733.99
817.32
848.17
931.91
984.59

**61** **10**.

**Figure S** . LC-MS analysis of compound

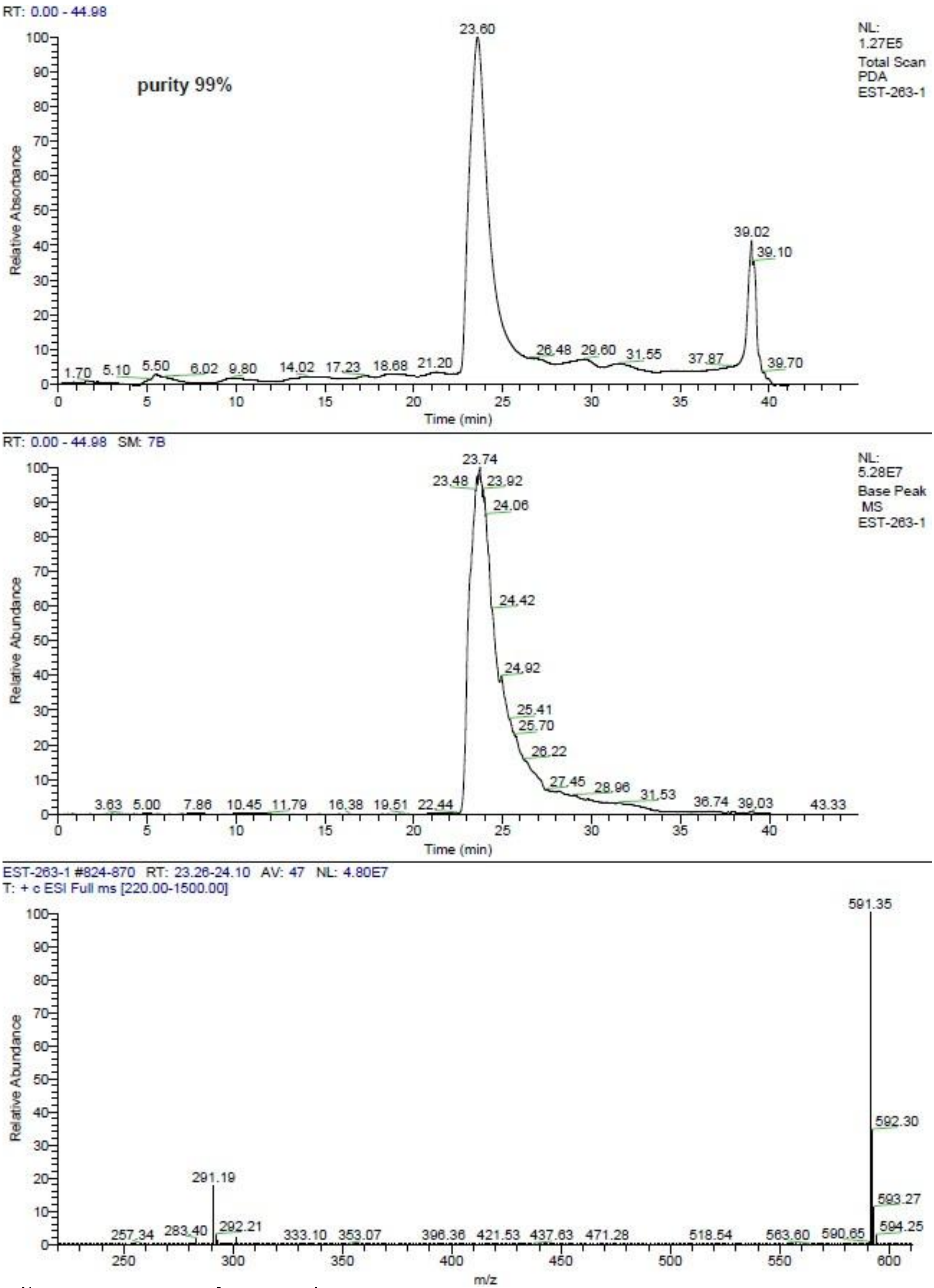

RT: 0.00 - 44.98
NL: 1.27E5
Total Scan
PDA
EST-263-1
purity 99%
Relative Absorbance
23.60
39.02
39.10
1.70 5.10 5.50 6.02 9.80 14.02 17.23 18.68 21.20 26.48 29.60 31.55 37.87 39.70
Time (min)
RT: 0.00 - 44.98 SM: 7B
NL: 5.28E7
Base Peak
MS
EST-263-1
Relative Abundance
23.74
23.48 23.92
24.06
24.42
24.92
25.41
25.70
26.22
27.45 28.96 31.53 36.74 39.03 43.33
3.63 5.00 7.86 10.45 11.79 16.38 19.51 22.44
Time (min)
EST-263-1 #824-870 RT: 23.26-24.10 AV: 47 NL: 4.80E7
T: + c ESI Full ms [220.00-1500.00]
Relative Abundance
591.35
592.30
593.27
594.25
291.19
257.34 283.40 292.21 333.10 353.07 396.36 421.53 437.63 471.28 518.54 563.60 590.65
m/z

[illegible]

**62** **11**.

**Figure S** . LC-MS analysis of compound

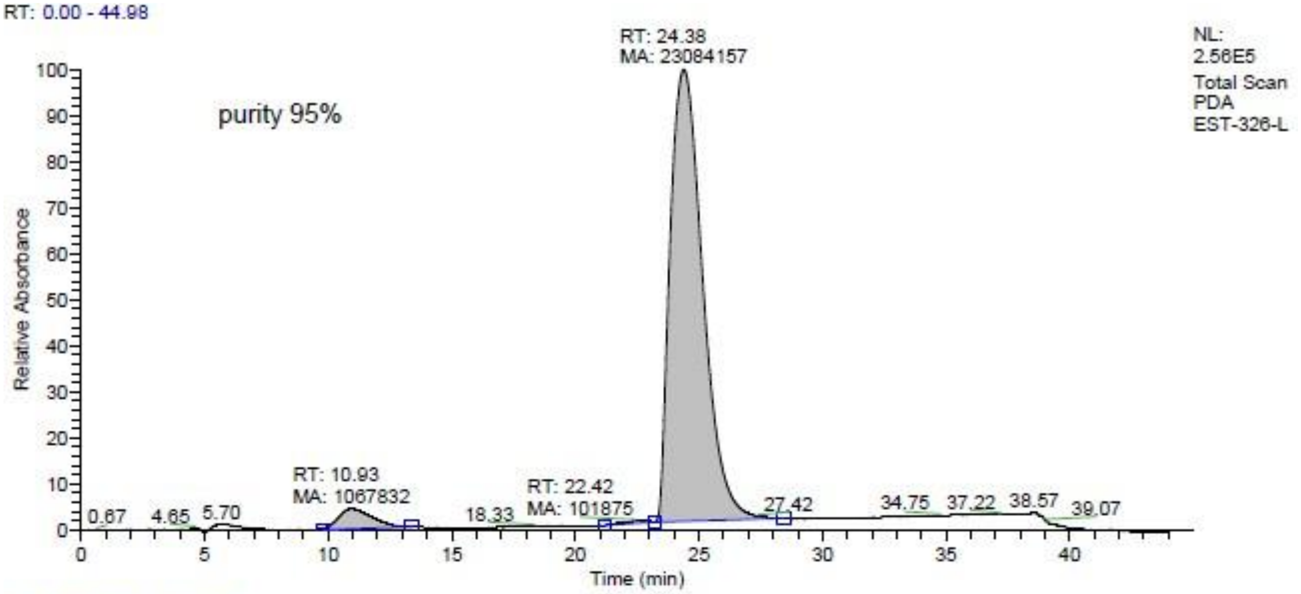
RT: 0.00 - 44.98
RT: 24.38
MA: 23084157
NL:
2.56E5
Total Scan
PDA
EST-326-L
purity 95%
Relative Absorbance
RT: 10.93
MA: 1067832
RT: 22.42
MA: 101875
0.67 4.65 5.70 18.33 27.42 34.75 37.22 38.57 39.07
Time (min)

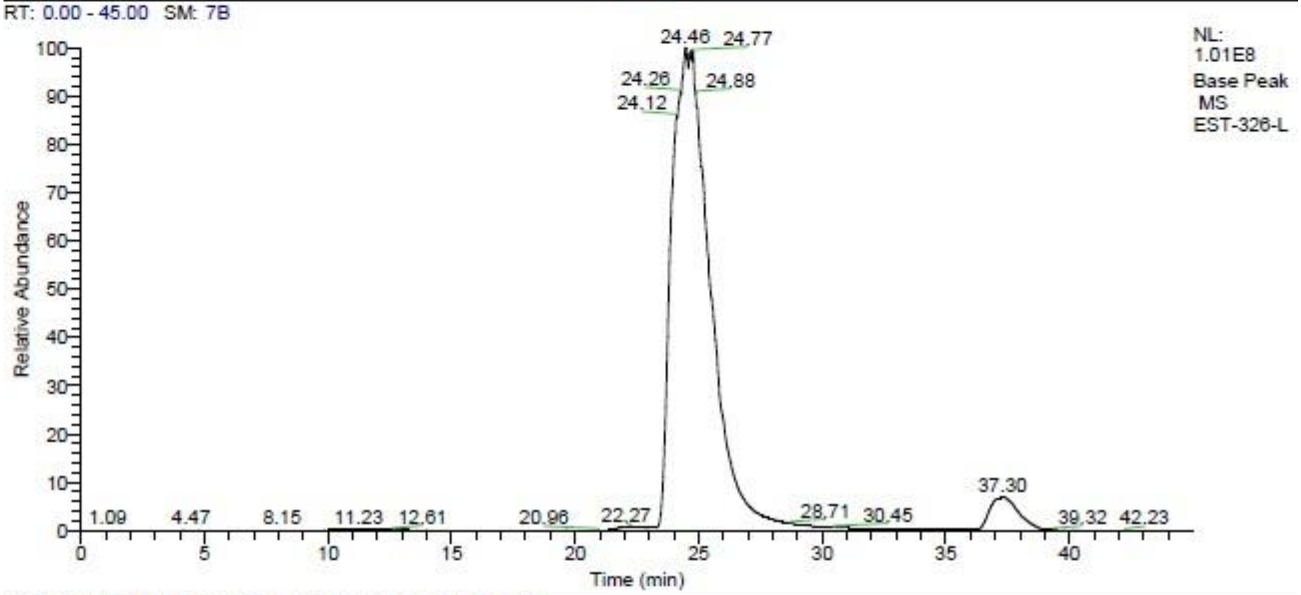
RT: 0.00 - 45.00 SM: 7B
24.46 24.77
24.26 24.88
24.12
NL:
1.01E8
Base Peak
MS
EST-326-L
Relative Abundance
1.09 4.47 8.15 11.23 12.61 20.96 22.27 28.71 30.45 37.30 39.32 42.23
Time (min)

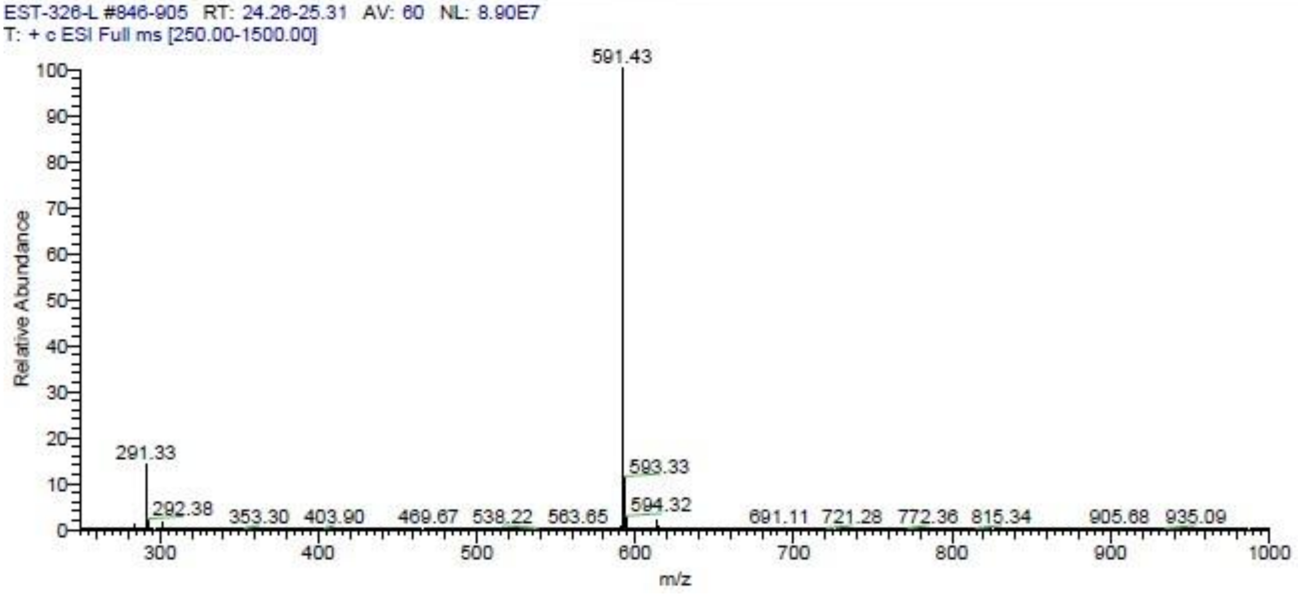
EST-326-L #846-905 RT: 24.26-25.31 AV: 60 NL: 8.90E7
T: + c ESI Full ms [250.00-1500.00]
591.43
291.33
593.33
292.38 353.30 403.90 469.67 538.22 563.65
594.32
691.11 721.28 772.36 815.34 905.68 935.09
Relative Abundance
m/z

**63** **12**.

**Figure S** . LC-MS analysis of compound

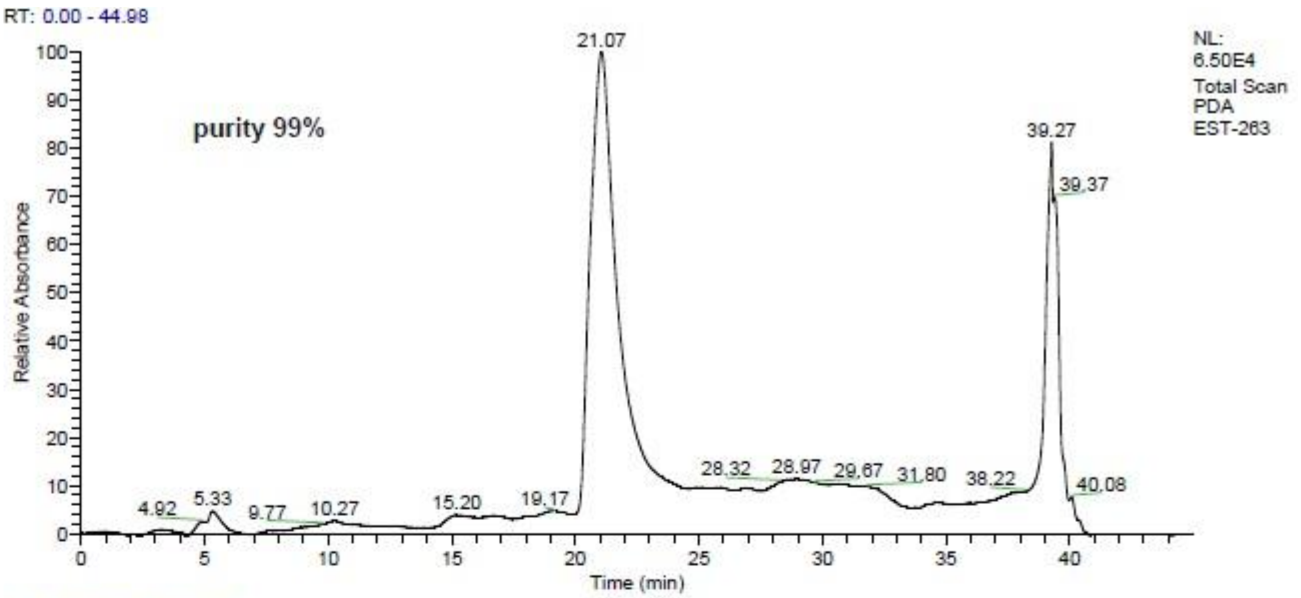
RT: 0.00 - 44.98
purity 99%
NL: 6.50E4 Total Scan PDA EST-263
21.07
39.27
39.37
4.92 5.33 9.77 10.27 15.20 19.17 28.32 28.97 29.67 31.80 38.22 40.08
Relative Absorbance
Time (min)

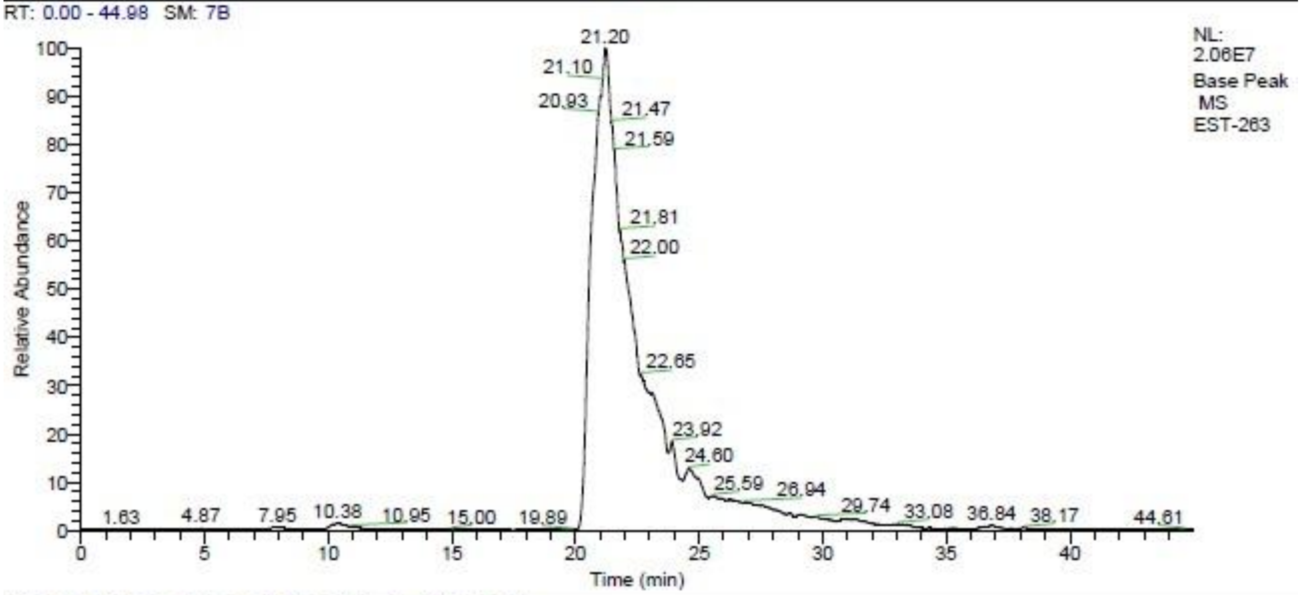
RT: 0.00 - 44.98 SM: 7B
NL: 2.06E7 Base Peak MS EST-263
21.20
21.10
20.93 21.47
21.59
21.81
22.00
22.65
23.92
24.60
25.59 26.94
29.74
1.63 4.87 7.95 10.38 10.95 15.00 19.89 33.08 36.84 38.17 44.61
Relative Abundance
Time (min)

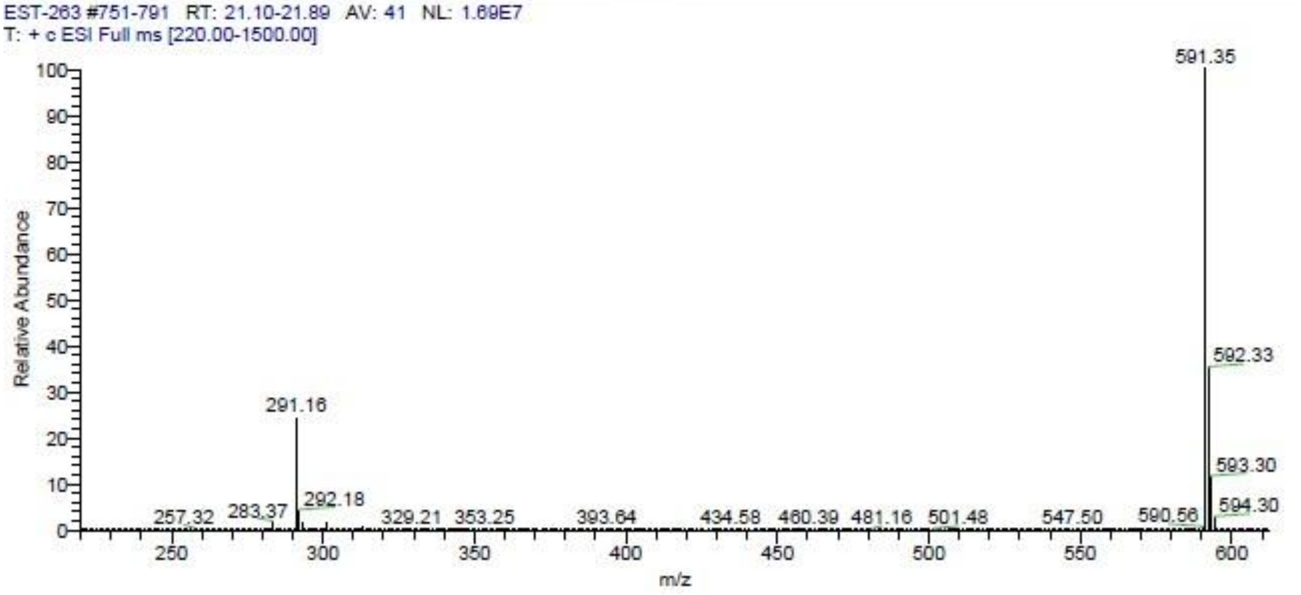
EST-263 #751-791 RT: 21.10-21.89 AV: 41 NL: 1.69E7
T: + c ESI Full ms [220.00-1500.00]
591.35
592.33
291.16
593.30
292.18
594.30
257.32 283.37 329.21 353.25 393.64 434.58 460.39 481.16 501.48 547.50 590.56
Relative Abundance
m/z

**64** **13**.

**Figure S** . LC-MS analysis of compound

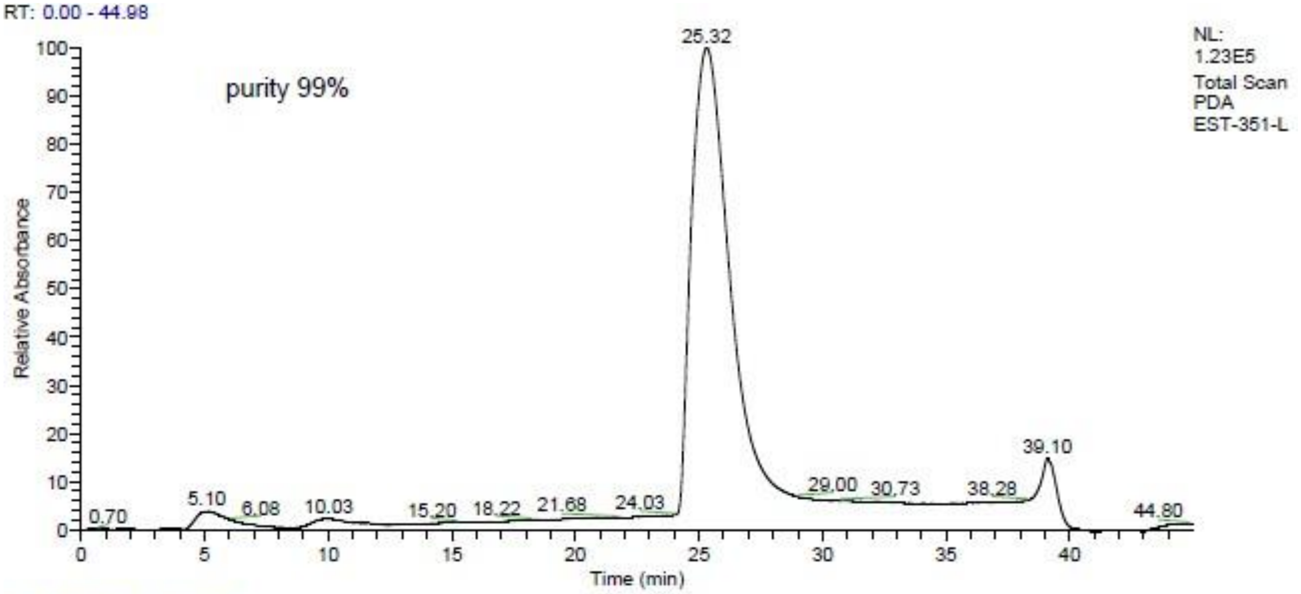
RT: 0.00 - 44.98
purity 99%
NL:
1.23E5
Total Scan
PDA
EST-351-L
Relative Absorbance
Time (min)
25.32
39.10
5.10
6.08
10.03
0.70
15.20
18.22
21.68
24.03
29.00
30.73
38.28
44.80

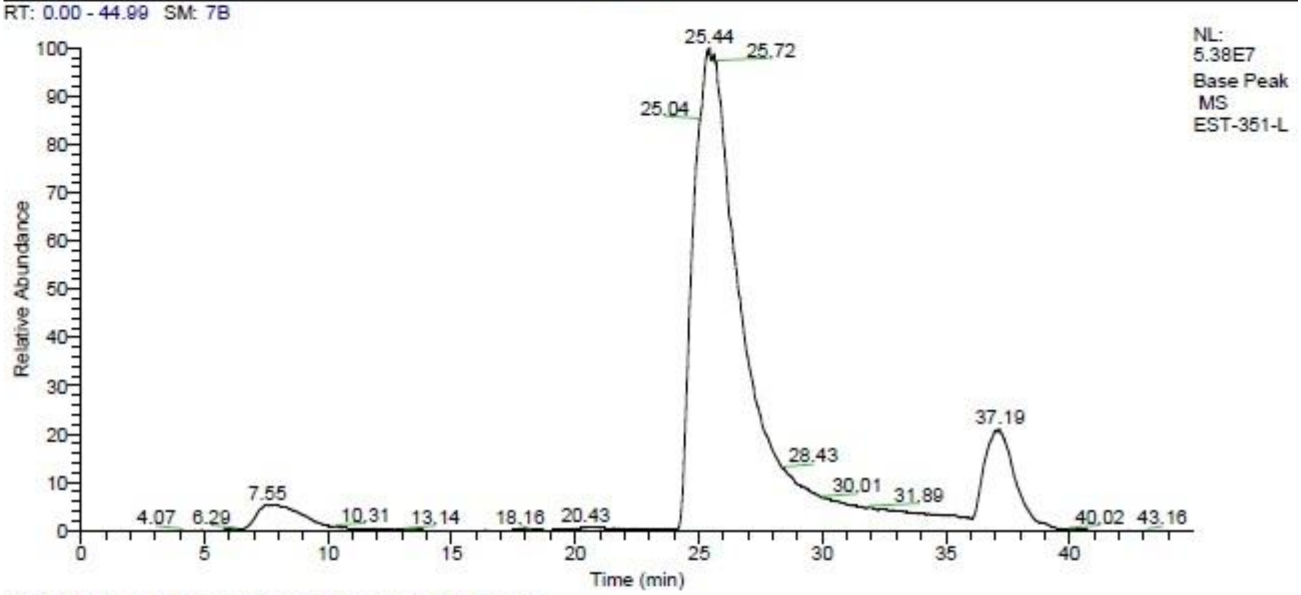
RT: 0.00 - 44.99 SM: 7B
NL:
5.38E7
Base Peak
MS
EST-351-L
Relative Abundance
Time (min)
25.44
25.72
25.04
37.19
28.43
30.01
31.89
7.55
4.07
6.29
10.31
13.14
18.16
20.43
40.02
43.16

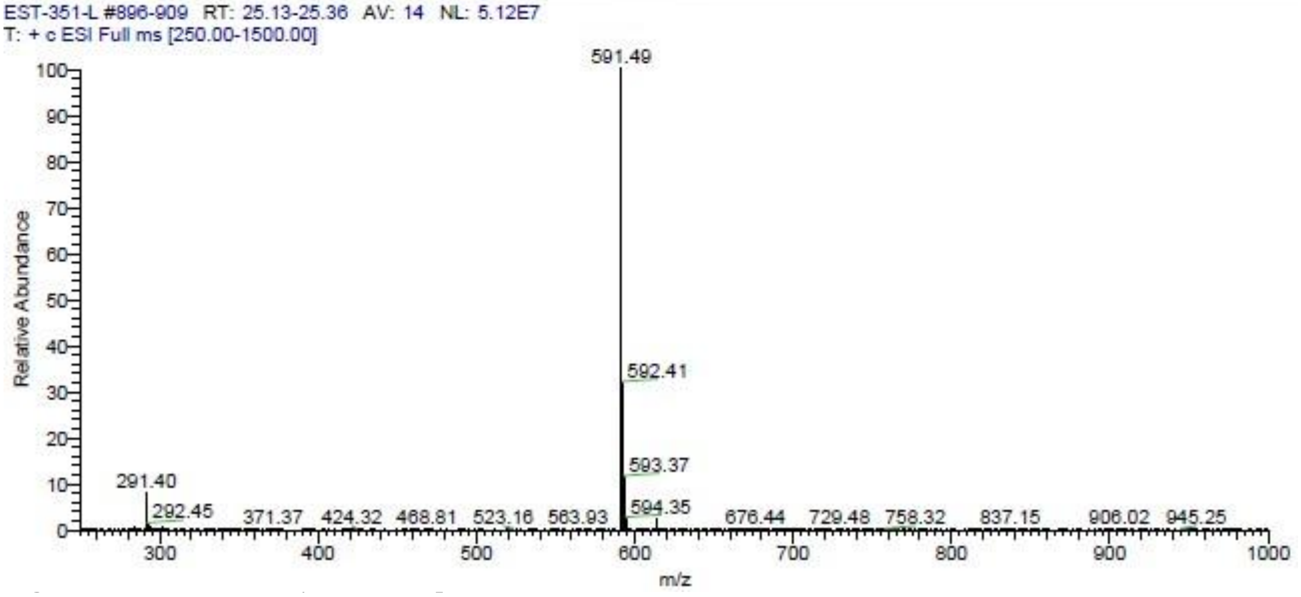
EST-351-L #896-909 RT: 25.13-25.36 AV: 14 NL: 5.12E7
T: + c ESI Full ms [250.00-1500.00]
Relative Abundance
m/z
591.49
592.41
593.37
594.35
291.40
292.45
371.37
424.32
468.81
523.16
563.93
676.44
729.48
758.32
837.15
906.02
945.25

**65** **14**.

**Figure S** . LC-MS analysis of compound

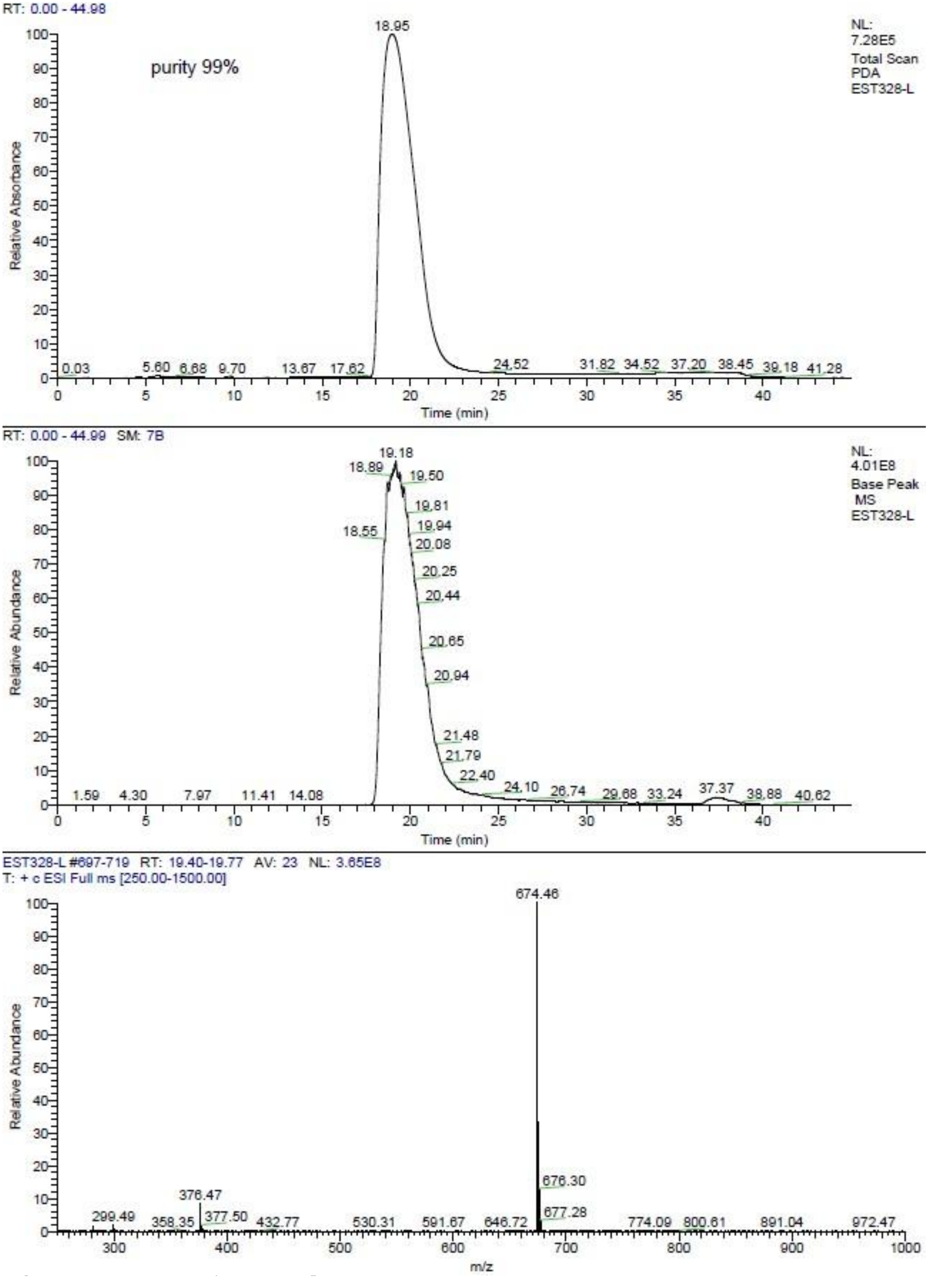
RT: 0.00 - 44.98
purity 99%
NL: 7.28E5 Total Scan PDA EST328-L
Relative Absorbance
Time (min)
18.95
0.03 5.60 6.68 9.70 13.67 17.62 24.52 31.82 34.52 37.20 38.45 39.18 41.28
RT: 0.00 - 44.99 SM: 7B
NL: 4.01E8 Base Peak MS EST328-L
Relative Abundance
Time (min)
19.18
18.89 19.50 19.81 19.94 18.55 20.08 20.25 20.44 20.65 20.94 21.48 21.79 22.40 24.10 26.74 29.68 33.24 37.37 38.88 40.62
1.59 4.30 7.97 11.41 14.08
EST328-L #697-719 RT: 19.40-19.77 AV: 23 NL: 3.65E8
T: + c ESI Full ms [250.00-1500.00]
Relative Abundance
m/z
674.46
676.30 677.28
376.47
299.49 358.35 377.50 432.77 530.31 591.67 646.72 774.09 800.61 891.04 972.47

**66** **15**.

**Figure S** . LC-MS analysis of compound

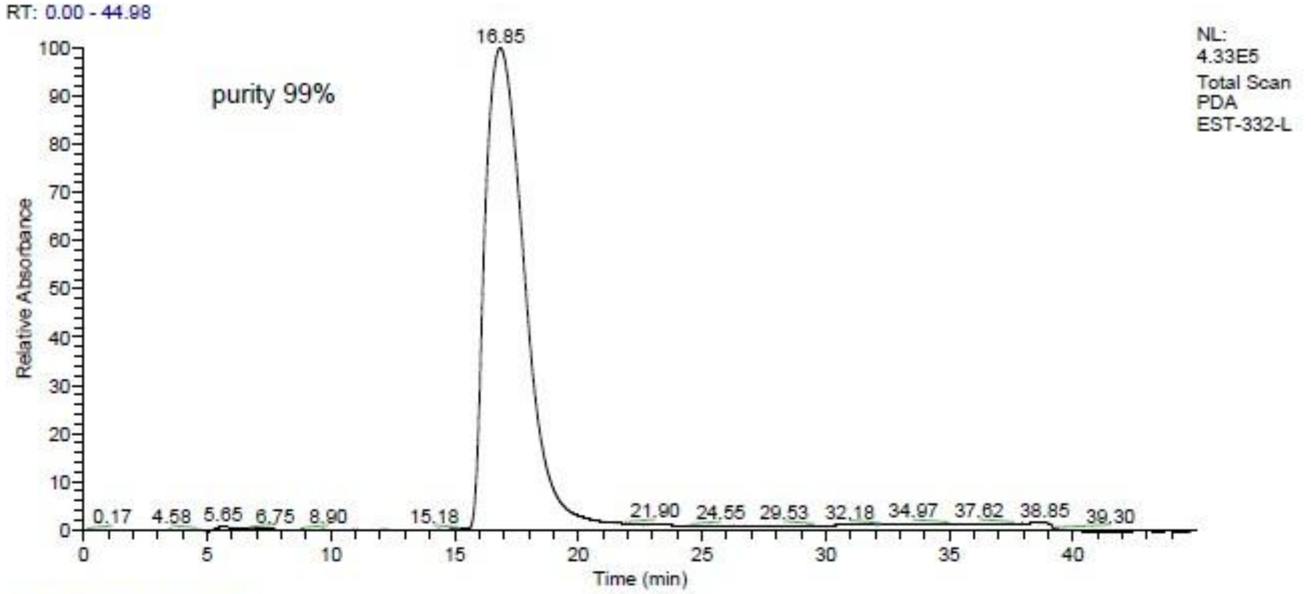
RT: 0.00 - 44.98
purity 99%
NL: 4.33E5
Total Scan PDA EST-332-L
16.85
0.17 4.58 5.65 6.75 8.90 15.18 21.90 24.55 29.53 32.18 34.97 37.62 38.85 39.30
Relative Absorbance
Time (min)

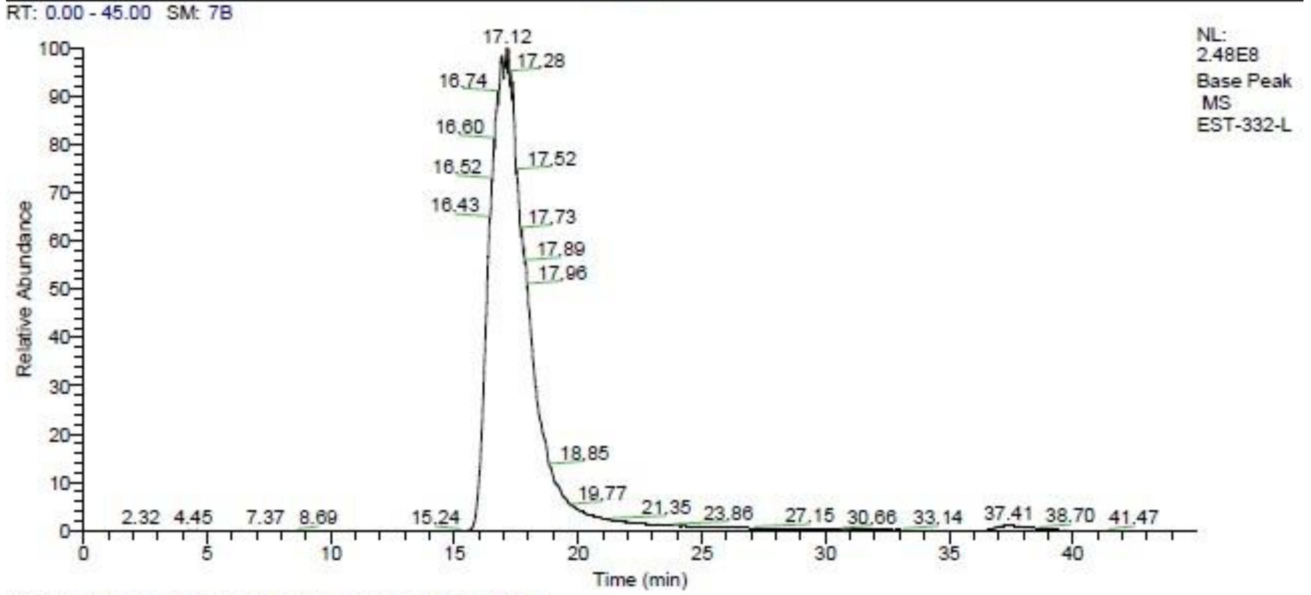
RT: 0.00 - 45.00 SM: 7B
NL: 2.48E8
Base Peak MS EST-332-L
17.12
17.28
16.74
16.60
16.52
16.43
17.52
17.73
17.89
17.96
18.85
19.77
2.32 4.45 7.37 8.69 15.24 21.35 23.86 27.15 30.66 33.14 37.41 38.70 41.47
Relative Abundance
Time (min)

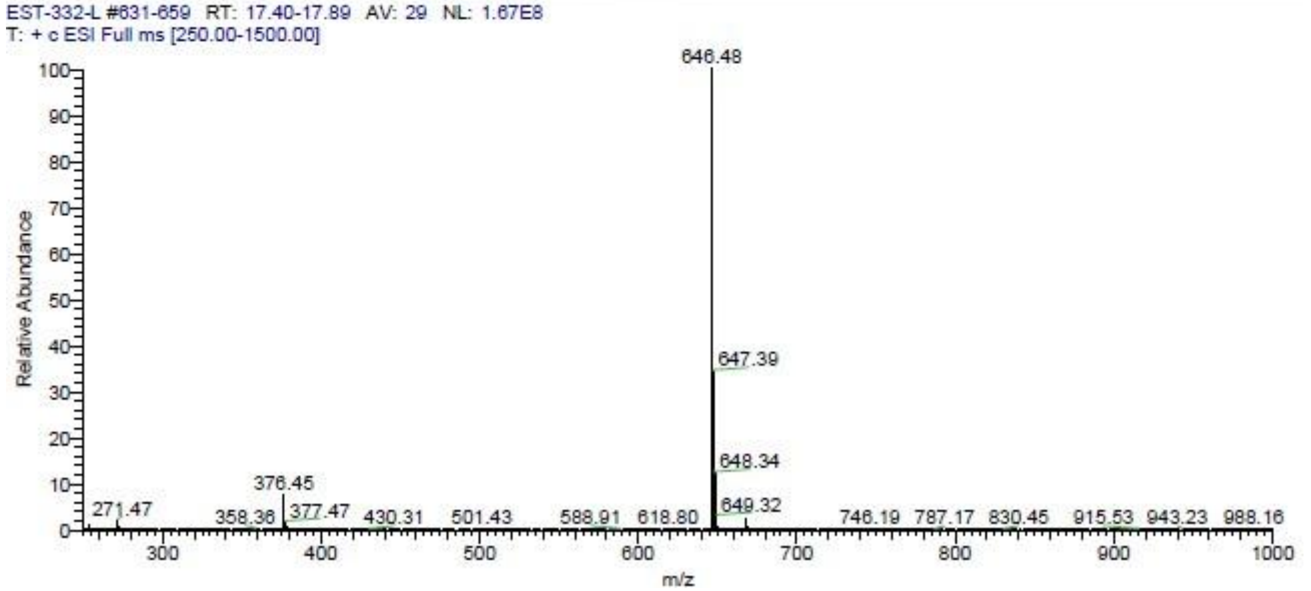
EST-332-L #631-659 RT: 17.40-17.89 AV: 29 NL: 1.67E8
T: + c ESI Full ms [250.00-1500.00]
646.48
647.39
648.34
649.32
376.45
271.47 358.36 377.47 430.31 501.43 588.91 618.80 746.19 787.17 830.45 915.53 943.23 988.16
Relative Abundance
m/z

**Figure S67** **16**.

**Figure S** . LC-MS analysis of compound

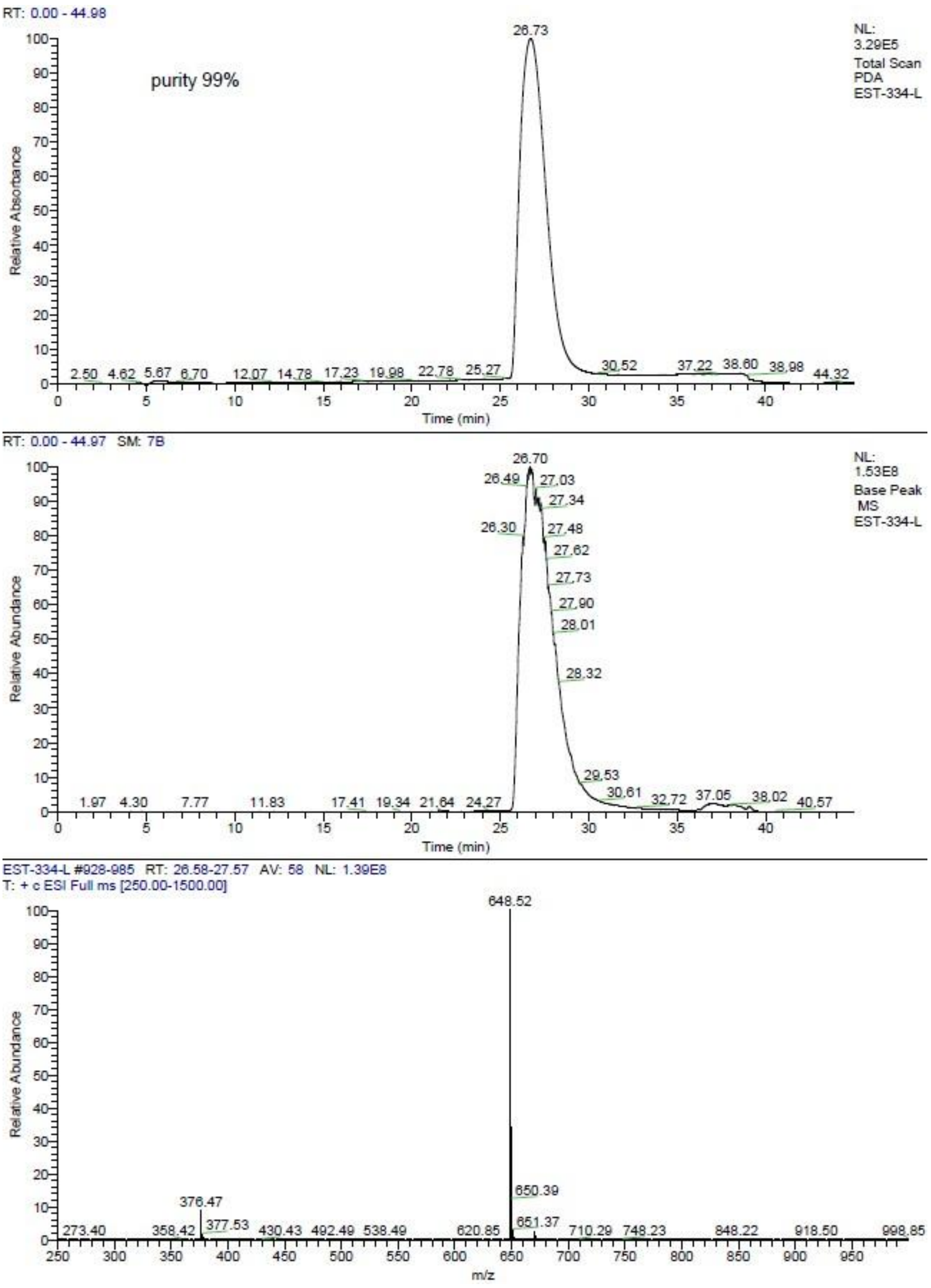
RT: 0.00 - 44.98
purity 99%
NL:
3.29E5
Total Scan
PDA
EST-334-L
26.73
Relative Absorbance
Time (min)
RT: 0.00 - 44.97 SM: 7B
NL:
1.53E8
Base Peak
MS
EST-334-L
26.70
Relative Abundance
Time (min)
EST-334-L #928-985 RT: 26.58-27.57 AV: 58 NL: 1.39E8
T: + c ESI Full ms [250.00-1500.00]
648.52
650.39
376.47
Relative Abundance
m/z

**68** **17**.

**Figure S** . LC-MS analysis of compound

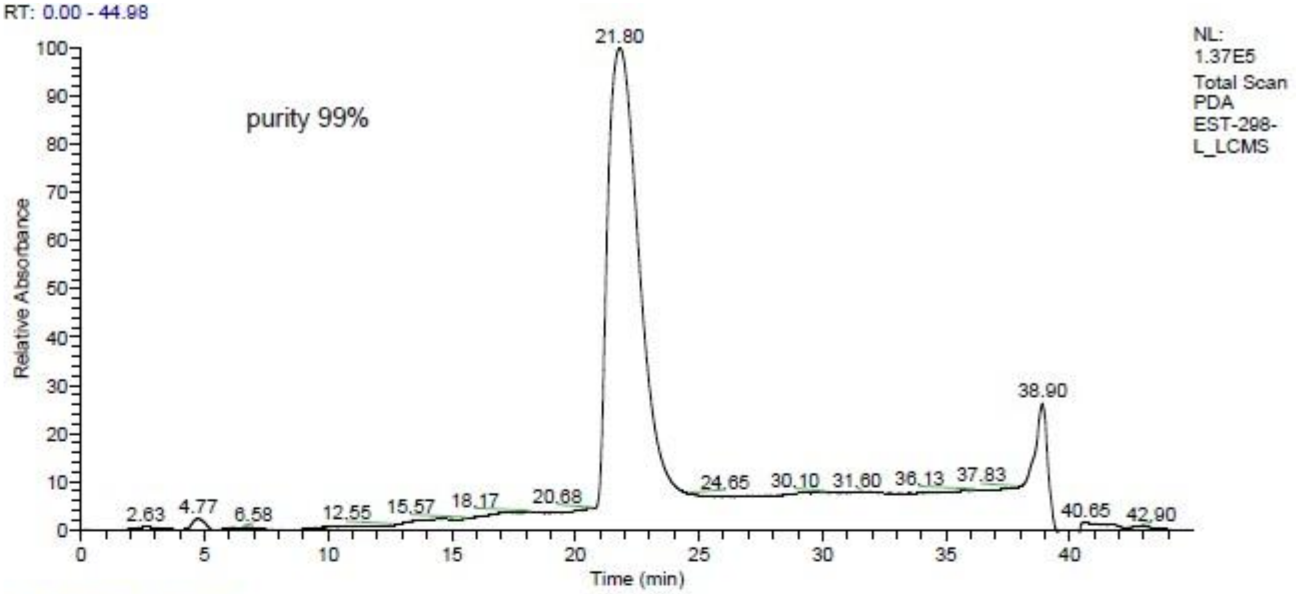
RT: 0.00 - 44.98
purity 99%
NL: 1.37E5 Total Scan PDA EST-298-L_LCMS
Relative Absorbance
Time (min)
21.80
38.90
2.63 4.77 6.58 12.55 15.57 18.17 20.68 24.65 30.10 31.60 36.13 37.83 40.65 42.90

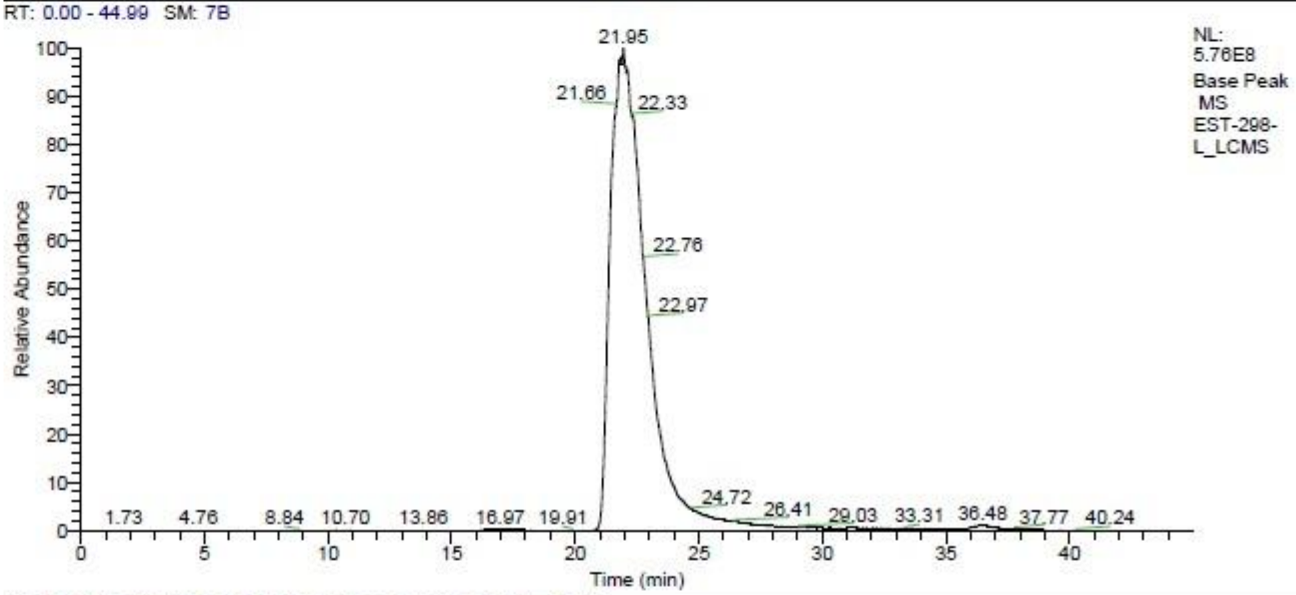
RT: 0.00 - 44.99 SM: 7B
NL: 5.76E8 Base Peak MS EST-298-L_LCMS
Relative Abundance
Time (min)
21.95
21.66 22.33 22.76 22.97
1.73 4.76 8.84 10.70 13.86 16.97 19.91 24.72 26.41 29.03 33.31 36.48 37.77 40.24

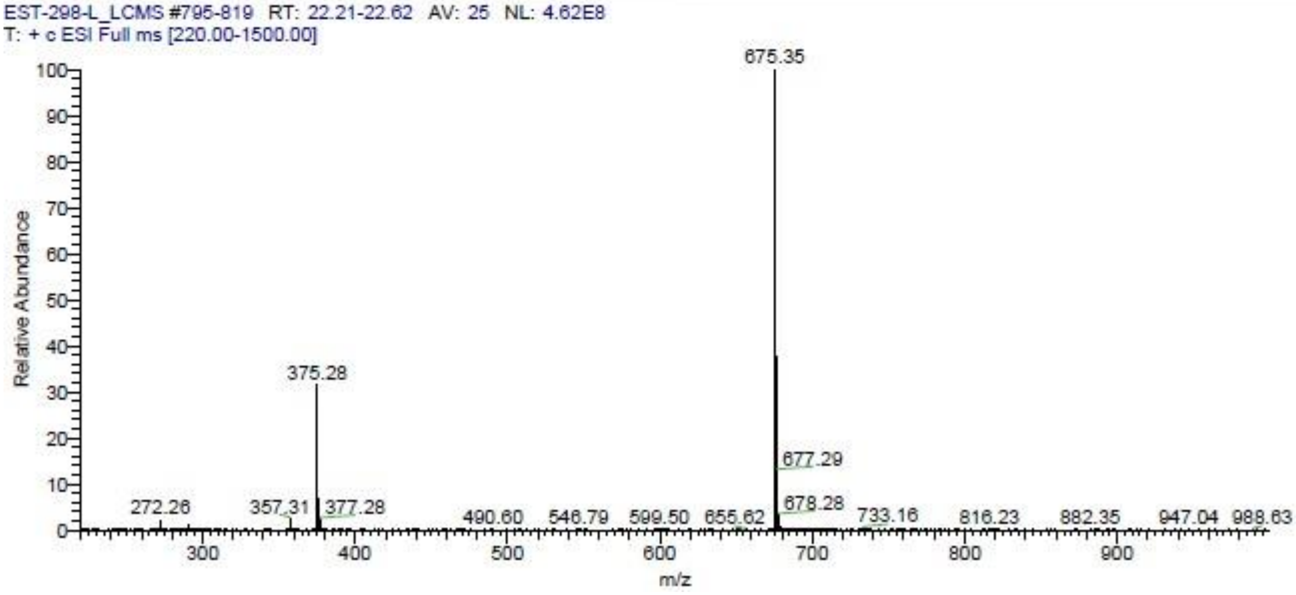
EST-298-L_LCMS #795-819 RT: 22.21-22.62 AV: 25 NL: 4.62E8
T: + c ESI Full ms [220.00-1500.00]
Relative Abundance
m/z
675.35
375.28
677.29
678.28
272.26 357.31 377.28 490.60 546.79 599.50 655.62 733.16 816.23 882.35 947.04 988.63

**69** **18**.

**Figure S** . LC-MS analysis of compound

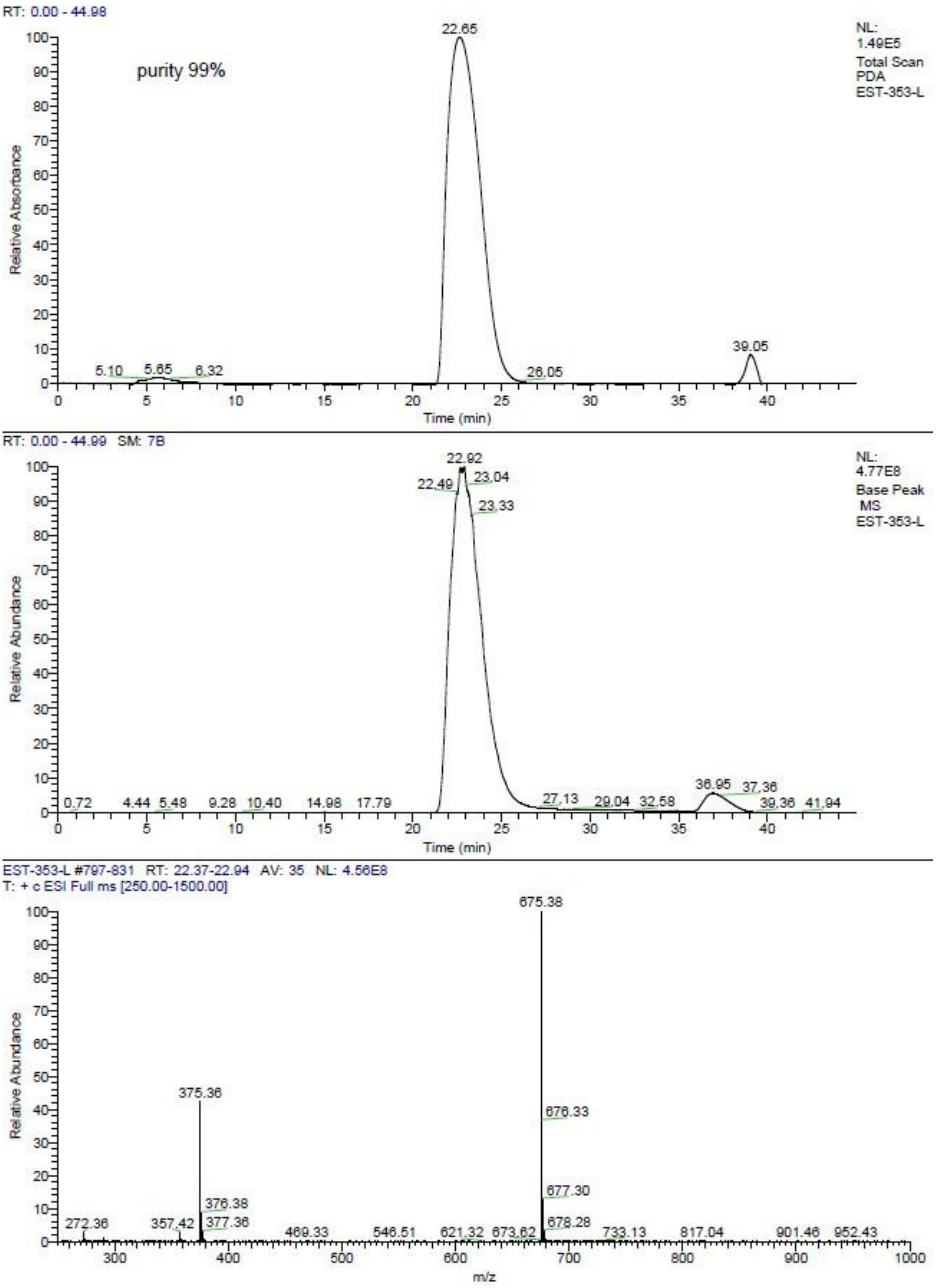
RT: 0.00 - 44.98
purity 99%
NL:
1.49E5
Total Scan
PDA
EST-353-L
Relative Absorbance
Time (min)
RT: 0.00 - 44.99 SM: 7B
NL:
4.77E8
Base Peak
MS
EST-353-L
Relative Abundance
Time (min)
EST-353-L #797-831 RT: 22.37-22.94 AV: 35 NL: 4.56E8
T: + c ESI Full ms [250.00-1500.00]
Relative Abundance
m/z

**70** **19**.

**Figure S** . LC-MS analysis of compound

| | |
|---|---|
| Box | : EST-398 |
| Position | : EST-398 |
| Method Filename | : 4_PFPP-120 - grad_35.lcm |
| Vial # | : 1-12 |
| Injection Volume | : 0.6 uL |
| Date Acquired | : 3/5/2026 4:13:23 PM |

Comment : Gradient, A: H2O, MeOH, FA (950:50:1); B: ACN; 35-100-35, 0.5 ml/min
Column: Shim-pack Scepter PFPP-120, 1.9um, 2.1x75 mm (Shimadzu)
ELSD:
Gain:10
Temperature: 50 °C

ELSD

| Ret. Time | Area | Area% |
|---|---|---|
| 3.250 | 74022 | 100.000 |
| | 74022 | 100.000 |

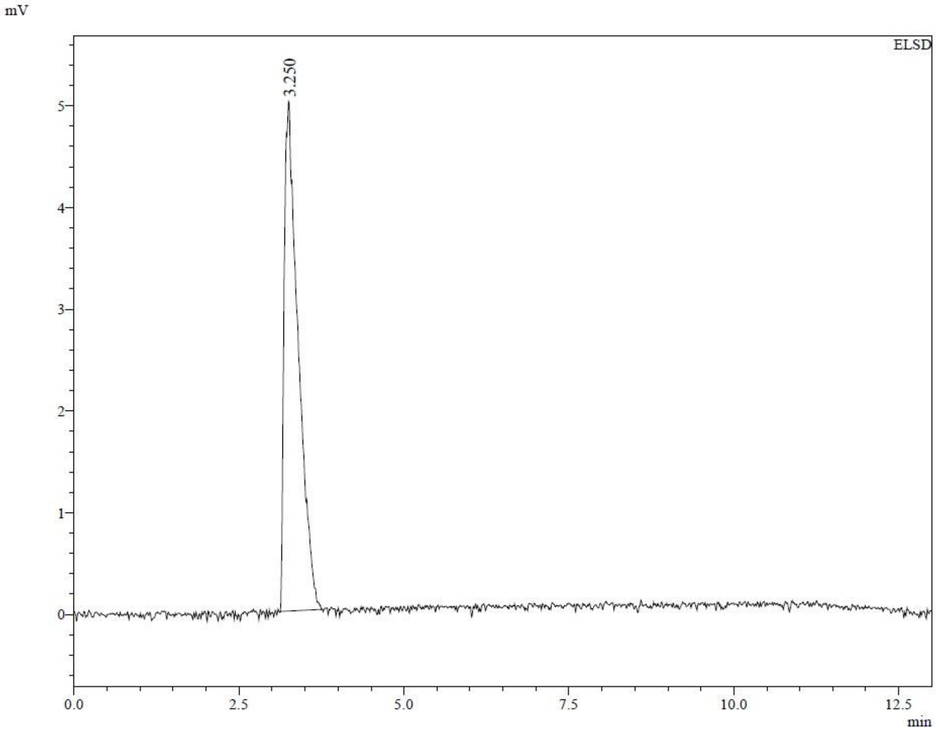

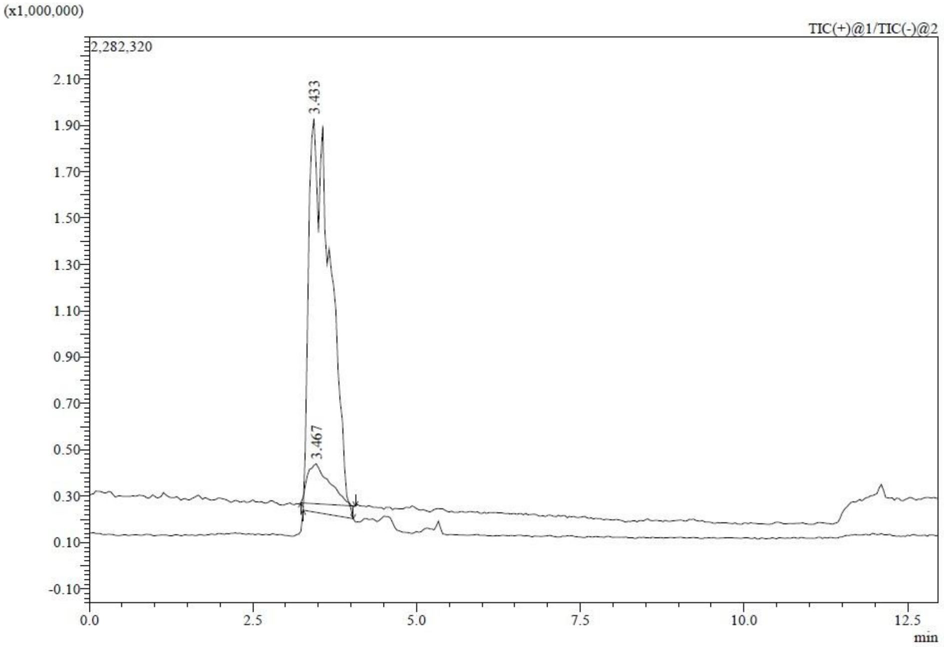

**Figure S71** **20**.

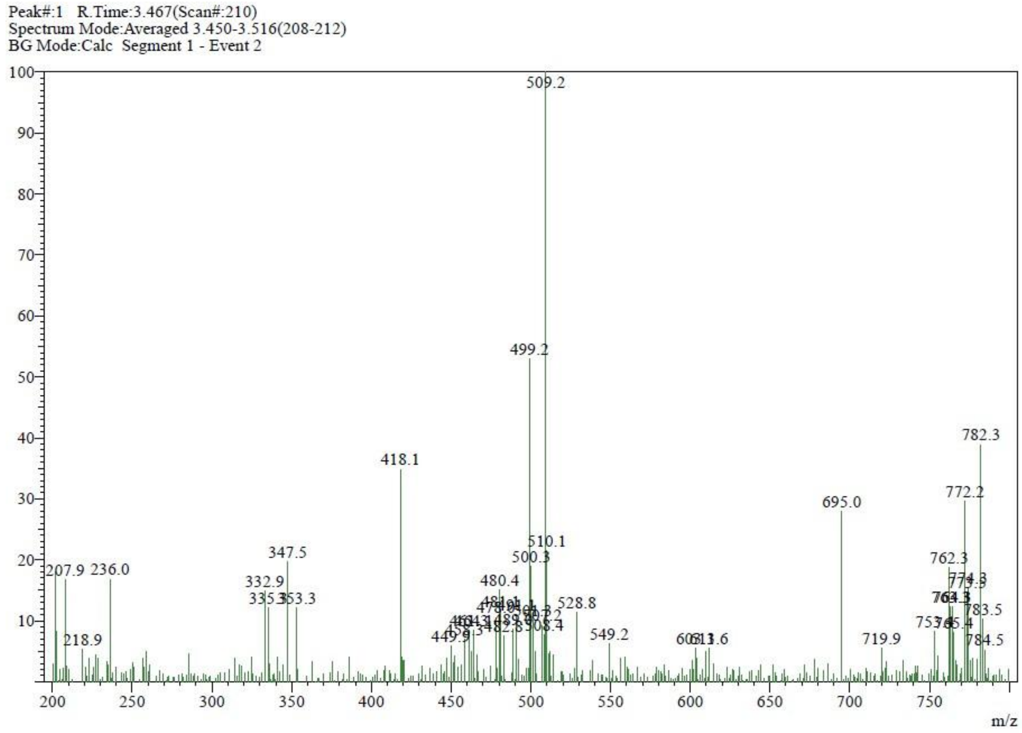

**Figure S71 (continued)**. LC-MS analysis of compound **20**.

. LC-MS analysis of compound

| | |
|---|---|
| Box | : EST-394 |
| Position | : EST-394 |
| Method Filename | : 4_PFPP-120 - grad_35.lcm |
| Vial # | : 1-11 |
| Injection Volume | : 0.6 uL |
| Date Acquired | : 3/5/2026 4:00:01 PM |

Comment : Gradient, A: H2O, MeOH, FA (950:50:1); B: ACN; 35-100-35, 0.5 ml/min
Column: Shim-pack Scepter PFPP-120, 1.9um, 2.1x75 mm (Shimadzu)
ELSD:
Gain:10
Temperature: 50 °C

ELSD

| Ret. Time | Area | Area% |
|---|---|---|
| 3.433 | 13111 | 100.000 |
| | 13111 | 100.000 |

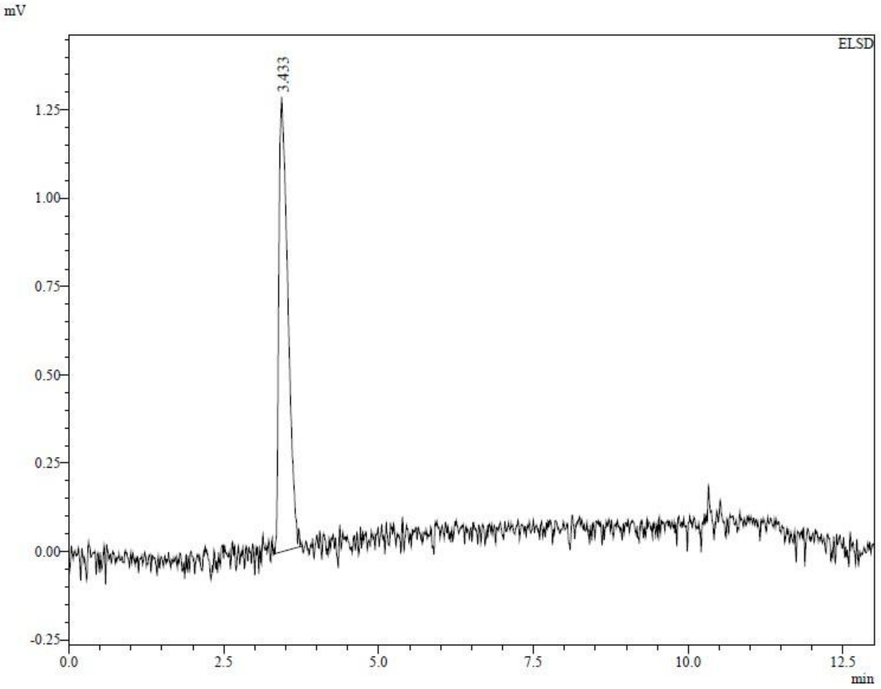

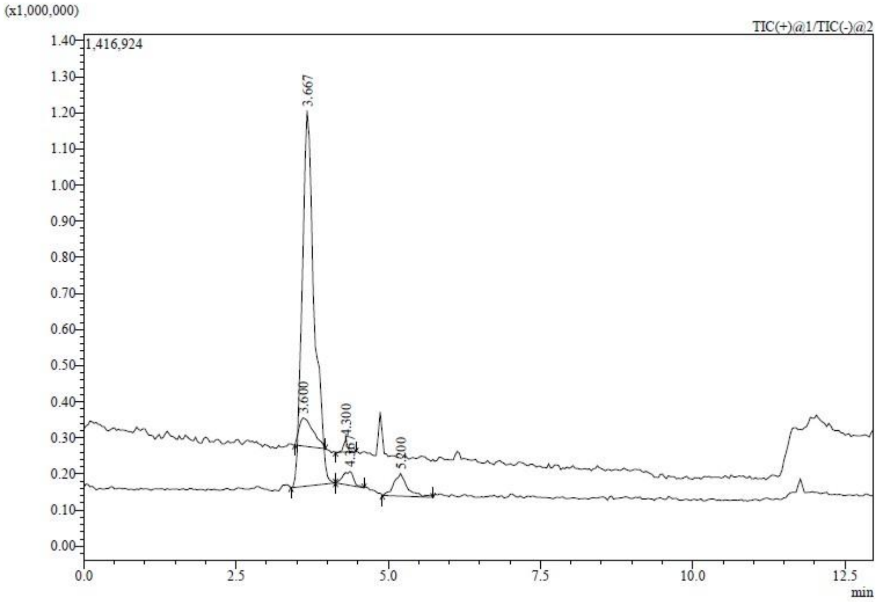

**Figure S72** **21**.

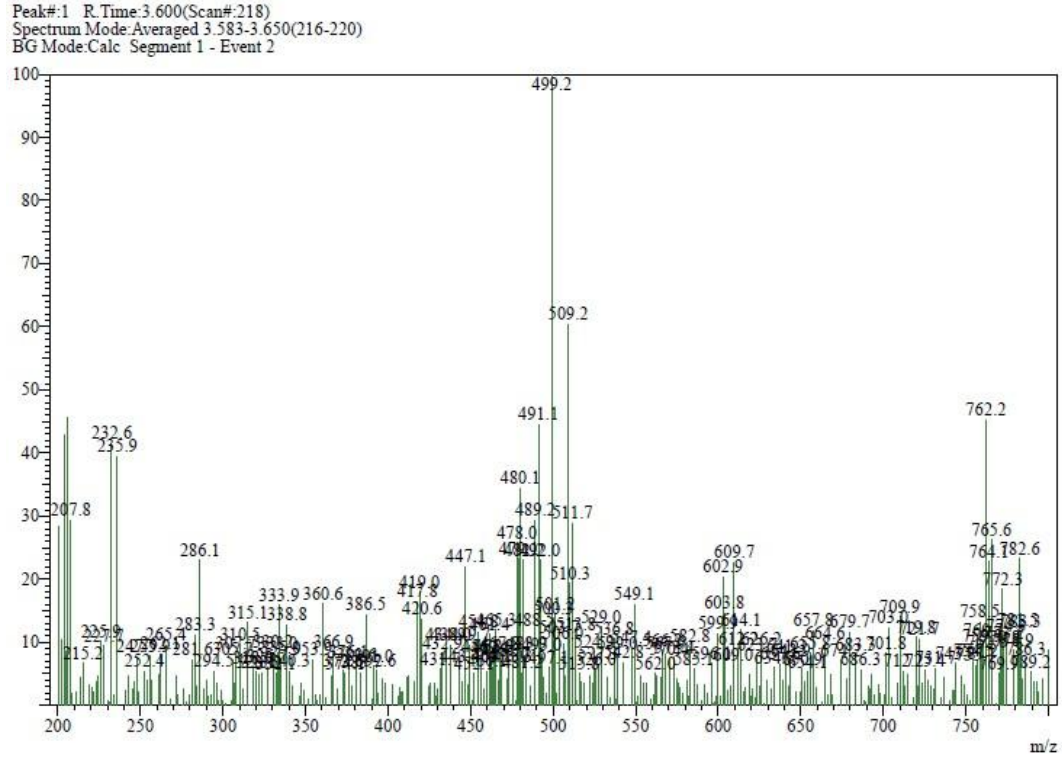

**Figure S72 (continued)**. LC-MS analysis of compound **21**.

. LC-MS analysis of compound

| | | |
|---|---|---|
| Box | : | EST-389 |
| Position | : | EST-389 |
| Method Filename | : | grad_C3_C4.lcm |
| Vial # | : | 1-20 |
| Injection Volume | : | 1 uL |
| Date Acquired | : | 1/15/2026 11:32:11 AM |

Comment : Gradient, A: H2O, MeOH, FA (950:50:1); B: ACN; 35-100-35, 0.6 ml/min
Column: Shim-pack Scepter C4-300, 1.9um, 2.1x100mm (Shimadzu)
ELSD:
Gain:10
Temperature: 50 °C

ELSD

| Ret. Time | Area | Area% |
|---|---|---|
| 4.983 | 95314 | 100.000 |
| | 95314 | 100.000 |

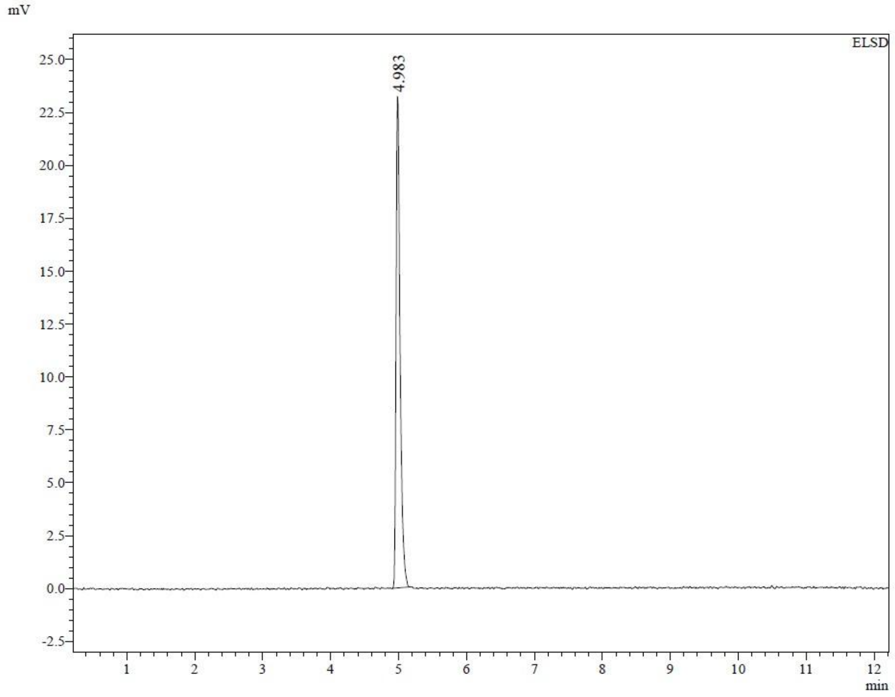

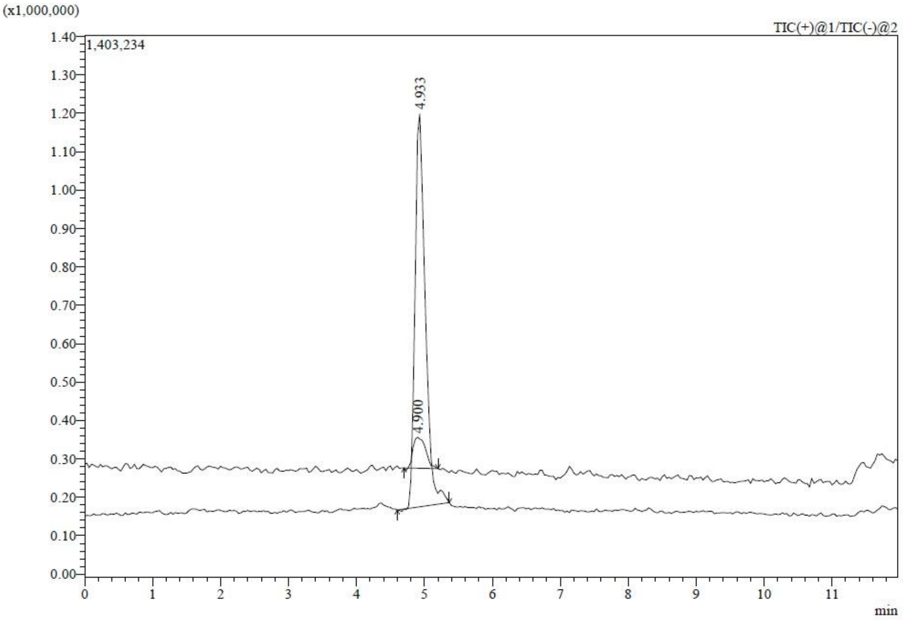

**Figure S73** **22**.

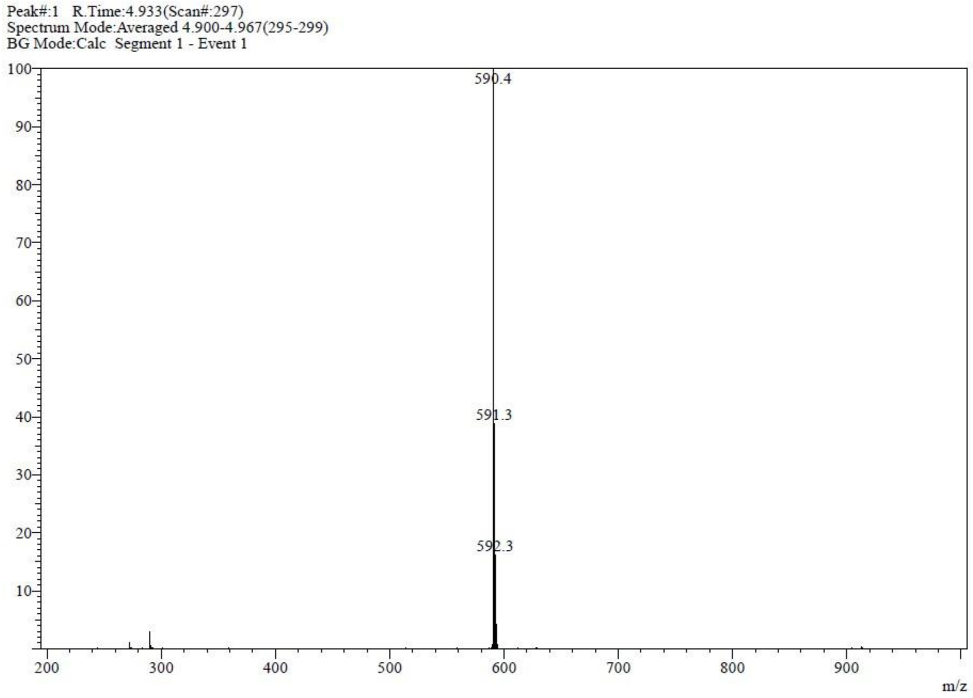

**Figure S73 (continued)**. LC-MS analysis of compound **22**.

. LC-MS analysis of compound
